# Hippurate hydrolases encoded by *Klebsiella pneumoniae* and *Klebsiella oxytoca* are functional

**DOI:** 10.64898/2026.07.27.739834

**Authors:** Devarsh Dave, Het Gajjar, Sarah Mawer, Anne L. McCartney, Jack C. Leo, David Negus, Lesley Hoyles

## Abstract

Human urine is non-sterile, harbouring its own microbiota (urobiota). *Klebsiella* species are commensals of the urobiota but can also cause urinary tract infections (UTIs). Gut-microbiota–host co-metabolites (MHCMs) such as hippurate are major components of urine. We previously showed UTI-associated strains of *Klebsiella pneumoniae* break down hippurate in human urine, though the enzymes responsible for this bioconversion are unknown. Here we sought to confirm that *K. pneumoniae* hydrolyses hippurate, and to determine whether the enzyme(s) responsible for this bioconversion are present and/or active in other *Klebsiella* species.

Ninhydrin assays confirmed that strains of *K. pneumoniae* (n=3) and, albeit to a lesser extent, *Klebsiella oxytoca* (n=4) isolated from human urine could produce glycine from hippurate. Comparative protein (KEGG, UniProt, phylogenetic) and structural (AlphaFold, ChimeraX) analyses were used to predict putative hippurate hydrolases (HHs) encoded by *Klebsiella* species. Their gene products (n=2 *K. pneumoniae*; n=3 *K. oxytoca*) were cloned, expressed and purified. The proteins belonged to three distinct groups: only group 1 and group 2 HHs hydrolysed hippurate under conditions used in this study. Both HHs co-occurred with high prevalence in 15/20 (75 %) *Klebsiella* species (2885/3012 genomes, 95.8 %).

Most *Klebsiella* species encode two distinct HHs that release glycine and benzoate from hippurate. The functional group 2 HH is predicted to facilitate delivery of amino acids such as glycine to *K. pneumoniae* cells in the nutrient-limited environment of urine, and its activity demonstrates that members of the urobiota and opportunistic pathogens can contribute to catabolism of MHCMs in human urine.

## Introduction

Hippurate (C_9_H_9_N_3_), also known as hippuric acid, *N*-benzoylglycine or 2-benzamidoacetic acid, is a microbiota–host co-metabolite (MHCM). The mammalian gut microbiota converts polyphenols present in ingested fruits, vegetables, tea, coffee, and diets containing phenylalanine, quinic acid and shikimic acid to benzoic acid ^1,2^. This benzoic acid is taken up from the gut and transported to the liver and kidneys, where it is modified via two different reactions within the mitochondrial matrix. The first reaction results in the production of benzoyl adenylate, the second reaction results in the substitution of the adenylate with coenzyme A (CoA), followed by glycine reacting with the benzoyl-CoA to form hippurate ^1,3^. This hippurate is excreted from the body in urine. Human urine is a rich source of hippurate (27.92–932.66 μmol/mmol creatinine) in healthy individuals ^3,4^, with high urine concentrations considered an indicator of metabolic health ^1,2^.

*Klebsiella* species are rod-shaped, Gram-negative bacteria found in a range of different environments, are human commensals and are often associated with opportunistic and nosocomial infections ^5–8^. The ESKAPE pathogen *Klebsiella pneumoniae* is the best known species of the genus *Klebsiella*, with carbapenem-resistant *K. pneumoniae* recently listed as a high-priority pathogen by the World Health Organization because of the threat it poses to public health ^9^. Other species that belong to the *K. pneumoniae* species complex (e.g. *Klebsiella variicola*, *Klebsiella quasipneumoniae*) are of increasing interest because of their contribution to the global burden of antimicrobial resistance and clinical infections ^7^. *K. pneumoniae* is a minor commensal of the urobiota in healthy women ^10^ but also causes urinary tract infections (UTIs) ^7,11^. Based on our recent metabolomics work, all tested *K. pneumoniae* isolates (*n*=4) recovered from catheter-associated UTIs were able to break down hippurate present in human urine ^12^. Beyond limited work done on *K. aerogenes* (positive for hippurate hydrolysis) ^13^ and *K. pneumoniae* (mixed results) ^14^ over 60 years ago, no data are available on hippurate hydrolysis by *Klebsiella* species, with the enzymes responsible for the bioconversion of hippurate to glycine and benzoate by *Klebsiella* species unstudied.

Microbial hippurate hydrolysis can be assessed using a ninhydrin assay, where a deep-purple colour (Ruhemann’s purple) is produced by glycine released via the activity of hippurate hydrolase (HH) (benzoylglycine amidohydrolase; EC 3.5.1.32; <u>KEGG orthology</u> <u>K01451</u>; MEROPS MH clan of peptidases, family M20) ^13,15^. This bioconversion has been used to differentiate streptococci: group B strains are positive in the ninhydrin assay, while groups A, C, D and F are negative ^16^. The ninhydrin assay has also been used to differentiate *Campylobacter jejuni* (positive for hippurate hydrolysis) from other *Campylobacter* species ^15,17^. The well-characterized cytoplasmic HH of *C. jejuni* is a zinc-containing homotetramer (193□±□11□kDa; monomer 42.4□±□0.8□kDa) that functions as a carboxypeptidase with optimal activity at pH 7.5 and 50 °C, and a broad substrate range^18,19^.

Our research focuses on understanding the functionality of *Klebsiella* species, alongside determining how bacteria contribute to MHCM bioconversions ^12,20–22^. The aims of the current study were to use the ninhydrin assay to confirm *Klebsiella* strains isolated from urine could hydrolyse hippurate, and to identify the genes responsible for hippurate hydrolysis. While much is known about how MHCMs are produced and excreted in urine, how bacteria found in urine act upon these metabolites has been largely ignored. These overlooked bioconversions potentially have implications for how members of the urobiota and opportunistic pathogens can persist in the urinary tract and bladder, and contribute to its microenvironment ^23^.

## Methods

### Strains used in this study

Details of the strains used in this study can be found in **Table 1**. The study of anonymised clinical isolates (Ko prefix) provided by the Nottingham University Hospitals NHS Trust (NUH) Pathogen Bank was approved by NUH Research and Innovation (19MI001). Whole-genome sequence data for the *K. pneumoniae* strains are available from BioProject PRJNA917129 ^5^; *K. oxytoca* data are available from BioProject PRJNA1439749. Genes present in genomes were predicted using Bakta v1.9.3 (database v5.1, full) ^24^.

**Table 1.** Strains used in this study.

| Strain ID | Species | Source | Reference |
| --- | --- | --- | --- |
| Ko6 | *Klebsiella oxytoca* (ST37) | Urine | 6 |
| Ko25 | *Klebsiella oxytoca* (ST612) | Clean-catch urine | 6 |
| Ko26 | *Klebsiella oxytoca* (ST176) | Urine | 6 |
| Ko55 | *Klebsiella oxytoca* (ST36) | Mid-stream urine | 6 |
| PS_Kpn11 | *Klebsiella pneumoniae* (K107:O2v2, ST258) | Urine | 5 |
| PS_Kpn29 | *Klebsiella pneumoniae* (K2:O1v1, ST14) | Urine | 5 |
| PS_Kpn36 | *Klebsiella pneumoniae* (K24:O2v1, ST11) | Urine | 5 |
| ATCC 14928 | *Streptococcus agalactiae* | – |  |
| NCTC 9994 | *Streptococcus pyogenes* | – |  |
| DH5α | *Escherichia coli* | – |  |
| BL21 (DE3) | *Escherichia coli* | – |  |

### Plate-based ninhydrin assay

Modified from the method of Hwang & Ederer ^16^. Overnight cultures were prepared by inoculating 5 mL brain–heart infusion (BHI) broth (Thermo Scientific) and aerobic incubation at 37 °C. Following incubation, cultures were centrifuged at 5000 ***g*** for 10 min. The supernatant was discarded. The pellet was washed and resuspended to a 3.0 McFarland standard using sterile distilled water.

A 100 mM sodium hippurate (Thermo Scientific) solution was prepared in sterile distilled water. For the ninhydrin assay, 100 µL of the McFarland-adjusted culture was added into individual wells of a 96-well plate (Corning Clear Polystyrene 96-Well Microplate; Fisher Scientific). Then, 100 µL of the sodium hippurate solution was added to each well. The plates were incubated under aerobic (21 % O_2_) and microaerobic (5 % O_2_, 10 % CO_2_, 85 % N_2_; Don Whitley DG250 workstation) conditions at 37□°C for 24 h. Following incubation, 50 µL ninhydrin reagent [3.5 % ninhydrin (Thermo Scientific) in 1:1 (v/v) mixture of acetone and butanol] were added to each well. The plates were re-incubated under their respective conditions at 37 °C for an additional 30 min. *Streptococcus agalactiae* ATCC 14928 was used as the positive control, and *Streptococcus pyogenes* NCTC 9994 and uninoculated sterile water were used as negative controls ^25^. Plates were read at *A*_560_ using a BioTek Cytation |^3^ imaging reader spectrophotometer to quantify the intensity of the purple colour produced. The assay was also performed with 10 µL of Trace Metal Mix A5 with Co (Merck) solution added to samples for both conditions (prepared by adding 10 µL of the trace metal stock solution to 1 mL sterile distilled water). Three biological replicates with three technical replicates were performed for each experiment.

### Identification and analyses of HHs encoded by *Klebsiella* species

A search for “hippurate hydrolase” in UniProt v2024_06 (accessed January 2025) gave one entry that had been manually annotated and reviewed: UniProt entry P45493, for *C. jejuni* ^18,19^. This was used as the reference protein sequence for comparative analyses. Filtering the search result for *“Klebsiella*”, 12 UniProt entries were identified: *K. quasipneumoniae* A0A1C3PY10, A0A1C3Q1U6 and A0AB34SYT7; *K. michiganensis* A0A7H5A841, A0A7H5ABG0 and A0A443V063; *K. grimontii* A0A285B0F0, A0A285B2R9 and A0A285B568; *K. oxytoca* A0A318FWX7, A0A318F7I2 and A0A318FI22. The protein sequences were used to create a multiple-sequence alignment (MSA) (Clustal Omega v1.2.2) and associated pairwise identity matrix (Geneious Prime v2025.2.1).

The tertiary structures of the 13 above-mentioned HHs were predicted using AlphaFold2 (ColabFold implementation) ^26^. The protein structures were visualized and analysed using ChimeraX v1.9 ^27^. The predicted protein structures of each of the 12 *Klebsiella* HHs were compared to the structure of *C. jejuni*’s confirmed HH using the MatchMaker command of ChimeraX. Root mean square deviation (RMSD) values were calculated to assess the similarity of the compared structures.

HH genes were identified in the Kyoto Encyclopedia of Genes and Genomes (KEGG) database (KEGG orthology K01451; accessed January 2025), and the sequences retrieved using KEGGREST v1.48.1 ^28^. These sequences were combined with those we identified in UniProt to create a BLASTP database, used to identify putative HHs encoded by our seven *Klebsiella* strains (**Table 1**).

All identified HHs were included in a MSA, created using Clustal Omega v1.2.2 in Geneious Prime v2025.2.1: KEGG (n=1684), UniProt *Klebsiella* and *C. jejuni* (n=13), and sequences predicted for our seven *Klebsiella* strains (n=18). The MSA was exported from Geneious in phylip format and used to create a phylogenetic tree: ModelFinder was first run in iqtree v3.1.2 ^29^ to determine the best-fit model to use for tree construction, then a bootstrapped (UFBoot ^30^, 1000 replicates) maximum-likelihood tree was created using iqtree and implementing the Q.PFAM+I+R10 model. The resulting tree file was visualized and annotated using iTOL v7.2 ^31^.

The above-mentioned phylogenetic analysis identified three groups of potential *Klebsiella* HHs. Only two of these were characterized in a wider *Klebsiella*-specific analysis (refer to Results for reasoning). The UniProt *Klebsiella* group 1 and group 2 sequences were used to create a BLASTP database against which the Bakta-predicted proteomes of 3012 *Klebsiella* isolates (high-quality whole-genome sequence data from ^5,11,20,22,32–34^, GenBank and this study) were searched. Results were filtered based on >70 % aa identity and 100 % query coverage (392 aa, group 1; 389 aa, group 2). For each isolate’s proteome, group 1 and group 2 hits were concatenated (separated by 10 Ns) and used to create an MSA (Clustal Omega v1.2.2 in Geneious Prime v2025.2.1), which was exported from Geneious in phylip format and run through iqtree as described above to find the best-fit model. A bootstrapped (UFBoot ^30^, 1000 replicates) maximum-likelihood tree was created using iqtree and implementing the Q.BIRD+I+R3 model. The resulting tree file was visualized and annotated using iTOL v7.2 ^31^.

Species-and group-specific analyses were undertaken for the genes flanking those encoding the group 1 and group 2 HHs. The R package rtracklayer v1.70.1 ^35^ was used to extract contig and coordinate information for both the group 1 and group 2 proteins from the Bakta-annotated gff3 files (n=2885). Biostrings v2.78.0 ^36^ was used to import the corresponding fna files into R, and the information from the gff3 files was used to extract sequences 5000 bp up-and down-stream of the HH-encoding genes, where possible, with sequences reverse-complemented when the HH was encoded on the negative strand. For each *Klebsiella* species and HH group combination, a non-redundant set of sequences was exported from R, and used with mafft v7.526 ^37^ to create MSAs. The MSAs were imported into Geneious Prime. For each MSA, a representative annotated genome was selected and the MSA was mapped to the reference (Geneious Mapper). Gene annotations were transferred to the consensus sequence, and sequences within both protein groups were compared to one another. In an effort to standardize functional annotations across all species, <u>FoldSeek</u> ^38^ was used to check proteins annotated in the consensus sequences. The gene alignments were exported from Geneious Prime and annotated manually using Adobe Illustrator.

### Cloning of predicted HHs and transformation

Genomic DNA of the *Klebsiella* strains used in this study (**Table 1**) was extracted using the Puregene Yeast/Bact. Kit B (Qiagen) following the manufacturer’s instructions. DNA concentration was measured (NanoDrop 2000 v1.6.198) and stocks were diluted to 100 ng/µL using nuclease-free water and stored at -20 °C until required. The pET26b(+) (Novagen) plasmid sequence was imported into the NEBuilder programme v2.10.8 as a circular vector and linearized via PCR at the multiple cloning site using the pET26b_forward (5′-CACCACCACCACCACCACTGAGATCCGGCTGCTAACAA-3′) and reverse (5’-ATGTATATCTCCTTCTTAAAGTTAAACAAAATTATTTC-3’) primers. Coding sequences (without stop codons for the introduction of the C-terminal fusion His_6_ tags) of the three Ko6 and two PS_Kpn11 genes predicted to be responsible for hippurate hydrolysis were included as inserts. The programme generated compatible primer pairs for HiFi DNA Assembly (**Table 2**) to insert the genes into the pET26b(+) vector.

**Table 2.**
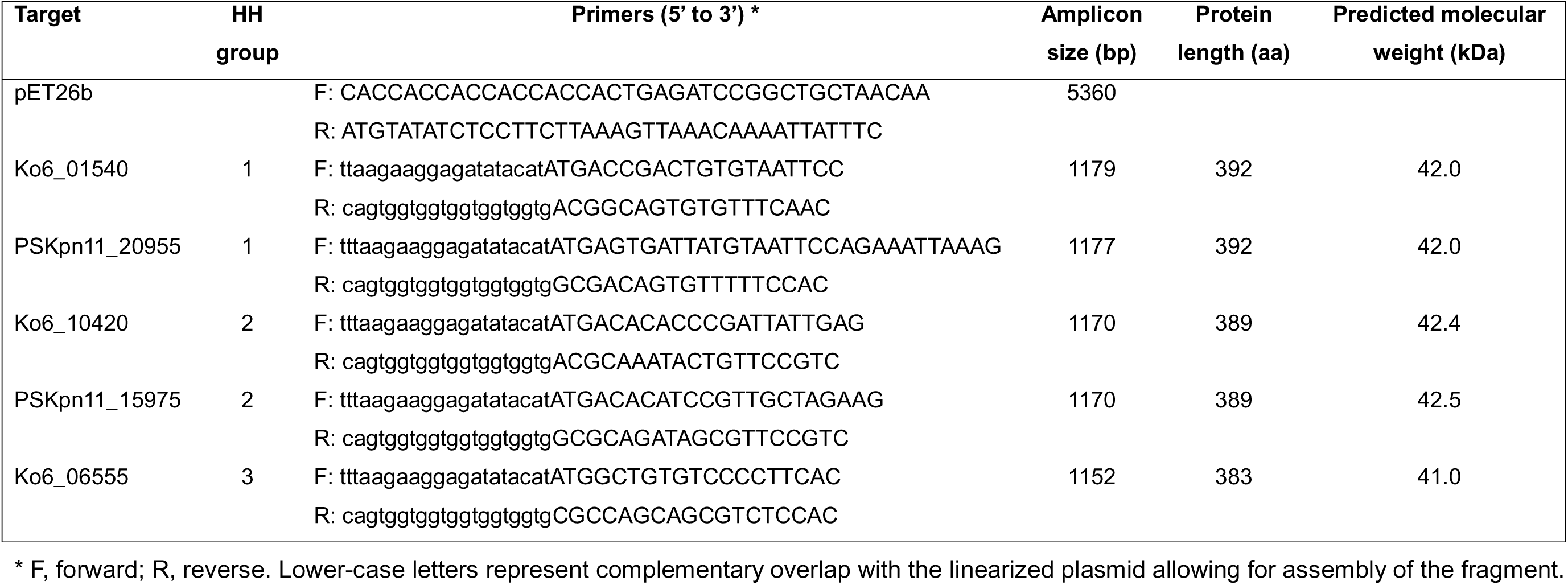
Primers designed for use in this study, with additional information for *Klebsiella* predicted HH gene products. * F, forward; R, reverse. Lower-case letters represent complementary overlap with the linearized plasmid allowing for assembly of the fragment.

Primers (Macrogen Inc., South Korea) were diluted in sterile nuclease-free water (10 pmol/µL), and their specificity tested via conventional PCR (protocols are described in **Supplementary Tables 1–3**). PCR products were visualized on 1 % agarose gels incorporating SYBR Safe DNA Gel Stain (Invitrogen) alongside a 1 kb DNA ladder (New England Biolabs).

Gene inserts were cleaned and purified from the PCR while the linearized plasmid was extracted from gel using the GeneJET Gel Extraction and DNA Cleanup Micro Kit (ThermoFisher Scientific). DNA concentrations were measured (NanoDrop 2000 v1.6.198). Gibson assembly was performed using the NEBuilder HiFi protocol with a 1:2 vector-to-insert molar ratio and incubation at 50 °C for 20 min. The assembled products were transformed into chemically competent *E. coli* DH5α cells via heat shock at 42 °C for 30 s and recovered in SOC medium (New England Biolabs) at 37 °C for 60 min. An aliquot (100 µL) of transformed cells was plated onto lysogeny broth (LB)-kanamycin agar (kanamycin at 50 µg/mL) and incubated overnight at 37□°C. Four colonies per construct were transferred in separate universals containing LB-kanamycin broth (kanamycin at 50 µg/mL) for overnight incubation at 37 °C.

The transformed cells were screened via PCR using T7 forward (5′-TAATACGACTCACTATAGGG-3′) and T7 terminator (5′-GCTAGTTATTGCTCAGCGG-3′). PCR protocols are described in **Supplementary Tables 4–5**. The PCR products were visualized on 1 % agarose gel stained with SYBR Safe DNA Gel Stain alongside the 1 kb DNA ladder with the amplicon sizes expected to be 1.2 kb. Two positive clones per gene were selected for downstream processing.

Glycerol stocks were prepared for PCR-positive clones and stored at –80 °C. PCR products of the selected clones were cleaned using the GeneJET Gel Extraction and DNA Micro kit, and their DNA concentration measured using NanoDrop 2000 v1.6.198. Products were adjusted to 10□ng/µL in a final volume of 15□µL sterile nuclease-free water and were sent for Sanger sequencing (Source BioScience; Cambridge). From the same overnight cultures, plasmids were extracted using the Monarch Plasmid Miniprep Kit (New England Biolabs). Their DNA was quantified using NanoDrop 2000 v1.6.198 and the plasmids were stored at – 20□°C.

Plasmids were transformed into chemically competent *E. coli* BL21 (DE3) cells via heat shock at 42 °C for 45 s and recovered in SOC medium. An aliquot (100□µL) was plated on LB-kanamycin agar and incubated overnight at 37 °C. A single colony from each plate was transferred in LB-kanamycin broth for overnight incubation at 37 °C. Following the incubation, glycerol stocks were prepared and stored at –80 °C.

### Production of protein

A single colony of *E. coli* BL21 (DE3) containing an HH expression plasmid was inoculated into LB-kanamycin and incubated overnight at 37 °C. An aliquot (500 µL) of the overnight culture was used to inoculate 50 mL of fresh LB-kanamycin broth, which was incubated at 37 °C until OD_600_ ∼0.6 was reached. Protein production was induced by addition of isopropyl β-D-thiogalactoside (IPTG) to a final concentration of 1 mM. The culture was incubated with shaking at 24 °C and 200 rpm for 16 h. Cells were harvested by centrifugation (6000□***g***) and resuspended in sterile phosphate-buffered saline (PBS). Cell lysis was performed by sonication on ice (10□×□30 s bursts, 15 Lµm). The insoluble material was collected by centrifugation (30 min, 20000 ***g***, 4 °C). The supernatant was filter-sterilized by passing through 0.45 µm and then 0.22 µm filters (Sartorius) to obtain the soluble fraction. The remaining pellet was resuspended in 5 mL of PBS, transferred to a new tube, and labelled as the insoluble fraction.

### Immobilized metal-ion affinity chromatography, protein concentration and purification

Protein purification was performed using the Proteus Ni-IDA midi kit (Protein Ark). Fresh chromatography buffers were prepared in PBS (pH 7) containing imidazole at final concentrations of 10 mM (binding buffer), 30 mM (wash buffer), and 300 mM (elution buffer). The soluble fraction was loaded onto a nickel affinity column pre-equilibrated with binding buffer. After washing with the imidazole-containing wash buffer, bound proteins were eluted using the high-imidazole elution buffer. The eluted protein was collected for subsequent analysis.

The eluted protein was concentrated, and the buffer was exchanged using a Vivaspin□20 ultrafiltration device (10 kDa MWCO) (Cytiva) with PBS until the final volume was approximately 1 mL. The purified protein was collected and stored in the freezer.

Samples collected at different stages (BI, before induction; CL, crude lysate; IF, insoluble fraction; SF, soluble fraction; EP, eluted protein; PP, purified protein; FT, flow-through) were mixed with Laemmli sample buffer, denatured at 95 °C, and run on a 4–12 % Tris-glycine gel. The gel was stained with with PageBlue (Thermo Fisher), destained overnight, and stored in water.

The protein concentration in the samples was determined using the Bradford assay (Bio-Rad) with BSA standards, which were serially diluted from 2 mg/mL (Pierce Bovine Serum Albumin Standard Ampules; Thermo Scientific).

The DNA sequence of each gene was translated into the corresponding protein sequence using the <u>Expasy Translate</u> tool. The resulting protein sequences were analysed using the <u>Expasy Compute pI/Mw</u> tool to determine the theoretical isoelectric point and molecular weight of each protein.

### Ninhydrin assay with purified proteins

Sodium hippurate (100 mM) and ninhydrin reagent solutions were prepared as described above. For the assay, 100 µL of diluted purified protein at different concentrations was added into individual wells of a 96-well plate in duplicates. Then, 100 µL of the sodium hippurate solution was added to each well. The plates were incubated under aerobic conditions at 37L°C for 24 h. Following incubation, 50 µL of ninhydrin reagent was added to each well. The plates were re-incubated under respective conditions at 37 °C for an additional 30 min. Empty vector control was used as the negative control, expressed and purified under the same conditions as the proteins. *A*_560_ was measured using the microplate reader to quantify the reaction intensity. Three biological replicates were completed (two technical replicates each).

### Michaelis–Menten curve

Group 1 proteins Ko6_01540 and PSKpn11_20955 were included in enzyme assays, carried out as follows. Hippurate solutions (in duplicate) were prepared in deep-well plates (BRAND; Fisher Scientific) to give final concentrations of 0, 5, 10, 15 and 20 mM sodium hippurate in 1 mL volumes. The plates were incubated at 37 °C overnight to equilibrate. Purified protein (10 µL) was added per reaction: 5 µg of Ko6_01540 and 0.5 µg for PSKpn11_20955. The plates were immediately incubated to 37 °C to allow the reactions to proceed. Aliquots (100 µL) were removed at 5, 10, 15, 30, 45 and 60 min after enzyme addition. For each substrate concentration and timepoint, a corresponding substrate control (sodium hippurate solution only) was processed in parallel and used as a blank to correct for non-enzymatic background. Each 100 µL reaction aliquot was mixed with 150 µL ninhydrin reagent in a 96-well microplate (technical duplicates for each timepoint and condition). Plates were incubated at 70 °C for 30 min to allow colour development, then cooled at room temperature for 20 min. Absorbance was measured at 560 nm using the microplate reader.

For each substrate concentration, timepoint and enzyme, the mean absorbance of duplicate wells was calculated and corrected for background (i.e. blank) readings. For determination of initial reaction rates, the 10-min timepoint was used (within the linear phase of product formation, based on the full time-course) for PSKpn11_20955, whereas the 45-min timepoint was used for Ko6_01540. The approximate initial reaction rate at each substrate concentration [S] was calculated as *v* = *A*_corr_ [S] / time, where *v* is the rate of reaction giving rates in absorbance units per minute (*A*/min). For each enzyme, initial reaction rates (*v*) were plotted against [S] (mM) in GraphPad Prism. Non-linear regression was performed using the built-in Michaelis–Menten model, V=V_max_[S]/(Kffi+[S]), with [S] as the *x* variable and *v* as the *y* variable. Default settings and the constraint Kffi>0 were used. The best-fit values of *V*_max_ and *K*_m_, along with 95 % confidence intervals and goodness-of-fit statistics (*R*²) were reported for each protein.

## Results

### Ninhydrin assays

All seven of our *Klebsiella* strains isolated from urine (**Table 1**) could hydrolyse hippurate, with the release of glycine detected by formation of the characteristic deep-purple colour in the ninhydrin assay (**Figure 1**). *K. oxytoca* and *K. pneumoniae* strains were better able to hydrolyse hippurate under microaerobic than aerobic conditions. *K. pneumoniae* PS_Kpn11 exhibited significantly (adjusted P value 0.0158) higher activity under microaerobic than aerobic conditions. Sodium hippurate promoted the growth of *K. oxytoca* and *K. pneumoniae* (**Supplementary Figure 1**).

**Figure 1.**
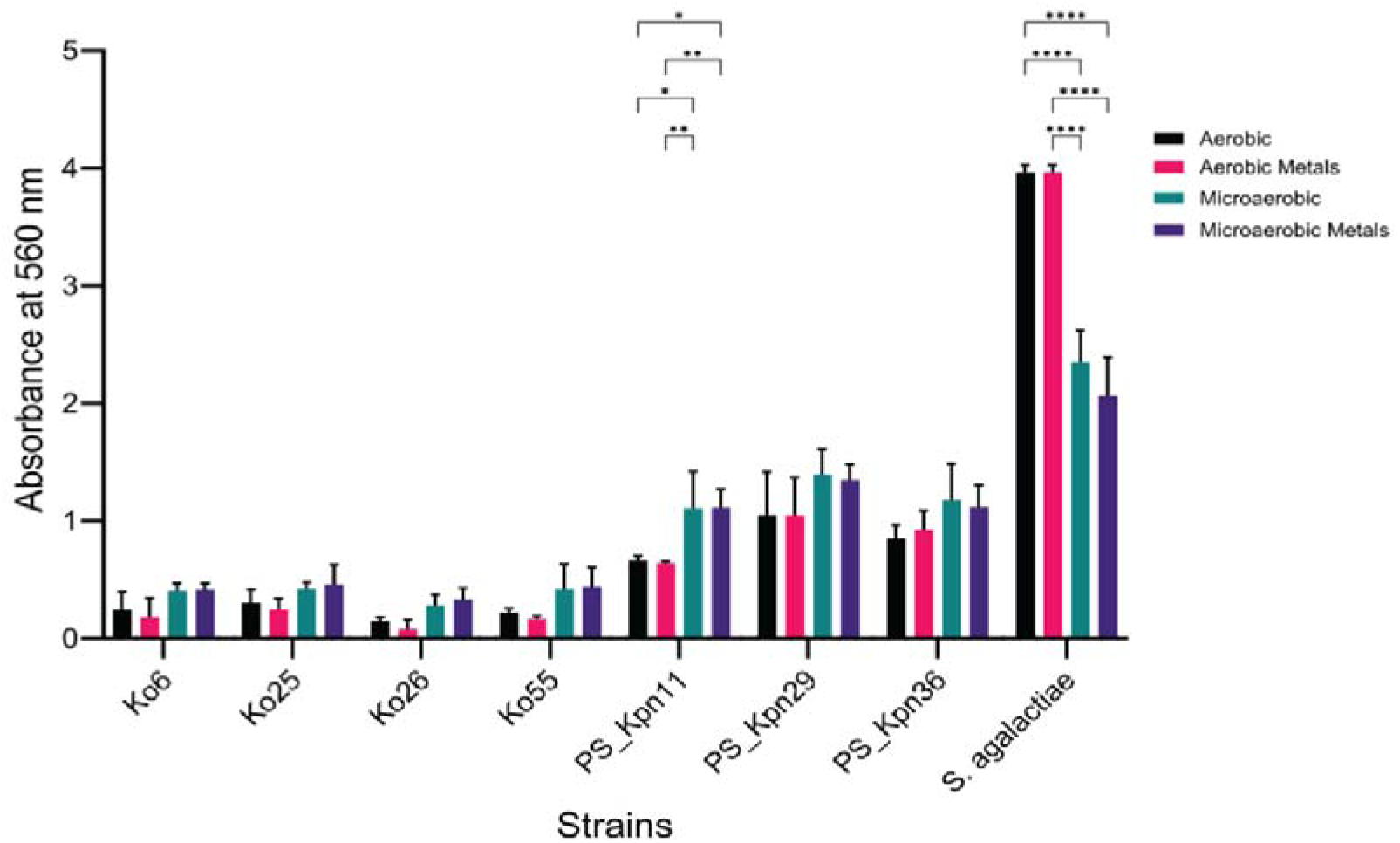
Ninhydrin assay to assess ability of *Klebsiella* strains to hydrolyse hippurate. Hippurate hydrolysis was quantified using a 96-well-based assay. Data are shown for the mean + standard deviation for three biological replicates (three technical replicates each), including the positive control *S. agalactiae*. Values for the negative control (*S. pyogenes*) have been subtracted from data. Statistical analysis was undertaken using two-way ANOVA with Tukey’s *post hoc* test testing. Adjusted P values: *, <0.05; **, <0.01; ****, <0.0001.

We undertook the ninhydrin assay in the presence and absence of trace metals, because analyses of the protein sequences predicted to encode HHs in our strains showed them to include metal-binding sites (**Supplementary Figure 2**) and the carboxypeptidase HHs of *C. jejuni* and *Pseudomonas* species have been shown to be inhibited by some metal ions ^18,39,40^. The presence of the trace metal mixture had no effect on the activity of *Klebsiella* isolates (**Figure 1**).

### Comparative sequence analyses

UniProt was used to search for HHs predicted to be encoded by bacteria. The only manually annotated and reviewed record retrieved belonged to *C. jejuni* subsp. *jejuni* ATCC 700819 serotype O:2 (UniProt P45493). The UniProt sequences of *Klebsiella* species identified as HHs shared 37.7–45.3 % amino acid (aa) identity with P45493. *Klebsiella* species sequences shared 36.2–97.7 % aa identity among themselves, with three distinct groups of predicted HH sequences represented in the data (**Figure 2A**).

**Figure 2.**
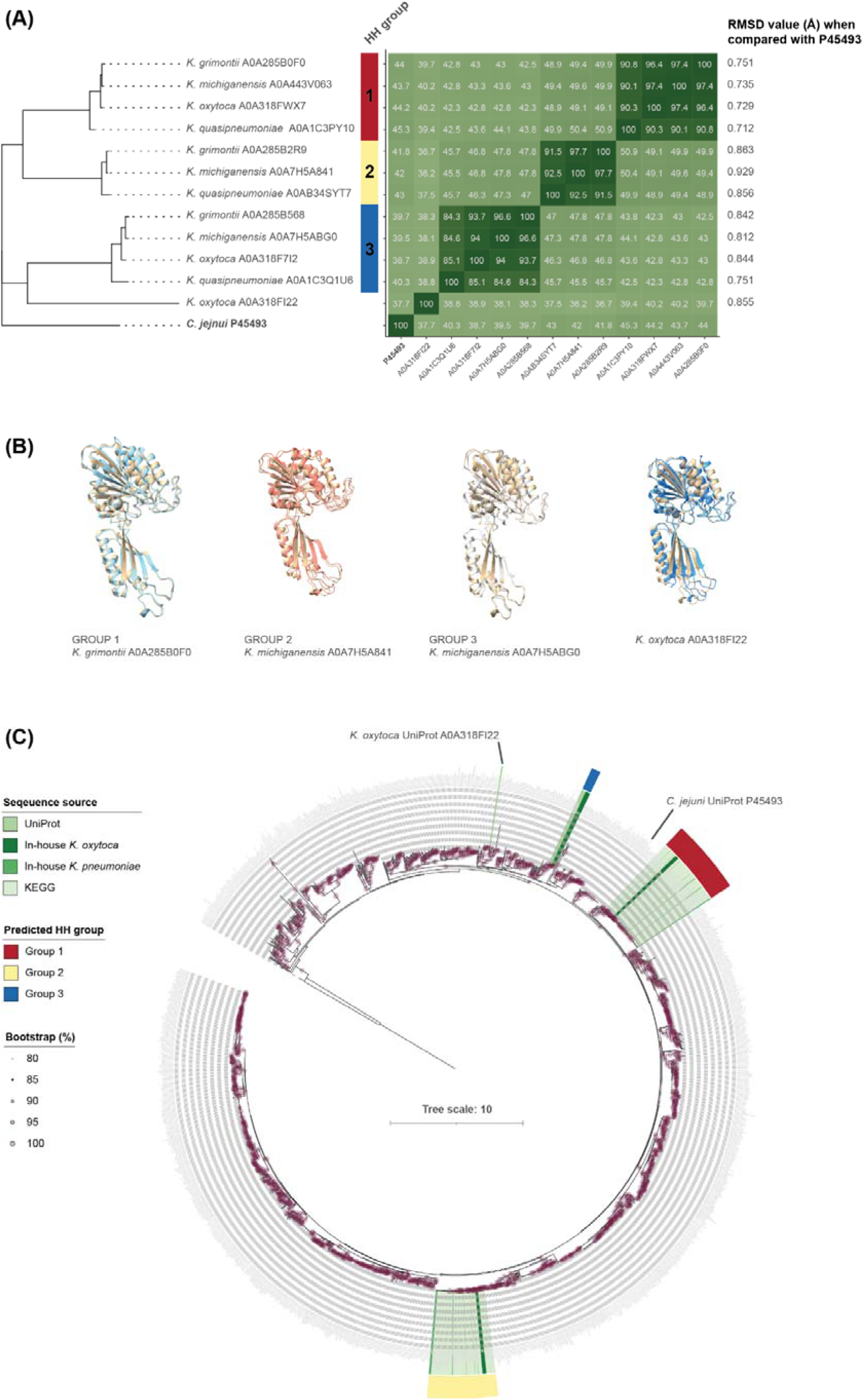
Identification of potential hippurate hydrolases (HHs) among *Klebsiella* species. **(A)** Comparison of *Klebsiella* sequences identified in UniProt with the *C. jejuni* reference sequence. The matrix shows amino acid identity values (%) among the sequences. The RMSD data were generated from ChimeraX (MatchMaker) comparisons of the AlphaFold-predicted structures of the *Klebsiella* proteins with the structure of the *C. jejuni* protein. **(B)** AlphaFold-predicted structures for representative *Klebsiella* sequences overlaid onto the *C. jejuni* protein structure (shown in the cream colour in each image). **(C)** Comparison of UniProt, KEGG orthology K01451 and in-house predicted HH protein sequences, confirming the grouping of *Klebsiella* sequence data into three distinct groups of proteins. The phylogenetic tree, rooted at the midpoint, was constructed using iqtree (Q.PFAM+I+R10 model) with bootstrap values presented as a percentage of 1000 replicates (UFBoot). Information on all bacterial protein sequences included in the analysis can be found in **Supplementary Table 6**.

We used AlphaFold2 to predict the structures of the proteins, and compared their structures using ChimeraX. The structural alignments indicated that the *Klebsiella* and *C. jejuni* proteins had near identical structures (**Figure 2B**), with C_α_ RMSD values of <2 Å (**Figure 2A**) suggesting they potentially had the same function – i.e. they could all hydrolyse hippurate. This interpretation is supported by the conservation of active site residues (**Supplementary Figure 2**).

HH KEGG orthology K01451 (*N*-benzoylglycine amidohydrolase) included 1684 entries across 1022 genomes at the time the analysis was undertaken (n=10 eukaryotes, n=1012 bacteria; **Supplementary Table 6**). Members of the phylum *Pseudomonadota* dominated the dataset (*Gammaproteobacteria* > *Betaproteobacteria* > *Alphaproteobacteria*). Within the *Enterobacteriaceae* (class *Gammaproteobacteria*, order *Enterobacterales*), *Klebsiella* represented 52/126 (41.3 %) genomes (**Supplementary Figure 3**): n=32 *K. pneumoniae* species complex; n=9 *K. oxytoca* species complex; n=11 other *Klebsiella*. *Klebsiella* and *Pluralibacter* genomes encoded an average of two copies of HH each according to the KEGG-based analysis, whereas all other *Enterobacteriaceae* encoded one copy of HH (**Supplementary Table 6**).

The UniProt and KEGG *Klebsiella* sequence data were used to generate a BLASTP database against which we searched the Bakta-predicted protein sequences for our seven *Klebsiella* isolates. Three and two different putative HHs were detected in the *K. oxytoca* and *K. pneumoniae* strains, respectively (**Supplementary Figure 2**; aa identity values are given in **Supplementary Figure 4**). Using the group labelling given in **Figure 2A**, group 1 included three *K. pneumoniae* (PSKpn11_20955, PSKpn36_24080, PSKpn29_03320) and four *K. oxytoca* (Ko25_02535, Ko6_01540, Ko26_10830, Ko55_25585) sequences, group 2 included three *K. pneumoniae* (PSKpn11_15975, PSKpn29_10165, PSKpn36_15870) and four *K. oxytoca* (Ko25_09835, Ko55_27995, Ko6_10420, Ko26_13045) sequences, and group 3 comprised only *K. oxytoca* sequences (Ko26_04715, Ko25_05940, Ko6_06555, Ko55_16140).

Comparison with the KEGG HH sequence data confirmed the clustering of our *Klebsiella* sequence data into three distinct clades, each with 100 % bootstrap support (**Figure 2C**). Group 1 comprised a clade of 64 sequences that included a *Pluralibacter gergoviae* sequence (KEGG pge_LG71_16955) and one from *Enterobacteriaceae* bacterium S05 (KEGG ebu_CUC76_08835). The Group 2 clade included 61 sequences, one of which belonged to *Enterobacteriaceae* bacterium S05 (KEGG ebu_CUC76_04190). PubMLST ^41^ and skani ^42^ analyses identified *Enterobacteriaceae* bacterium S05 as *K. quasipneumoniae* (100 % PubMLST; 96.6 % average nucleotide identity with the type strain of *K. quasipneumoniae*, genome accession GCF_020525925.1). The Group 3 clade comprised only the four UniProt and four in-house *K. oxytoca* sequences described above, with these sequences closely related (67.9–70.9 % aa identity) to three *Pantoea ananatis* sequences (KEGG plf_PANA5342_3025, paj_PAJ_0581 and paq_PAGR_g2899). *K. oxytoca* UniProt A0A318FI22 formed a clade (100 % bootstrap support; 64.0–65.8 % aa identity) with one *Paenibacillus* (KEGG psab_PSAB_05390) and three *Bacillus* (KEGG bqy_MUS_0512, bami_KSO_016960 and baq_BACAU_0478) sequences, distantly related to all other *Klebsiella* sequences. Because no other *Klebsiella* sequences were associated with A0A318FI22, we did not consider this sequence in further analyses.

### Cloning, transformation and assessment of protein activity

Strains *K. oxytoca* Ko6 and *K. pneumoniae* PS_Kpn11 were randomly selected as sources of proteins for functional characterization work, with each protein predicted to encode a HH in both strains (**Table 2**) cloned, transformed and produced. Group 1 and group 3 proteins could be purified as soluble proteins, whereas group 2 proteins were found predominantly in the insoluble fraction (**Figure 3A**). The group 1 proteins showed the highest activity in the ninhydrin assay, with the group 3 protein showing no detectable activity (**Figure 3B**). Group 2 proteins showed high HH activity in the insoluble and soluble fractions despite not being successfully purified (**Figure 3C**). The results for all three groups were consistent under both aerobic and microaerobic conditions, indicating that oxygen availability did not significantly influence enzyme activity (data not shown).

**Figure 3.**
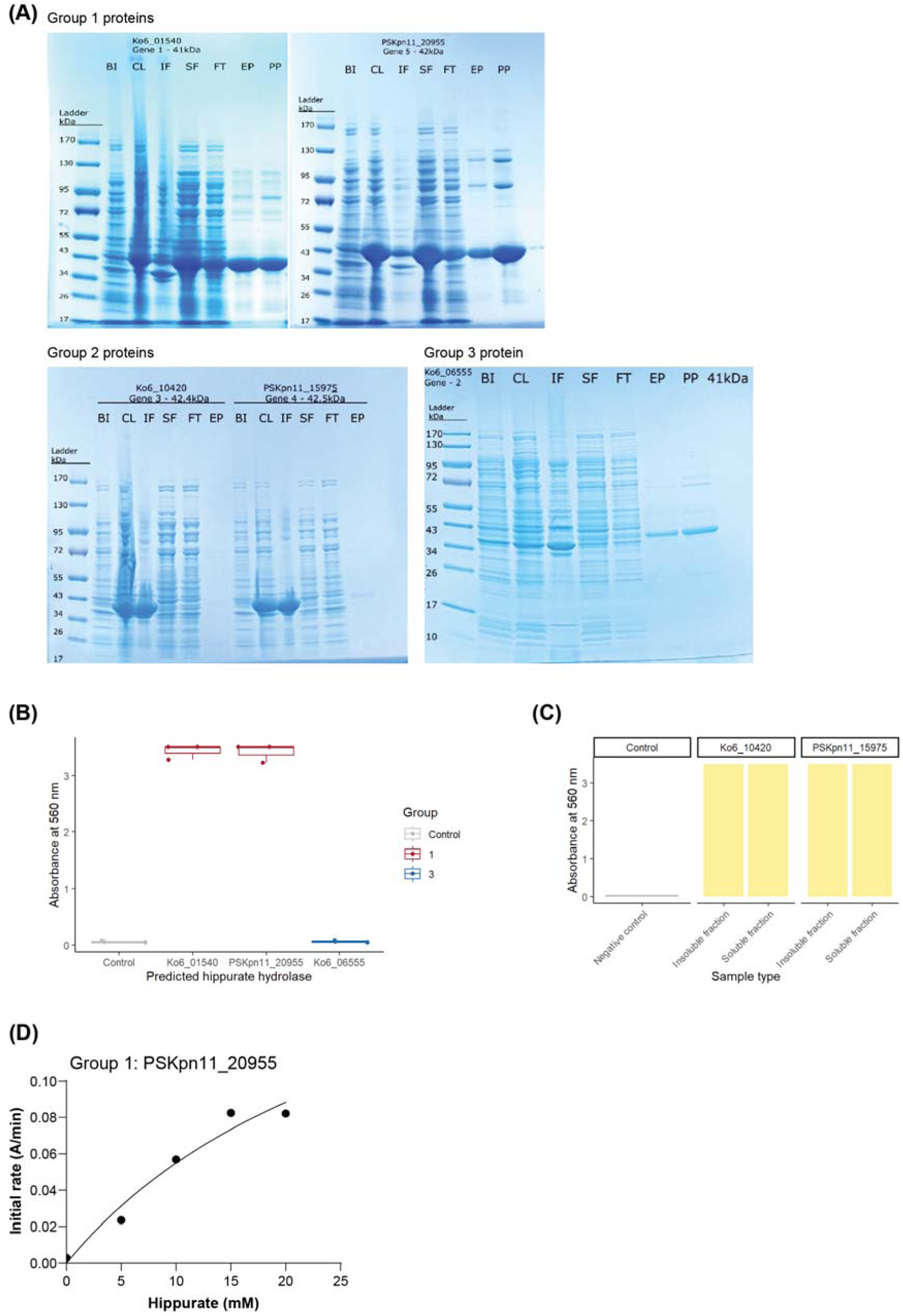
Purification of predicted HH proteins and assessment of their activity. **(A)** Images of SDS-PAGE gels for proteins belonging to each of the three predicted HH groups. BI, before induction; CL, crude lysate; IF, insoluble fraction; SF, soluble fraction; FT, flow-through; EP, eluted protein; PP, purified protein. **(B)** Assessment of activity of purified group 1 and group 3 proteins. Data are shown for three biological replicates. Amount of purified protein included in each assay: negative control, 0.017 mg/mL; Ko6_01540, 0.05 mg/mL; PSKpn11_20955, 0.05 mg/mL; Ko6_06555, 0.007 mg/mL. **(C)** Assessment of activity of insoluble and soluble fractions of group 2 proteins. Data are shown as mean of two biological replicates. Amount of protein included in each assay: negative control, 0.017 mg/mL; Ko6_10420, 0.018 mg/mL (insoluble fraction) and 0.017 mg/mL (soluble fraction); PSKpn11_15975, 0.013 mg/mL (insoluble fraction) and 0.025 mg/mL (soluble fraction). **(D)** Michaelis–Menten curve generated for the group 1 protein PSKpn11_20955 (0.05 mg/mL); three biological replicates with duplicate technical replicates.

Group 1 protein PSKpn11_20955 gave a reproducible kinetic profile in the Michaelis–Menten analysis (**Figure 3D**), with a *V*_max_ of 0.23 A/min and an apparent *K*_m_ of 30.99 mM and a high goodness of fit (*R*^2^=0.962). Group 1 protein Ko6_01540 produced a moderate goodness of fit (*R*^2^=0.9483), with large estimated *V*_max_ and *K*_m_ values and wide confidence intervals (**Supplementary Table 7**). However, it should be noted that assays with Ko6_01540 were hampered (across a range of protein concentrations) by the ninhydrin assay becoming cloudy.

### Identification of HHs across the genus *Klebsiella*

We sought to determine the prevalence of the group 1 and group 2 proteins across the genus *Klebsiella*. Proteome data for 3012 genomes representing 20 species of the genus *Klebsiella* were included in the BLASTP-based analysis (**Supplementary Table 8**). The majority (2885/3012, 95.8 %) of the genomes encoded full-length group 1 (392 aa) and group 2 (389 aa) proteins, with the concatenated sequences clustering based on species affiliation (**Figure 4**).

**Figure 4.**
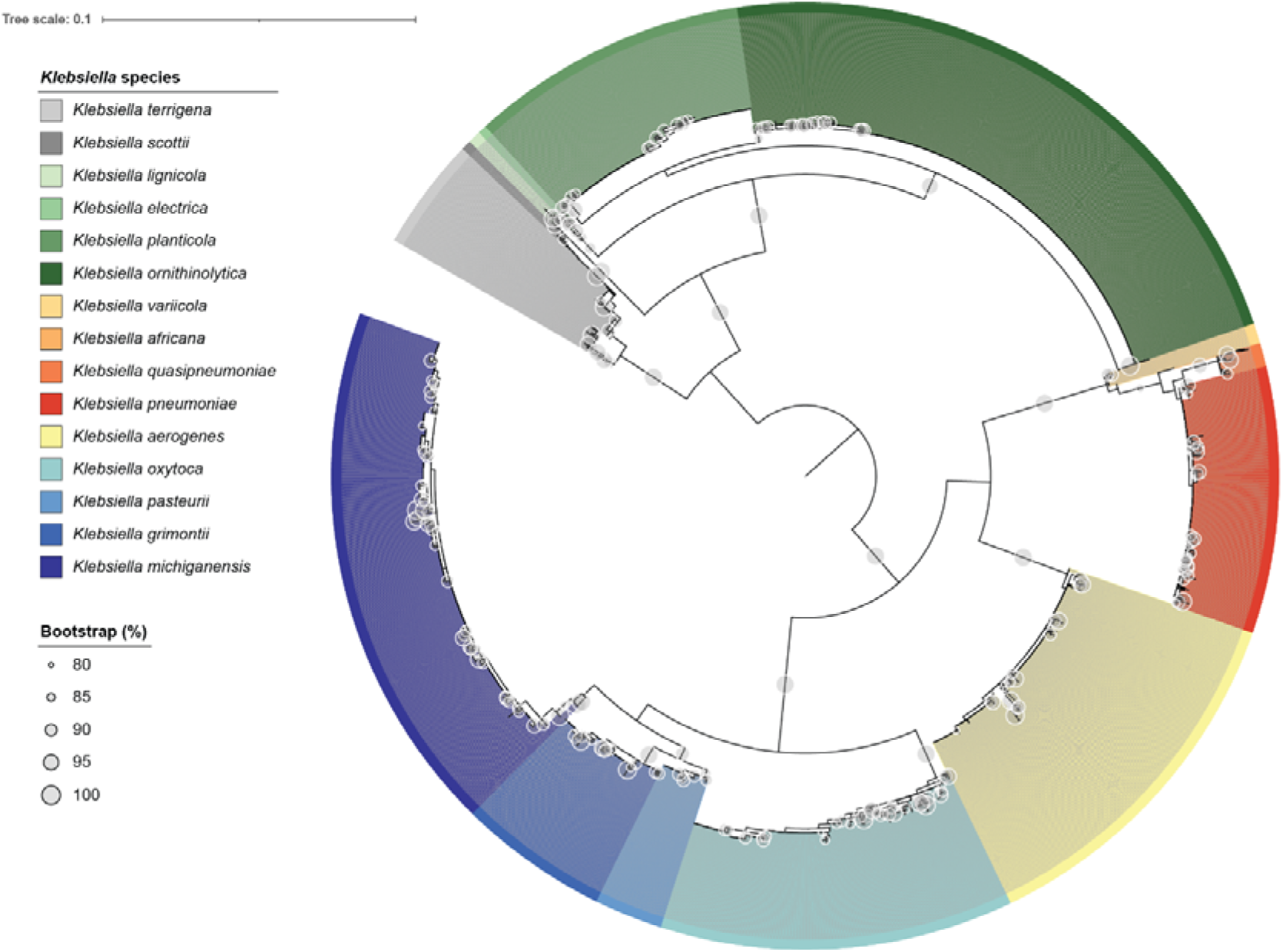
Hippurate hydrolases are prevalent among many *Klebsiella* species. Sequence data derived from 2885/3012 *Klebsiella* genomes are represented in the phylogenetic tree. Both group 1 and group 2 proteins were found in genomic data of 15/20 (75 %) of *Klebsiella* species. Number of genomes across entire dataset (2885/3012) that included both proteins: *Klebsiella africana* – 1/1, 100.0 %; *Klebsiella electrica* – 10/11, 90.9 %; *Klebsiellal scottii* – 10/11, 90.9 %; *Klebsiella lignicola* – 13/13, 100.0 %; *Klebsiella variicola* – 19/20, 95.0 %; *Klebsiella quasipneumoniae* – 23/23, 100.0 %; *Klebsiella pasteurii* – 68/68, 100.0 %; *Klebsiella terrigena* – 113/114, 99.1 %; *Klebsiella grimontii* – 153/154, 99.4 %; *Klebsiella pneumoniae* – 270/285, 94.7 %; *Klebsiella planticola* – 280/309, 90.6 %; *Klebsiella oxytoca* – 358/383, 93.5 %; *Klebsiella aerogenes* – 372/381, 97.6 %; *Klebsiella michiganensis* – 539/549, 98.2 %; *Klebsiella ornithinolytica* – 656/672, 97.6 %. The phylogenetic tree, rooted at the midpoint, was constructed using iqtree (Q.BIRD+I+R3 model) with bootstrap values presented as a percentage of 1000 replicates (UFBoot).

Only a small number of genomes did not encode a copy of the group 1 protein (46/3012, 1.5 %) or the group 2 protein (16/3012, 0.5 %). Four of the five *K. huaxiensis* genomes encoded only the group 1 protein; the other *K. huaxiensis* genome (accession GCF_036441055) returned no HH hits. The five *K. indica* genomes encoded only the group 1 protein. All six *K. spallanzanii* genomes encoded the group 1 protein, with five lacking the group 2 protein and one (accession GCF_902158595) encoding a truncated variant of the group 2 protein. The sole representatives of “*Klebsiella quasivariicola*” and a novel *Klebsiella* sp. ^43^ included in the analysis lacked group 1 and group 2 proteins, respectively (**Supplementary Table 8**).

## Discussion

Hippurate is an MHCM found in high abundance in human urine, formed by conjugation of gut-microbiota-derived benzoate and naturally available benzoate with glycine ^1–3^. Our previous metabolomics work showed that UTI-associated *K. pneumoniae* could degrade hippurate present in human urine ^12^. Little is known about hippurate hydrolysis in *Klebsiella*, with limited phenotypic data available for *K. aerogenes* and *K. pneumoniae* ^13,14^. In contrast, hippurate hydrolysis has been extensively studied in *Campylobacter* species and streptococci for differentiation and identification purposes ^13,15,17–19,25^. Cytoplasmic HHs from *Pseudomonas* sp. KT 801, *Pseudomonas putida* and *Rhodococcus equi* have been purified and functionally characterized, but their sequences have not been described ^39,40,44^. The aim of this work was to confirm that *Klebsiella* species hydrolyse hippurate and to identify the genes encoding the enzyme(s) responsible for this bioconversion.

In this study strains of *K. oxytoca* (n=4) and *K. pneumoniae* (n=3) isolated from urine hydrolysed hippurate, releasing glycine from hippurate in ninhydrin assays to produce the characteristic deep-purple colour indicative of amidohydrolase activity (**Figure 1**). *K. pneumoniae* strains were better able to hydrolyse hippurate than the *K. oxytoca* strains under aerobic and microaerobic conditions, with the latter representative of the hypoxic bladder microenvironment. Strain-specific differences in hippurate breakdown have been observed for *K. pneumoniae* and *C. jejuni*, but the factor(s) influencing these difference are unknown ^12,14,45^. Due to their detection limits the ninhydrin-based (millimolar) and NMR-based (micromolar) approaches we have used to characterize hippurate catabolism lack the sensitivity required to reliably quantify strain-and species-specific differences among *Klebsiella*, so will be replaced by a targeted mass spectrometry-based assay in our future work.

Structural analyses identified potential HHs encoded by *Klebsiella* species and our strains, with the proteins sharing low amino acid identify (37.7–45.3 %) with each other and the *C. jejuni* reference sequence (**Figure 2**). Our *Klebsiella* strains encoded three different potential HHs all related to sequences associated with KEGG orthology K01451, representing HH (reaction *N*-benzoylglycine amidohydrolase). Group 1 (392 aa) and group 2 (389 aa) proteins were found in all *K. pneumoniae* and *K. oxytoca* strains, while group 3 proteins (383 aa) were only found in *K. oxytoca* strains. The molecular weights of our predicted HHs (group 1, 42.0 kDa; group 2, 42.4–42.5 kDa; group 3, 41.0 kDa; **Table 2**) were in agreement for those of other carboxypeptidases (40–48 kDa) ^18^.

We cloned, produced and purified the group 1 and group 3 proteins, demonstrating the group 1 proteins were functional under the conditions tested here. Enzyme activity of carboxypeptidases such as HH is known to be greatly affected by incubation temperature ^40,46^. The group 3 protein may be functional at higher temperatures or hippurate may not be its substrate, but due to time constraints we were not able to assess these parameters. Enzyme kinetic data were generated for the group 1 protein PSKpn11_20955. All attempts to generate reliable kinetic data for group 1 protein Ko6_01540 failed, due to the ninhydrin assay developing a cloudy appearance that prevented measurement of absorbance values. This cloudiness could have been caused by protein denaturation, phase separation, or high salt concentrations may have remained in the purified protein causing the precipitation of some compounds ^47^. Cloudiness has been reported previously for the ninhydrin assay ^47^.

We were unable to purify the group 2 HHs, but protein present in the soluble and insoluble fractions was able to hydrolyse hippurate. Expression of the medically important *Pseudomonas* sp. RS-16 carboxypeptidase G2 (glucarpidase) in *E. coli* leads to the formation of inclusion bodies ^48^. Gel images for our group 2 HHs (**Figure 3A**) match those for carboxypeptidase G2 produced in *E. coli* ^48^, suggesting our group 2 proteins also form inclusion bodies. It is likely the expressed recombinant group 2 proteins are present as stable, insoluble homomultimers. Optimization of the production of recombinant carboxypeptidase G2 has included identification of an optimal signal peptide to export the expressed protein into the periplasmic space to allow proper protein folding ^49^. Its production has also been improved by incubation at 20–25 °C, along with inclusion of glycerol, tryptone and peptone in the growth medium ^48,49^. Further work is required to determine whether adoption of approaches outlined above will allow us to recover abundant group 2 soluble proteins.

Whether our proteins are functional in homomultimer forms of four to six subunits ^40,44,46,50^ remains to be determined empirically. SWISS-MODEL predicts the *C. jejuni* HH and our group 1 protein PSKpn11_20955 to be a homotetramer, by comparison with the crystal structure of a putative aminohydrolase (M20 family) from methicillin-resistant *Staphylococcus aureus* (accession A0A0H2WZV8). The crystal structure of a *K. pneumoniae* aminobenzoyl-glutamate utilization protein (hydrolase), identified by inputting our UniProt sequences into SWISS-MODEL and Phyre 2.2 (intensive mode) ^51^, is a homotetramer (https://doi.org/10.2210/pdb3IO1/pdb). Its 436 aa monomer sequence shares 25.3–26.3 % aa identity with our group 1–3 sequences and 23.3 % aa identity with the *C. jejuni* reference sequence, but its homotetramer structure is highly similar to that of the *C. jejuni* HH. The identification of this additional M20 family protein, A0A318FI22, the group 3 proteins, and variants of the group 1 and group 2 proteins (**Supplementary Table 8**) suggests the genus *Klebsiella* may harbour a diverse range of M20 family proteins that remain to be characterized.

Group 1 and group 2 HHs were prevalent across the genus *Klebsiella*, representing extended (*K. electrica*, *K. scottii*, *K. planticola*, *K. oxytoca*) and soft (*K. africana*, *K. lignicola*, *K. variicola*, *K. quasipneumoniae*, *K. pasteurii*, *K. terrigena*, *K. grimontii*, *K. pneumoniae*, *K. aerogenes*, *K. michiganensis*, *K. ornithinolytica*) core genes, present in >90 % and >95 % of genomes, respectively, in 15/20 *Klebsiella* species (**Figure 4**). The HH of *C. jejuni* has a broad substrate range [including *N*-benzoylglycine (hippurate), *N*-benzoylmethionine (methylhippurate), *N*-benzoylalanine, *N*-benzoylglutamate, *p*-aminobenzoyl-glutamate, *N*-benzoylvaline, *N*-benzoyl-phenylalanine, *N*-acetyl-glycine and *N*-acetyl-alanine] ^18^; therefore, hippurate hydrolysis may not be the core function of the amidohydrolases identified in *Klebsiella* species. Beyond hippurate, only *N*-acetyl-alanine and *N*-acetyl-glycine have been detected in human urine at micromolar quantities, representing endogenous amino acid derivatives ^52^. Therefore, they may act as additional substrates for HHs in human urine.

The group 1 HH-encoding gene was followed by genes encoding a HisJ-like periplasmic binding protein and a PdxI-like aldo/keto reductase (**Figure 5**). Organization of genes flanking these three core proteins differed across *Klebsiella* species and species complexes. In *E. coli*, HisJ binds to histidine and interacts with the HisMPQ ATP-binding cassette transporter complex for histidine uptake ^53^; therefore, the HisJ-like protein encoded adjacent to the group 1 HH gene could be a hippurate-binding protein. It is possible that the HisJ-like protein binds to a permease encoded elsewhere in the genome and delivers hippurate that is then broken down by HipO1 in the cytoplasm into benzoic acid and glycine, followed by conversion of the benzoic acid into benzaldehyde/phenol by the PdxI-like protein. The FepD/FepC-like siderophore transporter is unlikely to be the hippurate permease, as this is not always present and also comes with its own periplasmic binding protein.

**Figure 5.**
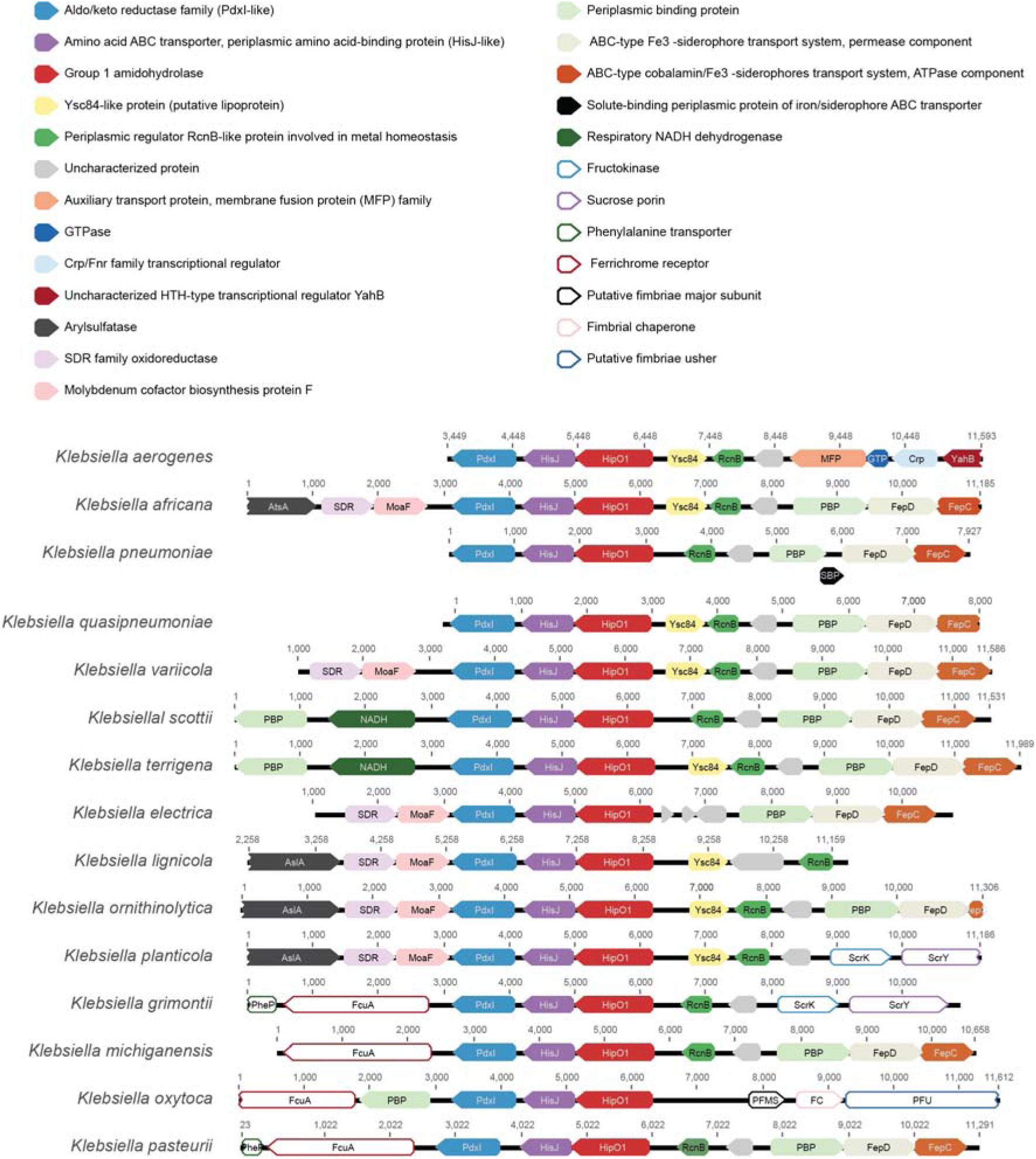
Visualization of genes flanking (+/-5000 bp) the gene encoding the group 1 HH (amidohydrolase) identified in this study. Annotated consensus sequences are shown, listed from top to bottom in terms of their affiliation with recognized *Klebsiella* species complexes ^43^ (number of non-redundant sequences included in consensus analysis, derived from *n* genomes): *K. aerogenes* (169, 372); *K. pneumoniae* species complex – *K. africana* (1, 1), *K. pneumoniae* (51, 270), *K. quasipneumoniae* (3, 23), *K. variicola* (6, 19); *K. terrigena* species complex – *K. scottii* (7, 10), *K. terrigena* (57, 113); *K. planticola* species complex – *K. electrica* (7, 10), *K. lignicola* (10, 13), *K. ornithinolytica* (319, 656), *K. planticola* (127, 280); *K. oxytoca* species complex – *K. grimontii* (88, 153), *K. michiganensis* (224, 539), *K. oxytoca* (139, 358), *K. pasteurii* (48, 68).

Arrangement of genes flanking the group 2 HH-encoding gene was highly conserved across 15 species of *Klebsiella* (**Figure 6**). In general, genes encoding components of the Sap ^54^ peptide transport system were followed by genes encoding an AraC family transcriptional regulator, the HH, a NorB-like quinolone resistance protein, the phage shock protein (PSP) system and a YcxJ family protein. All species encoded *pspFABCD*, with members of the *K. oxytoca* complex encoding *pspFABCDE*. In *E. coli*, the PSP system stabilizes the cell membrane, protects against envelope stress and maintains the proton motive force across the inner membrane of the cell ^55,56^. PspA and PspF are among the most conserved proteins in the *Enterobacteriaceae*, constituting the minimal PSP system; the *E. coli psp* operon comprises the regulon (*pspABCDE* with *pspF* and *pspG*) and the two-component *arcA*–*arcB* system, with *ycjX* and *ycjF* part of the operon in some species of bacteria ^56^. ArcA, activated by phosphorylation of ArcB, influences the strength and effectiveness of the transcriptional response to the inducing stress conditions in microaerobiosis in *E. coli* ^57^, and promotes fitness of *K. pneumoniae* in low oxygen conditions, decreased iron availability and host-mediated membrane damage in bacteraemia ^58^. Induction of the PSP via ArcAB in *Klebsiella* may partially explain why we observed higher HH activity under microaerobic than aerobic conditions. Work to determine whether the two distinct HHs are constitutively expressed or induced in the presence of relevant substrates, and to examine the overall transcriptomic responses of *K. pneumoniae* strains to hippurate exposure under hypoxia, is ongoing. However, based on preliminary qPCR data we predict the group 1 protein to be constitutively expressed and the group 2 protein to be induced in the hypoxic bladder microenvironment, facilitating scavenging of glycine from hippurate in nutrient-limited urine.

**Figure 6.**
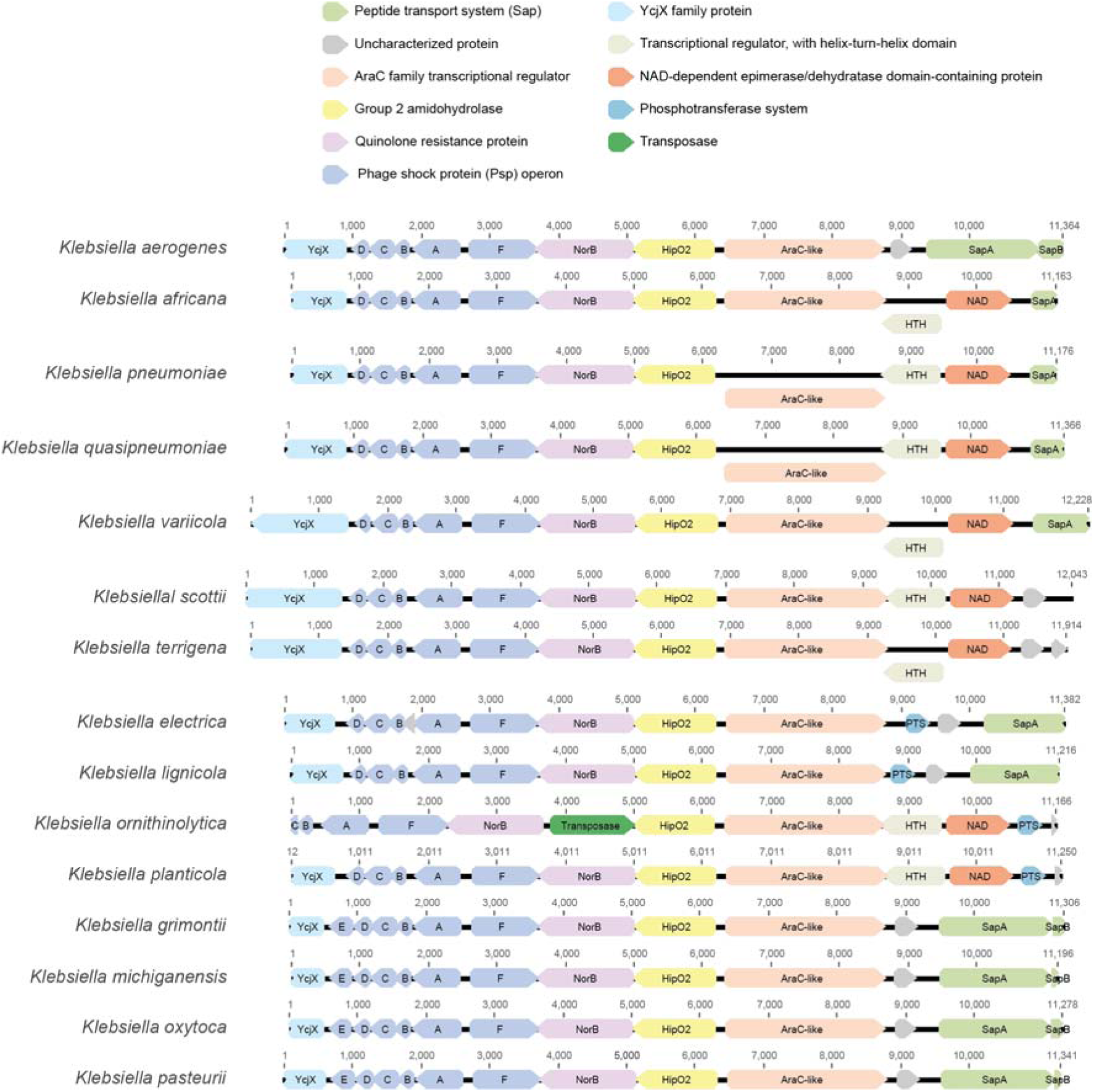
Visualization of genes flanking (+/-5000 bp) the gene encoding the group 2 HH (amidohydrolase) identified in this study. Annotated consensus sequences are shown, listed from top to bottom in terms of their affiliation with recognized *Klebsiella* species complexes ^43^ (number of non-redundant sequences included in consensus analysis, derived from *n* genomes): *K. aerogenes* (194, 372); *K. pneumoniae* species complex – *K. africana* (1, 1), *K. pneumoniae* (179, 270), *K. quasipneumoniae* (20, 23), *K. variicola* (16, 19); *K. terrigena* species complex – *K. scottii* (7, 10), *K. terrigena* (57, 113); *K. planticola* species complex – *K. electrica* (8, 10), *K. lignicola* (10, 13), *K. ornithinolytica* (337, 656), *K. planticola* (134, 280); *K. oxytoca* species complex – *K. grimontii* (80, 153), *K. michiganensis* (221, 539), *K. oxytoca* (126, 358), *K. pasteurii* (42, 68).

## Acknowledgements

LH designed the study. DD, HG and LH and visualized the data. DD undertook this work as part of an MRes research project, carrying out a comprehensive literature review, all phenotypic screening and initial bioinformatics work, primer design, and initial cloning and expression studies. HG purified and characterized the proteins. HG and SM optimized the ninhydrin assay for use with the purified proteins. ALM generated whole-genome sequence data for the study. LH undertook extensive bioinformatics analyses of the group 1 and group 2 proteins. JCL facilitated interpretation of gene data. LH and DN supervised the study. DD, HG and LH wrote the manuscript; all authors reviewed and approved the final version.

## Data availability statement

Whole-genome sequence data for strains included in this study are available from BioProject PRJNA917129 and BioProject PRJNA1439749.

## Abbreviations

Aa: amino acid
HH: hippurate hydrolase
KEGG: Kyoto Encyclopedia of Genes and Genomes
MHCM: microbiota–host co-metabolite
MSA: multiple-sequence alignment
PBS: phosphate-buffered saline
PSP: phage shock protein
RMSD: root mean square deviation
UTI: urinary tract infection.

