## Supplementary Material for "Hippurate hydrolases encoded by *Klebsiella pneumoniae* and *Klebsiella oxytoca* are functional"

**Supplementary Table 1.** Conventional PCR mix for cloning work.

| **Component** | **Reaction mix** |
| --- | --- |
| 5 x Phusion Plus buffer | 10 µL |
| 5 x GC enhancer | 10 µL |
| dNTPs | 1 µL |
| Forward Primer | 2.5 µL |
| Reverse Primer | 2.5 µL |
| Template DNA | 1 µL |
| Phusion Plus DNA Polymerase | 0.5 µL |
| Nuclease Free Water | 22.5 µL |

**Supplementary Table 2.** PCR run protocol for the genes of interest.

| **Step** | **Temperature (°C)** | **Time** | **Cycles** |
| --- | --- | --- | --- |
| Initial denaturation | 98 | 30 s | 1 |
| Denaturation  Annealing  Extension | 98  60  72 | 10 s  10 s  1 min | 35 |
| Final extension | 72 | 5 min | 1 |

**Supplementary Table 3.** PCR run protocol for the pET26b plasmid.

| **Step** | **Temperature (°C)** | **Time** | **Cycles** |
| --- | --- | --- | --- |
| Initial denaturation | 98 | 30 s | 1 |
| Denaturation  Annealing  Extension | 98  60  72 | 10 s  10 s  2.5 min | 35 |
| Final extension | 72 | 5 min | 1 |

**Supplementary Table 4.** T7 PCR mix.

| **Component** | **Reaction mix** |
| --- | --- |
| Mango Mix | 25 µL |
| T7 Forward Primer | 2.5 µL |
| T7 Reverse Primer | 2.5 µL |
| Template DNA (100 ng/µL) | 1 µL |
| Nuclease Free Water | 19 µL |

**Supplementary Table 5.** T7 PCR run protocol.

| **Step** | **Temperature (°C)** | **Time** | **Cycles** |
| --- | --- | --- | --- |
| Initial denaturation | 95 | 30 s | 1 |
| Denaturation  Annealing  Extension | 95  56  72 | 30 s  30 s  1.5 min | 35 |
| Final extension | 72 | 5 min | 1 |

**Supplementary Table 6.** Species represented in KEGG orthology K01451 (hippurate hydrolase).

| **Species** | **KEGG organism code** | **No. of copies of hippurate hydrolase** | **Type** | **Phylum** | **Class** | **Order** | **Family** | **Genus** |
| --- | --- | --- | --- | --- | --- | --- | --- | --- |
| Alternaria alternata | aalt | 1 | Eukaryote | Dothideomycetes | Pleosporomycetidae | Pleosporales | Pleosporineae | Alternaria |
| Penicillium oxalicum | pou | 1 | Eukaryote | Eurotiomycetes | Eurotiomycetidae | Eurotiales | Aspergillaceae | Penicillium |
| Coccidioides posadasii | cpw | 1 | Eukaryote | leotiomyceta | Eurotiomycetes | Eurotiomycetidae | Onygenales | Coccidioides |
| Pyrenophora tritici-repentis | ptrr | 1 | Eukaryote | Pleosporomycetidae | Pleosporales | Pleosporineae | Pleosporaceae | Pyrenophora |
| Mollisia scopiformis | psco | 1 | Eukaryote | sordariomyceta | Leotiomycetes | Helotiales | Mollisiaceae | Mollisia |
| Metarhizium brunneum | mbrn | 1 | Eukaryote | Sordariomycetes | Hypocreomycetidae | Hypocreales | Clavicipitaceae | Metarhizium |
| Cordyceps militaris | cmt | 1 | Eukaryote | Sordariomycetes | Hypocreomycetidae | Hypocreales | Cordycipitaceae | Cordyceps |
| Fusarium oxysporum | fox | 1 | Eukaryote | Sordariomycetes | Hypocreomycetidae | Hypocreales | Nectriaceae | Fusarium |
| Fusarium verticillioides | fvr | 1 | Eukaryote | Sordariomycetes | Hypocreomycetidae | Hypocreales | Nectriaceae | Fusarium |
| Pochonia chlamydosporia | pchm | 1 | Eukaryote | Sordariomycetes | Hypocreomycetidae | Hypocreales | Clavicipitaceae | Pochonia |
| Klebsiella aerogenes EA1509E | ear | 2 | Bacteria | Pseudomonadota | Gammaproteobacteria | Enterobacterales | Enterobacteriaceae | Klebsiella |
| Klebsiella aerogenes KCTC 2190 | eae | 2 | Bacteria | Pseudomonadota | Gammaproteobacteria | Enterobacterales | Enterobacteriaceae | Klebsiella |
| Klebsiella electrica | ree | 2 | Bacteria | Pseudomonadota | Gammaproteobacteria | Enterobacterales | Enterobacteriaceae | Klebsiella |
| Klebsiella ornithinolytica B6 | ror | 2 | Bacteria | Pseudomonadota | Gammaproteobacteria | Enterobacterales | Enterobacteriaceae | Klebsiella |
| Klebsiella ornithinolytica S12 | ron | 2 | Bacteria | Pseudomonadota | Gammaproteobacteria | Enterobacterales | Enterobacteriaceae | Klebsiella |
| Klebsiella planticola | rpln | 2 | Bacteria | Pseudomonadota | Gammaproteobacteria | Enterobacterales | Enterobacteriaceae | Klebsiella |
| Klebsiella sp. CTHL.F3a | klc | 2 | Bacteria | Pseudomonadota | Gammaproteobacteria | Enterobacterales | Enterobacteriaceae | Klebsiella |
| Klebsiella sp. LTGPAF-6F | kll | 2 | Bacteria | Pseudomonadota | Gammaproteobacteria | Enterobacterales | Enterobacteriaceae | Klebsiella |
| Klebsiella sp. M5al | klm | 2 | Bacteria | Pseudomonadota | Gammaproteobacteria | Enterobacterales | Enterobacteriaceae | Klebsiella |
| Klebsiella sp. X13 | rao | 2 | Bacteria | Pseudomonadota | Gammaproteobacteria | Enterobacterales | Enterobacteriaceae | Klebsiella |
| Klebsiella terrigena | rtg | 2 | Bacteria | Pseudomonadota | Gammaproteobacteria | Enterobacterales | Enterobacteriaceae | Klebsiella |
| Klebsiella grimontii | kgr | 2 | Bacteria | Pseudomonadota | Gammaproteobacteria | Enterobacterales | Enterobacteriaceae | Klebsiella |
| Klebsiella huaxiensis | klw | 1 | Bacteria | Pseudomonadota | Gammaproteobacteria | Enterobacterales | Enterobacteriaceae | Klebsiella |
| Klebsiella michiganensis E718 | koe | 2 | Bacteria | Pseudomonadota | Gammaproteobacteria | Enterobacterales | Enterobacteriaceae | Klebsiella |
| Klebsiella michiganensis HKOPL1 | koy | 2 | Bacteria | Pseudomonadota | Gammaproteobacteria | Enterobacterales | Enterobacteriaceae | Klebsiella |
| Klebsiella michiganensis KCTC 1686 | kox | 2 | Bacteria | Pseudomonadota | Gammaproteobacteria | Enterobacterales | Enterobacteriaceae | Klebsiella |
| Klebsiella michiganensis M1 | kom | 2 | Bacteria | Pseudomonadota | Gammaproteobacteria | Enterobacterales | Enterobacteriaceae | Klebsiella |
| Klebsiella oxytoca CAV1374 | koc | 2 | Bacteria | Pseudomonadota | Gammaproteobacteria | Enterobacterales | Enterobacteriaceae | Klebsiella |
| Klebsiella oxytoca KONIH1 | kok | 2 | Bacteria | Pseudomonadota | Gammaproteobacteria | Enterobacterales | Enterobacteriaceae | Klebsiella |
| Klebsiella pasteurii | kpas | 2 | Bacteria | Pseudomonadota | Gammaproteobacteria | Enterobacterales | Enterobacteriaceae | Klebsiella |
| Klebsiella africana | kar | 2 | Bacteria | Pseudomonadota | Gammaproteobacteria | Enterobacterales | Enterobacteriaceae | Klebsiella |
| Klebsiella pneumoniae 30660/NJST258_1 | kpa | 2 | Bacteria | Pseudomonadota | Gammaproteobacteria | Enterobacterales | Enterobacteriaceae | Klebsiella |
| Klebsiella pneumoniae 30684/NJST258_2 | kps | 2 | Bacteria | Pseudomonadota | Gammaproteobacteria | Enterobacterales | Enterobacteriaceae | Klebsiella |
| Klebsiella pneumoniae 32192 | kpne | 2 | Bacteria | Pseudomonadota | Gammaproteobacteria | Enterobacterales | Enterobacteriaceae | Klebsiella |
| Klebsiella pneumoniae 34618 | kpnu | 2 | Bacteria | Pseudomonadota | Gammaproteobacteria | Enterobacterales | Enterobacteriaceae | Klebsiella |
| Klebsiella pneumoniae blaNDM-1 | kpb | 2 | Bacteria | Pseudomonadota | Gammaproteobacteria | Enterobacterales | Enterobacteriaceae | Klebsiella |
| Klebsiella pneumoniae CG43 | kpi | 2 | Bacteria | Pseudomonadota | Gammaproteobacteria | Enterobacterales | Enterobacteriaceae | Klebsiella |
| Klebsiella pneumoniae JM45 | kpj | 1 | Bacteria | Pseudomonadota | Gammaproteobacteria | Enterobacterales | Enterobacteriaceae | Klebsiella |
| Klebsiella pneumoniae KCTC 2242 | kpo | 2 | Bacteria | Pseudomonadota | Gammaproteobacteria | Enterobacterales | Enterobacteriaceae | Klebsiella |
| Klebsiella pneumoniae Kp52.145 | kpnk | 2 | Bacteria | Pseudomonadota | Gammaproteobacteria | Enterobacterales | Enterobacteriaceae | Klebsiella |
| Klebsiella pneumoniae PMK1 | kpx | 2 | Bacteria | Pseudomonadota | Gammaproteobacteria | Enterobacterales | Enterobacteriaceae | Klebsiella |
| Klebsiella pneumoniae subsp. pneumoniae 1084 (serotype K1) | kpp | 2 | Bacteria | Pseudomonadota | Gammaproteobacteria | Enterobacterales | Enterobacteriaceae | Klebsiella |
| Klebsiella pneumoniae subsp. pneumoniae ATCC 43816 KPPR1 | kpt | 2 | Bacteria | Pseudomonadota | Gammaproteobacteria | Enterobacterales | Enterobacteriaceae | Klebsiella |
| Klebsiella pneumoniae subsp. pneumoniae HS11286 | kpm | 2 | Bacteria | Pseudomonadota | Gammaproteobacteria | Enterobacterales | Enterobacteriaceae | Klebsiella |
| Klebsiella pneumoniae subsp. pneumoniae KPNIH10 | kpc | 2 | Bacteria | Pseudomonadota | Gammaproteobacteria | Enterobacterales | Enterobacteriaceae | Klebsiella |
| Klebsiella pneumoniae subsp. pneumoniae KPNIH24 | kph | 2 | Bacteria | Pseudomonadota | Gammaproteobacteria | Enterobacterales | Enterobacteriaceae | Klebsiella |
| Klebsiella pneumoniae subsp. pneumoniae KPNIH27 | kpz | 2 | Bacteria | Pseudomonadota | Gammaproteobacteria | Enterobacterales | Enterobacteriaceae | Klebsiella |
| Klebsiella pneumoniae subsp. pneumoniae KPNIH29 | kpv | 2 | Bacteria | Pseudomonadota | Gammaproteobacteria | Enterobacterales | Enterobacteriaceae | Klebsiella |
| Klebsiella pneumoniae subsp. pneumoniae KPNIH30 | kpw | 2 | Bacteria | Pseudomonadota | Gammaproteobacteria | Enterobacterales | Enterobacteriaceae | Klebsiella |
| Klebsiella pneumoniae subsp. pneumoniae KPNIH31 | kpy | 2 | Bacteria | Pseudomonadota | Gammaproteobacteria | Enterobacterales | Enterobacteriaceae | Klebsiella |
| Klebsiella pneumoniae subsp. pneumoniae KPNIH32 | kpg | 2 | Bacteria | Pseudomonadota | Gammaproteobacteria | Enterobacterales | Enterobacteriaceae | Klebsiella |
| Klebsiella pneumoniae subsp. pneumoniae KPR0928 | kpq | 2 | Bacteria | Pseudomonadota | Gammaproteobacteria | Enterobacterales | Enterobacteriaceae | Klebsiella |
| Klebsiella pneumoniae subsp. pneumoniae MGH 78578 (serotype K52) | kpn | 2 | Bacteria | Pseudomonadota | Gammaproteobacteria | Enterobacterales | Enterobacteriaceae | Klebsiella |
| Klebsiella pneumoniae subsp. pneumoniae NTUH-K2044 (serotype K1) | kpu | 2 | Bacteria | Pseudomonadota | Gammaproteobacteria | Enterobacterales | Enterobacteriaceae | Klebsiella |
| Klebsiella pneumoniae subsp. rhinoscleromatis SB3432 | kpr | 2 | Bacteria | Pseudomonadota | Gammaproteobacteria | Enterobacterales | Enterobacteriaceae | Klebsiella |
| Klebsiella quasipneumoniae | kqu | 2 | Bacteria | Pseudomonadota | Gammaproteobacteria | Enterobacterales | Enterobacteriaceae | Klebsiella |
| Klebsiella quasivariicola | kqv | 1 | Bacteria | Pseudomonadota | Gammaproteobacteria | Enterobacterales | Enterobacteriaceae | Klebsiella |
| Klebsiella variicola 342 | kpe | 2 | Bacteria | Pseudomonadota | Gammaproteobacteria | Enterobacterales | Enterobacteriaceae | Klebsiella |
| Klebsiella variicola At-22 | kva | 2 | Bacteria | Pseudomonadota | Gammaproteobacteria | Enterobacterales | Enterobacteriaceae | Klebsiella |
| Klebsiella variicola DSM 15968 | kvq | 2 | Bacteria | Pseudomonadota | Gammaproteobacteria | Enterobacterales | Enterobacteriaceae | Klebsiella |
| Klebsiella variicola DX120E | kvd | 2 | Bacteria | Pseudomonadota | Gammaproteobacteria | Enterobacterales | Enterobacteriaceae | Klebsiella |
| Klebsiella variicola KP5-1 | kpk | 2 | Bacteria | Pseudomonadota | Gammaproteobacteria | Enterobacterales | Enterobacteriaceae | Klebsiella |
| Streptomyces aurantiacus | sgm | 1 | Bacteria | Actinomycetota | Actinomycetes | Kitasatosporales | Streptomycetaceae | Streptomyces |
| Streptomyces griseofuscus | sgf | 1 | Bacteria | Actinomycetota | Actinomycetes | Kitasatosporales | Streptomycetaceae | Streptomyces |
| Streptomyces leeuwenhoekii | sle | 2 | Bacteria | Actinomycetota | Actinomycetes | Kitasatosporales | Streptomycetaceae | Streptomyces |
| Streptomyces lydicus A02 | sld | 1 | Bacteria | Actinomycetota | Actinomycetes | Kitasatosporales | Streptomycetaceae | Streptomyces |
| Streptomyces nigrescens | snig | 1 | Bacteria | Actinomycetota | Actinomycetes | Kitasatosporales | Streptomycetaceae | Streptomyces |
| Streptomyces noursei NK660 | salu | 1 | Bacteria | Actinomycetota | Actinomycetes | Kitasatosporales | Streptomycetaceae | Streptomyces |
| Streptomyces xanthophaeus | sxt | 1 | Bacteria | Actinomycetota | Actinomycetes | Kitasatosporales | Streptomycetaceae | Streptomyces |
| Agreia sp. COWG | agx | 1 | Bacteria | Actinomycetota | Actinomycetes | Micrococcales | Microbacteriaceae | Agreia |
| Arthrobacter sp. Hiyo4 | arty | 1 | Bacteria | Actinomycetota | Actinomycetes | Micrococcales | Micrococcaceae | Arthrobacter |
| Arthrobacter sp. Hiyo8 | arh | 1 | Bacteria | Actinomycetota | Actinomycetes | Micrococcales | Micrococcaceae | Arthrobacter |
| Arthrobacter sp. Rue61a | arr | 1 | Bacteria | Actinomycetota | Actinomycetes | Micrococcales | Micrococcaceae | Arthrobacter |
| Corynebacterium glutamicum ATCC 13032 (Bielefeld) | cgb | 1 | Bacteria | Actinomycetota | Actinomycetes | Mycobacteriales | Corynebacteriaceae | Corynebacterium |
| Corynebacterium marinum | cmq | 1 | Bacteria | Actinomycetota | Actinomycetes | Mycobacteriales | Corynebacteriaceae | Corynebacterium |
| Corynebacterium maris | cmd | 1 | Bacteria | Actinomycetota | Actinomycetes | Mycobacteriales | Corynebacteriaceae | Corynebacterium |
| Corynebacterium renale | crl | 1 | Bacteria | Actinomycetota | Actinomycetes | Mycobacteriales | Corynebacteriaceae | Corynebacterium |
| Mycobacteroides abscessus subsp. bolletii 50594 | mabb | 1 | Bacteria | Actinomycetota | Actinomycetes | Mycobacteriales | Mycobacteriaceae | Mycobacteroides |
| Mycolicibacillus koreensis | mkr | 1 | Bacteria | Actinomycetota | Actinomycetes | Mycobacteriales | Mycobacteriaceae | Mycolicibacillus |
| Tsukamurella pulmonis | tpul | 1 | Bacteria | Actinomycetota | Actinomycetes | Mycobacteriales | Tsukamurellaceae | Tsukamurella |
| Micropruina glycogenica | mgg | 1 | Bacteria | Actinomycetota | Actinomycetes | Propionibacteriales | Nocardioidaceae | Micropruina |
| Propionibacterium freudenreichii subsp. freudenreichii DSM 20271 | pfre | 2 | Bacteria | Actinomycetota | Actinomycetes | Propionibacteriales | Propionibacteriaceae | Propionibacterium |
| Amycolatopsis keratiniphila | aoi | 3 | Bacteria | Actinomycetota | Actinomycetes | Pseudonocardiales | Pseudonocardiaceae | Amycolatopsis |
| Amycolatopsis mediterranei RB | amz | 2 | Bacteria | Actinomycetota | Actinomycetes | Pseudonocardiales | Pseudonocardiaceae | Amycolatopsis |
| Amycolatopsis mediterranei S699 | amm | 2 | Bacteria | Actinomycetota | Actinomycetes | Pseudonocardiales | Pseudonocardiaceae | Amycolatopsis |
| Amycolatopsis mediterranei S699 | amn | 2 | Bacteria | Actinomycetota | Actinomycetes | Pseudonocardiales | Pseudonocardiaceae | Amycolatopsis |
| Amycolatopsis mediterranei U32 | amd | 2 | Bacteria | Actinomycetota | Actinomycetes | Pseudonocardiales | Pseudonocardiaceae | Amycolatopsis |
| Amycolatopsis methanolica | amq | 2 | Bacteria | Actinomycetota | Actinomycetes | Pseudonocardiales | Pseudonocardiaceae | Amycolatopsis |
| Amycolatopsis sp. CA-230715 | amyz | 1 | Bacteria | Actinomycetota | Actinomycetes | Pseudonocardiales | Pseudonocardiaceae | Amycolatopsis |
| Kutzneria sp. CA-103260 | kut | 3 | Bacteria | Actinomycetota | Actinomycetes | Pseudonocardiales | Pseudonocardiaceae | Kutzneria |
| Pseudonocardia autotrophica | paut | 2 | Bacteria | Actinomycetota | Actinomycetes | Pseudonocardiales | Pseudonocardiaceae | Pseudonocardia |
| Streptosporangium roseum | sro | 1 | Bacteria | Actinomycetota | Actinomycetes | Streptosporangiales | Streptosporangiaceae | Streptosporangium |
| Geobacillus kaustophilus | gka | 1 | Bacteria | Bacillota | Bacilli | Caryophanales | Anoxybacillaceae | Geobacillus |
| Alkalihalophilus pseudofirmus | bpf | 1 | Bacteria | Bacillota | Bacilli | Caryophanales | Bacillaceae | Alkalihalophilus |
| Bacillus amyloliquefaciens IT-45 | bami | 1 | Bacteria | Bacillota | Bacilli | Caryophanales | Bacillaceae | Bacillus |
| Bacillus amyloliquefaciens XH7 | bxh | 1 | Bacteria | Bacillota | Bacilli | Caryophanales | Bacillaceae | Bacillus |
| Bacillus sonorensis | bson | 1 | Bacteria | Bacillota | Bacilli | Caryophanales | Bacillaceae | Bacillus |
| Bacillus toyonensis biovar Thuringiensis | btm | 1 | Bacteria | Bacillota | Bacilli | Caryophanales | Bacillaceae | Bacillus |
| Bacillus velezensis CAU B946 | baq | 1 | Bacteria | Bacillota | Bacilli | Caryophanales | Bacillaceae | Bacillus |
| Bacillus velezensis YAU B9601-Y2 | bqy | 1 | Bacteria | Bacillota | Bacilli | Caryophanales | Bacillaceae | Bacillus |
| Halalkalibacterium halodurans | bha | 1 | Bacteria | Bacillota | Bacilli | Caryophanales | Bacillaceae | Halalkalibacterium |
| Oceanobacillus iheyensis | oih | 1 | Bacteria | Bacillota | Bacilli | Caryophanales | Bacillaceae | Oceanobacillus |
| Priestia aryabhattai | parh | 1 | Bacteria | Bacillota | Bacilli | Caryophanales | Bacillaceae | Priestia |
| Listeria monocytogenes ATCC 19117 (serotype 4d) | lmoa | 1 | Bacteria | Bacillota | Bacilli | Caryophanales | Listeriaceae | Listeria |
| Listeria monocytogenes L312 (serotype 4b) | lmol | 1 | Bacteria | Bacillota | Bacilli | Caryophanales | Listeriaceae | Listeria |
| Listeria monocytogenes L99 (serotype 4a) | lml | 1 | Bacteria | Bacillota | Bacilli | Caryophanales | Listeriaceae | Listeria |
| Listeria monocytogenes serotype 7 SLCC2482 | lmz | 1 | Bacteria | Bacillota | Bacilli | Caryophanales | Listeriaceae | Listeria |
| Listeria monocytogenes SLCC2372 (serotype 1/2c) | lmx | 1 | Bacteria | Bacillota | Bacilli | Caryophanales | Listeriaceae | Listeria |
| Listeria monocytogenes SLCC2376 (serotype 4c) | lmon | 1 | Bacteria | Bacillota | Bacilli | Caryophanales | Listeriaceae | Listeria |
| Listeria monocytogenes SLCC2378 (serotype 4e) | lmoo | 1 | Bacteria | Bacillota | Bacilli | Caryophanales | Listeriaceae | Listeria |
| Listeria monocytogenes SLCC2479 (serotype 3c) | lmoy | 1 | Bacteria | Bacillota | Bacilli | Caryophanales | Listeriaceae | Listeria |
| Listeria monocytogenes SLCC2540 (serotype 3b) | lmot | 1 | Bacteria | Bacillota | Bacilli | Caryophanales | Listeriaceae | Listeria |
| Listeria monocytogenes SLCC2755 (serotype 1/2b) | lmw | 1 | Bacteria | Bacillota | Bacilli | Caryophanales | Listeriaceae | Listeria |
| Listeria monocytogenes SLCC5850 (serotype 1/2a) | lmoc | 1 | Bacteria | Bacillota | Bacilli | Caryophanales | Listeriaceae | Listeria |
| Listeria monocytogenes SLCC7179 (serotype 3a) | lmos | 1 | Bacteria | Bacillota | Bacilli | Caryophanales | Listeriaceae | Listeria |
| Paenibacillus nuruki | pnk | 1 | Bacteria | Bacillota | Bacilli | Caryophanales | Paenibacillaceae | Paenibacillus |
| Paenibacillus sabinae | psab | 1 | Bacteria | Bacillota | Bacilli | Caryophanales | Paenibacillaceae | Paenibacillus |
| Macrococcoides caseolyticum | mcl | 1 | Bacteria | Bacillota | Bacilli | Caryophanales | Staphylococcaceae | Macrococcoides |
| Mammaliicoccus stepanovicii | sste | 1 | Bacteria | Bacillota | Bacilli | Caryophanales | Staphylococcaceae | Mammaliicoccus |
| Staphylococcus aureus Bmb9393 (MRSA) | saur | 1 | Bacteria | Bacillota | Bacilli | Caryophanales | Staphylococcaceae | Staphylococcus |
| Staphylococcus aureus subsp. aureus ECT-R 2 (MSSA) | suc | 1 | Bacteria | Bacillota | Bacilli | Caryophanales | Staphylococcaceae | Staphylococcus |
| Staphylococcus aureus subsp. aureus Mu3 (MRSA/hetero-VISA) | saw | 1 | Bacteria | Bacillota | Bacilli | Caryophanales | Staphylococcaceae | Staphylococcus |
| Staphylococcus aureus subsp. aureus Mu50 (MRSA/VISA) | sav | 1 | Bacteria | Bacillota | Bacilli | Caryophanales | Staphylococcaceae | Staphylococcus |
| Staphylococcus aureus subsp. aureus MW2 (CA-MRSA) | sam | 1 | Bacteria | Bacillota | Bacilli | Caryophanales | Staphylococcaceae | Staphylococcus |
| Staphylococcus aureus subsp. aureus N315 (MRSA/VSSA) | sau | 1 | Bacteria | Bacillota | Bacilli | Caryophanales | Staphylococcaceae | Staphylococcus |
| Staphylococcus aureus subsp. aureus Newman | sae | 1 | Bacteria | Bacillota | Bacilli | Caryophanales | Staphylococcaceae | Staphylococcus |
| Staphylococcus aureus subsp. aureus ST398 (MRSA) | sug | 1 | Bacteria | Bacillota | Bacilli | Caryophanales | Staphylococcaceae | Staphylococcus |
| Staphylococcus aureus subsp. aureus USA300_FPR3757 (CA-MRSA) | saa | 1 | Bacteria | Bacillota | Bacilli | Caryophanales | Staphylococcaceae | Staphylococcus |
| Staphylococcus aureus subsp. aureus USA300_TCH1516 (CA-MRSA) | sax | 1 | Bacteria | Bacillota | Bacilli | Caryophanales | Staphylococcaceae | Staphylococcus |
| Staphylococcus aureus subsp. aureus VC40 | suv | 1 | Bacteria | Bacillota | Bacilli | Caryophanales | Staphylococcaceae | Staphylococcus |
| Staphylococcus epidermidis ATCC 12228 | sep | 1 | Bacteria | Bacillota | Bacilli | Caryophanales | Staphylococcaceae | Staphylococcus |
| Staphylococcus haemolyticus JCSC1435 | sha | 1 | Bacteria | Bacillota | Bacilli | Caryophanales | Staphylococcaceae | Staphylococcus |
| Staphylococcus warneri | swa | 1 | Bacteria | Bacillota | Bacilli | Caryophanales | Staphylococcaceae | Staphylococcus |
| Enterococcus faecalis 62 | efl | 1 | Bacteria | Bacillota | Bacilli | Lactobacillales | Enterococcaceae | Enterococcus |
| Enterococcus faecalis D32 | efd | 1 | Bacteria | Bacillota | Bacilli | Lactobacillales | Enterococcaceae | Enterococcus |
| Weissella soli | wso | 2 | Bacteria | Bacillota | Bacilli | Lactobacillales | Lactobacillaceae | Weissella |
| Lactococcus lactis subsp. cremoris MG1363 | llm | 2 | Bacteria | Bacillota | Bacilli | Lactobacillales | Streptococcaceae | Lactococcus |
| Lactococcus lactis subsp. cremoris NZ9000 | lln | 2 | Bacteria | Bacillota | Bacilli | Lactobacillales | Streptococcaceae | Lactococcus |
| Lactococcus lactis subsp. cremoris UC509.9 | lli | 2 | Bacteria | Bacillota | Bacilli | Lactobacillales | Streptococcaceae | Lactococcus |
| Streptococcus australis | saup | 1 | Bacteria | Bacillota | Bacilli | Lactobacillales | Streptococcaceae | Streptococcus |
| Streptococcus cristatus | soi | 1 | Bacteria | Bacillota | Bacilli | Lactobacillales | Streptococcaceae | Streptococcus |
| Streptococcus ferus | sfer | 2 | Bacteria | Bacillota | Bacilli | Lactobacillales | Streptococcaceae | Streptococcus |
| Streptococcus gallolyticus subsp. gallolyticus ATCC 43143 | sgt | 1 | Bacteria | Bacillota | Bacilli | Lactobacillales | Streptococcaceae | Streptococcus |
| Streptococcus gallolyticus subsp. gallolyticus ATCC BAA-2069 | sgg | 1 | Bacteria | Bacillota | Bacilli | Lactobacillales | Streptococcaceae | Streptococcus |
| Streptococcus gallolyticus UCN34 | sga | 2 | Bacteria | Bacillota | Bacilli | Lactobacillales | Streptococcaceae | Streptococcus |
| Streptococcus gordonii | sgo | 1 | Bacteria | Bacillota | Bacilli | Lactobacillales | Streptococcaceae | Streptococcus |
| Streptococcus infantarius subsp. infantarius | sif | 2 | Bacteria | Bacillota | Bacilli | Lactobacillales | Streptococcaceae | Streptococcus |
| Streptococcus lutetiensis | slu | 2 | Bacteria | Bacillota | Bacilli | Lactobacillales | Streptococcaceae | Streptococcus |
| Streptococcus macedonicus | smn | 1 | Bacteria | Bacillota | Bacilli | Lactobacillales | Streptococcaceae | Streptococcus |
| Streptococcus mitis | smb | 1 | Bacteria | Bacillota | Bacilli | Lactobacillales | Streptococcaceae | Streptococcus |
| Streptococcus mutans GS-5 (serotype c) | smut | 2 | Bacteria | Bacillota | Bacilli | Lactobacillales | Streptococcaceae | Streptococcus |
| Streptococcus mutans LJ23 (serotype k) | smj | 2 | Bacteria | Bacillota | Bacilli | Lactobacillales | Streptococcaceae | Streptococcus |
| Streptococcus mutans NN2025 (serotype c) | smc | 2 | Bacteria | Bacillota | Bacilli | Lactobacillales | Streptococcaceae | Streptococcus |
| Streptococcus mutans UA159 (serotype c) | smu | 2 | Bacteria | Bacillota | Bacilli | Lactobacillales | Streptococcaceae | Streptococcus |
| Streptococcus mutans UA159-FR | smua | 2 | Bacteria | Bacillota | Bacilli | Lactobacillales | Streptococcaceae | Streptococcus |
| Streptococcus pasteurianus | stb | 1 | Bacteria | Bacillota | Bacilli | Lactobacillales | Streptococcaceae | Streptococcus |
| Streptococcus pneumoniae R6 (avirulent serotype 2) | spr | 1 | Bacteria | Bacillota | Bacilli | Lactobacillales | Streptococcaceae | Streptococcus |
| Streptococcus salivarius 57.I | stf | 1 | Bacteria | Bacillota | Bacilli | Lactobacillales | Streptococcaceae | Streptococcus |
| Streptococcus salivarius JIM8777 | stj | 2 | Bacteria | Bacillota | Bacilli | Lactobacillales | Streptococcaceae | Streptococcus |
| Streptococcus sanguinis | ssa | 1 | Bacteria | Bacillota | Bacilli | Lactobacillales | Streptococcaceae | Streptococcus |
| Streptococcus suis 05ZYH33 (serotype 2) | ssu | 1 | Bacteria | Bacillota | Bacilli | Lactobacillales | Streptococcaceae | Streptococcus |
| Streptococcus suis 98HAH33 (serotype 2) | ssv | 1 | Bacteria | Bacillota | Bacilli | Lactobacillales | Streptococcaceae | Streptococcus |
| Streptococcus suis A7 (serotype 2) | ssf | 1 | Bacteria | Bacillota | Bacilli | Lactobacillales | Streptococcaceae | Streptococcus |
| Streptococcus suis D12 (serotype 9) | ssk | 1 | Bacteria | Bacillota | Bacilli | Lactobacillales | Streptococcaceae | Streptococcus |
| Streptococcus suis D9 (serotype 7) | ssq | 1 | Bacteria | Bacillota | Bacilli | Lactobacillales | Streptococcaceae | Streptococcus |
| Streptococcus suis GZ1 (serotype 2) | ssw | 1 | Bacteria | Bacillota | Bacilli | Lactobacillales | Streptococcaceae | Streptococcus |
| Streptococcus suis JS14 (serotype 14) | sui | 1 | Bacteria | Bacillota | Bacilli | Lactobacillales | Streptococcaceae | Streptococcus |
| Streptococcus suis S735 (serotype 2) | sup | 1 | Bacteria | Bacillota | Bacilli | Lactobacillales | Streptococcaceae | Streptococcus |
| Streptococcus suis SC070731 (serotype 2) | ssus | 1 | Bacteria | Bacillota | Bacilli | Lactobacillales | Streptococcaceae | Streptococcus |
| Streptococcus suis SS12 (serotype 1/2) | suo | 1 | Bacteria | Bacillota | Bacilli | Lactobacillales | Streptococcaceae | Streptococcus |
| Streptococcus suis ST1 (serotype 1) | srp | 1 | Bacteria | Bacillota | Bacilli | Lactobacillales | Streptococcaceae | Streptococcus |
| Streptococcus suis ST3 (serotype 3) | sst | 1 | Bacteria | Bacillota | Bacilli | Lactobacillales | Streptococcaceae | Streptococcus |
| Streptococcus suis T15 | ssui | 1 | Bacteria | Bacillota | Bacilli | Lactobacillales | Streptococcaceae | Streptococcus |
| Streptococcus thermophilus CNRZ1066 | stc | 3 | Bacteria | Bacillota | Bacilli | Lactobacillales | Streptococcaceae | Streptococcus |
| Streptococcus thermophilus JIM 8232 | stu | 2 | Bacteria | Bacillota | Bacilli | Lactobacillales | Streptococcaceae | Streptococcus |
| Streptococcus thermophilus LMG 18311 | stl | 2 | Bacteria | Bacillota | Bacilli | Lactobacillales | Streptococcaceae | Streptococcus |
| Streptococcus thermophilus ND03 | stn | 1 | Bacteria | Bacillota | Bacilli | Lactobacillales | Streptococcaceae | Streptococcus |
| Streptococcus troglodytae | strg | 2 | Bacteria | Bacillota | Bacilli | Lactobacillales | Streptococcaceae | Streptococcus |
| Streptococcus urinalis | surn | 1 | Bacteria | Bacillota | Bacilli | Lactobacillales | Streptococcaceae | Streptococcus |
| Streptococcus vestibularis | svb | 1 | Bacteria | Bacillota | Bacilli | Lactobacillales | Streptococcaceae | Streptococcus |
| Streptococcus viridans | svf | 1 | Bacteria | Bacillota | Bacilli | Lactobacillales | Streptococcaceae | Streptococcus |
| Turicibacter sp. H121 | tur | 1 | Bacteria | Bacillota | Erysipelotrichia | Erysipelotrichales | Turicibacteraceae | Turicibacter |
| Megasphaera elsdenii | med | 1 | Bacteria | Bacillota | Negativicutes | Veillonellales | Veillonellaceae | Megasphaera |
| Petrimonas sp. IBARAKI | pet | 1 | Bacteria | Bacteroidota | Bacteroidia | Bacteroidales | Dysgonomonadaceae | Petrimonas |
| Porphyromonas somerae | psoe | 1 | Bacteria | Bacteroidota | Bacteroidia | Bacteroidales | Porphyromonadaceae | Porphyromonas |
| Hoylesella buccalis | pbuc | 1 | Bacteria | Bacteroidota | Bacteroidia | Bacteroidales | Prevotellaceae | Hoylesella |
| Prevotella bivia | pbiv | 1 | Bacteria | Bacteroidota | Bacteroidia | Bacteroidales | Prevotellaceae | Prevotella |
| Aquirufa nivalisilvae | psez | 2 | Bacteria | Bacteroidota | Cytophagia | Cytophagales | Flectobacillaceae | Aquirufa |
| Nibribacter ruber | nib | 1 | Bacteria | Bacteroidota | Cytophagia | Cytophagales | Hymenobacteraceae | Nibribacter |
| Flavobacteriaceae bacterium 3519-10 | fba | 1 | Bacteria | Bacteroidota | Flavobacteriia | Flavobacteriales | Flavobacteriaceae | Flavobacteriaceae |
| Faecalibacter bovis | fbo | 1 | Bacteria | Bacteroidota | Flavobacteriia | Flavobacteriales | Weeksellaceae | Faecalibacter |
| Marnyiella aurantia | cbau | 1 | Bacteria | Bacteroidota | Flavobacteriia | Flavobacteriales | Weeksellaceae | Marnyiella |
| Chryseobacterium gotjawalense | chrc | 1 | Bacteria | Bacteroidota | Flavobacteriia | Flavobacteriales | Weeksellaceae | Chryseobacterium |
| Chryseobacterium sp. 6424 | chrz | 1 | Bacteria | Bacteroidota | Flavobacteriia | Flavobacteriales | Weeksellaceae | Chryseobacterium |
| Chryseobacterium suipulveris | csup | 1 | Bacteria | Bacteroidota | Flavobacteriia | Flavobacteriales | Weeksellaceae | Chryseobacterium |
| Kaistella daneshvariae | kda | 1 | Bacteria | Bacteroidota | Flavobacteriia | Flavobacteriales | Weeksellaceae | Kaistella |
| Kaistella faecalis | cfae | 1 | Bacteria | Bacteroidota | Flavobacteriia | Flavobacteriales | Weeksellaceae | Kaistella |
| Sphingobacterium sp. ML3W | sht | 1 | Bacteria | Bacteroidota | Sphingobacteriia | Sphingobacteriales | Sphingobacteriaceae | Sphingobacterium |
| Campylobacter avium | cavi | 1 | Bacteria | Campylobacterota | Epsilonproteobacteria | Campylobacterales | Campylobacteraceae | Campylobacter |
| Campylobacter hepaticus | chw | 1 | Bacteria | Campylobacterota | Epsilonproteobacteria | Campylobacterales | Campylobacteraceae | Campylobacter |
| Campylobacter jejuni 32488 | cjz | 1 | Bacteria | Campylobacterota | Epsilonproteobacteria | Campylobacterales | Campylobacteraceae | Campylobacter |
| Campylobacter jejuni 4031 | cjx | 1 | Bacteria | Campylobacterota | Epsilonproteobacteria | Campylobacterales | Campylobacteraceae | Campylobacter |
| Campylobacter jejuni RM1221 | cjr | 1 | Bacteria | Campylobacterota | Epsilonproteobacteria | Campylobacterales | Campylobacteraceae | Campylobacter |
| Campylobacter jejuni subsp. doylei 269.97 | cjd | 1 | Bacteria | Campylobacterota | Epsilonproteobacteria | Campylobacterales | Campylobacteraceae | Campylobacter |
| Campylobacter jejuni subsp. jejuni 00-1597 | cjl | 1 | Bacteria | Campylobacterota | Epsilonproteobacteria | Campylobacterales | Campylobacteraceae | Campylobacter |
| Campylobacter jejuni subsp. jejuni 00-2425 | cjei | 1 | Bacteria | Campylobacterota | Epsilonproteobacteria | Campylobacterales | Campylobacteraceae | Campylobacter |
| Campylobacter jejuni subsp. jejuni 00-2426 | cjej | 1 | Bacteria | Campylobacterota | Epsilonproteobacteria | Campylobacterales | Campylobacteraceae | Campylobacter |
| Campylobacter jejuni subsp. jejuni 00-2538 | cjeu | 1 | Bacteria | Campylobacterota | Epsilonproteobacteria | Campylobacterales | Campylobacteraceae | Campylobacter |
| Campylobacter jejuni subsp. jejuni 00-2544 | cjen | 1 | Bacteria | Campylobacterota | Epsilonproteobacteria | Campylobacterales | Campylobacteraceae | Campylobacter |
| Campylobacter jejuni subsp. jejuni 00-6200 | cjw | 1 | Bacteria | Campylobacterota | Epsilonproteobacteria | Campylobacterales | Campylobacteraceae | Campylobacter |
| Campylobacter jejuni subsp. jejuni 35925B2 | cjq | 1 | Bacteria | Campylobacterota | Epsilonproteobacteria | Campylobacterales | Campylobacteraceae | Campylobacter |
| Campylobacter jejuni subsp. jejuni 81116 | cju | 1 | Bacteria | Campylobacterota | Epsilonproteobacteria | Campylobacterales | Campylobacteraceae | Campylobacter |
| Campylobacter jejuni subsp. jejuni 81-176 | cjj | 1 | Bacteria | Campylobacterota | Epsilonproteobacteria | Campylobacterales | Campylobacteraceae | Campylobacter |
| Campylobacter jejuni subsp. jejuni IA3902 | cji | 1 | Bacteria | Campylobacterota | Epsilonproteobacteria | Campylobacterales | Campylobacteraceae | Campylobacter |
| Campylobacter jejuni subsp. jejuni ICDCCJ07001 | cjn | 1 | Bacteria | Campylobacterota | Epsilonproteobacteria | Campylobacterales | Campylobacteraceae | Campylobacter |
| Campylobacter jejuni subsp. jejuni M1 | cjm | 1 | Bacteria | Campylobacterota | Epsilonproteobacteria | Campylobacterales | Campylobacteraceae | Campylobacter |
| Campylobacter jejuni subsp. jejuni MTVDSCj20 | cjv | 1 | Bacteria | Campylobacterota | Epsilonproteobacteria | Campylobacterales | Campylobacteraceae | Campylobacter |
| Campylobacter jejuni subsp. jejuni NCTC 11168 = ATCC 700819 | cje | 1 | Bacteria | Campylobacterota | Epsilonproteobacteria | Campylobacterales | Campylobacteraceae | Campylobacter |
| Campylobacter jejuni subsp. jejuni NCTC 11168-BN148 | cjb | 1 | Bacteria | Campylobacterota | Epsilonproteobacteria | Campylobacterales | Campylobacteraceae | Campylobacter |
| Campylobacter jejuni subsp. jejuni PT14 | cjp | 1 | Bacteria | Campylobacterota | Epsilonproteobacteria | Campylobacterales | Campylobacteraceae | Campylobacter |
| Campylobacter jejuni subsp. jejuni R14 | cjer | 1 | Bacteria | Campylobacterota | Epsilonproteobacteria | Campylobacterales | Campylobacteraceae | Campylobacter |
| Campylobacter jejuni subsp. jejuni S3 | cjs | 1 | Bacteria | Campylobacterota | Epsilonproteobacteria | Campylobacterales | Campylobacteraceae | Campylobacter |
| Campylobacter jejuni subsp. jejuni YH001 | cjy | 1 | Bacteria | Campylobacterota | Epsilonproteobacteria | Campylobacterales | Campylobacteraceae | Campylobacter |
| Helicobacter ailurogastricus | hail | 1 | Bacteria | Campylobacterota | Epsilonproteobacteria | Campylobacterales | Helicobacteraceae | Helicobacter |
| Helicobacter felis | hfe | 1 | Bacteria | Campylobacterota | Epsilonproteobacteria | Campylobacterales | Helicobacteraceae | Helicobacter |
| Helicobacter heilmannii | hhm | 1 | Bacteria | Campylobacterota | Epsilonproteobacteria | Campylobacterales | Helicobacteraceae | Helicobacter |
| Cyanobium sp. NIES-981 | cyi | 1 | Bacteria | Cyanobacteriota | Cyanophyceae | Synechococcales | Prochlorococcaceae | Cyanobium |
| Malacoplasma iowae | miw | 1 | Bacteria | Mycoplasmatota |  | Mycoplasmoidales | Mycoplasmoidaceae | Malacoplasma |
| Malacoplasma penetrans | mpe | 1 | Bacteria | Mycoplasmatota |  | Mycoplasmoidales | Mycoplasmoidaceae | Malacoplasma |
| Anaeromyxobacter oryzae | aory | 1 | Bacteria | Myxococcota | Myxococcia | Myxococcales | Cystobacterineae | Anaeromyxobacter |
| Anaeromyxobacter paludicola | apau | 1 | Bacteria | Myxococcota | Myxococcia | Myxococcales | Cystobacterineae | Anaeromyxobacter |
| Nannocystis punicea | nann | 1 | Bacteria | Myxococcota | Polyangia | Nannocystales | Nannocystaceae | Nannocystis |
| Asaia bogorensis | abg | 1 | Bacteria | Pseudomonadota | Alphaproteobacteria | Acetobacterales | Acetobacteraceae | Asaia |
| Roseomonas marmotae | rmt | 1 | Bacteria | Pseudomonadota | Alphaproteobacteria | Acetobacterales | Roseomonadaceae | Roseomonas |
| Allobosea sp. Tri-49 | boi | 1 | Bacteria | Pseudomonadota | Alphaproteobacteria | Hyphomicrobiales | Alloboseaceae | Allobosea |
| Antarcticirhabdus aurantiaca | jav | 1 | Bacteria | Pseudomonadota | Alphaproteobacteria | Hyphomicrobiales | Aurantimonadaceae | Antarcticirhabdus |
| Martelella endophytica | mey | 1 | Bacteria | Pseudomonadota | Alphaproteobacteria | Hyphomicrobiales | Aurantimonadaceae | Martelella |
| Ditibartonella apihabitans | bapa | 4 | Bacteria | Pseudomonadota | Alphaproteobacteria | Hyphomicrobiales | Bartonellaceae | Ditibartonella |
| Ditibartonella choladocola | bapi | 4 | Bacteria | Pseudomonadota | Alphaproteobacteria | Hyphomicrobiales | Bartonellaceae | Ditibartonella |
| Pseudochrobactrum algeriensis | pscq | 3 | Bacteria | Pseudomonadota | Alphaproteobacteria | Hyphomicrobiales | Brucellaceae | Pseudochrobactrum |
| Brucella anthropi ATCC 49188 | oan | 1 | Bacteria | Pseudomonadota | Alphaproteobacteria | Hyphomicrobiales | Brucellaceae | Brucella |
| Brucella anthropi OAB | oah | 1 | Bacteria | Pseudomonadota | Alphaproteobacteria | Hyphomicrobiales | Brucellaceae | Brucella |
| Brucella inopinata | bio | 1 | Bacteria | Pseudomonadota | Alphaproteobacteria | Hyphomicrobiales | Brucellaceae | Brucella |
| Brucella intermedia | oin | 2 | Bacteria | Pseudomonadota | Alphaproteobacteria | Hyphomicrobiales | Brucellaceae | Brucella |
| Brucella melitensis bv. 1 16M | bmel | 1 | Bacteria | Pseudomonadota | Alphaproteobacteria | Hyphomicrobiales | Brucellaceae | Brucella |
| Brucella pseudogrignonensis | ops | 2 | Bacteria | Pseudomonadota | Alphaproteobacteria | Hyphomicrobiales | Brucellaceae | Brucella |
| Brucella sp. 09RB8471 | brj | 1 | Bacteria | Pseudomonadota | Alphaproteobacteria | Hyphomicrobiales | Brucellaceae | Brucella |
| Brucella sp. JSBI001 | bruc | 1 | Bacteria | Pseudomonadota | Alphaproteobacteria | Hyphomicrobiales | Brucellaceae | Brucella |
| Ochrobactrum quorumnocens | och | 2 | Bacteria | Pseudomonadota | Alphaproteobacteria | Hyphomicrobiales | Brucellaceae | Ochrobactrum |
| Ochrobactrum sp. BTU1 | occ | 2 | Bacteria | Pseudomonadota | Alphaproteobacteria | Hyphomicrobiales | Brucellaceae | Ochrobactrum |
| Ochrobactrum sp. MT180101 | ocr | 1 | Bacteria | Pseudomonadota | Alphaproteobacteria | Hyphomicrobiales | Brucellaceae | Ochrobactrum |
| Devosia neptuniae | dnp | 1 | Bacteria | Pseudomonadota | Alphaproteobacteria | Hyphomicrobiales | Devosiaceae | Devosia |
| Maritalea myrionectae | mmyr | 2 | Bacteria | Pseudomonadota | Alphaproteobacteria | Hyphomicrobiales | Devosiaceae | Maritalea |
| Candidatus Tokpelaia hoelldoblerii | thd | 2 | Bacteria | Pseudomonadota | Alphaproteobacteria | Hyphomicrobiales | Hyphomicrobiales incertae sedis | Candidatus Tokpelaia |
| Nordella sp. HKS 07 | noh | 1 | Bacteria | Pseudomonadota | Alphaproteobacteria | Hyphomicrobiales | Hyphomicrobiales incertae sedis | Nordella |
| Methylobacterium currus | mee | 1 | Bacteria | Pseudomonadota | Alphaproteobacteria | Hyphomicrobiales | Methylobacteriaceae | Methylobacterium |
| Methylobacterium indicum | mind | 2 | Bacteria | Pseudomonadota | Alphaproteobacteria | Hyphomicrobiales | Methylobacteriaceae | Methylobacterium |
| Microvirga lotononidis | mld | 1 | Bacteria | Pseudomonadota | Alphaproteobacteria | Hyphomicrobiales | Methylobacteriaceae | Microvirga |
| Microvirga ossetica | moc | 1 | Bacteria | Pseudomonadota | Alphaproteobacteria | Hyphomicrobiales | Methylobacteriaceae | Microvirga |
| Bradyrhizobium cosmicum | brs | 1 | Bacteria | Pseudomonadota | Alphaproteobacteria | Hyphomicrobiales | Nitrobacteraceae | Bradyrhizobium |
| Bradyrhizobium diazoefficiens USDA 110 | bja | 1 | Bacteria | Pseudomonadota | Alphaproteobacteria | Hyphomicrobiales | Nitrobacteraceae | Bradyrhizobium |
| Bradyrhizobium elkanii | bel | 3 | Bacteria | Pseudomonadota | Alphaproteobacteria | Hyphomicrobiales | Nitrobacteraceae | Bradyrhizobium |
| Bradyrhizobium japonicum USDA 6 | bju | 1 | Bacteria | Pseudomonadota | Alphaproteobacteria | Hyphomicrobiales | Nitrobacteraceae | Bradyrhizobium |
| Bradyrhizobium oligotrophicum | aol | 1 | Bacteria | Pseudomonadota | Alphaproteobacteria | Hyphomicrobiales | Nitrobacteraceae | Bradyrhizobium |
| Bradyrhizobium sp. BF49 | brad | 1 | Bacteria | Pseudomonadota | Alphaproteobacteria | Hyphomicrobiales | Nitrobacteraceae | Bradyrhizobium |
| Bradyrhizobium sp. ORS 278 | bra | 5 | Bacteria | Pseudomonadota | Alphaproteobacteria | Hyphomicrobiales | Nitrobacteraceae | Bradyrhizobium |
| Bradyrhizobium sp. ORS 285 | bro | 5 | Bacteria | Pseudomonadota | Alphaproteobacteria | Hyphomicrobiales | Nitrobacteraceae | Bradyrhizobium |
| Bradyrhizobium vignae | bvz | 3 | Bacteria | Pseudomonadota | Alphaproteobacteria | Hyphomicrobiales | Nitrobacteraceae | Bradyrhizobium |
| Phreatobacter stygius | pstg | 1 | Bacteria | Pseudomonadota | Alphaproteobacteria | Hyphomicrobiales | Phreatobacteraceae | Phreatobacter |
| Aminobacter aminovorans | aak | 1 | Bacteria | Pseudomonadota | Alphaproteobacteria | Hyphomicrobiales | Phyllobacteriaceae | Aminobacter |
| Aminobacter niigataensis | anj | 3 | Bacteria | Pseudomonadota | Alphaproteobacteria | Hyphomicrobiales | Phyllobacteriaceae | Aminobacter |
| Mesorhizobium japonicum MAFF 303099 | mlo | 2 | Bacteria | Pseudomonadota | Alphaproteobacteria | Hyphomicrobiales | Phyllobacteriaceae | Mesorhizobium |
| Mesorhizobium sp. 8 | mesr | 1 | Bacteria | Pseudomonadota | Alphaproteobacteria | Hyphomicrobiales | Phyllobacteriaceae | Mesorhizobium |
| Mesorhizobium sp. Pch-S | mesp | 1 | Bacteria | Pseudomonadota | Alphaproteobacteria | Hyphomicrobiales | Phyllobacteriaceae | Mesorhizobium |
| Phyllobacterium sp. 628 | phyl | 1 | Bacteria | Pseudomonadota | Alphaproteobacteria | Hyphomicrobiales | Phyllobacteriaceae | Phyllobacterium |
| Phyllobacterium zundukense | pht | 1 | Bacteria | Pseudomonadota | Alphaproteobacteria | Hyphomicrobiales | Phyllobacteriaceae | Phyllobacterium |
| Arminella rhododendri | rrho | 1 | Bacteria | Pseudomonadota | Alphaproteobacteria | Hyphomicrobiales | Rhizobiaceae | Arminella |
| Arminella tumorigenes | rtu | 1 | Bacteria | Pseudomonadota | Alphaproteobacteria | Hyphomicrobiales | Rhizobiaceae | Arminella |
| Gillisella vitis VAR03-1 | avf | 3 | Bacteria | Pseudomonadota | Alphaproteobacteria | Hyphomicrobiales | Rhizobiaceae | Gillisella |
| Gillisella vitis VAT03-9 | avv | 4 | Bacteria | Pseudomonadota | Alphaproteobacteria | Hyphomicrobiales | Rhizobiaceae | Gillisella |
| Martinezella jaguaris | rjg | 2 | Bacteria | Pseudomonadota | Alphaproteobacteria | Hyphomicrobiales | Rhizobiaceae | Martinezella |
| Martinezella lusitana | rls | 2 | Bacteria | Pseudomonadota | Alphaproteobacteria | Hyphomicrobiales | Rhizobiaceae | Martinezella |
| Martinezella rhizogenes K599 | aro | 2 | Bacteria | Pseudomonadota | Alphaproteobacteria | Hyphomicrobiales | Rhizobiaceae | Martinezella |
| Peteryoungia desertarenae | pdes | 1 | Bacteria | Pseudomonadota | Alphaproteobacteria | Hyphomicrobiales | Rhizobiaceae | Peteryoungia |
| Rhizobium acidisoli | rad | 1 | Bacteria | Pseudomonadota | Alphaproteobacteria | Hyphomicrobiales | Rhizobiaceae | Rhizobium |
| Rhizobium anhuiense bv. trifolii | raw | 2 | Bacteria | Pseudomonadota | Alphaproteobacteria | Hyphomicrobiales | Rhizobiaceae | Rhizobium |
| Rhizobium bangladeshense | rban | 1 | Bacteria | Pseudomonadota | Alphaproteobacteria | Hyphomicrobiales | Rhizobiaceae | Rhizobium |
| Rhizobium brockwellii | rbw | 2 | Bacteria | Pseudomonadota | Alphaproteobacteria | Hyphomicrobiales | Rhizobiaceae | Rhizobium |
| Rhizobium esperanzae | rez | 3 | Bacteria | Pseudomonadota | Alphaproteobacteria | Hyphomicrobiales | Rhizobiaceae | Rhizobium |
| Rhizobium etli bv. mimosae IE4771 | rei | 4 | Bacteria | Pseudomonadota | Alphaproteobacteria | Hyphomicrobiales | Rhizobiaceae | Rhizobium |
| Rhizobium etli bv. mimosae Mim1 | rel | 3 | Bacteria | Pseudomonadota | Alphaproteobacteria | Hyphomicrobiales | Rhizobiaceae | Rhizobium |
| Rhizobium etli bv. phaseoli IE4803 | rep | 4 | Bacteria | Pseudomonadota | Alphaproteobacteria | Hyphomicrobiales | Rhizobiaceae | Rhizobium |
| Rhizobium etli CFN 42 | ret | 2 | Bacteria | Pseudomonadota | Alphaproteobacteria | Hyphomicrobiales | Rhizobiaceae | Rhizobium |
| Rhizobium etli CIAT 652 | rec | 2 | Bacteria | Pseudomonadota | Alphaproteobacteria | Hyphomicrobiales | Rhizobiaceae | Rhizobium |
| Rhizobium favelukesii | rhl | 3 | Bacteria | Pseudomonadota | Alphaproteobacteria | Hyphomicrobiales | Rhizobiaceae | Rhizobium |
| Rhizobium gallicum bv. gallicum | rga | 5 | Bacteria | Pseudomonadota | Alphaproteobacteria | Hyphomicrobiales | Rhizobiaceae | Rhizobium |
| Rhizobium grahamii | rgr | 1 | Bacteria | Pseudomonadota | Alphaproteobacteria | Hyphomicrobiales | Rhizobiaceae | Rhizobium |
| Rhizobium hidalgonense | rhid | 1 | Bacteria | Pseudomonadota | Alphaproteobacteria | Hyphomicrobiales | Rhizobiaceae | Rhizobium |
| Rhizobium indicum | rii | 2 | Bacteria | Pseudomonadota | Alphaproteobacteria | Hyphomicrobiales | Rhizobiaceae | Rhizobium |
| Rhizobium johnstonii | rle | 2 | Bacteria | Pseudomonadota | Alphaproteobacteria | Hyphomicrobiales | Rhizobiaceae | Rhizobium |
| Rhizobium laguerreae | rlw | 2 | Bacteria | Pseudomonadota | Alphaproteobacteria | Hyphomicrobiales | Rhizobiaceae | Rhizobium |
| Rhizobium leguminosarum bv. trifolii CB782 | rlu | 1 | Bacteria | Pseudomonadota | Alphaproteobacteria | Hyphomicrobiales | Rhizobiaceae | Rhizobium |
| Rhizobium leguminosarum bv. trifolii WSM1325 | rlg | 1 | Bacteria | Pseudomonadota | Alphaproteobacteria | Hyphomicrobiales | Rhizobiaceae | Rhizobium |
| Rhizobium leguminosarum bv. trifolii WSM1689 | rlb | 1 | Bacteria | Pseudomonadota | Alphaproteobacteria | Hyphomicrobiales | Rhizobiaceae | Rhizobium |
| Rhizobium leguminosarum bv. trifolii WSM2304 | rlt | 4 | Bacteria | Pseudomonadota | Alphaproteobacteria | Hyphomicrobiales | Rhizobiaceae | Rhizobium |
| Rhizobium lentis | rln | 1 | Bacteria | Pseudomonadota | Alphaproteobacteria | Hyphomicrobiales | Rhizobiaceae | Rhizobium |
| Rhizobium oryzihabitans | roy | 1 | Bacteria | Pseudomonadota | Alphaproteobacteria | Hyphomicrobiales | Rhizobiaceae | Rhizobium |
| Rhizobium phaseoli | rpha | 4 | Bacteria | Pseudomonadota | Alphaproteobacteria | Hyphomicrobiales | Rhizobiaceae | Rhizobium |
| Rhizobium rhizogenes K84 | ara | 5 | Bacteria | Pseudomonadota | Alphaproteobacteria | Hyphomicrobiales | Rhizobiaceae | Rhizobium |
| Rhizobium rosettiformans | rros | 1 | Bacteria | Pseudomonadota | Alphaproteobacteria | Hyphomicrobiales | Rhizobiaceae | Rhizobium |
| Rhizobium ruizarguesonis | rrg | 2 | Bacteria | Pseudomonadota | Alphaproteobacteria | Hyphomicrobiales | Rhizobiaceae | Rhizobium |
| Rhizobium sp. 11515TR | rhr | 2 | Bacteria | Pseudomonadota | Alphaproteobacteria | Hyphomicrobiales | Rhizobiaceae | Rhizobium |
| Rhizobium sp. AB2/73 | rhib | 2 | Bacteria | Pseudomonadota | Alphaproteobacteria | Hyphomicrobiales | Rhizobiaceae | Rhizobium |
| Rhizobium sp. Kim5 | rhk | 4 | Bacteria | Pseudomonadota | Alphaproteobacteria | Hyphomicrobiales | Rhizobiaceae | Rhizobium |
| Rhizobium sp. N1341 | rhn | 3 | Bacteria | Pseudomonadota | Alphaproteobacteria | Hyphomicrobiales | Rhizobiaceae | Rhizobium |
| Rhizobium sp. N731 | rhx | 4 | Bacteria | Pseudomonadota | Alphaproteobacteria | Hyphomicrobiales | Rhizobiaceae | Rhizobium |
| Rhizobium sp. S41 | rhv | 2 | Bacteria | Pseudomonadota | Alphaproteobacteria | Hyphomicrobiales | Rhizobiaceae | Rhizobium |
| Rhizobium sullae | rsul | 1 | Bacteria | Pseudomonadota | Alphaproteobacteria | Hyphomicrobiales | Rhizobiaceae | Rhizobium |
| Rhizobium tropici | rtr | 2 | Bacteria | Pseudomonadota | Alphaproteobacteria | Hyphomicrobiales | Rhizobiaceae | Rhizobium |
| Shinella oryzae | soj | 1 | Bacteria | Pseudomonadota | Alphaproteobacteria | Hyphomicrobiales | Rhizobiaceae | Shinella |
| Shinella sp. HZN7 | shz | 1 | Bacteria | Pseudomonadota | Alphaproteobacteria | Hyphomicrobiales | Rhizobiaceae | Shinella |
| Shinella sumterensis | ssum | 1 | Bacteria | Pseudomonadota | Alphaproteobacteria | Hyphomicrobiales | Rhizobiaceae | Shinella |
| Shinella zoogloeoides | szo | 1 | Bacteria | Pseudomonadota | Alphaproteobacteria | Hyphomicrobiales | Rhizobiaceae | Shinella |
| Terrirhizobium terrae | asit | 1 | Bacteria | Pseudomonadota | Alphaproteobacteria | Hyphomicrobiales | Rhizobiaceae | Terrirhizobium |
| Pseudorhizobium banfieldiae | rht | 3 | Bacteria | Pseudomonadota | Alphaproteobacteria | Hyphomicrobiales | Rhizobiaceae | Pseudorhizobium |
| Pseudorhizobium flavum | rfv | 1 | Bacteria | Pseudomonadota | Alphaproteobacteria | Hyphomicrobiales | Rhizobiaceae | Pseudorhizobium |
| Sinorhizobium alkalisoli | eak | 1 | Bacteria | Pseudomonadota | Alphaproteobacteria | Hyphomicrobiales | Rhizobiaceae | Sinorhizobium |
| Sinorhizobium americanum | same | 3 | Bacteria | Pseudomonadota | Alphaproteobacteria | Hyphomicrobiales | Rhizobiaceae | Sinorhizobium |
| Sinorhizobium fredii HH103 | sfh | 1 | Bacteria | Pseudomonadota | Alphaproteobacteria | Hyphomicrobiales | Rhizobiaceae | Sinorhizobium |
| Sinorhizobium fredii NGR234 | rhi | 1 | Bacteria | Pseudomonadota | Alphaproteobacteria | Hyphomicrobiales | Rhizobiaceae | Sinorhizobium |
| Sinorhizobium fredii USDA 257 | sfd | 2 | Bacteria | Pseudomonadota | Alphaproteobacteria | Hyphomicrobiales | Rhizobiaceae | Sinorhizobium |
| Sinorhizobium garamanticum | egi | 1 | Bacteria | Pseudomonadota | Alphaproteobacteria | Hyphomicrobiales | Rhizobiaceae | Sinorhizobium |
| Sinorhizobium kummerowiae | skm | 1 | Bacteria | Pseudomonadota | Alphaproteobacteria | Hyphomicrobiales | Rhizobiaceae | Sinorhizobium |
| Sinorhizobium medicae | smd | 1 | Bacteria | Pseudomonadota | Alphaproteobacteria | Hyphomicrobiales | Rhizobiaceae | Sinorhizobium |
| Sinorhizobium meliloti 1021 | sme | 3 | Bacteria | Pseudomonadota | Alphaproteobacteria | Hyphomicrobiales | Rhizobiaceae | Sinorhizobium |
| Sinorhizobium meliloti 2011 | smel | 3 | Bacteria | Pseudomonadota | Alphaproteobacteria | Hyphomicrobiales | Rhizobiaceae | Sinorhizobium |
| Sinorhizobium meliloti AK83 | smk | 1 | Bacteria | Pseudomonadota | Alphaproteobacteria | Hyphomicrobiales | Rhizobiaceae | Sinorhizobium |
| Sinorhizobium meliloti BL225C | smq | 1 | Bacteria | Pseudomonadota | Alphaproteobacteria | Hyphomicrobiales | Rhizobiaceae | Sinorhizobium |
| Sinorhizobium meliloti GR4 | smeg | 1 | Bacteria | Pseudomonadota | Alphaproteobacteria | Hyphomicrobiales | Rhizobiaceae | Sinorhizobium |
| Sinorhizobium meliloti Rm41 | smi | 3 | Bacteria | Pseudomonadota | Alphaproteobacteria | Hyphomicrobiales | Rhizobiaceae | Sinorhizobium |
| Sinorhizobium meliloti RMO17 | smer | 1 | Bacteria | Pseudomonadota | Alphaproteobacteria | Hyphomicrobiales | Rhizobiaceae | Sinorhizobium |
| Sinorhizobium meliloti SM11 | smx | 2 | Bacteria | Pseudomonadota | Alphaproteobacteria | Hyphomicrobiales | Rhizobiaceae | Sinorhizobium |
| Sinorhizobium mexicanum | emx | 1 | Bacteria | Pseudomonadota | Alphaproteobacteria | Hyphomicrobiales | Rhizobiaceae | Sinorhizobium |
| Sinorhizobium numidicum | enu | 1 | Bacteria | Pseudomonadota | Alphaproteobacteria | Hyphomicrobiales | Rhizobiaceae | Sinorhizobium |
| Sinorhizobium sojae | esj | 1 | Bacteria | Pseudomonadota | Alphaproteobacteria | Hyphomicrobiales | Rhizobiaceae | Sinorhizobium |
| Sinorhizobium sp. CCBAU 05631 | sino | 1 | Bacteria | Pseudomonadota | Alphaproteobacteria | Hyphomicrobiales | Rhizobiaceae | Sinorhizobium |
| Sinorhizobium sp. RAC02 | six | 2 | Bacteria | Pseudomonadota | Alphaproteobacteria | Hyphomicrobiales | Rhizobiaceae | Sinorhizobium |
| Sinorhizobium terangae | steg | 1 | Bacteria | Pseudomonadota | Alphaproteobacteria | Hyphomicrobiales | Rhizobiaceae | Sinorhizobium |
| Agrobacterium cucumeris | acuc | 2 | Bacteria | Pseudomonadota | Alphaproteobacteria | Hyphomicrobiales | Rhizobiaceae | Agrobacterium |
| Agrobacterium fabrum | atu | 4 | Bacteria | Pseudomonadota | Alphaproteobacteria | Hyphomicrobiales | Rhizobiaceae | Agrobacterium |
| Agrobacterium larrymoorei | alf | 3 | Bacteria | Pseudomonadota | Alphaproteobacteria | Hyphomicrobiales | Rhizobiaceae | Agrobacterium |
| Agrobacterium leguminum | aleg | 2 | Bacteria | Pseudomonadota | Alphaproteobacteria | Hyphomicrobiales | Rhizobiaceae | Agrobacterium |
| Agrobacterium pusense CFBP5875 | rpus | 2 | Bacteria | Pseudomonadota | Alphaproteobacteria | Hyphomicrobiales | Rhizobiaceae | Agrobacterium |
| Agrobacterium pusense IRBG74 | rir | 7 | Bacteria | Pseudomonadota | Alphaproteobacteria | Hyphomicrobiales | Rhizobiaceae | Agrobacterium |
| Agrobacterium rubi | arui | 2 | Bacteria | Pseudomonadota | Alphaproteobacteria | Hyphomicrobiales | Rhizobiaceae | Agrobacterium |
| Agrobacterium salinitolerans | asal | 2 | Bacteria | Pseudomonadota | Alphaproteobacteria | Hyphomicrobiales | Rhizobiaceae | Agrobacterium |
| Agrobacterium sp. 33MFTa1.1 | agt | 2 | Bacteria | Pseudomonadota | Alphaproteobacteria | Hyphomicrobiales | Rhizobiaceae | Agrobacterium |
| Agrobacterium sp. CGMCC 11546 | agrc | 2 | Bacteria | Pseudomonadota | Alphaproteobacteria | Hyphomicrobiales | Rhizobiaceae | Agrobacterium |
| Agrobacterium sp. RAC06 | agc | 1 | Bacteria | Pseudomonadota | Alphaproteobacteria | Hyphomicrobiales | Rhizobiaceae | Agrobacterium |
| Agrobacterium tumefaciens Ach5 | atf | 4 | Bacteria | Pseudomonadota | Alphaproteobacteria | Hyphomicrobiales | Rhizobiaceae | Agrobacterium |
| Agrobacterium tumefaciens H13-3 | agr | 5 | Bacteria | Pseudomonadota | Alphaproteobacteria | Hyphomicrobiales | Rhizobiaceae | Agrobacterium |
| Agrobacterium tumefaciens S33 | ata | 2 | Bacteria | Pseudomonadota | Alphaproteobacteria | Hyphomicrobiales | Rhizobiaceae | Agrobacterium |
| Agrobacterium vaccinii | avq | 2 | Bacteria | Pseudomonadota | Alphaproteobacteria | Hyphomicrobiales | Rhizobiaceae | Agrobacterium |
| Allorhizobium ampelinum | avi | 4 | Bacteria | Pseudomonadota | Alphaproteobacteria | Hyphomicrobiales | Rhizobiaceae | Allorhizobium |
| Neorhizobium galegae bv. officinalis bv. officinalis HAMBI 1141 | ngl | 5 | Bacteria | Pseudomonadota | Alphaproteobacteria | Hyphomicrobiales | Rhizobiaceae | Neorhizobium |
| Neorhizobium galegae bv. orientalis HAMBI 540 | ngg | 5 | Bacteria | Pseudomonadota | Alphaproteobacteria | Hyphomicrobiales | Rhizobiaceae | Neorhizobium |
| Neorhizobium petrolearium | npm | 1 | Bacteria | Pseudomonadota | Alphaproteobacteria | Hyphomicrobiales | Rhizobiaceae | Neorhizobium |
| Neorhizobium sp. NCHU2750 | nen | 5 | Bacteria | Pseudomonadota | Alphaproteobacteria | Hyphomicrobiales | Rhizobiaceae | Neorhizobium |
| Neorhizobium sp. SOG26 | neo | 2 | Bacteria | Pseudomonadota | Alphaproteobacteria | Hyphomicrobiales | Rhizobiaceae | Neorhizobium |
| Ensifer adhaerens Casida A | eah | 2 | Bacteria | Pseudomonadota | Alphaproteobacteria | Hyphomicrobiales | Rhizobiaceae | Ensifer |
| Ensifer adhaerens OV14 | ead | 4 | Bacteria | Pseudomonadota | Alphaproteobacteria | Hyphomicrobiales | Rhizobiaceae | Ensifer |
| Ensifer canadensis | ecaa | 4 | Bacteria | Pseudomonadota | Alphaproteobacteria | Hyphomicrobiales | Rhizobiaceae | Ensifer |
| Ensifer sp. PDNC004 | enp | 3 | Bacteria | Pseudomonadota | Alphaproteobacteria | Hyphomicrobiales | Rhizobiaceae | Ensifer |
| Labrenzia sp. CP4 | lap | 1 | Bacteria | Pseudomonadota | Alphaproteobacteria | Hyphomicrobiales | Stappiaceae | Labrenzia |
| Labrenzia sp. PHM005 | labp | 1 | Bacteria | Pseudomonadota | Alphaproteobacteria | Hyphomicrobiales | Stappiaceae | Labrenzia |
| Labrenzia sp. THAF35 | labt | 1 | Bacteria | Pseudomonadota | Alphaproteobacteria | Hyphomicrobiales | Stappiaceae | Labrenzia |
| Labrenzia sp. VG12 | labr | 1 | Bacteria | Pseudomonadota | Alphaproteobacteria | Hyphomicrobiales | Stappiaceae | Labrenzia |
| Pannonibacter phragmitetus | pphr | 1 | Bacteria | Pseudomonadota | Alphaproteobacteria | Hyphomicrobiales | Stappiaceae | Pannonibacter |
| Roseibium algicola | lagg | 1 | Bacteria | Pseudomonadota | Alphaproteobacteria | Hyphomicrobiales | Stappiaceae | Roseibium |
| Ancylobacter polymorphus | apol | 1 | Bacteria | Pseudomonadota | Alphaproteobacteria | Hyphomicrobiales | Xanthobacteraceae | Ancylobacter |
| Labrys sp. KNU-23 | lne | 1 | Bacteria | Pseudomonadota | Alphaproteobacteria | Hyphomicrobiales | Xanthobacteraceae | Labrys |
| Frigidibacter mobilis | daa | 1 | Bacteria | Pseudomonadota | Alphaproteobacteria | Rhodobacterales | Paracoccaceae | Frigidibacter |
| Haematobacter massiliensis | hml | 1 | Bacteria | Pseudomonadota | Alphaproteobacteria | Rhodobacterales | Paracoccaceae | Haematobacter |
| Paracoccus aminophilus | pami | 4 | Bacteria | Pseudomonadota | Alphaproteobacteria | Rhodobacterales | Paracoccaceae | Paracoccus |
| Paracoccus aminovorans | pamn | 1 | Bacteria | Pseudomonadota | Alphaproteobacteria | Rhodobacterales | Paracoccaceae | Paracoccus |
| Paracoccus kondratievae | pkd | 1 | Bacteria | Pseudomonadota | Alphaproteobacteria | Rhodobacterales | Paracoccaceae | Paracoccus |
| Paracoccus marcusii | pmau | 3 | Bacteria | Pseudomonadota | Alphaproteobacteria | Rhodobacterales | Paracoccaceae | Paracoccus |
| Paracoccus pantotrophus | ppan | 1 | Bacteria | Pseudomonadota | Alphaproteobacteria | Rhodobacterales | Paracoccaceae | Paracoccus |
| Paracoccus saliphilus | psap | 1 | Bacteria | Pseudomonadota | Alphaproteobacteria | Rhodobacterales | Paracoccaceae | Paracoccus |
| Paroceanicella profunda | ppru | 1 | Bacteria | Pseudomonadota | Alphaproteobacteria | Rhodobacterales | Paracoccaceae | Paroceanicella |
| Planktomarina temperata | ptp | 3 | Bacteria | Pseudomonadota | Alphaproteobacteria | Rhodobacterales | Paracoccaceae | Planktomarina |
| Pseudooceanicola algae | palw | 4 | Bacteria | Pseudomonadota | Alphaproteobacteria | Rhodobacterales | Paracoccaceae | Pseudooceanicola |
| Rhodovulum sp. P5 | rhc | 2 | Bacteria | Pseudomonadota | Alphaproteobacteria | Rhodobacterales | Paracoccaceae | Rhodovulum |
| Rhodovulum sulfidophilum | rsu | 1 | Bacteria | Pseudomonadota | Alphaproteobacteria | Rhodobacterales | Paracoccaceae | Rhodovulum |
| Rhodobacter capsulatus | rcp | 1 | Bacteria | Pseudomonadota | Alphaproteobacteria | Rhodobacterales | Rhodobactergroup | Rhodobacter |
| Celeribacter indicus | cid | 1 | Bacteria | Pseudomonadota | Alphaproteobacteria | Rhodobacterales | Roseobacteraceae | Celeribacter |
| Celeribacter marinus | cmar | 1 | Bacteria | Pseudomonadota | Alphaproteobacteria | Rhodobacterales | Roseobacteraceae | Celeribacter |
| Ketogulonicigenium robustum | kro | 3 | Bacteria | Pseudomonadota | Alphaproteobacteria | Rhodobacterales | Roseobacteraceae | Ketogulonicigenium |
| Ketogulonicigenium vulgare WSH-001 | kvl | 1 | Bacteria | Pseudomonadota | Alphaproteobacteria | Rhodobacterales | Roseobacteraceae | Ketogulonicigenium |
| Ketogulonicigenium vulgare Y25 | kvu | 1 | Bacteria | Pseudomonadota | Alphaproteobacteria | Rhodobacterales | Roseobacteraceae | Ketogulonicigenium |
| Lentibacter algarum | lalg | 3 | Bacteria | Pseudomonadota | Alphaproteobacteria | Rhodobacterales | Roseobacteraceae | Lentibacter |
| Octadecabacter antarcticus | oat | 2 | Bacteria | Pseudomonadota | Alphaproteobacteria | Rhodobacterales | Roseobacteraceae | Octadecabacter |
| Octadecabacter arcticus | oar | 2 | Bacteria | Pseudomonadota | Alphaproteobacteria | Rhodobacterales | Roseobacteraceae | Octadecabacter |
| Phaeobacter inhibens 2.10 | pgl | 2 | Bacteria | Pseudomonadota | Alphaproteobacteria | Rhodobacterales | Roseobacteraceae | Phaeobacter |
| Phaeobacter inhibens DSM 17395 | pga | 2 | Bacteria | Pseudomonadota | Alphaproteobacteria | Rhodobacterales | Roseobacteraceae | Phaeobacter |
| Phaeobacter piscinae | ppic | 2 | Bacteria | Pseudomonadota | Alphaproteobacteria | Rhodobacterales | Roseobacteraceae | Phaeobacter |
| Phaeobacter porticola | php | 2 | Bacteria | Pseudomonadota | Alphaproteobacteria | Rhodobacterales | Roseobacteraceae | Phaeobacter |
| Roseicyclus elongatus | red | 2 | Bacteria | Pseudomonadota | Alphaproteobacteria | Rhodobacterales | Roseobacteraceae | Roseicyclus |
| Roseobacter fucihabitans | rfu | 3 | Bacteria | Pseudomonadota | Alphaproteobacteria | Rhodobacterales | Roseobacteraceae | Roseobacter |
| Roseobacter litoralis | rli | 2 | Bacteria | Pseudomonadota | Alphaproteobacteria | Rhodobacterales | Roseobacteraceae | Roseobacter |
| Roseovarius sp. AK1035 | rok | 1 | Bacteria | Pseudomonadota | Alphaproteobacteria | Rhodobacterales | Roseobacteraceae | Roseovarius |
| Ruegeria pomeroyi | sil | 1 | Bacteria | Pseudomonadota | Alphaproteobacteria | Rhodobacterales | Roseobacteraceae | Ruegeria |
| Salipiger abyssi | paby | 4 | Bacteria | Pseudomonadota | Alphaproteobacteria | Rhodobacterales | Roseobacteraceae | Salipiger |
| Salipiger profundus | tpro | 3 | Bacteria | Pseudomonadota | Alphaproteobacteria | Rhodobacterales | Roseobacteraceae | Salipiger |
| Sulfitobacter indolifex | sinl | 5 | Bacteria | Pseudomonadota | Alphaproteobacteria | Rhodobacterales | Roseobacteraceae | Sulfitobacter |
| Azospirillum baldaniorum Sp245 | abs | 2 | Bacteria | Pseudomonadota | Alphaproteobacteria | Rhodospirillales | Azospirillaceae | Azospirillum |
| Azospirillum lipoferum | ali | 1 | Bacteria | Pseudomonadota | Alphaproteobacteria | Rhodospirillales | Azospirillaceae | Azospirillum |
| Azospirillum sp. B510 | azl | 1 | Bacteria | Pseudomonadota | Alphaproteobacteria | Rhodospirillales | Azospirillaceae | Azospirillum |
| Magnetospira sp. QH-2 | magq | 1 | Bacteria | Pseudomonadota | Alphaproteobacteria | Rhodospirillales | Thalassospiraceae | Magnetospira |
| Thalassospira xiamenensis | txi | 1 | Bacteria | Pseudomonadota | Alphaproteobacteria | Rhodospirillales | Thalassospiraceae | Thalassospira |
| Porphyrobacter sp. LM 6 | porl | 1 | Bacteria | Pseudomonadota | Alphaproteobacteria | Sphingomonadales | Erythrobacteraceae | Porphyrobacter |
| Erythrobacter litoralis DSM 8509 | elq | 2 | Bacteria | Pseudomonadota | Alphaproteobacteria | Sphingomonadales | Erythrobacteraceae | Erythrobacter |
| Sphingobium sp. AntQ-1 | spag | 2 | Bacteria | Pseudomonadota | Alphaproteobacteria | Sphingomonadales | Sphingobiaceae | Sphingobium |
| Sphingobium sp. EP60837 | sphb | 2 | Bacteria | Pseudomonadota | Alphaproteobacteria | Sphingomonadales | Sphingobiaceae | Sphingobium |
| Sphingobium sp. MI1205 | spmi | 1 | Bacteria | Pseudomonadota | Alphaproteobacteria | Sphingomonadales | Sphingobiaceae | Sphingobium |
| Sphingobium sp. TKS | spht | 1 | Bacteria | Pseudomonadota | Alphaproteobacteria | Sphingomonadales | Sphingobiaceae | Sphingobium |
| Sphingobium sp. YG1 | spyg | 2 | Bacteria | Pseudomonadota | Alphaproteobacteria | Sphingomonadales | Sphingobiaceae | Sphingobium |
| Novosphingobium sp. PP1Y | npp | 4 | Bacteria | Pseudomonadota | Alphaproteobacteria | Sphingomonadales | Sphingomonadaceae | Novosphingobium |
| Sphingomonas taxi | stax | 1 | Bacteria | Pseudomonadota | Alphaproteobacteria | Sphingomonadales | Sphingomonadaceae | Sphingomonas |
| Sphingopyxis fribergensis | sphk | 1 | Bacteria | Pseudomonadota | Alphaproteobacteria | Sphingomonadales | Sphingopyxidaceae | Sphingopyxis |
| Sphingopyxis sp. EG6 | speg | 1 | Bacteria | Pseudomonadota | Alphaproteobacteria | Sphingomonadales | Sphingopyxidaceae | Sphingopyxis |
| Sphingopyxis sp. FD7 | spfd | 1 | Bacteria | Pseudomonadota | Alphaproteobacteria | Sphingomonadales | Sphingopyxidaceae | Sphingopyxis |
| Uncultured Sphingopyxis sp. UC10 | sphu | 1 | Bacteria | Pseudomonadota | Alphaproteobacteria | Sphingomonadales | Sphingopyxidaceae | Sphingopyxis |
| Achromobacter aegrifaciens | aaeg | 2 | Bacteria | Pseudomonadota | Betaproteobacteria | Burkholderiales | Alcaligenaceae | Achromobacter |
| Achromobacter insolitus | ais | 2 | Bacteria | Pseudomonadota | Betaproteobacteria | Burkholderiales | Alcaligenaceae | Achromobacter |
| Achromobacter pestifer | apes | 2 | Bacteria | Pseudomonadota | Betaproteobacteria | Burkholderiales | Alcaligenaceae | Achromobacter |
| Achromobacter sp. 77 | acho | 1 | Bacteria | Pseudomonadota | Betaproteobacteria | Burkholderiales | Alcaligenaceae | Achromobacter |
| Achromobacter sp. AONIH1 | achr | 2 | Bacteria | Pseudomonadota | Betaproteobacteria | Burkholderiales | Alcaligenaceae | Achromobacter |
| Achromobacter xylosoxidans A8 | axy | 1 | Bacteria | Pseudomonadota | Betaproteobacteria | Burkholderiales | Alcaligenaceae | Achromobacter |
| Advenella alkanexedens | aaln | 1 | Bacteria | Pseudomonadota | Betaproteobacteria | Burkholderiales | Alcaligenaceae | Advenella |
| Advenella mimigardefordensis | amim | 2 | Bacteria | Pseudomonadota | Betaproteobacteria | Burkholderiales | Alcaligenaceae | Advenella |
| Bordetella avium | bav | 1 | Bacteria | Pseudomonadota | Betaproteobacteria | Burkholderiales | Alcaligenaceae | Bordetella |
| Bordetella bronchiseptica 253 | bbh | 1 | Bacteria | Pseudomonadota | Betaproteobacteria | Burkholderiales | Alcaligenaceae | Bordetella |
| Bordetella bronchiseptica MO149 | bbm | 1 | Bacteria | Pseudomonadota | Betaproteobacteria | Burkholderiales | Alcaligenaceae | Bordetella |
| Bordetella bronchiseptica RB50 | bbr | 1 | Bacteria | Pseudomonadota | Betaproteobacteria | Burkholderiales | Alcaligenaceae | Bordetella |
| Bordetella bronchiseptica S798 | bbx | 1 | Bacteria | Pseudomonadota | Betaproteobacteria | Burkholderiales | Alcaligenaceae | Bordetella |
| Bordetella hinzii | bhz | 1 | Bacteria | Pseudomonadota | Betaproteobacteria | Burkholderiales | Alcaligenaceae | Bordetella |
| Bordetella parapertussis 12822 | bpa | 1 | Bacteria | Pseudomonadota | Betaproteobacteria | Burkholderiales | Alcaligenaceae | Bordetella |
| Bordetella pseudohinzii | bpdz | 1 | Bacteria | Pseudomonadota | Betaproteobacteria | Burkholderiales | Alcaligenaceae | Bordetella |
| Bordetella trematum | btrm | 1 | Bacteria | Pseudomonadota | Betaproteobacteria | Burkholderiales | Alcaligenaceae | Bordetella |
| Castellaniella defragrans | cdn | 1 | Bacteria | Pseudomonadota | Betaproteobacteria | Burkholderiales | Alcaligenaceae | Castellaniella |
| Neopusillimonas aestuarii | pud | 1 | Bacteria | Pseudomonadota | Betaproteobacteria | Burkholderiales | Alcaligenaceae | Neopusillimonas |
| Neopusillimonas aromaticivorans | narm | 1 | Bacteria | Pseudomonadota | Betaproteobacteria | Burkholderiales | Alcaligenaceae | Neopusillimonas |
| Orrella dioscoreae | odi | 1 | Bacteria | Pseudomonadota | Betaproteobacteria | Burkholderiales | Alcaligenaceae | Orrella |
| Paralcaligenes sp. KSB-10 | park | 1 | Bacteria | Pseudomonadota | Betaproteobacteria | Burkholderiales | Alcaligenaceae | Paralcaligenes |
| Pigmentiphaga aceris | pacr | 1 | Bacteria | Pseudomonadota | Betaproteobacteria | Burkholderiales | Alcaligenaceae | Pigmentiphaga |
| Pigmentiphaga sp. H8 | pig | 1 | Bacteria | Pseudomonadota | Betaproteobacteria | Burkholderiales | Alcaligenaceae | Pigmentiphaga |
| Burkholderia aenigmatica | baen | 5 | Bacteria | Pseudomonadota | Betaproteobacteria | Burkholderiales | Burkholderiaceae | Burkholderia |
| Burkholderia ambifaria AMMD | bam | 5 | Bacteria | Pseudomonadota | Betaproteobacteria | Burkholderiales | Burkholderiaceae | Burkholderia |
| Burkholderia ambifaria MC40-6 | bac | 5 | Bacteria | Pseudomonadota | Betaproteobacteria | Burkholderiales | Burkholderiaceae | Burkholderia |
| Burkholderia anthina | bann | 3 | Bacteria | Pseudomonadota | Betaproteobacteria | Burkholderiales | Burkholderiaceae | Burkholderia |
| Burkholderia arboris | bari | 4 | Bacteria | Pseudomonadota | Betaproteobacteria | Burkholderiales | Burkholderiaceae | Burkholderia |
| Burkholderia cenocepacia DDS 22E-1 | bcen | 4 | Bacteria | Pseudomonadota | Betaproteobacteria | Burkholderiales | Burkholderiaceae | Burkholderia |
| Burkholderia cenocepacia DWS 37E-2 | bcew | 4 | Bacteria | Pseudomonadota | Betaproteobacteria | Burkholderiales | Burkholderiaceae | Burkholderia |
| Burkholderia cenocepacia H111 | bceo | 4 | Bacteria | Pseudomonadota | Betaproteobacteria | Burkholderiales | Burkholderiaceae | Burkholderia |
| Burkholderia cenocepacia HI2424 | bch | 4 | Bacteria | Pseudomonadota | Betaproteobacteria | Burkholderiales | Burkholderiaceae | Burkholderia |
| Burkholderia cenocepacia J2315 | bcj | 3 | Bacteria | Pseudomonadota | Betaproteobacteria | Burkholderiales | Burkholderiaceae | Burkholderia |
| Burkholderia cepacia ATCC 25416 | bcep | 5 | Bacteria | Pseudomonadota | Betaproteobacteria | Burkholderiales | Burkholderiaceae | Burkholderia |
| Burkholderia cepacia DDS 7H-2 | bced | 3 | Bacteria | Pseudomonadota | Betaproteobacteria | Burkholderiales | Burkholderiaceae | Burkholderia |
| Burkholderia cepacia GG4 | bct | 3 | Bacteria | Pseudomonadota | Betaproteobacteria | Burkholderiales | Burkholderiaceae | Burkholderia |
| Burkholderia contaminans | bcon | 4 | Bacteria | Pseudomonadota | Betaproteobacteria | Burkholderiales | Burkholderiaceae | Burkholderia |
| Burkholderia diffusa | bdf | 3 | Bacteria | Pseudomonadota | Betaproteobacteria | Burkholderiales | Burkholderiaceae | Burkholderia |
| Burkholderia dolosa | bdl | 2 | Bacteria | Pseudomonadota | Betaproteobacteria | Burkholderiales | Burkholderiaceae | Burkholderia |
| Burkholderia gladioli ATCC 10248 | bgo | 3 | Bacteria | Pseudomonadota | Betaproteobacteria | Burkholderiales | Burkholderiaceae | Burkholderia |
| Burkholderia gladioli BSR3 | bgd | 3 | Bacteria | Pseudomonadota | Betaproteobacteria | Burkholderiales | Burkholderiaceae | Burkholderia |
| Burkholderia glumae BGR1 | bgl | 1 | Bacteria | Pseudomonadota | Betaproteobacteria | Burkholderiales | Burkholderiaceae | Burkholderia |
| Burkholderia glumae LMG 2196 = ATCC 33617 | bgu | 1 | Bacteria | Pseudomonadota | Betaproteobacteria | Burkholderiales | Burkholderiaceae | Burkholderia |
| Burkholderia humptydooensis 2002721687 | bul | 1 | Bacteria | Pseudomonadota | Betaproteobacteria | Burkholderiales | Burkholderiaceae | Burkholderia |
| Burkholderia humptydooensis FDAARGOS_899 | bhg | 1 | Bacteria | Pseudomonadota | Betaproteobacteria | Burkholderiales | Burkholderiaceae | Burkholderia |
| Burkholderia humptydooensis MSMB43 | bud | 1 | Bacteria | Pseudomonadota | Betaproteobacteria | Burkholderiales | Burkholderiaceae | Burkholderia |
| Burkholderia lata | bur | 5 | Bacteria | Pseudomonadota | Betaproteobacteria | Burkholderiales | Burkholderiaceae | Burkholderia |
| Burkholderia latens | blat | 4 | Bacteria | Pseudomonadota | Betaproteobacteria | Burkholderiales | Burkholderiaceae | Burkholderia |
| Burkholderia mallei 2000031063 | bmai | 1 | Bacteria | Pseudomonadota | Betaproteobacteria | Burkholderiales | Burkholderiaceae | Burkholderia |
| Burkholderia mallei 2002734299 | bmab | 2 | Bacteria | Pseudomonadota | Betaproteobacteria | Burkholderiales | Burkholderiaceae | Burkholderia |
| Burkholderia mallei 23344 | bmal | 1 | Bacteria | Pseudomonadota | Betaproteobacteria | Burkholderiales | Burkholderiaceae | Burkholderia |
| Burkholderia mallei 6 | bmae | 2 | Bacteria | Pseudomonadota | Betaproteobacteria | Burkholderiales | Burkholderiaceae | Burkholderia |
| Burkholderia mallei ATCC 23344 | bma | 2 | Bacteria | Pseudomonadota | Betaproteobacteria | Burkholderiales | Burkholderiaceae | Burkholderia |
| Burkholderia mallei BMQ | bmaq | 1 | Bacteria | Pseudomonadota | Betaproteobacteria | Burkholderiales | Burkholderiaceae | Burkholderia |
| Burkholderia mallei FMH 23344 | bmaf | 2 | Bacteria | Pseudomonadota | Betaproteobacteria | Burkholderiales | Burkholderiaceae | Burkholderia |
| Burkholderia mallei NCTC 10229 | bml | 2 | Bacteria | Pseudomonadota | Betaproteobacteria | Burkholderiales | Burkholderiaceae | Burkholderia |
| Burkholderia mallei NCTC 10247 | bmaz | 2 | Bacteria | Pseudomonadota | Betaproteobacteria | Burkholderiales | Burkholderiaceae | Burkholderia |
| Burkholderia mallei NCTC 10247 | bmn | 2 | Bacteria | Pseudomonadota | Betaproteobacteria | Burkholderiales | Burkholderiaceae | Burkholderia |
| Burkholderia mallei SAVP1 | bmv | 1 | Bacteria | Pseudomonadota | Betaproteobacteria | Burkholderiales | Burkholderiaceae | Burkholderia |
| Burkholderia mayonis | buu | 2 | Bacteria | Pseudomonadota | Betaproteobacteria | Burkholderiales | Burkholderiaceae | Burkholderia |
| Burkholderia metallica | bmec | 5 | Bacteria | Pseudomonadota | Betaproteobacteria | Burkholderiales | Burkholderiaceae | Burkholderia |
| Burkholderia multivorans ATCC 17616 (JGI) | bmu | 4 | Bacteria | Pseudomonadota | Betaproteobacteria | Burkholderiales | Burkholderiaceae | Burkholderia |
| Burkholderia multivorans ATCC 17616 (Tohoku) | bmj | 5 | Bacteria | Pseudomonadota | Betaproteobacteria | Burkholderiales | Burkholderiaceae | Burkholderia |
| Burkholderia multivorans ATCC BAA-247 | bmul | 4 | Bacteria | Pseudomonadota | Betaproteobacteria | Burkholderiales | Burkholderiaceae | Burkholderia |
| Burkholderia multivorans DDS 15A-1 | bmk | 4 | Bacteria | Pseudomonadota | Betaproteobacteria | Burkholderiales | Burkholderiaceae | Burkholderia |
| Burkholderia oklahomensis C6786 | boc | 2 | Bacteria | Pseudomonadota | Betaproteobacteria | Burkholderiales | Burkholderiaceae | Burkholderia |
| Burkholderia oklahomensis EO147 | bok | 2 | Bacteria | Pseudomonadota | Betaproteobacteria | Burkholderiales | Burkholderiaceae | Burkholderia |
| Burkholderia orbicola | bcm | 4 | Bacteria | Pseudomonadota | Betaproteobacteria | Burkholderiales | Burkholderiaceae | Burkholderia |
| Burkholderia plantarii ATCC 43733 | bpla | 3 | Bacteria | Pseudomonadota | Betaproteobacteria | Burkholderiales | Burkholderiaceae | Burkholderia |
| Burkholderia plantarii PG1 | bgp | 3 | Bacteria | Pseudomonadota | Betaproteobacteria | Burkholderiales | Burkholderiaceae | Burkholderia |
| Burkholderia pseudomallei 1026b | bpz | 2 | Bacteria | Pseudomonadota | Betaproteobacteria | Burkholderiales | Burkholderiaceae | Burkholderia |
| Burkholderia pseudomallei 1106a | bpl | 2 | Bacteria | Pseudomonadota | Betaproteobacteria | Burkholderiales | Burkholderiaceae | Burkholderia |
| Burkholderia pseudomallei 1710b | bpm | 2 | Bacteria | Pseudomonadota | Betaproteobacteria | Burkholderiales | Burkholderiaceae | Burkholderia |
| Burkholderia pseudomallei 668 | bpd | 2 | Bacteria | Pseudomonadota | Betaproteobacteria | Burkholderiales | Burkholderiaceae | Burkholderia |
| Burkholderia pseudomallei A79A | bpso | 2 | Bacteria | Pseudomonadota | Betaproteobacteria | Burkholderiales | Burkholderiaceae | Burkholderia |
| Burkholderia pseudomallei BPC006 | bpq | 2 | Bacteria | Pseudomonadota | Betaproteobacteria | Burkholderiales | Burkholderiaceae | Burkholderia |
| Burkholderia pseudomallei HBPUB10134a | bpsh | 2 | Bacteria | Pseudomonadota | Betaproteobacteria | Burkholderiales | Burkholderiaceae | Burkholderia |
| Burkholderia pseudomallei K96243 | bps | 2 | Bacteria | Pseudomonadota | Betaproteobacteria | Burkholderiales | Burkholderiaceae | Burkholderia |
| Burkholderia pseudomallei MSHR146 | bpsu | 2 | Bacteria | Pseudomonadota | Betaproteobacteria | Burkholderiales | Burkholderiaceae | Burkholderia |
| Burkholderia pseudomallei MSHR305 | bpse | 2 | Bacteria | Pseudomonadota | Betaproteobacteria | Burkholderiales | Burkholderiaceae | Burkholderia |
| Burkholderia pseudomallei MSHR511 | bpsm | 2 | Bacteria | Pseudomonadota | Betaproteobacteria | Burkholderiales | Burkholderiaceae | Burkholderia |
| Burkholderia pseudomallei MSHR520 | bpsd | 2 | Bacteria | Pseudomonadota | Betaproteobacteria | Burkholderiales | Burkholderiaceae | Burkholderia |
| Burkholderia pseudomallei NAU35A-3 | bpsa | 2 | Bacteria | Pseudomonadota | Betaproteobacteria | Burkholderiales | Burkholderiaceae | Burkholderia |
| Burkholderia pseudomallei NCTC 13179 | bpk | 2 | Bacteria | Pseudomonadota | Betaproteobacteria | Burkholderiales | Burkholderiaceae | Burkholderia |
| Burkholderia pseudomallei TSV202 | but | 2 | Bacteria | Pseudomonadota | Betaproteobacteria | Burkholderiales | Burkholderiaceae | Burkholderia |
| Burkholderia pseudomultivorans | bpsl | 4 | Bacteria | Pseudomonadota | Betaproteobacteria | Burkholderiales | Burkholderiaceae | Burkholderia |
| Burkholderia pyrrocinia | bpyr | 4 | Bacteria | Pseudomonadota | Betaproteobacteria | Burkholderiales | Burkholderiaceae | Burkholderia |
| Burkholderia savannae | bsav | 2 | Bacteria | Pseudomonadota | Betaproteobacteria | Burkholderiales | Burkholderiaceae | Burkholderia |
| Burkholderia semiarida | bser | 4 | Bacteria | Pseudomonadota | Betaproteobacteria | Burkholderiales | Burkholderiaceae | Burkholderia |
| Burkholderia seminalis | bsem | 5 | Bacteria | Pseudomonadota | Betaproteobacteria | Burkholderiales | Burkholderiaceae | Burkholderia |
| Burkholderia sp. CCGE1001 | bug | 3 | Bacteria | Pseudomonadota | Betaproteobacteria | Burkholderiales | Burkholderiaceae | Burkholderia |
| Burkholderia sp. CCGE1003 | bgf | 1 | Bacteria | Pseudomonadota | Betaproteobacteria | Burkholderiales | Burkholderiaceae | Burkholderia |
| Burkholderia sp. HB1 | buq | 2 | Bacteria | Pseudomonadota | Betaproteobacteria | Burkholderiales | Burkholderiaceae | Burkholderia |
| Burkholderia sp. JP2-270 | burk | 3 | Bacteria | Pseudomonadota | Betaproteobacteria | Burkholderiales | Burkholderiaceae | Burkholderia |
| Burkholderia sp. KJ006 | buk | 2 | Bacteria | Pseudomonadota | Betaproteobacteria | Burkholderiales | Burkholderiaceae | Burkholderia |
| Burkholderia sp. PAMC 26561 | bum | 3 | Bacteria | Pseudomonadota | Betaproteobacteria | Burkholderiales | Burkholderiaceae | Burkholderia |
| Burkholderia sp. PAMC 28687 | bui | 1 | Bacteria | Pseudomonadota | Betaproteobacteria | Burkholderiales | Burkholderiaceae | Burkholderia |
| Burkholderia sp. YI23 | byi | 2 | Bacteria | Pseudomonadota | Betaproteobacteria | Burkholderiales | Burkholderiaceae | Burkholderia |
| Burkholderia stabilis | bstl | 3 | Bacteria | Pseudomonadota | Betaproteobacteria | Burkholderiales | Burkholderiaceae | Burkholderia |
| Burkholderia stagnalis | bstg | 4 | Bacteria | Pseudomonadota | Betaproteobacteria | Burkholderiales | Burkholderiaceae | Burkholderia |
| Burkholderia territorii | btei | 4 | Bacteria | Pseudomonadota | Betaproteobacteria | Burkholderiales | Burkholderiaceae | Burkholderia |
| Burkholderia thailandensis 2002721643 | bthl | 1 | Bacteria | Pseudomonadota | Betaproteobacteria | Burkholderiales | Burkholderiaceae | Burkholderia |
| Burkholderia thailandensis 2002721723 | btq | 1 | Bacteria | Pseudomonadota | Betaproteobacteria | Burkholderiales | Burkholderiaceae | Burkholderia |
| Burkholderia thailandensis 2003015869 | btha | 2 | Bacteria | Pseudomonadota | Betaproteobacteria | Burkholderiales | Burkholderiaceae | Burkholderia |
| Burkholderia thailandensis E254 | bthe | 1 | Bacteria | Pseudomonadota | Betaproteobacteria | Burkholderiales | Burkholderiaceae | Burkholderia |
| Burkholderia thailandensis E264 | bte | 1 | Bacteria | Pseudomonadota | Betaproteobacteria | Burkholderiales | Burkholderiaceae | Burkholderia |
| Burkholderia thailandensis E444 | btj | 1 | Bacteria | Pseudomonadota | Betaproteobacteria | Burkholderiales | Burkholderiaceae | Burkholderia |
| Burkholderia thailandensis H0587 | btz | 2 | Bacteria | Pseudomonadota | Betaproteobacteria | Burkholderiales | Burkholderiaceae | Burkholderia |
| Burkholderia thailandensis MSMB121 | btd | 2 | Bacteria | Pseudomonadota | Betaproteobacteria | Burkholderiales | Burkholderiaceae | Burkholderia |
| Burkholderia thailandensis MSMB59 | btv | 1 | Bacteria | Pseudomonadota | Betaproteobacteria | Burkholderiales | Burkholderiaceae | Burkholderia |
| Burkholderia thailandensis USAMRU Malaysia #20 | bthm | 1 | Bacteria | Pseudomonadota | Betaproteobacteria | Burkholderiales | Burkholderiaceae | Burkholderia |
| Burkholderia ubonensis | bub | 6 | Bacteria | Pseudomonadota | Betaproteobacteria | Burkholderiales | Burkholderiaceae | Burkholderia |
| Burkholderia vietnamiensis G4 | bvi | 2 | Bacteria | Pseudomonadota | Betaproteobacteria | Burkholderiales | Burkholderiaceae | Burkholderia |
| Burkholderia vietnamiensis LMG 10929 | bve | 2 | Bacteria | Pseudomonadota | Betaproteobacteria | Burkholderiales | Burkholderiaceae | Burkholderia |
| Caballeronia cordobensis | bue | 2 | Bacteria | Pseudomonadota | Betaproteobacteria | Burkholderiales | Burkholderiaceae | Caballeronia |
| Caballeronia insecticola | buo | 2 | Bacteria | Pseudomonadota | Betaproteobacteria | Burkholderiales | Burkholderiaceae | Caballeronia |
| Caballeronia sp. NK8 | cabk | 3 | Bacteria | Pseudomonadota | Betaproteobacteria | Burkholderiales | Burkholderiaceae | Caballeronia |
| Caballeronia sp. SBC2 | caba | 5 | Bacteria | Pseudomonadota | Betaproteobacteria | Burkholderiales | Burkholderiaceae | Caballeronia |
| Cupriavidus basilensis | cbw | 3 | Bacteria | Pseudomonadota | Betaproteobacteria | Burkholderiales | Burkholderiaceae | Cupriavidus |
| Cupriavidus campinensis | ccam | 2 | Bacteria | Pseudomonadota | Betaproteobacteria | Burkholderiales | Burkholderiaceae | Cupriavidus |
| Cupriavidus cauae | ccax | 2 | Bacteria | Pseudomonadota | Betaproteobacteria | Burkholderiales | Burkholderiaceae | Cupriavidus |
| Cupriavidus gilardii | cgd | 2 | Bacteria | Pseudomonadota | Betaproteobacteria | Burkholderiales | Burkholderiaceae | Cupriavidus |
| Cupriavidus malaysiensis | cup | 6 | Bacteria | Pseudomonadota | Betaproteobacteria | Burkholderiales | Burkholderiaceae | Cupriavidus |
| Cupriavidus metallidurans | rme | 4 | Bacteria | Pseudomonadota | Betaproteobacteria | Burkholderiales | Burkholderiaceae | Cupriavidus |
| Cupriavidus nantongensis | cnan | 2 | Bacteria | Pseudomonadota | Betaproteobacteria | Burkholderiales | Burkholderiaceae | Cupriavidus |
| Cupriavidus necator H16 | reh | 4 | Bacteria | Pseudomonadota | Betaproteobacteria | Burkholderiales | Burkholderiaceae | Cupriavidus |
| Cupriavidus necator N-1 | cnc | 5 | Bacteria | Pseudomonadota | Betaproteobacteria | Burkholderiales | Burkholderiaceae | Cupriavidus |
| Cupriavidus necator NH9 | cuh | 4 | Bacteria | Pseudomonadota | Betaproteobacteria | Burkholderiales | Burkholderiaceae | Cupriavidus |
| Cupriavidus oxalaticus | cox | 5 | Bacteria | Pseudomonadota | Betaproteobacteria | Burkholderiales | Burkholderiaceae | Cupriavidus |
| Cupriavidus pauculus | cpau | 2 | Bacteria | Pseudomonadota | Betaproteobacteria | Burkholderiales | Burkholderiaceae | Cupriavidus |
| Cupriavidus pinatubonensis JMP134 | reu | 4 | Bacteria | Pseudomonadota | Betaproteobacteria | Burkholderiales | Burkholderiaceae | Cupriavidus |
| Cupriavidus sp. EM10 | cupe | 2 | Bacteria | Pseudomonadota | Betaproteobacteria | Burkholderiales | Burkholderiaceae | Cupriavidus |
| Cupriavidus sp. ISTL7 | cupr | 1 | Bacteria | Pseudomonadota | Betaproteobacteria | Burkholderiales | Burkholderiaceae | Cupriavidus |
| Cupriavidus sp. KK10 | cuk | 4 | Bacteria | Pseudomonadota | Betaproteobacteria | Burkholderiales | Burkholderiaceae | Cupriavidus |
| Cupriavidus sp. USMAA2-4 | cuu | 5 | Bacteria | Pseudomonadota | Betaproteobacteria | Burkholderiales | Burkholderiaceae | Cupriavidus |
| Cupriavidus sp. USMAHM13 | ccup | 5 | Bacteria | Pseudomonadota | Betaproteobacteria | Burkholderiales | Burkholderiaceae | Cupriavidus |
| Cupriavidus taiwanensis | cti | 5 | Bacteria | Pseudomonadota | Betaproteobacteria | Burkholderiales | Burkholderiaceae | Cupriavidus |
| Pandoraea apista | papi | 2 | Bacteria | Pseudomonadota | Betaproteobacteria | Burkholderiales | Burkholderiaceae | Pandoraea |
| Pandoraea commovens | pcom | 1 | Bacteria | Pseudomonadota | Betaproteobacteria | Burkholderiales | Burkholderiaceae | Pandoraea |
| Pandoraea faecigallinarum | pfg | 1 | Bacteria | Pseudomonadota | Betaproteobacteria | Burkholderiales | Burkholderiaceae | Pandoraea |
| Pandoraea fibrosis | pfib | 1 | Bacteria | Pseudomonadota | Betaproteobacteria | Burkholderiales | Burkholderiaceae | Pandoraea |
| Pandoraea norimbergensis | pnr | 1 | Bacteria | Pseudomonadota | Betaproteobacteria | Burkholderiales | Burkholderiaceae | Pandoraea |
| Pandoraea oxalativorans | pox | 1 | Bacteria | Pseudomonadota | Betaproteobacteria | Burkholderiales | Burkholderiaceae | Pandoraea |
| Pandoraea pnomenusa | prb | 1 | Bacteria | Pseudomonadota | Betaproteobacteria | Burkholderiales | Burkholderiaceae | Pandoraea |
| Pandoraea pnomenusa 3kgm | ppk | 1 | Bacteria | Pseudomonadota | Betaproteobacteria | Burkholderiales | Burkholderiaceae | Pandoraea |
| Pandoraea pnomenusa DSM 16536 | ppnm | 1 | Bacteria | Pseudomonadota | Betaproteobacteria | Burkholderiales | Burkholderiaceae | Pandoraea |
| Pandoraea pnomenusa RB38 | ppno | 1 | Bacteria | Pseudomonadota | Betaproteobacteria | Burkholderiales | Burkholderiaceae | Pandoraea |
| Pandoraea pulmonicola | ppul | 2 | Bacteria | Pseudomonadota | Betaproteobacteria | Burkholderiales | Burkholderiaceae | Pandoraea |
| Pandoraea sp. NE5 | pann | 2 | Bacteria | Pseudomonadota | Betaproteobacteria | Burkholderiales | Burkholderiaceae | Pandoraea |
| Pandoraea sp. XJJ-1 | panx | 1 | Bacteria | Pseudomonadota | Betaproteobacteria | Burkholderiales | Burkholderiaceae | Pandoraea |
| Pandoraea sp. XY-2 | pand | 1 | Bacteria | Pseudomonadota | Betaproteobacteria | Burkholderiales | Burkholderiaceae | Pandoraea |
| Pandoraea sputorum | pspu | 2 | Bacteria | Pseudomonadota | Betaproteobacteria | Burkholderiales | Burkholderiaceae | Pandoraea |
| Pandoraea thiooxydans | ptx | 2 | Bacteria | Pseudomonadota | Betaproteobacteria | Burkholderiales | Burkholderiaceae | Pandoraea |
| Pandoraea vervacti | pve | 1 | Bacteria | Pseudomonadota | Betaproteobacteria | Burkholderiales | Burkholderiaceae | Pandoraea |
| Paraburkholderia acidiphila | pacp | 4 | Bacteria | Pseudomonadota | Betaproteobacteria | Burkholderiales | Burkholderiaceae | Paraburkholderia |
| Paraburkholderia acidisoli | pacs | 4 | Bacteria | Pseudomonadota | Betaproteobacteria | Burkholderiales | Burkholderiaceae | Paraburkholderia |
| Paraburkholderia aromaticivorans | parb | 3 | Bacteria | Pseudomonadota | Betaproteobacteria | Burkholderiales | Burkholderiaceae | Paraburkholderia |
| Paraburkholderia atlantica CCGE1002 | bge | 2 | Bacteria | Pseudomonadota | Betaproteobacteria | Burkholderiales | Burkholderiaceae | Paraburkholderia |
| Paraburkholderia bryophila | pbry | 2 | Bacteria | Pseudomonadota | Betaproteobacteria | Burkholderiales | Burkholderiaceae | Paraburkholderia |
| Paraburkholderia caffeinilytica | pcaf | 3 | Bacteria | Pseudomonadota | Betaproteobacteria | Burkholderiales | Burkholderiaceae | Paraburkholderia |
| Paraburkholderia caledonica | pcj | 1 | Bacteria | Pseudomonadota | Betaproteobacteria | Burkholderiales | Burkholderiaceae | Paraburkholderia |
| Paraburkholderia caribensis | bcai | 2 | Bacteria | Pseudomonadota | Betaproteobacteria | Burkholderiales | Burkholderiaceae | Paraburkholderia |
| Paraburkholderia dioscoreae | pdio | 3 | Bacteria | Pseudomonadota | Betaproteobacteria | Burkholderiales | Burkholderiaceae | Paraburkholderia |
| Paraburkholderia edwinii | pew | 1 | Bacteria | Pseudomonadota | Betaproteobacteria | Burkholderiales | Burkholderiaceae | Paraburkholderia |
| Paraburkholderia fungorum | bfn | 3 | Bacteria | Pseudomonadota | Betaproteobacteria | Burkholderiales | Burkholderiaceae | Paraburkholderia |
| Paraburkholderia ginsengisoli | pgis | 4 | Bacteria | Pseudomonadota | Betaproteobacteria | Burkholderiales | Burkholderiaceae | Paraburkholderia |
| Paraburkholderia graminis | pgp | 2 | Bacteria | Pseudomonadota | Betaproteobacteria | Burkholderiales | Burkholderiaceae | Paraburkholderia |
| Paraburkholderia hospita | phs | 2 | Bacteria | Pseudomonadota | Betaproteobacteria | Burkholderiales | Burkholderiaceae | Paraburkholderia |
| Paraburkholderia kirstenboschensis | pkf | 1 | Bacteria | Pseudomonadota | Betaproteobacteria | Burkholderiales | Burkholderiaceae | Paraburkholderia |
| Paraburkholderia kururiensis | phun | 1 | Bacteria | Pseudomonadota | Betaproteobacteria | Burkholderiales | Burkholderiaceae | Paraburkholderia |
| Paraburkholderia largidicola | plad | 2 | Bacteria | Pseudomonadota | Betaproteobacteria | Burkholderiales | Burkholderiaceae | Paraburkholderia |
| Paraburkholderia megapolitana | pmeg | 2 | Bacteria | Pseudomonadota | Betaproteobacteria | Burkholderiales | Burkholderiaceae | Paraburkholderia |
| Paraburkholderia pallida | ppai | 7 | Bacteria | Pseudomonadota | Betaproteobacteria | Burkholderiales | Burkholderiaceae | Paraburkholderia |
| Paraburkholderia phenoliruptrix | bpx | 4 | Bacteria | Pseudomonadota | Betaproteobacteria | Burkholderiales | Burkholderiaceae | Paraburkholderia |
| Paraburkholderia phymatum | bph | 2 | Bacteria | Pseudomonadota | Betaproteobacteria | Burkholderiales | Burkholderiaceae | Paraburkholderia |
| Paraburkholderia phytofirmans OLGA172 | buz | 2 | Bacteria | Pseudomonadota | Betaproteobacteria | Burkholderiales | Burkholderiaceae | Paraburkholderia |
| Paraburkholderia phytofirmans PsJN | bpy | 2 | Bacteria | Pseudomonadota | Betaproteobacteria | Burkholderiales | Burkholderiaceae | Paraburkholderia |
| Paraburkholderia sabiae | psaa | 2 | Bacteria | Pseudomonadota | Betaproteobacteria | Burkholderiales | Burkholderiaceae | Paraburkholderia |
| Paraburkholderia sp. SOS3 | para | 1 | Bacteria | Pseudomonadota | Betaproteobacteria | Burkholderiales | Burkholderiaceae | Paraburkholderia |
| Paraburkholderia sprentiae | pspw | 1 | Bacteria | Pseudomonadota | Betaproteobacteria | Burkholderiales | Burkholderiaceae | Paraburkholderia |
| Paraburkholderia terrae | pter | 2 | Bacteria | Pseudomonadota | Betaproteobacteria | Burkholderiales | Burkholderiaceae | Paraburkholderia |
| Paraburkholderia terricola | pts | 1 | Bacteria | Pseudomonadota | Betaproteobacteria | Burkholderiales | Burkholderiaceae | Paraburkholderia |
| Paraburkholderia tropica | ptro | 5 | Bacteria | Pseudomonadota | Betaproteobacteria | Burkholderiales | Burkholderiaceae | Paraburkholderia |
| Paraburkholderia xenovorans LB400 | bxb | 2 | Bacteria | Pseudomonadota | Betaproteobacteria | Burkholderiales | Burkholderiaceae | Paraburkholderia |
| Paraburkholderia xenovorans LB400 | bxe | 2 | Bacteria | Pseudomonadota | Betaproteobacteria | Burkholderiales | Burkholderiaceae | Paraburkholderia |
| Paucimonas lemoignei | plg | 1 | Bacteria | Pseudomonadota | Betaproteobacteria | Burkholderiales | Burkholderiaceae | Paucimonas |
| Polynucleobacter wuianus | pwu | 1 | Bacteria | Pseudomonadota | Betaproteobacteria | Burkholderiales | Burkholderiaceae | Polynucleobacter |
| Ralstonia insidiosa | rin | 2 | Bacteria | Pseudomonadota | Betaproteobacteria | Burkholderiales | Burkholderiaceae | Ralstonia |
| Ralstonia mannitolilytica | rmn | 1 | Bacteria | Pseudomonadota | Betaproteobacteria | Burkholderiales | Burkholderiaceae | Ralstonia |
| Ralstonia pickettii 12D | rpf | 1 | Bacteria | Pseudomonadota | Betaproteobacteria | Burkholderiales | Burkholderiaceae | Ralstonia |
| Ralstonia pickettii 12J | rpi | 1 | Bacteria | Pseudomonadota | Betaproteobacteria | Burkholderiales | Burkholderiaceae | Ralstonia |
| Ralstonia pickettii DTP0602 | rpj | 5 | Bacteria | Pseudomonadota | Betaproteobacteria | Burkholderiales | Burkholderiaceae | Ralstonia |
| Ralstonia pseudosolanacearum GMI1000 | rso | 1 | Bacteria | Pseudomonadota | Betaproteobacteria | Burkholderiales | Burkholderiaceae | Ralstonia |
| Ralstonia solanacearum CFBP2957 | rsc | 2 | Bacteria | Pseudomonadota | Betaproteobacteria | Burkholderiales | Burkholderiaceae | Ralstonia |
| Ralstonia solanacearum CMR15 | rsm | 3 | Bacteria | Pseudomonadota | Betaproteobacteria | Burkholderiales | Burkholderiaceae | Ralstonia |
| Ralstonia solanacearum Po82 | rsn | 3 | Bacteria | Pseudomonadota | Betaproteobacteria | Burkholderiales | Burkholderiaceae | Ralstonia |
| Ralstonia solanacearum PSI07 | rsl | 3 | Bacteria | Pseudomonadota | Betaproteobacteria | Burkholderiales | Burkholderiaceae | Ralstonia |
| Ralstonia syzygii subsp. celebesensis | rsg | 1 | Bacteria | Pseudomonadota | Betaproteobacteria | Burkholderiales | Burkholderiaceae | Ralstonia |
| Ralstonia wenshanensis | rwe | 1 | Bacteria | Pseudomonadota | Betaproteobacteria | Burkholderiales | Burkholderiaceae | Ralstonia |
| Trinickia caryophylli | tcar | 1 | Bacteria | Pseudomonadota | Betaproteobacteria | Burkholderiales | Burkholderiaceae | Trinickia |
| Trinickia violacea | tvl | 2 | Bacteria | Pseudomonadota | Betaproteobacteria | Burkholderiales | Burkholderiaceae | Trinickia |
| Thiomonas arsenitoxydans | thi | 1 | Bacteria | Pseudomonadota | Betaproteobacteria | Burkholderiales | Burkholderiales genera incertae sedis | Thiomonas |
| Xylophilus rhododendri | xyk | 1 | Bacteria | Pseudomonadota | Betaproteobacteria | Burkholderiales | Burkholderiales genera incertae sedis | Xylophilus |
| Acidovorax carolinensis NA2 | acid | 1 | Bacteria | Pseudomonadota | Betaproteobacteria | Burkholderiales | Comamonadaceae | Acidovorax |
| Acidovorax sp. 1608163 | acio | 1 | Bacteria | Pseudomonadota | Betaproteobacteria | Burkholderiales | Comamonadaceae | Acidovorax |
| Acidovorax sp. DW039 | aciq | 2 | Bacteria | Pseudomonadota | Betaproteobacteria | Burkholderiales | Comamonadaceae | Acidovorax |
| Acidovorax sp. KKS102 | ack | 1 | Bacteria | Pseudomonadota | Betaproteobacteria | Burkholderiales | Comamonadaceae | Acidovorax |
| Acidovorax sp. RAC01 | acra | 1 | Bacteria | Pseudomonadota | Betaproteobacteria | Burkholderiales | Comamonadaceae | Acidovorax |
| Alicycliphilus denitrificans BC | adn | 4 | Bacteria | Pseudomonadota | Betaproteobacteria | Burkholderiales | Comamonadaceae | Alicycliphilus |
| Alicycliphilus denitrificans K601 | adk | 4 | Bacteria | Pseudomonadota | Betaproteobacteria | Burkholderiales | Comamonadaceae | Alicycliphilus |
| Comamonas antarctica | aant | 2 | Bacteria | Pseudomonadota | Betaproteobacteria | Burkholderiales | Comamonadaceae | Comamonas |
| Comamonas endophytica | cenp | 2 | Bacteria | Pseudomonadota | Betaproteobacteria | Burkholderiales | Comamonadaceae | Comamonas |
| Comamonas odontotermitis | codo | 2 | Bacteria | Pseudomonadota | Betaproteobacteria | Burkholderiales | Comamonadaceae | Comamonas |
| Comamonas resistens | crj | 1 | Bacteria | Pseudomonadota | Betaproteobacteria | Burkholderiales | Comamonadaceae | Comamonas |
| Comamonas serinivorans | cser | 1 | Bacteria | Pseudomonadota | Betaproteobacteria | Burkholderiales | Comamonadaceae | Comamonas |
| Comamonas testosteroni TK102 | ctes | 3 | Bacteria | Pseudomonadota | Betaproteobacteria | Burkholderiales | Comamonadaceae | Comamonas |
| Comamonas thiooxydans | ctt | 2 | Bacteria | Pseudomonadota | Betaproteobacteria | Burkholderiales | Comamonadaceae | Comamonas |
| Delftia acidovorans | dac | 3 | Bacteria | Pseudomonadota | Betaproteobacteria | Burkholderiales | Comamonadaceae | Delftia |
| Delftia lacustris | dla | 3 | Bacteria | Pseudomonadota | Betaproteobacteria | Burkholderiales | Comamonadaceae | Delftia |
| Delftia sp. Cs1-4 | del | 3 | Bacteria | Pseudomonadota | Betaproteobacteria | Burkholderiales | Comamonadaceae | Delftia |
| Delftia sp. HK171 | dhk | 2 | Bacteria | Pseudomonadota | Betaproteobacteria | Burkholderiales | Comamonadaceae | Delftia |
| Delftia tsuruhatensis | dts | 2 | Bacteria | Pseudomonadota | Betaproteobacteria | Burkholderiales | Comamonadaceae | Delftia |
| Diaphorobacter aerolatus | daer | 2 | Bacteria | Pseudomonadota | Betaproteobacteria | Burkholderiales | Comamonadaceae | Diaphorobacter |
| Diaphorobacter limosus | dls | 1 | Bacteria | Pseudomonadota | Betaproteobacteria | Burkholderiales | Comamonadaceae | Diaphorobacter |
| Hydrogenophaga crocea | hcz | 1 | Bacteria | Pseudomonadota | Betaproteobacteria | Burkholderiales | Comamonadaceae | Hydrogenophaga |
| Hydrogenophaga pseudoflava | hpse | 1 | Bacteria | Pseudomonadota | Betaproteobacteria | Burkholderiales | Comamonadaceae | Hydrogenophaga |
| Hydrogenophaga sp. RAC07 | hyr | 1 | Bacteria | Pseudomonadota | Betaproteobacteria | Burkholderiales | Comamonadaceae | Hydrogenophaga |
| Hydrogenophaga taeniospiralis | htn | 1 | Bacteria | Pseudomonadota | Betaproteobacteria | Burkholderiales | Comamonadaceae | Hydrogenophaga |
| Limnohabitans sp. 103DPR2 | lim | 3 | Bacteria | Pseudomonadota | Betaproteobacteria | Burkholderiales | Comamonadaceae | Limnohabitans |
| Limnohabitans sp. 63ED37-2 | lih | 3 | Bacteria | Pseudomonadota | Betaproteobacteria | Burkholderiales | Comamonadaceae | Limnohabitans |
| Paenacidovorax monticola | amon | 1 | Bacteria | Pseudomonadota | Betaproteobacteria | Burkholderiales | Comamonadaceae | Paenacidovorax |
| Paracidovorax citrulli | aav | 1 | Bacteria | Pseudomonadota | Betaproteobacteria | Burkholderiales | Comamonadaceae | Paracidovorax |
| Polaromonas sp. JS666 | pol | 1 | Bacteria | Pseudomonadota | Betaproteobacteria | Burkholderiales | Comamonadaceae | Polaromonas |
| Polaromonas vacuolata | pvac | 2 | Bacteria | Pseudomonadota | Betaproteobacteria | Burkholderiales | Comamonadaceae | Polaromonas |
| Ramlibacter tataouinensis | rta | 2 | Bacteria | Pseudomonadota | Betaproteobacteria | Burkholderiales | Comamonadaceae | Ramlibacter |
| Rhodoferax koreensis | rhy | 1 | Bacteria | Pseudomonadota | Betaproteobacteria | Burkholderiales | Comamonadaceae | Rhodoferax |
| Rhodoferax lithotrophicus | rlh | 1 | Bacteria | Pseudomonadota | Betaproteobacteria | Burkholderiales | Comamonadaceae | Rhodoferax |
| Rhodoferax mekongensis | rmk | 1 | Bacteria | Pseudomonadota | Betaproteobacteria | Burkholderiales | Comamonadaceae | Rhodoferax |
| Rhodoferax sediminis CHu59-6-5 | rhf | 2 | Bacteria | Pseudomonadota | Betaproteobacteria | Burkholderiales | Comamonadaceae | Rhodoferax |
| Serpentinimonas raichei | cbaa | 1 | Bacteria | Pseudomonadota | Betaproteobacteria | Burkholderiales | Comamonadaceae | Serpentinimonas |
| Variovorax boronicumulans | vbo | 4 | Bacteria | Pseudomonadota | Betaproteobacteria | Burkholderiales | Comamonadaceae | Variovorax |
| Variovorax paradoxus B4 | vpd | 2 | Bacteria | Pseudomonadota | Betaproteobacteria | Burkholderiales | Comamonadaceae | Variovorax |
| Variovorax paradoxus EPS | vpe | 2 | Bacteria | Pseudomonadota | Betaproteobacteria | Burkholderiales | Comamonadaceae | Variovorax |
| Variovorax paradoxus S110 | vap | 1 | Bacteria | Pseudomonadota | Betaproteobacteria | Burkholderiales | Comamonadaceae | Variovorax |
| Variovorax sp. 38R | vav | 4 | Bacteria | Pseudomonadota | Betaproteobacteria | Burkholderiales | Comamonadaceae | Variovorax |
| Variovorax sp. PAMC26660 | vaz | 3 | Bacteria | Pseudomonadota | Betaproteobacteria | Burkholderiales | Comamonadaceae | Variovorax |
| Variovorax sp. PMC12 | vam | 2 | Bacteria | Pseudomonadota | Betaproteobacteria | Burkholderiales | Comamonadaceae | Variovorax |
| Verminephrobacter eiseniae | vei | 1 | Bacteria | Pseudomonadota | Betaproteobacteria | Burkholderiales | Comamonadaceae | Verminephrobacter |
| Collimonas arenae | care | 2 | Bacteria | Pseudomonadota | Betaproteobacteria | Burkholderiales | Oxalobacteraceae | Collimonas |
| Herbaspirillum frisingense | hfr | 1 | Bacteria | Pseudomonadota | Betaproteobacteria | Burkholderiales | Oxalobacteraceae | Herbaspirillum |
| Herbaspirillum hiltneri | hht | 1 | Bacteria | Pseudomonadota | Betaproteobacteria | Burkholderiales | Oxalobacteraceae | Herbaspirillum |
| Herbaspirillum huttiense | hhf | 2 | Bacteria | Pseudomonadota | Betaproteobacteria | Burkholderiales | Oxalobacteraceae | Herbaspirillum |
| Herbaspirillum rubrisubalbicans | hrb | 1 | Bacteria | Pseudomonadota | Betaproteobacteria | Burkholderiales | Oxalobacteraceae | Herbaspirillum |
| Herbaspirillum seropedicae SmR1 | hse | 2 | Bacteria | Pseudomonadota | Betaproteobacteria | Burkholderiales | Oxalobacteraceae | Herbaspirillum |
| Herbaspirillum seropedicae Z67 | hsz | 1 | Bacteria | Pseudomonadota | Betaproteobacteria | Burkholderiales | Oxalobacteraceae | Herbaspirillum |
| Herbaspirillum sp. DW155 | hew | 1 | Bacteria | Pseudomonadota | Betaproteobacteria | Burkholderiales | Oxalobacteraceae | Herbaspirillum |
| Herbaspirillum sp. meg3 | hee | 1 | Bacteria | Pseudomonadota | Betaproteobacteria | Burkholderiales | Oxalobacteraceae | Herbaspirillum |
| Herminiimonas arsenicoxydans | har | 1 | Bacteria | Pseudomonadota | Betaproteobacteria | Burkholderiales | Oxalobacteraceae | Herminiimonas |
| Janthinobacterium sp. HH102 | jah | 1 | Bacteria | Pseudomonadota | Betaproteobacteria | Burkholderiales | Oxalobacteraceae | Janthinobacterium |
| Janthinobacterium sp. Marseille | mms | 2 | Bacteria | Pseudomonadota | Betaproteobacteria | Burkholderiales | Oxalobacteraceae | Janthinobacterium |
| Aquincola tertiaricarbonis | ater | 2 | Bacteria | Pseudomonadota | Betaproteobacteria | Burkholderiales | Sphaerotilaceae | Aquincola |
| Methylibium petroleiphilum | mpt | 1 | Bacteria | Pseudomonadota | Betaproteobacteria | Burkholderiales | Sphaerotilaceae | Methylibium |
| Sphaerotilus microaerophilus | smio | 3 | Bacteria | Pseudomonadota | Betaproteobacteria | Burkholderiales | Sphaerotilaceae | Sphaerotilus |
| Sphaerotilus sulfidivorans | snn | 1 | Bacteria | Pseudomonadota | Betaproteobacteria | Burkholderiales | Sphaerotilaceae | Sphaerotilus |
| Leptothrix cholodnii | lch | 1 | Bacteria | Pseudomonadota | Betaproteobacteria | Burkholderiales | Sphaerotilaceae | Leptothrix |
| Sutterella megalosphaeroides | sutt | 1 | Bacteria | Pseudomonadota | Betaproteobacteria | Burkholderiales | Sutterellaceae | Sutterella |
| Aquitalea magnusonii | amah | 1 | Bacteria | Pseudomonadota | Betaproteobacteria | Neisseriales | Chromobacteriaceae | Aquitalea |
| Aquitalea sp. USM4 | aqs | 1 | Bacteria | Pseudomonadota | Betaproteobacteria | Neisseriales | Chromobacteriaceae | Aquitalea |
| Chromobacterium haemolyticum | chae | 2 | Bacteria | Pseudomonadota | Betaproteobacteria | Neisseriales | Chromobacteriaceae | Chromobacterium |
| Chromobacterium paludis | chrm | 2 | Bacteria | Pseudomonadota | Betaproteobacteria | Neisseriales | Chromobacteriaceae | Chromobacterium |
| Chromobacterium phragmitis IIBBL 112-1 | chri | 1 | Bacteria | Pseudomonadota | Betaproteobacteria | Neisseriales | Chromobacteriaceae | Chromobacterium |
| Chromobacterium phragmitis IIBBL 274-1 | chrb | 1 | Bacteria | Pseudomonadota | Betaproteobacteria | Neisseriales | Chromobacteriaceae | Chromobacterium |
| Chromobacterium rhizoryzae | crz | 2 | Bacteria | Pseudomonadota | Betaproteobacteria | Neisseriales | Chromobacteriaceae | Chromobacterium |
| Chromobacterium sp. ATCC 53434 | chro | 1 | Bacteria | Pseudomonadota | Betaproteobacteria | Neisseriales | Chromobacteriaceae | Chromobacterium |
| Chromobacterium vaccinii | cvc | 1 | Bacteria | Pseudomonadota | Betaproteobacteria | Neisseriales | Chromobacteriaceae | Chromobacterium |
| Chromobacterium violaceum | cvi | 2 | Bacteria | Pseudomonadota | Betaproteobacteria | Neisseriales | Chromobacteriaceae | Chromobacterium |
| Pseudogulbenkiania sp. NH8B | pse | 2 | Bacteria | Pseudomonadota | Betaproteobacteria | Neisseriales | Chromobacteriaceae | Pseudogulbenkiania |
| Alysiella crassa | acrs | 1 | Bacteria | Pseudomonadota | Betaproteobacteria | Neisseriales | Neisseriaceae | Alysiella |
| Alysiella filiformis | aff | 1 | Bacteria | Pseudomonadota | Betaproteobacteria | Neisseriales | Neisseriaceae | Alysiella |
| Neisseria brasiliensis | nbl | 1 | Bacteria | Pseudomonadota | Betaproteobacteria | Neisseriales | Neisseriaceae | Neisseria |
| Neisseria chenwenguii | nei | 1 | Bacteria | Pseudomonadota | Betaproteobacteria | Neisseriales | Neisseriaceae | Neisseria |
| Neisseria weixii | nwx | 1 | Bacteria | Pseudomonadota | Betaproteobacteria | Neisseriales | Neisseriaceae | Neisseria |
| Snodgrassella alvi | salv | 1 | Bacteria | Pseudomonadota | Betaproteobacteria | Neisseriales | Neisseriaceae | Snodgrassella |
| Snodgrassella communis | scom | 1 | Bacteria | Pseudomonadota | Betaproteobacteria | Neisseriales | Neisseriaceae | Snodgrassella |
| Vitreoscilla filiformis | vff | 1 | Bacteria | Pseudomonadota | Betaproteobacteria | Neisseriales | Neisseriaceae | Vitreoscilla |
| Vitreoscilla massiliensis | vms | 1 | Bacteria | Pseudomonadota | Betaproteobacteria | Neisseriales | Neisseriaceae | Vitreoscilla |
| Aromatoleum petrolei | apet | 2 | Bacteria | Pseudomonadota | Betaproteobacteria | Rhodocyclales | Rhodocyclaceae | Aromatoleum |
| Oryzomicrobium terrae | otr | 3 | Bacteria | Pseudomonadota | Betaproteobacteria | Rhodocyclales | Rhodocyclaceae | Oryzomicrobium |
| Azoarcus olearius BH72 | azo | 2 | Bacteria | Pseudomonadota | Betaproteobacteria | Rhodocyclales | Zoogloeaceae | Azoarcus |
| Azoarcus olearius DQS-4 | aoa | 2 | Bacteria | Pseudomonadota | Betaproteobacteria | Rhodocyclales | Zoogloeaceae | Azoarcus |
| Azoarcus sp. CIB | azi | 2 | Bacteria | Pseudomonadota | Betaproteobacteria | Rhodocyclales | Zoogloeaceae | Azoarcus |
| Azoarcus sp. DN11 | azd | 2 | Bacteria | Pseudomonadota | Betaproteobacteria | Rhodocyclales | Zoogloeaceae | Azoarcus |
| Azoarcus sp. KH32C | aza | 2 | Bacteria | Pseudomonadota | Betaproteobacteria | Rhodocyclales | Zoogloeaceae | Azoarcus |
| Thauera aminoaromatica | tmz | 2 | Bacteria | Pseudomonadota | Betaproteobacteria | Rhodocyclales | Zoogloeaceae | Thauera |
| Thauera humireducens Piv1 | thau | 2 | Bacteria | Pseudomonadota | Betaproteobacteria | Rhodocyclales | Zoogloeaceae | Thauera |
| Thauera humireducens SgZ-1 | thu | 1 | Bacteria | Pseudomonadota | Betaproteobacteria | Rhodocyclales | Zoogloeaceae | Thauera |
| Thauera sp. GDN1 | thag | 2 | Bacteria | Pseudomonadota | Betaproteobacteria | Rhodocyclales | Zoogloeaceae | Thauera |
| Oceanimonas pelagia | ope | 1 | Bacteria | Pseudomonadota | Gammaproteobacteria | Aeromonadales | Aeromonadaceae | Oceanimonas |
| Oceanimonas sp. GK1 | oce | 1 | Bacteria | Pseudomonadota | Gammaproteobacteria | Aeromonadales | Aeromonadaceae | Oceanimonas |
| Paraglaciecola mesophila | pmes | 1 | Bacteria | Pseudomonadota | Gammaproteobacteria | Alteromonadales | Alteromonadaceae | Paraglaciecola |
| Pseudoalteromonas agarivorans | paga | 1 | Bacteria | Pseudomonadota | Gammaproteobacteria | Alteromonadales | Pseudoalteromonadaceae | Pseudoalteromonas |
| Pseudoalteromonas arctica | part | 1 | Bacteria | Pseudomonadota | Gammaproteobacteria | Alteromonadales | Pseudoalteromonadaceae | Pseudoalteromonas |
| Pseudoalteromonas espejiana | pea | 1 | Bacteria | Pseudomonadota | Gammaproteobacteria | Alteromonadales | Pseudoalteromonadaceae | Pseudoalteromonas |
| Pseudoalteromonas nigrifaciens | png | 1 | Bacteria | Pseudomonadota | Gammaproteobacteria | Alteromonadales | Pseudoalteromonadaceae | Pseudoalteromonas |
| Pseudoalteromonas spongiae | pspo | 1 | Bacteria | Pseudomonadota | Gammaproteobacteria | Alteromonadales | Pseudoalteromonadaceae | Pseudoalteromonas |
| Pseudoalteromonas tetraodonis | ptd | 1 | Bacteria | Pseudomonadota | Gammaproteobacteria | Alteromonadales | Pseudoalteromonadaceae | Pseudoalteromonas |
| Pseudoalteromonas translucida KMM 520 | ptn | 1 | Bacteria | Pseudomonadota | Gammaproteobacteria | Alteromonadales | Pseudoalteromonadaceae | Pseudoalteromonas |
| Pseudoalteromonas tunicata | ptu | 1 | Bacteria | Pseudomonadota | Gammaproteobacteria | Alteromonadales | Pseudoalteromonadaceae | Pseudoalteromonas |
| Ignatzschineria larvae | ilv | 2 | Bacteria | Pseudomonadota | Gammaproteobacteria | Cardiobacteriales | Ignatzschineriaceae | Ignatzschineria |
| Ignatzschineria rhizosphaerae | ign | 2 | Bacteria | Pseudomonadota | Gammaproteobacteria | Cardiobacteriales | Ignatzschineriaceae | Ignatzschineria |
| Wohlfahrtiimonas chitiniclastica | wcn | 2 | Bacteria | Pseudomonadota | Gammaproteobacteria | Cardiobacteriales | Ignatzschineriaceae | Wohlfahrtiimonas |
| Marinagarivorans cellulosilyticus | marq | 1 | Bacteria | Pseudomonadota | Gammaproteobacteria | Cellvibrionales | Cellvibrionaceae | Marinagarivorans |
| Teredinibacter turnerae | ttu | 1 | Bacteria | Pseudomonadota | Gammaproteobacteria | Cellvibrionales | Cellvibrionaceae | Teredinibacter |
| Sodalis praecaptivus | sod | 1 | Bacteria | Pseudomonadota | Gammaproteobacteria | Enterobacterales | Bruguierivoracaceae | Sodalis |
| Leminorella grimontii | lgb | 1 | Bacteria | Pseudomonadota | Gammaproteobacteria | Enterobacterales | Budviciaceae | Leminorella |
| Leminorella richardii | lri | 1 | Bacteria | Pseudomonadota | Gammaproteobacteria | Enterobacterales | Budviciaceae | Leminorella |
| Limnobaculum parvum | lpv | 2 | Bacteria | Pseudomonadota | Gammaproteobacteria | Enterobacterales | Budviciaceae | Limnobaculum |
| Limnobaculum zhutongyuii | prag | 1 | Bacteria | Pseudomonadota | Gammaproteobacteria | Enterobacterales | Budviciaceae | Limnobaculum |
| Pragia fontium | pfq | 2 | Bacteria | Pseudomonadota | Gammaproteobacteria | Enterobacterales | Budviciaceae | Pragia |
| Atlantibacter hermannii | ahn | 1 | Bacteria | Pseudomonadota | Gammaproteobacteria | Enterobacterales | Enterobacteriaceae | Atlantibacter |
| Buttiauxella agrestis | bage | 1 | Bacteria | Pseudomonadota | Gammaproteobacteria | Enterobacterales | Enterobacteriaceae | Buttiauxella |
| Buttiauxella ferragutiae | bft | 1 | Bacteria | Pseudomonadota | Gammaproteobacteria | Enterobacterales | Enterobacteriaceae | Buttiauxella |
| Buttiauxella selenatireducens | bsel | 1 | Bacteria | Pseudomonadota | Gammaproteobacteria | Enterobacterales | Enterobacteriaceae | Buttiauxella |
| Buttiauxella sp. 3AFRM03 | buf | 1 | Bacteria | Pseudomonadota | Gammaproteobacteria | Enterobacterales | Enterobacteriaceae | Buttiauxella |
| Cedecea neteri M006 | cem | 1 | Bacteria | Pseudomonadota | Gammaproteobacteria | Enterobacterales | Enterobacteriaceae | Cedecea |
| Cedecea neteri ND14a | cen | 1 | Bacteria | Pseudomonadota | Gammaproteobacteria | Enterobacterales | Enterobacteriaceae | Cedecea |
| Citrobacter rodentium | cro | 1 | Bacteria | Pseudomonadota | Gammaproteobacteria | Enterobacterales | Enterobacteriaceae | Citrobacter |
| Enterobacter cloacae ECNIH2 | ecle | 1 | Bacteria | Pseudomonadota | Gammaproteobacteria | Enterobacterales | Enterobacteriaceae | Enterobacter |
| Enterobacter cloacae ECNIH5 | ecli | 1 | Bacteria | Pseudomonadota | Gammaproteobacteria | Enterobacterales | Enterobacteriaceae | Enterobacter |
| Enterobacter dykesii | edy | 1 | Bacteria | Pseudomonadota | Gammaproteobacteria | Enterobacterales | Enterobacteriaceae | Enterobacter |
| Enterobacter hormaechei CAV1176 | ehm | 1 | Bacteria | Pseudomonadota | Gammaproteobacteria | Enterobacterales | Enterobacteriaceae | Enterobacter |
| Enterobacter hormaechei NCTC 9394 | eclo | 1 | Bacteria | Pseudomonadota | Gammaproteobacteria | Enterobacterales | Enterobacteriaceae | Enterobacter |
| Enterobacter hormaechei subsp. hoffmannii ECNIH3 | ecla | 1 | Bacteria | Pseudomonadota | Gammaproteobacteria | Enterobacterales | Enterobacteriaceae | Enterobacter |
| Enterobacter hormaechei subsp. hoffmannii ECR091 | eclc | 1 | Bacteria | Pseudomonadota | Gammaproteobacteria | Enterobacterales | Enterobacteriaceae | Enterobacter |
| Enterobacter hormaechei subsp. hormaechei | eclz | 1 | Bacteria | Pseudomonadota | Gammaproteobacteria | Enterobacterales | Enterobacteriaceae | Enterobacter |
| Enterobacter hormaechei subsp. steigerwaltii | ecly | 1 | Bacteria | Pseudomonadota | Gammaproteobacteria | Enterobacterales | Enterobacteriaceae | Enterobacter |
| Enterobacter hormaechei subsp. xiangfangensis | exf | 1 | Bacteria | Pseudomonadota | Gammaproteobacteria | Enterobacterales | Enterobacteriaceae | Enterobacter |
| Enterobacter hormaechei subsp. xiangfangensis 34399 | eclx | 1 | Bacteria | Pseudomonadota | Gammaproteobacteria | Enterobacterales | Enterobacteriaceae | Enterobacter |
| Enterobacter lignolyticus G5 | kle | 1 | Bacteria | Pseudomonadota | Gammaproteobacteria | Enterobacterales | Enterobacteriaceae | Enterobacter |
| Enterobacter pasteurii | ept | 1 | Bacteria | Pseudomonadota | Gammaproteobacteria | Enterobacterales | Enterobacteriaceae | Enterobacter |
| Enterobacter sp. AN-K1 | enk | 1 | Bacteria | Pseudomonadota | Gammaproteobacteria | Enterobacterales | Enterobacteriaceae | Enterobacter |
| Enterobacter sp. DSM 30060 | ens | 1 | Bacteria | Pseudomonadota | Gammaproteobacteria | Enterobacterales | Enterobacteriaceae | Enterobacter |
| Enterobacter sp. N18-03635 | enb | 1 | Bacteria | Pseudomonadota | Gammaproteobacteria | Enterobacterales | Enterobacteriaceae | Enterobacter |
| Enterobacteriaceae bacterium bta3-1 | ebb | 1 | Bacteria | Pseudomonadota | Gammaproteobacteria | Enterobacterales | Enterobacteriaceae | Enterobacteriaceae |
| Enterobacteriaceae bacterium S05 | ebu | 2 | Bacteria | Pseudomonadota | Gammaproteobacteria | Enterobacterales | Enterobacteriaceae | Enterobacteriaceae |
| Escherichia albertii | eal | 1 | Bacteria | Pseudomonadota | Gammaproteobacteria | Enterobacterales | Enterobacteriaceae | Escherichia |
| Escherichia coli ABU 83972 | eab | 1 | Bacteria | Pseudomonadota | Gammaproteobacteria | Enterobacterales | Enterobacteriaceae | Escherichia |
| Escherichia coli APEC O1 (APEC) | ecv | 1 | Bacteria | Pseudomonadota | Gammaproteobacteria | Enterobacterales | Enterobacteriaceae | Escherichia |
| Escherichia coli clone D i14 | elc | 1 | Bacteria | Pseudomonadota | Gammaproteobacteria | Enterobacterales | Enterobacteriaceae | Escherichia |
| Escherichia coli clone D i2 | eld | 1 | Bacteria | Pseudomonadota | Gammaproteobacteria | Enterobacterales | Enterobacteriaceae | Escherichia |
| Escherichia coli JJ1886 | ecoj | 1 | Bacteria | Pseudomonadota | Gammaproteobacteria | Enterobacterales | Enterobacteriaceae | Escherichia |
| Escherichia coli LF82 | elf | 1 | Bacteria | Pseudomonadota | Gammaproteobacteria | Enterobacterales | Enterobacteriaceae | Escherichia |
| Escherichia coli NA114 (UPEC) | ena | 1 | Bacteria | Pseudomonadota | Gammaproteobacteria | Enterobacterales | Enterobacteriaceae | Escherichia |
| Escherichia coli O127:H6 E2348/69 (EPEC) | ecg | 1 | Bacteria | Pseudomonadota | Gammaproteobacteria | Enterobacterales | Enterobacteriaceae | Escherichia |
| Escherichia coli O145:H28 RM13514 (EHEC) | ecoo | 1 | Bacteria | Pseudomonadota | Gammaproteobacteria | Enterobacterales | Enterobacteriaceae | Escherichia |
| Escherichia coli O145:H28 RM13516 (EHEC) | ecoh | 1 | Bacteria | Pseudomonadota | Gammaproteobacteria | Enterobacterales | Enterobacteriaceae | Escherichia |
| Escherichia coli O150:H5 SE15 (commensal) | ese | 1 | Bacteria | Pseudomonadota | Gammaproteobacteria | Enterobacterales | Enterobacteriaceae | Escherichia |
| Escherichia coli O157:H7 EC4115 (EHEC) | ecf | 1 | Bacteria | Pseudomonadota | Gammaproteobacteria | Enterobacterales | Enterobacteriaceae | Escherichia |
| Escherichia coli O157:H7 EDL933 (EHEC) | ece | 1 | Bacteria | Pseudomonadota | Gammaproteobacteria | Enterobacterales | Enterobacteriaceae | Escherichia |
| Escherichia coli O157:H7 Sakai (EHEC) | ecs | 1 | Bacteria | Pseudomonadota | Gammaproteobacteria | Enterobacterales | Enterobacteriaceae | Escherichia |
| Escherichia coli O157:H7 TW14359 (EHEC) | etw | 1 | Bacteria | Pseudomonadota | Gammaproteobacteria | Enterobacterales | Enterobacteriaceae | Escherichia |
| Escherichia coli O157:H7 Xuzhou21 (EHEC) | elx | 1 | Bacteria | Pseudomonadota | Gammaproteobacteria | Enterobacterales | Enterobacteriaceae | Escherichia |
| Escherichia coli O18:K1 PMV-1 (ExPEC) | ecoi | 1 | Bacteria | Pseudomonadota | Gammaproteobacteria | Enterobacterales | Enterobacteriaceae | Escherichia |
| Escherichia coli O18:K1:H7 IHE3034 (ExPEC) | eih | 1 | Bacteria | Pseudomonadota | Gammaproteobacteria | Enterobacterales | Enterobacteriaceae | Escherichia |
| Escherichia coli O18:K1:H7 UTI89 (UPEC) | eci | 1 | Bacteria | Pseudomonadota | Gammaproteobacteria | Enterobacterales | Enterobacteriaceae | Escherichia |
| Escherichia coli O25b:K100:H4-ST131 EC958 (UPEC) | ecos | 1 | Bacteria | Pseudomonadota | Gammaproteobacteria | Enterobacterales | Enterobacteriaceae | Escherichia |
| Escherichia coli O45:K1:H7 S88 (ExPEC) | ecz | 1 | Bacteria | Pseudomonadota | Gammaproteobacteria | Enterobacterales | Enterobacteriaceae | Escherichia |
| Escherichia coli O55:H7 CB9615 (EPEC) | eok | 1 | Bacteria | Pseudomonadota | Gammaproteobacteria | Enterobacterales | Enterobacteriaceae | Escherichia |
| Escherichia coli O55:H7 RM12579 (EPEC) | elr | 1 | Bacteria | Pseudomonadota | Gammaproteobacteria | Enterobacterales | Enterobacteriaceae | Escherichia |
| Escherichia coli O6:K15:H31 536 (UPEC) | ecp | 1 | Bacteria | Pseudomonadota | Gammaproteobacteria | Enterobacterales | Enterobacteriaceae | Escherichia |
| Escherichia coli O6:K2:H1 CFT073 (UPEC) | ecc | 1 | Bacteria | Pseudomonadota | Gammaproteobacteria | Enterobacterales | Enterobacteriaceae | Escherichia |
| Escherichia coli O81 ED1a (commensal) | ecq | 1 | Bacteria | Pseudomonadota | Gammaproteobacteria | Enterobacterales | Enterobacteriaceae | Escherichia |
| Escherichia coli O83:H1 NRG 857C (AIEC) | eln | 1 | Bacteria | Pseudomonadota | Gammaproteobacteria | Enterobacterales | Enterobacteriaceae | Escherichia |
| Escherichia coli UM146 | elu | 1 | Bacteria | Pseudomonadota | Gammaproteobacteria | Enterobacterales | Enterobacteriaceae | Escherichia |
| Escherichia fergusonii | efe | 1 | Bacteria | Pseudomonadota | Gammaproteobacteria | Enterobacterales | Enterobacteriaceae | Escherichia |
| Escherichia marmotae | ema | 1 | Bacteria | Pseudomonadota | Gammaproteobacteria | Enterobacterales | Enterobacteriaceae | Escherichia |
| Escherichia ruysiae | eruy | 1 | Bacteria | Pseudomonadota | Gammaproteobacteria | Enterobacterales | Enterobacteriaceae | Escherichia |
| Escherichia sp. E4742 | esz | 1 | Bacteria | Pseudomonadota | Gammaproteobacteria | Enterobacterales | Enterobacteriaceae | Escherichia |
| Jejuibacter calystegiae | izh | 1 | Bacteria | Pseudomonadota | Gammaproteobacteria | Enterobacterales | Enterobacteriaceae | Jejuibacter |
| Kluyvera ascorbata | kas | 1 | Bacteria | Pseudomonadota | Gammaproteobacteria | Enterobacterales | Enterobacteriaceae | Kluyvera |
| Kluyvera intermedia | kie | 1 | Bacteria | Pseudomonadota | Gammaproteobacteria | Enterobacterales | Enterobacteriaceae | Kluyvera |
| Kluyvera sp. CRP | klu | 1 | Bacteria | Pseudomonadota | Gammaproteobacteria | Enterobacterales | Enterobacteriaceae | Kluyvera |
| Leclercia adecarboxylata | lax | 1 | Bacteria | Pseudomonadota | Gammaproteobacteria | Enterobacterales | Enterobacteriaceae | Leclercia |
| Leclercia pneumoniae | lpnu | 1 | Bacteria | Pseudomonadota | Gammaproteobacteria | Enterobacterales | Enterobacteriaceae | Leclercia |
| Leclercia sp. Colony189 | ler | 1 | Bacteria | Pseudomonadota | Gammaproteobacteria | Enterobacterales | Enterobacteriaceae | Leclercia |
| Leclercia sp. J807 | lea | 1 | Bacteria | Pseudomonadota | Gammaproteobacteria | Enterobacterales | Enterobacteriaceae | Leclercia |
| Leclercia sp. LSNIH1 | lei | 1 | Bacteria | Pseudomonadota | Gammaproteobacteria | Enterobacterales | Enterobacteriaceae | Leclercia |
| Leclercia sp. LSNIH3 | leh | 1 | Bacteria | Pseudomonadota | Gammaproteobacteria | Enterobacterales | Enterobacteriaceae | Leclercia |
| Leclercia sp. W17 | lee | 1 | Bacteria | Pseudomonadota | Gammaproteobacteria | Enterobacterales | Enterobacteriaceae | Leclercia |
| Leclercia sp. W6 | ley | 1 | Bacteria | Pseudomonadota | Gammaproteobacteria | Enterobacterales | Enterobacteriaceae | Leclercia |
| Pluralibacter gergoviae | pge | 2 | Bacteria | Pseudomonadota | Gammaproteobacteria | Enterobacterales | Enterobacteriaceae | Pluralibacter |
| Scandinavium goeteborgense | sgoe | 1 | Bacteria | Pseudomonadota | Gammaproteobacteria | Enterobacterales | Enterobacteriaceae | Scandinavium |
| Shimwellia blattae | ebt | 1 | Bacteria | Pseudomonadota | Gammaproteobacteria | Enterobacterales | Enterobacteriaceae | Shimwellia |
| Mixta calida | pcd | 1 | Bacteria | Pseudomonadota | Gammaproteobacteria | Enterobacterales | Erwiniaceae | Mixta |
| Mixta gaviniae | pgz | 1 | Bacteria | Pseudomonadota | Gammaproteobacteria | Enterobacterales | Erwiniaceae | Mixta |
| Mixta hanseatica | mhan | 1 | Bacteria | Pseudomonadota | Gammaproteobacteria | Enterobacterales | Erwiniaceae | Mixta |
| Mixta intestinalis | mint | 1 | Bacteria | Pseudomonadota | Gammaproteobacteria | Enterobacterales | Erwiniaceae | Mixta |
| Mixta theicola | mthi | 2 | Bacteria | Pseudomonadota | Gammaproteobacteria | Enterobacterales | Erwiniaceae | Mixta |
| Pantoea alhagi | palh | 1 | Bacteria | Pseudomonadota | Gammaproteobacteria | Enterobacterales | Erwiniaceae | Pantoea |
| Pantoea ananatis AJ13355 | paj | 2 | Bacteria | Pseudomonadota | Gammaproteobacteria | Enterobacterales | Erwiniaceae | Pantoea |
| Pantoea ananatis LMG 5342 | plf | 2 | Bacteria | Pseudomonadota | Gammaproteobacteria | Enterobacterales | Erwiniaceae | Pantoea |
| Pantoea ananatis PA13 | paq | 2 | Bacteria | Pseudomonadota | Gammaproteobacteria | Enterobacterales | Erwiniaceae | Pantoea |
| Pantoea phytobeneficialis | ppho | 1 | Bacteria | Pseudomonadota | Gammaproteobacteria | Enterobacterales | Erwiniaceae | Pantoea |
| Pantoea sp. At-9b | pao | 1 | Bacteria | Pseudomonadota | Gammaproteobacteria | Enterobacterales | Erwiniaceae | Pantoea |
| Pantoea sp. PSNIH2 | panp | 1 | Bacteria | Pseudomonadota | Gammaproteobacteria | Enterobacterales | Erwiniaceae | Pantoea |
| Tatumella citrea | tci | 2 | Bacteria | Pseudomonadota | Gammaproteobacteria | Enterobacterales | Erwiniaceae | Tatumella |
| Hafnia alvei | hav | 2 | Bacteria | Pseudomonadota | Gammaproteobacteria | Enterobacterales | Hafniaceae | Hafnia |
| Hafnia paralvei | hpar | 1 | Bacteria | Pseudomonadota | Gammaproteobacteria | Enterobacterales | Hafniaceae | Hafnia |
| Obesumbacterium proteus | opo | 1 | Bacteria | Pseudomonadota | Gammaproteobacteria | Enterobacterales | Hafniaceae | Obesumbacterium |
| Chania multitudinisentens | sfo | 1 | Bacteria | Pseudomonadota | Gammaproteobacteria | Enterobacterales | Yersiniaceae | Chania |
| Ewingella americana | eame | 1 | Bacteria | Pseudomonadota | Gammaproteobacteria | Enterobacterales | Yersiniaceae | Ewingella |
| Rahnella aceris Y9602 | rah | 1 | Bacteria | Pseudomonadota | Gammaproteobacteria | Enterobacterales | Yersiniaceae | Rahnella |
| Rahnella aceris ZF458 | race | 1 | Bacteria | Pseudomonadota | Gammaproteobacteria | Enterobacterales | Yersiniaceae | Rahnella |
| Rahnella aquatilis CIP 78.65 = ATCC 33071 | raq | 1 | Bacteria | Pseudomonadota | Gammaproteobacteria | Enterobacterales | Yersiniaceae | Rahnella |
| Rahnella aquatilis HX2 | raa | 1 | Bacteria | Pseudomonadota | Gammaproteobacteria | Enterobacterales | Yersiniaceae | Rahnella |
| Rahnella bonaserana | rbon | 1 | Bacteria | Pseudomonadota | Gammaproteobacteria | Enterobacterales | Yersiniaceae | Rahnella |
| Rahnella inusitata | riu | 1 | Bacteria | Pseudomonadota | Gammaproteobacteria | Enterobacterales | Yersiniaceae | Rahnella |
| Rahnella sikkimica | rox | 1 | Bacteria | Pseudomonadota | Gammaproteobacteria | Enterobacterales | Yersiniaceae | Rahnella |
| Rahnella victoriana | rvc | 1 | Bacteria | Pseudomonadota | Gammaproteobacteria | Enterobacterales | Yersiniaceae | Rahnella |
| Rouxiella badensis subsp. acadiensis | rbad | 1 | Bacteria | Pseudomonadota | Gammaproteobacteria | Enterobacterales | Yersiniaceae | Rouxiella |
| Rouxiella chamberiensis | rcb | 1 | Bacteria | Pseudomonadota | Gammaproteobacteria | Enterobacterales | Yersiniaceae | Rouxiella |
| Serratia entomophila | senp | 2 | Bacteria | Pseudomonadota | Gammaproteobacteria | Enterobacterales | Yersiniaceae | Serratia |
| Serratia ficaria | sfj | 1 | Bacteria | Pseudomonadota | Gammaproteobacteria | Enterobacterales | Yersiniaceae | Serratia |
| Serratia fonticola DSM 4576 | sfw | 1 | Bacteria | Pseudomonadota | Gammaproteobacteria | Enterobacterales | Yersiniaceae | Serratia |
| Serratia fonticola GS2 | sfg | 1 | Bacteria | Pseudomonadota | Gammaproteobacteria | Enterobacterales | Yersiniaceae | Serratia |
| Serratia grimesii | sgri | 1 | Bacteria | Pseudomonadota | Gammaproteobacteria | Enterobacterales | Yersiniaceae | Serratia |
| Serratia inhibens | sply | 1 | Bacteria | Pseudomonadota | Gammaproteobacteria | Enterobacterales | Yersiniaceae | Serratia |
| Serratia liquefaciens | slq | 1 | Bacteria | Pseudomonadota | Gammaproteobacteria | Enterobacterales | Yersiniaceae | Serratia |
| Serratia marcescens SM39 | smar | 2 | Bacteria | Pseudomonadota | Gammaproteobacteria | Enterobacterales | Yersiniaceae | Serratia |
| Serratia marcescens subsp. marcescens Db11 | smac | 1 | Bacteria | Pseudomonadota | Gammaproteobacteria | Enterobacterales | Yersiniaceae | Serratia |
| Serratia marcescens WW4 | smw | 2 | Bacteria | Pseudomonadota | Gammaproteobacteria | Enterobacterales | Yersiniaceae | Serratia |
| Serratia nematodiphila | snem | 2 | Bacteria | Pseudomonadota | Gammaproteobacteria | Enterobacterales | Yersiniaceae | Serratia |
| Serratia nevei | snev | 2 | Bacteria | Pseudomonadota | Gammaproteobacteria | Enterobacterales | Yersiniaceae | Serratia |
| Serratia plymuthica 4Rx13 | srl | 1 | Bacteria | Pseudomonadota | Gammaproteobacteria | Enterobacterales | Yersiniaceae | Serratia |
| Serratia plymuthica AS9 | srr | 1 | Bacteria | Pseudomonadota | Gammaproteobacteria | Enterobacterales | Yersiniaceae | Serratia |
| Serratia plymuthica S13 | sry | 1 | Bacteria | Pseudomonadota | Gammaproteobacteria | Enterobacterales | Yersiniaceae | Serratia |
| Serratia proteamaculans | spe | 1 | Bacteria | Pseudomonadota | Gammaproteobacteria | Enterobacterales | Yersiniaceae | Serratia |
| Serratia quinivorans | squ | 1 | Bacteria | Pseudomonadota | Gammaproteobacteria | Enterobacterales | Yersiniaceae | Serratia |
| Serratia rhizosphaerae | srhz | 1 | Bacteria | Pseudomonadota | Gammaproteobacteria | Enterobacterales | Yersiniaceae | Serratia |
| Serratia sarumanii | ssar | 2 | Bacteria | Pseudomonadota | Gammaproteobacteria | Enterobacterales | Yersiniaceae | Serratia |
| Serratia sp. AS12 | srs | 1 | Bacteria | Pseudomonadota | Gammaproteobacteria | Enterobacterales | Yersiniaceae | Serratia |
| Serratia sp. AS13 | sra | 1 | Bacteria | Pseudomonadota | Gammaproteobacteria | Enterobacterales | Yersiniaceae | Serratia |
| Serratia sp. FS14 | serf | 2 | Bacteria | Pseudomonadota | Gammaproteobacteria | Enterobacterales | Yersiniaceae | Serratia |
| Serratia sp. MYb239 | serm | 1 | Bacteria | Pseudomonadota | Gammaproteobacteria | Enterobacterales | Yersiniaceae | Serratia |
| Serratia sp. SCBI | sers | 1 | Bacteria | Pseudomonadota | Gammaproteobacteria | Enterobacterales | Yersiniaceae | Serratia |
| Serratia surfactantfaciens | ssur | 2 | Bacteria | Pseudomonadota | Gammaproteobacteria | Enterobacterales | Yersiniaceae | Serratia |
| Serratia ureilytica | suri | 1 | Bacteria | Pseudomonadota | Gammaproteobacteria | Enterobacterales | Yersiniaceae | Serratia |
| Celerinatantimonas diazotrophica | cdiz | 1 | Bacteria | Pseudomonadota | Gammaproteobacteria | Unclassified | Celerinatantimonadaceae | Celerinatantimonas |
| Stenotrophomonas geniculata | sgen | 1 | Bacteria | Pseudomonadota | Gammaproteobacteria | Lysobacterales | Lysobacteraceae | Stenotrophomonas |
| Stenotrophomonas indicatrix | sinc | 1 | Bacteria | Pseudomonadota | Gammaproteobacteria | Lysobacterales | Lysobacteraceae | Stenotrophomonas |
| Stenotrophomonas maltophilia D457 | smz | 1 | Bacteria | Pseudomonadota | Gammaproteobacteria | Lysobacterales | Lysobacteraceae | Stenotrophomonas |
| Stenotrophomonas maltophilia JV3 | buj | 1 | Bacteria | Pseudomonadota | Gammaproteobacteria | Lysobacterales | Lysobacteraceae | Stenotrophomonas |
| Stenotrophomonas maltophilia K279a | sml | 1 | Bacteria | Pseudomonadota | Gammaproteobacteria | Lysobacterales | Lysobacteraceae | Stenotrophomonas |
| Stenotrophomonas maltophilia R551-3 | smt | 1 | Bacteria | Pseudomonadota | Gammaproteobacteria | Lysobacterales | Lysobacteraceae | Stenotrophomonas |
| Stenotrophomonas oleivorans | stes | 1 | Bacteria | Pseudomonadota | Gammaproteobacteria | Lysobacterales | Lysobacteraceae | Stenotrophomonas |
| Stenotrophomonas pavanii | spaq | 1 | Bacteria | Pseudomonadota | Gammaproteobacteria | Lysobacterales | Lysobacteraceae | Stenotrophomonas |
| Stenotrophomonas sp. MYb57 | stem | 1 | Bacteria | Pseudomonadota | Gammaproteobacteria | Lysobacterales | Lysobacteraceae | Stenotrophomonas |
| Stenotrophomonas sp. WZN-1 | sten | 1 | Bacteria | Pseudomonadota | Gammaproteobacteria | Lysobacterales | Lysobacteraceae | Stenotrophomonas |
| Xanthomonas hortorum | xhr | 1 | Bacteria | Pseudomonadota | Gammaproteobacteria | Lysobacterales | Lysobacteraceae | Xanthomonas |
| Xanthomonas hortorum pv. gardneri | xga | 1 | Bacteria | Pseudomonadota | Gammaproteobacteria | Lysobacterales | Lysobacteraceae | Xanthomonas |
| Xanthomonas hyacinthi | xhy | 1 | Bacteria | Pseudomonadota | Gammaproteobacteria | Lysobacterales | Lysobacteraceae | Xanthomonas |
| Xanthomonas hydrangeae | xhd | 1 | Bacteria | Pseudomonadota | Gammaproteobacteria | Lysobacterales | Lysobacteraceae | Xanthomonas |
| Methylocaldum szegediense | msze | 1 | Bacteria | Pseudomonadota | Gammaproteobacteria | Methylococcales | Methylococcaceae | Methylocaldum |
| Methylotuvimicrobium alcaliphilum | mah | 2 | Bacteria | Pseudomonadota | Gammaproteobacteria | Methylococcales | Methylococcaceae | Methylotuvimicrobium |
| Moraxella haemolytica | mhac | 1 | Bacteria | Pseudomonadota | Gammaproteobacteria | Moraxellales | Moraxellaceae | Moraxella |
| Moraxella nasibovis | mnj | 1 | Bacteria | Pseudomonadota | Gammaproteobacteria | Moraxellales | Moraxellaceae | Moraxella |
| Moraxella ovis | moi | 1 | Bacteria | Pseudomonadota | Gammaproteobacteria | Moraxellales | Moraxellaceae | Moraxella |
| Cobetia sp. AM6 | cobe | 1 | Bacteria | Pseudomonadota | Gammaproteobacteria | Oceanospirillales | Halomonadaceae | Cobetia |
| Halomonas alkalicola | halw | 1 | Bacteria | Pseudomonadota | Gammaproteobacteria | Oceanospirillales | Halomonadaceae | Halomonas |
| Halomonas sp. A3H3 | haah | 1 | Bacteria | Pseudomonadota | Gammaproteobacteria | Oceanospirillales | Halomonadaceae | Halomonas |
| Halotalea alkalilenta | haa | 1 | Bacteria | Pseudomonadota | Gammaproteobacteria | Oceanospirillales | Halomonadaceae | Halotalea |
| Marinobacterium iners | mgeo | 1 | Bacteria | Pseudomonadota | Gammaproteobacteria | Oceanospirillales | Oceanospirillaceae | Marinobacterium |
| Marinobacterium rhizophilum | mrz | 1 | Bacteria | Pseudomonadota | Gammaproteobacteria | Oceanospirillales | Oceanospirillaceae | Marinobacterium |
| Marinobacterium sp. LSUCC0821 | marl | 1 | Bacteria | Pseudomonadota | Gammaproteobacteria | Oceanospirillales | Oceanospirillaceae | Marinobacterium |
| Neptunomonas japonica | njp | 1 | Bacteria | Pseudomonadota | Gammaproteobacteria | Oceanospirillales | Oceanospirillaceae | Neptunomonas |
| Gilliamella apicola | gap | 1 | Bacteria | Pseudomonadota | Gammaproteobacteria | Orbales | Orbaceae | Gilliamella |
| Gilliamella apis | gaq | 1 | Bacteria | Pseudomonadota | Gammaproteobacteria | Orbales | Orbaceae | Gilliamella |
| Zophobihabitans entericus | orb | 1 | Bacteria | Pseudomonadota | Gammaproteobacteria | Orbales | Orbaceae | Zophobihabitans |
| Marinobacter sp. BSs20148 | mbs | 1 | Bacteria | Pseudomonadota | Gammaproteobacteria | Pseudomonadales | Marinobacteraceae | Marinobacter |
| Marinobacter sp. JH2 | marj | 1 | Bacteria | Pseudomonadota | Gammaproteobacteria | Pseudomonadales | Marinobacteraceae | Marinobacter |
| Aquipseudomonas alcaligenes | palc | 1 | Bacteria | Pseudomonadota | Gammaproteobacteria | Pseudomonadales | Pseudomonadaceae | Aquipseudomonas |
| Metapseudomonas furukawaii | pfuw | 1 | Bacteria | Pseudomonadota | Gammaproteobacteria | Pseudomonadales | Pseudomonadaceae | Metapseudomonas |
| Metapseudomonas otitidis | poj | 1 | Bacteria | Pseudomonadota | Gammaproteobacteria | Pseudomonadales | Pseudomonadaceae | Metapseudomonas |
| Metapseudomonas resinovorans | pre | 2 | Bacteria | Pseudomonadota | Gammaproteobacteria | Pseudomonadales | Pseudomonadaceae | Metapseudomonas |
| Pseudomonas aeruginosa B136-33 | psg | 2 | Bacteria | Pseudomonadota | Gammaproteobacteria | Pseudomonadales | Pseudomonadaceae | Pseudomonas |
| Pseudomonas aeruginosa B18 | sech | 2 | Bacteria | Pseudomonadota | Gammaproteobacteria | Pseudomonadales | Pseudomonadaceae | Pseudomonas |
| Pseudomonas aeruginosa c7447m | paec | 2 | Bacteria | Pseudomonadota | Gammaproteobacteria | Pseudomonadales | Pseudomonadaceae | Pseudomonas |
| Pseudomonas aeruginosa DK2 | pdk | 2 | Bacteria | Pseudomonadota | Gammaproteobacteria | Pseudomonadales | Pseudomonadaceae | Pseudomonas |
| Pseudomonas aeruginosa LES431 | pael | 2 | Bacteria | Pseudomonadota | Gammaproteobacteria | Pseudomonadales | Pseudomonadaceae | Pseudomonas |
| Pseudomonas aeruginosa LESB58 | pag | 2 | Bacteria | Pseudomonadota | Gammaproteobacteria | Pseudomonadales | Pseudomonadaceae | Pseudomonas |
| Pseudomonas aeruginosa M18 | paf | 2 | Bacteria | Pseudomonadota | Gammaproteobacteria | Pseudomonadales | Pseudomonadaceae | Pseudomonas |
| Pseudomonas aeruginosa MTB-1 | paem | 2 | Bacteria | Pseudomonadota | Gammaproteobacteria | Pseudomonadales | Pseudomonadaceae | Pseudomonas |
| Pseudomonas aeruginosa NCGM 1900 | paeb | 2 | Bacteria | Pseudomonadota | Gammaproteobacteria | Pseudomonadales | Pseudomonadaceae | Pseudomonas |
| Pseudomonas aeruginosa NCGM2.S1 | pnc | 2 | Bacteria | Pseudomonadota | Gammaproteobacteria | Pseudomonadales | Pseudomonadaceae | Pseudomonas |
| Pseudomonas aeruginosa PA1 | paep | 2 | Bacteria | Pseudomonadota | Gammaproteobacteria | Pseudomonadales | Pseudomonadaceae | Pseudomonas |
| Pseudomonas aeruginosa PA1R | paer | 2 | Bacteria | Pseudomonadota | Gammaproteobacteria | Pseudomonadales | Pseudomonadaceae | Pseudomonas |
| Pseudomonas aeruginosa PA38182 | paeu | 2 | Bacteria | Pseudomonadota | Gammaproteobacteria | Pseudomonadales | Pseudomonadaceae | Pseudomonas |
| Pseudomonas aeruginosa PAO1 | pae | 2 | Bacteria | Pseudomonadota | Gammaproteobacteria | Pseudomonadales | Pseudomonadaceae | Pseudomonas |
| Pseudomonas aeruginosa PAO1-VE13 | paev | 2 | Bacteria | Pseudomonadota | Gammaproteobacteria | Pseudomonadales | Pseudomonadaceae | Pseudomonas |
| Pseudomonas aeruginosa PAO1-VE2 | paei | 2 | Bacteria | Pseudomonadota | Gammaproteobacteria | Pseudomonadales | Pseudomonadaceae | Pseudomonas |
| Pseudomonas aeruginosa PAO581 | paeo | 2 | Bacteria | Pseudomonadota | Gammaproteobacteria | Pseudomonadales | Pseudomonadaceae | Pseudomonas |
| Pseudomonas aeruginosa RP73 | prp | 2 | Bacteria | Pseudomonadota | Gammaproteobacteria | Pseudomonadales | Pseudomonadaceae | Pseudomonas |
| Pseudomonas aeruginosa SCV20265 | paes | 2 | Bacteria | Pseudomonadota | Gammaproteobacteria | Pseudomonadales | Pseudomonadaceae | Pseudomonas |
| Pseudomonas aeruginosa UCBPP-PA14 | pau | 2 | Bacteria | Pseudomonadota | Gammaproteobacteria | Pseudomonadales | Pseudomonadaceae | Pseudomonas |
| Pseudomonas aeruginosa YL84 | paeg | 2 | Bacteria | Pseudomonadota | Gammaproteobacteria | Pseudomonadales | Pseudomonadaceae | Pseudomonas |
| Pseudomonas asiatica | pasi | 1 | Bacteria | Pseudomonadota | Gammaproteobacteria | Pseudomonadales | Pseudomonadaceae | Pseudomonas |
| Pseudomonas batumici | pbam | 1 | Bacteria | Pseudomonadota | Gammaproteobacteria | Pseudomonadales | Pseudomonadaceae | Pseudomonas |
| Pseudomonas brassicacearum subsp. brassicacearum NFM421 | pba | 1 | Bacteria | Pseudomonadota | Gammaproteobacteria | Pseudomonadales | Pseudomonadaceae | Pseudomonas |
| Pseudomonas chlororaphis PA23 | pch | 2 | Bacteria | Pseudomonadota | Gammaproteobacteria | Pseudomonadales | Pseudomonadaceae | Pseudomonas |
| Pseudomonas chlororaphis PCL1606 | pcz | 1 | Bacteria | Pseudomonadota | Gammaproteobacteria | Pseudomonadales | Pseudomonadaceae | Pseudomonas |
| Pseudomonas chlororaphis subsp. aurantiaca | pcp | 1 | Bacteria | Pseudomonadota | Gammaproteobacteria | Pseudomonadales | Pseudomonadaceae | Pseudomonas |
| Pseudomonas chlororaphis subsp. piscium | pchp | 1 | Bacteria | Pseudomonadota | Gammaproteobacteria | Pseudomonadales | Pseudomonadaceae | Pseudomonas |
| Pseudomonas denitrificans (nom. rej.) | pden | 1 | Bacteria | Pseudomonadota | Gammaproteobacteria | Pseudomonadales | Pseudomonadaceae | Pseudomonas |
| Pseudomonas fluorescens A506 | pfc | 1 | Bacteria | Pseudomonadota | Gammaproteobacteria | Pseudomonadales | Pseudomonadaceae | Pseudomonas |
| Pseudomonas fluorescens Pf0-1 | pfo | 1 | Bacteria | Pseudomonadota | Gammaproteobacteria | Pseudomonadales | Pseudomonadaceae | Pseudomonas |
| Pseudomonas fluorescens UK4 | pfn | 1 | Bacteria | Pseudomonadota | Gammaproteobacteria | Pseudomonadales | Pseudomonadaceae | Pseudomonas |
| Pseudomonas fortuita | pfou | 1 | Bacteria | Pseudomonadota | Gammaproteobacteria | Pseudomonadales | Pseudomonadaceae | Pseudomonas |
| Pseudomonas glycinae | pgy | 1 | Bacteria | Pseudomonadota | Gammaproteobacteria | Pseudomonadales | Pseudomonadaceae | Pseudomonas |
| Pseudomonas hygromyciniae | phyg | 1 | Bacteria | Pseudomonadota | Gammaproteobacteria | Pseudomonadales | Pseudomonadaceae | Pseudomonas |
| Pseudomonas inefficax | pix | 1 | Bacteria | Pseudomonadota | Gammaproteobacteria | Pseudomonadales | Pseudomonadaceae | Pseudomonas |
| Pseudomonas izuensis | piz | 3 | Bacteria | Pseudomonadota | Gammaproteobacteria | Pseudomonadales | Pseudomonadaceae | Pseudomonas |
| Pseudomonas juntendi | pju | 1 | Bacteria | Pseudomonadota | Gammaproteobacteria | Pseudomonadales | Pseudomonadaceae | Pseudomonas |
| Pseudomonas kermanshahensis | pkm | 1 | Bacteria | Pseudomonadota | Gammaproteobacteria | Pseudomonadales | Pseudomonadaceae | Pseudomonas |
| Pseudomonas khavaziana | pkv | 1 | Bacteria | Pseudomonadota | Gammaproteobacteria | Pseudomonadales | Pseudomonadaceae | Pseudomonas |
| Pseudomonas kielensis | pkj | 1 | Bacteria | Pseudomonadota | Gammaproteobacteria | Pseudomonadales | Pseudomonadaceae | Pseudomonas |
| Pseudomonas knackmussii | pkc | 1 | Bacteria | Pseudomonadota | Gammaproteobacteria | Pseudomonadales | Pseudomonadaceae | Pseudomonas |
| Pseudomonas kribbensis | pke | 1 | Bacteria | Pseudomonadota | Gammaproteobacteria | Pseudomonadales | Pseudomonadaceae | Pseudomonas |
| Pseudomonas kurunegalensis | pkk | 1 | Bacteria | Pseudomonadota | Gammaproteobacteria | Pseudomonadales | Pseudomonadaceae | Pseudomonas |
| Pseudomonas lalucatii | plau | 1 | Bacteria | Pseudomonadota | Gammaproteobacteria | Pseudomonadales | Pseudomonadaceae | Pseudomonas |
| Pseudomonas lurida | pfx | 1 | Bacteria | Pseudomonadota | Gammaproteobacteria | Pseudomonadales | Pseudomonadaceae | Pseudomonas |
| Pseudomonas luteola | plul | 1 | Bacteria | Pseudomonadota | Gammaproteobacteria | Pseudomonadales | Pseudomonadaceae | Pseudomonas |
| Pseudomonas monteilii SB3078 | pmon | 1 | Bacteria | Pseudomonadota | Gammaproteobacteria | Pseudomonadales | Pseudomonadaceae | Pseudomonas |
| Pseudomonas monteilii SB3101 | pmot | 1 | Bacteria | Pseudomonadota | Gammaproteobacteria | Pseudomonadales | Pseudomonadaceae | Pseudomonas |
| Pseudomonas multiresinivorans | pmui | 1 | Bacteria | Pseudomonadota | Gammaproteobacteria | Pseudomonadales | Pseudomonadaceae | Pseudomonas |
| Pseudomonas nitroreducens | pnt | 1 | Bacteria | Pseudomonadota | Gammaproteobacteria | Pseudomonadales | Pseudomonadaceae | Pseudomonas |
| Pseudomonas oryzihabitans PRS08-11306 | ppsl | 1 | Bacteria | Pseudomonadota | Gammaproteobacteria | Pseudomonadales | Pseudomonadaceae | Pseudomonas |
| Pseudomonas oryzihabitans USDA-ARS-USMARC-56511 | por | 1 | Bacteria | Pseudomonadota | Gammaproteobacteria | Pseudomonadales | Pseudomonadaceae | Pseudomonas |
| Pseudomonas paraeruginosa | ppaa | 1 | Bacteria | Pseudomonadota | Gammaproteobacteria | Pseudomonadales | Pseudomonadaceae | Pseudomonas |
| Pseudomonas paraeruginosa PA7 | pap | 1 | Bacteria | Pseudomonadota | Gammaproteobacteria | Pseudomonadales | Pseudomonadaceae | Pseudomonas |
| Pseudomonas piscis | ppii | 3 | Bacteria | Pseudomonadota | Gammaproteobacteria | Pseudomonadales | Pseudomonadaceae | Pseudomonas |
| Pseudomonas plecoglossicida | ppj | 1 | Bacteria | Pseudomonadota | Gammaproteobacteria | Pseudomonadales | Pseudomonadaceae | Pseudomonas |
| Pseudomonas promysalinigenes | pprg | 1 | Bacteria | Pseudomonadota | Gammaproteobacteria | Pseudomonadales | Pseudomonadaceae | Pseudomonas |
| Pseudomonas protegens Cab57 | ppro | 1 | Bacteria | Pseudomonadota | Gammaproteobacteria | Pseudomonadales | Pseudomonadaceae | Pseudomonas |
| Pseudomonas protegens CHA0 | pprc | 2 | Bacteria | Pseudomonadota | Gammaproteobacteria | Pseudomonadales | Pseudomonadaceae | Pseudomonas |
| Pseudomonas protegens Pf-5 | pfl | 2 | Bacteria | Pseudomonadota | Gammaproteobacteria | Pseudomonadales | Pseudomonadaceae | Pseudomonas |
| Pseudomonas putida BIRD-1 | ppb | 1 | Bacteria | Pseudomonadota | Gammaproteobacteria | Pseudomonadales | Pseudomonadaceae | Pseudomonas |
| Pseudomonas putida DLL-E4 | ppud | 1 | Bacteria | Pseudomonadota | Gammaproteobacteria | Pseudomonadales | Pseudomonadaceae | Pseudomonas |
| Pseudomonas putida DOT-T1E | ppx | 1 | Bacteria | Pseudomonadota | Gammaproteobacteria | Pseudomonadales | Pseudomonadaceae | Pseudomonas |
| Pseudomonas putida F1 | ppf | 1 | Bacteria | Pseudomonadota | Gammaproteobacteria | Pseudomonadales | Pseudomonadaceae | Pseudomonas |
| Pseudomonas putida GB-1 | ppg | 1 | Bacteria | Pseudomonadota | Gammaproteobacteria | Pseudomonadales | Pseudomonadaceae | Pseudomonas |
| Pseudomonas putida H8234 | pput | 1 | Bacteria | Pseudomonadota | Gammaproteobacteria | Pseudomonadales | Pseudomonadaceae | Pseudomonas |
| Pseudomonas putida HB3267 | ppuh | 1 | Bacteria | Pseudomonadota | Gammaproteobacteria | Pseudomonadales | Pseudomonadaceae | Pseudomonas |
| Pseudomonas putida NBRC 14164 | ppun | 1 | Bacteria | Pseudomonadota | Gammaproteobacteria | Pseudomonadales | Pseudomonadaceae | Pseudomonas |
| Pseudomonas putida ND6 | ppi | 1 | Bacteria | Pseudomonadota | Gammaproteobacteria | Pseudomonadales | Pseudomonadaceae | Pseudomonas |
| Pseudomonas putida S16 | ppt | 1 | Bacteria | Pseudomonadota | Gammaproteobacteria | Pseudomonadales | Pseudomonadaceae | Pseudomonas |
| Pseudomonas putida W619 | ppw | 1 | Bacteria | Pseudomonadota | Gammaproteobacteria | Pseudomonadales | Pseudomonadaceae | Pseudomonas |
| Pseudomonas salmasensis | psam | 1 | Bacteria | Pseudomonadota | Gammaproteobacteria | Pseudomonadales | Pseudomonadaceae | Pseudomonas |
| Pseudomonas sessilinigenes | psep | 3 | Bacteria | Pseudomonadota | Gammaproteobacteria | Pseudomonadales | Pseudomonadaceae | Pseudomonas |
| Pseudomonas shahriarae | pshh | 1 | Bacteria | Pseudomonadota | Gammaproteobacteria | Pseudomonadales | Pseudomonadaceae | Pseudomonas |
| Pseudomonas silesiensis | psil | 1 | Bacteria | Pseudomonadota | Gammaproteobacteria | Pseudomonadales | Pseudomonadaceae | Pseudomonas |
| Pseudomonas solani Sm006 | psoa | 1 | Bacteria | Pseudomonadota | Gammaproteobacteria | Pseudomonadales | Pseudomonadaceae | Pseudomonas |
| Pseudomonas sp. HS6 | pshs | 2 | Bacteria | Pseudomonadota | Gammaproteobacteria | Pseudomonadales | Pseudomonadaceae | Pseudomonas |
| Pseudomonas sp. JY-Q | psjy | 1 | Bacteria | Pseudomonadota | Gammaproteobacteria | Pseudomonadales | Pseudomonadaceae | Pseudomonas |
| Pseudomonas sp. MRSN12121 | psem | 2 | Bacteria | Pseudomonadota | Gammaproteobacteria | Pseudomonadales | Pseudomonadaceae | Pseudomonas |
| Pseudomonas sp. NC02 | psnc | 2 | Bacteria | Pseudomonadota | Gammaproteobacteria | Pseudomonadales | Pseudomonadaceae | Pseudomonas |
| Pseudomonas sp. Os17 | psos | 1 | Bacteria | Pseudomonadota | Gammaproteobacteria | Pseudomonadales | Pseudomonadaceae | Pseudomonas |
| Pseudomonas sp. TCU-HL1 | pset | 1 | Bacteria | Pseudomonadota | Gammaproteobacteria | Pseudomonadales | Pseudomonadaceae | Pseudomonas |
| Pseudomonas sp. UW4 | ppuu | 1 | Bacteria | Pseudomonadota | Gammaproteobacteria | Pseudomonadales | Pseudomonadaceae | Pseudomonas |
| Pseudomonas sp. VLB120 | psv | 1 | Bacteria | Pseudomonadota | Gammaproteobacteria | Pseudomonadales | Pseudomonadaceae | Pseudomonas |
| Pseudomonas synxantha LBUM223 | pfb | 1 | Bacteria | Pseudomonadota | Gammaproteobacteria | Pseudomonadales | Pseudomonadaceae | Pseudomonas |
| Pseudomonas taiwanensis | ptai | 1 | Bacteria | Pseudomonadota | Gammaproteobacteria | Pseudomonadales | Pseudomonadaceae | Pseudomonas |
| Pseudomonas tohonis | ptw | 1 | Bacteria | Pseudomonadota | Gammaproteobacteria | Pseudomonadales | Pseudomonadaceae | Pseudomonas |
| Pseudomonas trivialis | ptv | 1 | Bacteria | Pseudomonadota | Gammaproteobacteria | Pseudomonadales | Pseudomonadaceae | Pseudomonas |
| Pseudomonas umsongensis | pum | 1 | Bacteria | Pseudomonadota | Gammaproteobacteria | Pseudomonadales | Pseudomonadaceae | Pseudomonas |
| Pseudomonas veronii | pvr | 2 | Bacteria | Pseudomonadota | Gammaproteobacteria | Pseudomonadales | Pseudomonadaceae | Pseudomonas |
| Pseudomonas versuta | ppsy | 1 | Bacteria | Pseudomonadota | Gammaproteobacteria | Pseudomonadales | Pseudomonadaceae | Pseudomonas |
| Pseudomonas viridiflava | pvd | 1 | Bacteria | Pseudomonadota | Gammaproteobacteria | Pseudomonadales | Pseudomonadaceae | Pseudomonas |
| Pseudomonas wayambapalatensis | pwy | 1 | Bacteria | Pseudomonadota | Gammaproteobacteria | Pseudomonadales | Pseudomonadaceae | Pseudomonas |
| Pseudomonas yamanorum | pym | 2 | Bacteria | Pseudomonadota | Gammaproteobacteria | Pseudomonadales | Pseudomonadaceae | Pseudomonas |
| Stutzerimonas frequens | psed | 1 | Bacteria | Pseudomonadota | Gammaproteobacteria | Pseudomonadales | Pseudomonadaceae | Stutzerimonas |
| Stutzerimonas stutzeri RCH2 | psh | 1 | Bacteria | Pseudomonadota | Gammaproteobacteria | Pseudomonadales | Pseudomonadaceae | Stutzerimonas |
| Francisella orientalis Toba 04 | fna | 1 | Bacteria | Pseudomonadota | Gammaproteobacteria | Thiotrichales | Francisellaceae | Francisella |
| Francisella philomiragia subsp. philomiragia ATCC 25017 | fph | 1 | Bacteria | Pseudomonadota | Gammaproteobacteria | Thiotrichales | Francisellaceae | Francisella |
| Vibrio furnissii | vfu | 1 | Bacteria | Pseudomonadota | Gammaproteobacteria | Vibrionales | Vibrionaceae | Vibrio |
| Vibrio natriegens | vna | 1 | Bacteria | Pseudomonadota | Gammaproteobacteria | Vibrionales | Vibrionaceae | Vibrio |
| Vibrio sp. EJY3 | vej | 1 | Bacteria | Pseudomonadota | Gammaproteobacteria | Vibrionales | Vibrionaceae | Vibrio |
| Desulfofustis limnaeus | dlm | 1 | Bacteria | Thermodesulfobacteriota | Desulfobulbia | Desulfobulbales | Desulfocapsaceae | Desulfofustis |
| Citrifermentans bremense | gbn | 1 | Bacteria | Thermodesulfobacteriota | Desulfuromonadia | Geobacterales | Geobacteraceae | Citrifermentans |

**Supplementary Table 7.** Results for Michaelis–Menten analysis (GraphPad Prism)

| **Michaelis-Menten** | **Ko6_01540** | **PSKpn11_20955** |
| --- | --- | --- |
| **Best-fit values** |  |  |
| Vmax | ~ 15316595606311 | 0.2251 |
| Km | ~ 1.601e+016 | 30.99 |
| **95% CI (profile likelihood)** | |  |
| Vmax | (Very wide) | 0.08786 to +infinity |
| Km | (Very wide) | 4.569 to +infinity |
| **Goodness of Fit** |  |  |
| Degrees of Freedom | 3 | 3 |
| R squared | 0.9483 | 0.962 |
| Sum of Squares | 0.00001492 | 0.0001923 |
| Sy.x | 0.00223 | 0.008006 |
| **Constraints** |  |  |
| Km | Km > 0 | Km > 0 |

**Supplementary Table 8.** Detection of group 1 and group 2 protein sequences in *Klebsiella* genome data

| **Name** | **Group1_aa** | **Group2_aa** | **Species** |
| --- | --- | --- | --- |
| ERR025623 | 392 | 389 | Klebsiella aerogenes |
| GCF_000215745 | 392 | 389 | Klebsiella aerogenes |
| GCF_000334515 | 392 | 389 | Klebsiella aerogenes |
| GCF_000383335 | 392 | 389 | Klebsiella aerogenes |
| GCF_000534075 | 392 | 389 | Klebsiella aerogenes |
| GCF_000534095 | 392 | 389 | Klebsiella aerogenes |
| GCF_000534115 | 392 | 389 | Klebsiella aerogenes |
| GCF_000534135 | 392 | 389 | Klebsiella aerogenes |
| GCF_000534235 | 392 | 389 | Klebsiella aerogenes |
| GCF_000534255 | 392 | 389 | Klebsiella aerogenes |
| GCF_000534315 | 392 | 389 | Klebsiella aerogenes |
| GCF_000534335 | 392 | 389 | Klebsiella aerogenes |
| GCF_000692155 | 392 | 389 | Klebsiella aerogenes |
| GCF_000692175 | 392 | 389 | Klebsiella aerogenes |
| GCF_000692195 | 392 | 389 | Klebsiella aerogenes |
| GCF_000692215 | 392 | 389 | Klebsiella aerogenes |
| GCF_000755545 | 392 | 389 | Klebsiella aerogenes |
| GCF_000802765 | 392 | 389 | Klebsiella aerogenes |
| GCF_000956775 | 392 | 389 | Klebsiella aerogenes |
| GCF_000956895 | 392 | 389 | Klebsiella aerogenes |
| GCF_000957485 | 392 | 389 | Klebsiella aerogenes |
| GCF_000957525 | 392 | 389 | Klebsiella aerogenes |
| GCF_000957705 | 392 | 389 | Klebsiella aerogenes |
| GCF_000957715 | 392 | 389 | Klebsiella aerogenes |
| GCF_000957725 | 392 | 389 | Klebsiella aerogenes |
| GCF_000958075 | 392 | 389 | Klebsiella aerogenes |
| GCF_000958305 | 392 | 389 | Klebsiella aerogenes |
| GCF_001006555 | 392 | 389 | Klebsiella aerogenes |
| GCF_001011175 | 392 | 244 | Klebsiella aerogenes |
| GCF_001011185 | 392 | 389 | Klebsiella aerogenes |
| GCF_001011195 | 392 | 389 | Klebsiella aerogenes |
| GCF_001011205 | 392 | 389 | Klebsiella aerogenes |
| GCF_001011255 | 392 | 389 | Klebsiella aerogenes |
| GCF_001011265 | 392 | 389 | Klebsiella aerogenes |
| GCF_001011295 | 392 | 389 | Klebsiella aerogenes |
| GCF_001011305 | 392 | 389 | Klebsiella aerogenes |
| GCF_001011335 | 392 | 244 | Klebsiella aerogenes |
| GCF_001011355 | 392 | 389 | Klebsiella aerogenes |
| GCF_001011375 | 392 | 389 | Klebsiella aerogenes |
| GCF_001011385 | 392 | 389 | Klebsiella aerogenes |
| GCF_001011415 | 392 | 389 | Klebsiella aerogenes |
| GCF_001011425 | 392 | 389 | Klebsiella aerogenes |
| GCF_001011455 | 392 | 389 | Klebsiella aerogenes |
| GCF_001011475 | 67 | 389 | Klebsiella aerogenes |
| GCF_001011475 | 392 | 389 | Klebsiella aerogenes |
| GCF_001011495 | 392 | 389 | Klebsiella aerogenes |
| GCF_001011515 | 392 | 244 | Klebsiella aerogenes |
| GCF_001011535 | 392 | 389 | Klebsiella aerogenes |
| GCF_001011545 | 392 | 389 | Klebsiella aerogenes |
| GCF_001011575 | 392 | 389 | Klebsiella aerogenes |
| GCF_001011595 | 392 | 389 | Klebsiella aerogenes |
| GCF_001011615 | 392 | 244 | Klebsiella aerogenes |
| GCF_001011625 | 392 | 244 | Klebsiella aerogenes |
| GCF_001011645 | 94 | 389 | Klebsiella aerogenes |
| GCF_001011645 | 392 | 389 | Klebsiella aerogenes |
| GCF_001011795 | 392 | 244 | Klebsiella aerogenes |
| GCF_001011815 | 392 | 389 | Klebsiella aerogenes |
| GCF_001011915 | 392 | 244 | Klebsiella aerogenes |
| GCF_001011935 | 392 | 389 | Klebsiella aerogenes |
| GCF_001021995 | 392 | 389 | Klebsiella aerogenes |
| GCF_001030055 | 392 | 389 | Klebsiella aerogenes |
| GCF_001030125 | 392 | 389 | Klebsiella aerogenes |
| GCF_001030165 | 392 | 389 | Klebsiella aerogenes |
| GCF_001030185 | 392 | 389 | Klebsiella aerogenes |
| GCF_001052095 | 392 | 389 | Klebsiella aerogenes |
| GCF_001052565 | 392 | 389 | Klebsiella aerogenes |
| GCF_001053235 | 392 | 389 | Klebsiella aerogenes |
| GCF_001053595 | 392 | 389 | Klebsiella aerogenes |
| GCF_001054275 | 392 | 389 | Klebsiella aerogenes |
| GCF_001054315 | 392 | 389 | Klebsiella aerogenes |
| GCF_001054405 | 392 | 389 | Klebsiella aerogenes |
| GCF_001055555 | 392 | 389 | Klebsiella aerogenes |
| GCF_001057945 | 392 | 389 | Klebsiella aerogenes |
| GCF_001058645 | 392 | 389 | Klebsiella aerogenes |
| GCF_001059975 | 392 | 389 | Klebsiella aerogenes |
| GCF_001472155 | 392 | 389 | Klebsiella aerogenes |
| GCF_001472275 | 392 | 389 | Klebsiella aerogenes |
| GCF_001472415 | 392 | 389 | Klebsiella aerogenes |
| GCF_001472725 | 392 | 389 | Klebsiella aerogenes |
| GCF_001472755 | 392 | 389 | Klebsiella aerogenes |
| GCF_001472935 | 392 | 389 | Klebsiella aerogenes |
| GCF_001473095 | 392 | 389 | Klebsiella aerogenes |
| GCF_001518035 | 392 | 389 | Klebsiella aerogenes |
| GCF_001518115 | 392 | 389 | Klebsiella aerogenes |
| GCF_001518125 | 392 | 389 | Klebsiella aerogenes |
| GCF_001518435 | 392 | 389 | Klebsiella aerogenes |
| GCF_001518675 | 392 | 389 | Klebsiella aerogenes |
| GCF_001525245 | 392 | 389 | Klebsiella aerogenes |
| GCF_001525265 | 392 | 389 | Klebsiella aerogenes |
| GCF_001525505 | 392 | 389 | Klebsiella aerogenes |
| GCF_001559215 | 392 | 389 | Klebsiella aerogenes |
| GCF_001571545 | 392 | 389 | Klebsiella aerogenes |
| GCF_001593585 | 392 | 389 | Klebsiella aerogenes |
| GCF_001631185 | 392 | 389 | Klebsiella aerogenes |
| GCF_001631215 | 392 | 389 | Klebsiella aerogenes |
| GCF_001631305 | 392 | 389 | Klebsiella aerogenes |
| GCF_001631315 | 392 | 389 | Klebsiella aerogenes |
| GCF_001631345 | 392 | 389 | Klebsiella aerogenes |
| GCF_001631565 | 392 | 389 | Klebsiella aerogenes |
| GCF_001631605 | 392 | 389 | Klebsiella aerogenes |
| GCF_001631645 | 392 | 389 | Klebsiella aerogenes |
| GCF_001631655 | 392 | 389 | Klebsiella aerogenes |
| GCF_001631725 | 392 | 389 | Klebsiella aerogenes |
| GCF_001631735 | 392 | 389 | Klebsiella aerogenes |
| GCF_001631755 | 392 | 389 | Klebsiella aerogenes |
| GCF_001631845 | 392 | 389 | Klebsiella aerogenes |
| GCF_001631895 | 392 | 389 | Klebsiella aerogenes |
| GCF_001649605 | 392 | 389 | Klebsiella aerogenes |
| GCF_001662695 | 392 | 389 | Klebsiella aerogenes |
| GCF_001662705 | 392 | 389 | Klebsiella aerogenes |
| GCF_001662715 | 392 | 389 | Klebsiella aerogenes |
| GCF_001662765 | 392 | 389 | Klebsiella aerogenes |
| GCF_001939895 | 392 | 389 | Klebsiella aerogenes |
| GCF_001974865 | 392 | 389 | Klebsiella aerogenes |
| GCF_002003705 | 392 | 389 | Klebsiella aerogenes |
| GCF_002152895 | 392 | 389 | Klebsiella aerogenes |
| GCF_002152915 | 392 | 389 | Klebsiella aerogenes |
| GCF_002152925 | 392 | 389 | Klebsiella aerogenes |
| GCF_002182075 | 392 | 389 | Klebsiella aerogenes |
| GCF_002184575 | 392 | 389 | Klebsiella aerogenes |
| GCF_002184625 | 392 | 389 | Klebsiella aerogenes |
| GCF_002185345 | 392 | 389 | Klebsiella aerogenes |
| GCF_002591115 | 392 | 389 | Klebsiella aerogenes |
| GCF_002796405 | 392 | 389 | Klebsiella aerogenes |
| GCF_002796425 | 392 | 389 | Klebsiella aerogenes |
| GCF_002796525 | 392 | 389 | Klebsiella aerogenes |
| GCF_002849555 | 392 | 389 | Klebsiella aerogenes |
| GCF_002852865 | 392 | 389 | Klebsiella aerogenes |
| GCF_002863955 | 392 | 389 | Klebsiella aerogenes |
| GCF_002863975 | 392 | 389 | Klebsiella aerogenes |
| GCF_002891215 | 392 | 389 | Klebsiella aerogenes |
| GCF_002918815 | 392 | 389 | Klebsiella aerogenes |
| GCF_002948835 | 392 | 389 | Klebsiella aerogenes |
| GCF_003057155 | 392 | 389 | Klebsiella aerogenes |
| GCF_003057175 | 392 | 389 | Klebsiella aerogenes |
| GCF_003057195 | 392 | 389 | Klebsiella aerogenes |
| GCF_003057215 | 392 | 389 | Klebsiella aerogenes |
| GCF_003057225 | 392 | 389 | Klebsiella aerogenes |
| GCF_003057255 | 392 | 389 | Klebsiella aerogenes |
| GCF_003071285 | 392 | 389 | Klebsiella aerogenes |
| GCF_003075515 | 392 | 389 | Klebsiella aerogenes |
| GCF_003225895 | 392 | 389 | Klebsiella aerogenes |
| GCF_003227775 | 392 | 389 | Klebsiella aerogenes |
| GCF_003227795 | 392 | 389 | Klebsiella aerogenes |
| GCF_003227835 | 392 | 64 | Klebsiella aerogenes |
| GCF_003227835 | 392 | 389 | Klebsiella aerogenes |
| GCF_003400515 | 392 | 389 | Klebsiella aerogenes |
| GCF_003400785 | 392 | 389 | Klebsiella aerogenes |
| GCF_003400825 | 392 | 389 | Klebsiella aerogenes |
| GCF_003401085 | 392 | 389 | Klebsiella aerogenes |
| GCF_003401285 | 392 | 389 | Klebsiella aerogenes |
| GCF_003401315 | 392 | 389 | Klebsiella aerogenes |
| GCF_003401425 | 392 | 389 | Klebsiella aerogenes |
| GCF_003401475 | 392 | 389 | Klebsiella aerogenes |
| GCF_003417445 | 392 | 389 | Klebsiella aerogenes |
| GCF_003425735 | 392 | 389 | Klebsiella aerogenes |
| GCF_003546885 | 392 | 389 | Klebsiella aerogenes |
| GCF_003812185 | 392 | 389 | Klebsiella aerogenes |
| GCF_003950535 | 392 | 389 | Klebsiella aerogenes |
| GCF_003950545 | 392 | 389 | Klebsiella aerogenes |
| GCF_003950575 | 392 | 389 | Klebsiella aerogenes |
| GCF_003950595 | 392 | 389 | Klebsiella aerogenes |
| GCF_003950615 | 392 | 389 | Klebsiella aerogenes |
| GCF_003950635 | 392 | 389 | Klebsiella aerogenes |
| GCF_003951115 | 392 | 389 | Klebsiella aerogenes |
| GCF_003951125 | 392 | 389 | Klebsiella aerogenes |
| GCF_003951175 | 392 | 389 | Klebsiella aerogenes |
| GCF_003951185 | 392 | 389 | Klebsiella aerogenes |
| GCF_003951195 | 392 | 389 | Klebsiella aerogenes |
| GCF_003951235 | 392 | 389 | Klebsiella aerogenes |
| GCF_003951245 | 392 | 389 | Klebsiella aerogenes |
| GCF_003951535 | 392 | 389 | Klebsiella aerogenes |
| GCF_003951575 | 392 | 389 | Klebsiella aerogenes |
| GCF_003951605 | 392 | 389 | Klebsiella aerogenes |
| GCF_003951615 | 392 | 389 | Klebsiella aerogenes |
| GCF_003951635 | 392 | 389 | Klebsiella aerogenes |
| GCF_003952035 | 392 | 389 | Klebsiella aerogenes |
| GCF_003952045 | 392 | 64 | Klebsiella aerogenes |
| GCF_003952045 | 392 | 389 | Klebsiella aerogenes |
| GCF_003952055 | 392 | 389 | Klebsiella aerogenes |
| GCF_003952105 | 392 | 389 | Klebsiella aerogenes |
| GCF_003952125 | 392 | 389 | Klebsiella aerogenes |
| GCF_003952145 | 392 | 389 | Klebsiella aerogenes |
| GCF_003952905 | 392 | 389 | Klebsiella aerogenes |
| GCF_003952915 | 392 | 389 | Klebsiella aerogenes |
| GCF_004127295 | 392 | 389 | Klebsiella aerogenes |
| GCF_004127305 | 392 | 389 | Klebsiella aerogenes |
| GCF_006517605 | 392 | 389 | Klebsiella aerogenes |
| GCF_006874725 | 392 | 389 | Klebsiella aerogenes |
| GCF_007558345 | 392 | 389 | Klebsiella aerogenes |
| GCF_007558425 | 392 | 389 | Klebsiella aerogenes |
| GCF_007558485 | 392 | 389 | Klebsiella aerogenes |
| GCF_007558625 | 392 | 389 | Klebsiella aerogenes |
| GCF_007558645 | 392 | 389 | Klebsiella aerogenes |
| GCF_007558665 | 392 | 389 | Klebsiella aerogenes |
| GCF_007558675 | 392 | 389 | Klebsiella aerogenes |
| GCF_007571405 | 392 | 389 | Klebsiella aerogenes |
| GCF_007632255 | 392 | 389 | Klebsiella aerogenes |
| GCF_008082115 | 392 | 389 | Klebsiella aerogenes |
| GCF_008364345 | 392 | 389 | Klebsiella aerogenes |
| GCF_008693885 | 392 | 389 | Klebsiella aerogenes |
| GCF_008727695 | 392 | 389 | Klebsiella aerogenes |
| GCF_008931665 | 229 | 389 | Klebsiella aerogenes |
| GCF_008931665 | 392 | 389 | Klebsiella aerogenes |
| GCF_009732795 | 392 | 389 | Klebsiella aerogenes |
| GCF_009732815 | 392 | 389 | Klebsiella aerogenes |
| GCF_009732835 | 392 | 389 | Klebsiella aerogenes |
| GCF_009757245 | 392 | 64 | Klebsiella aerogenes |
| GCF_009757245 | 392 | 389 | Klebsiella aerogenes |
| GCF_009909445 | 392 | 389 | Klebsiella aerogenes |
| GCF_010509815 | 392 | 390 | Klebsiella aerogenes |
| GCF_010588245 | 392 | 389 | Klebsiella aerogenes |
| GCF_010589945 | 392 | 389 | Klebsiella aerogenes |
| GCF_010590305 | 392 | 389 | Klebsiella aerogenes |
| GCF_010590365 | 392 | 389 | Klebsiella aerogenes |
| GCF_010590725 | 392 | 389 | Klebsiella aerogenes |
| GCF_010591065 | 392 | 389 | Klebsiella aerogenes |
| GCF_010592485 | 392 | 389 | Klebsiella aerogenes |
| GCF_010592605 | 392 | 389 | Klebsiella aerogenes |
| GCF_010592765 | 392 | 389 | Klebsiella aerogenes |
| GCF_010592775 | 392 | 389 | Klebsiella aerogenes |
| GCF_010597765 | 392 | 389 | Klebsiella aerogenes |
| GCF_010597785 | 392 | 389 | Klebsiella aerogenes |
| GCF_010597805 | 392 | 389 | Klebsiella aerogenes |
| GCF_010597985 | 392 | 389 | Klebsiella aerogenes |
| GCF_010598245 | 392 | 389 | Klebsiella aerogenes |
| GCF_010598485 | 392 | 389 | Klebsiella aerogenes |
| GCF_010598495 | 392 | 389 | Klebsiella aerogenes |
| GCF_010598905 | 392 | 389 | Klebsiella aerogenes |
| GCF_011006785 | 392 | 389 | Klebsiella aerogenes |
| GCF_011006935 | 392 | 389 | Klebsiella aerogenes |
| GCF_011067245 | 392 | 389 | Klebsiella aerogenes |
| GCF_011392705 | 392 | 389 | Klebsiella aerogenes |
| GCF_011392925 | 392 | 389 | Klebsiella aerogenes |
| GCF_011604725 | 392 | 389 | Klebsiella aerogenes |
| GCF_013166315 | 392 | 389 | Klebsiella aerogenes |
| GCF_013166325 | 392 | 389 | Klebsiella aerogenes |
| GCF_013166335 | 392 | 389 | Klebsiella aerogenes |
| GCF_013166345 | 392 | 389 | Klebsiella aerogenes |
| GCF_013166415 | 392 | 389 | Klebsiella aerogenes |
| GCF_013166435 | 392 | 389 | Klebsiella aerogenes |
| GCF_013166445 | 392 | 389 | Klebsiella aerogenes |
| GCF_013403435 | 392 | 389 | Klebsiella aerogenes |
| GCF_013727635 | 392 | 389 | Klebsiella aerogenes |
| GCF_013925105 | 392 | 389 | Klebsiella aerogenes |
| GCF_014169215 | 392 | 389 | Klebsiella aerogenes |
| GCF_014235545 | 392 | 389 | Klebsiella aerogenes |
| GCF_014333355 | 392 | 389 | Klebsiella aerogenes |
| GCF_014856475 | 392 | 389 | Klebsiella aerogenes |
| GCF_014856485 | 392 | 389 | Klebsiella aerogenes |
| GCF_014856495 | 392 | 389 | Klebsiella aerogenes |
| GCF_014901435 | 392 | 389 | Klebsiella aerogenes |
| GCF_014901455 | 392 | 389 | Klebsiella aerogenes |
| GCF_014901595 | 392 | 389 | Klebsiella aerogenes |
| GCF_014901605 | 392 | 389 | Klebsiella aerogenes |
| GCF_014901995 | 392 | 389 | Klebsiella aerogenes |
| GCF_014902045 | 392 | 389 | Klebsiella aerogenes |
| GCF_014902465 | 392 | 389 | Klebsiella aerogenes |
| GCF_014902735 | 392 | 389 | Klebsiella aerogenes |
| GCF_014902765 | 392 | 389 | Klebsiella aerogenes |
| GCF_014902795 | 392 | 389 | Klebsiella aerogenes |
| GCF_014902855 | 392 | 389 | Klebsiella aerogenes |
| GCF_015679435 | 392 | 389 | Klebsiella aerogenes |
| GCF_015701185 | 392 | 389 | Klebsiella aerogenes |
| GCF_015701195 | 392 | 389 | Klebsiella aerogenes |
| GCF_015701235 | 392 | 389 | Klebsiella aerogenes |
| GCF_015718665 | 392 | 389 | Klebsiella aerogenes |
| GCF_016527315 | 392 | 389 | Klebsiella aerogenes |
| GCF_016527875 | 392 | 389 | Klebsiella aerogenes |
| GCF_016529455 | 392 | 389 | Klebsiella aerogenes |
| GCF_016554165 | 392 | 389 | Klebsiella aerogenes |
| GCF_016643805 | 392 | 389 | Klebsiella aerogenes |
| GCF_016643825 | 392 | 64 | Klebsiella aerogenes |
| GCF_016643825 | 392 | 389 | Klebsiella aerogenes |
| GCF_016644025 | 392 | 389 | Klebsiella aerogenes |
| GCF_016644035 | 392 | 389 | Klebsiella aerogenes |
| GCF_016939495 | 392 | 389 | Klebsiella aerogenes |
| GCF_017742775 | 392 | 389 | Klebsiella aerogenes |
| GCF_018069155 | 392 | 389 | Klebsiella aerogenes |
| GCF_018278705 | 392 | 389 | Klebsiella aerogenes |
| GCF_018420135 | 392 | 389 | Klebsiella aerogenes |
| GCF_018420295 | 392 | 389 | Klebsiella aerogenes |
| GCF_018420355 | 392 | 389 | Klebsiella aerogenes |
| GCF_018420415 | 392 | 389 | Klebsiella aerogenes |
| GCF_018420895 | 392 | 389 | Klebsiella aerogenes |
| GCF_018420935 | 392 | 389 | Klebsiella aerogenes |
| GCF_018420955 | 392 | 389 | Klebsiella aerogenes |
| GCF_018420975 | 392 | 389 | Klebsiella aerogenes |
| GCF_018421075 | 392 | 389 | Klebsiella aerogenes |
| GCF_018421225 | 392 | 389 | Klebsiella aerogenes |
| GCF_018421335 | 392 | 389 | Klebsiella aerogenes |
| GCF_018421675 | 392 | 389 | Klebsiella aerogenes |
| GCF_018421795 | 392 | 389 | Klebsiella aerogenes |
| GCF_018422045 | 392 | 389 | Klebsiella aerogenes |
| GCF_018422565 | 392 | 389 | Klebsiella aerogenes |
| GCF_018422575 | 392 | 389 | Klebsiella aerogenes |
| GCF_018422885 | 392 | 389 | Klebsiella aerogenes |
| GCF_018422935 | 392 | 389 | Klebsiella aerogenes |
| GCF_018441025 | 392 | 389 | Klebsiella aerogenes |
| GCF_018443245 | 392 | 389 | Klebsiella aerogenes |
| GCF_018445415 | 392 | 389 | Klebsiella aerogenes |
| GCF_018445685 | 392 | 389 | Klebsiella aerogenes |
| GCF_018445765 | 392 | 389 | Klebsiella aerogenes |
| GCF_018445795 | 392 | 389 | Klebsiella aerogenes |
| GCF_018445865 | 392 | 389 | Klebsiella aerogenes |
| GCF_018445905 | 392 | 389 | Klebsiella aerogenes |
| GCF_018445925 | 392 | 389 | Klebsiella aerogenes |
| GCF_018446235 | 392 | 389 | Klebsiella aerogenes |
| GCF_018446415 | 392 | 389 | Klebsiella aerogenes |
| GCF_018446755 | 392 | 389 | Klebsiella aerogenes |
| GCF_018446775 | 392 | 389 | Klebsiella aerogenes |
| GCF_018446975 | 392 | 389 | Klebsiella aerogenes |
| GCF_018447155 | 392 | 389 | Klebsiella aerogenes |
| GCF_018447335 | 392 | 389 | Klebsiella aerogenes |
| GCF_018447575 | 392 | 389 | Klebsiella aerogenes |
| GCF_018447655 | 392 | 389 | Klebsiella aerogenes |
| GCF_019047925 | 392 | 389 | Klebsiella aerogenes |
| GCF_019048125 | 392 | 389 | Klebsiella aerogenes |
| GCF_019048885 | 392 | 389 | Klebsiella aerogenes |
| GCF_019104165 | 392 | 64 | Klebsiella aerogenes |
| GCF_019104165 | 392 | 389 | Klebsiella aerogenes |
| GCF_019166045 | 392 | 389 | Klebsiella aerogenes |
| GCF_019166205 | 392 | 389 | Klebsiella aerogenes |
| GCF_019731535 | 392 | 389 | Klebsiella aerogenes |
| GCF_019733755 | 392 | 389 | Klebsiella aerogenes |
| GCF_019772015 | 392 | 389 | Klebsiella aerogenes |
| GCF_019797685 | 392 | 389 | Klebsiella aerogenes |
| GCF_019931695 | 392 | 389 | Klebsiella aerogenes |
| GCF_020035255 | 392 | 389 | Klebsiella aerogenes |
| GCF_020035275 | 392 | 389 | Klebsiella aerogenes |
| GCF_020075485 | 392 | 389 | Klebsiella aerogenes |
| GCF_020075555 | 392 | 389 | Klebsiella aerogenes |
| GCF_020075625 | 392 | 389 | Klebsiella aerogenes |
| GCF_020143225 | 392 | 389 | Klebsiella aerogenes |
| GCF_020251765 | 392 | 389 | Klebsiella aerogenes |
| GCF_020511695 | 392 | 389 | Klebsiella aerogenes |
| GCF_020511705 | 392 | 389 | Klebsiella aerogenes |
| GCF_020592115 | 392 | 389 | Klebsiella aerogenes |
| GCF_020592195 | 392 | 389 | Klebsiella aerogenes |
| GCF_020592275 | 392 | 389 | Klebsiella aerogenes |
| GCF_020982565 | 392 | 389 | Klebsiella aerogenes |
| GCF_021416125 | 392 | 389 | Klebsiella aerogenes |
| GCF_021416145 | 392 | 389 | Klebsiella aerogenes |
| GCF_021416165 | 392 | 389 | Klebsiella aerogenes |
| GCF_022341225 | 392 | 389 | Klebsiella aerogenes |
| GCF_022700775 | 392 | 389 | Klebsiella aerogenes |
| GCF_022759585 | 392 | 389 | Klebsiella aerogenes |
| GCF_022759605 | 392 | 389 | Klebsiella aerogenes |
| GCF_023647495 | 392 | 389 | Klebsiella aerogenes |
| GCF_023955405 | 392 | 389 | Klebsiella aerogenes |
| GCF_024168945 | 392 | 389 | Klebsiella aerogenes |
| GCF_024661995 | 392 | 389 | Klebsiella aerogenes |
| GCF_024677785 | 392 | 389 | Klebsiella aerogenes |
| GCF_024677805 | 392 | 389 | Klebsiella aerogenes |
| GCF_024677865 | 392 | 389 | Klebsiella aerogenes |
| GCF_024917575 | 392 | 389 | Klebsiella aerogenes |
| GCF_024918355 | 392 | 389 | Klebsiella aerogenes |
| GCF_025143515 | 392 | 389 | Klebsiella aerogenes |
| GCF_025144035 | 392 | 389 | Klebsiella aerogenes |
| GCF_025146975 | 392 | 389 | Klebsiella aerogenes |
| GCF_025150155 | 392 | 389 | Klebsiella aerogenes |
| GCF_025150205 | 392 | 389 | Klebsiella aerogenes |
| GCF_025150225 | 392 | 389 | Klebsiella aerogenes |
| GCF_025150285 | 392 | 389 | Klebsiella aerogenes |
| GCF_025183105 | 392 | 389 | Klebsiella aerogenes |
| GCF_025183505 | 392 | 389 | Klebsiella aerogenes |
| GCF_025183515 | 392 | 389 | Klebsiella aerogenes |
| GCF_025183645 | 392 | 389 | Klebsiella aerogenes |
| GCF_025218545 | 392 | 389 | Klebsiella aerogenes |
| GCF_025308935 | 392 | 389 | Klebsiella aerogenes |
| GCF_025308955 | 392 | 389 | Klebsiella aerogenes |
| GCF_025377565 | 342 | 389 | Klebsiella aerogenes |
| GCF_025535995 | 392 | 389 | Klebsiella aerogenes |
| GCF_025536055 | 392 | 389 | Klebsiella aerogenes |
| GCF_026702945 | 392 | 389 | Klebsiella aerogenes |
| GCF_026712585 | 392 | 389 | Klebsiella aerogenes |
| GCF_027595505 | 392 | 389 | Klebsiella aerogenes |
| GCF_027595565 | 392 | 389 | Klebsiella aerogenes |
| GCF_027595705 | 392 | 389 | Klebsiella aerogenes |
| GCF_027668365 | 392 | 389 | Klebsiella aerogenes |
| GCF_027885685 | 392 | 389 | Klebsiella aerogenes |
| GCF_028994925 | 392 | 389 | Klebsiella aerogenes |
| GCF_029521985 | 392 | 389 | Klebsiella aerogenes |
| GCF_029717425 | 392 | 389 | Klebsiella aerogenes |
| GCF_029717585 | 392 | 389 | Klebsiella aerogenes |
| GCF_029843285 | 392 | 389 | Klebsiella aerogenes |
| GCF_029955765 | 392 | 389 | Klebsiella aerogenes |
| GCF_029956065 | 392 | 389 | Klebsiella aerogenes |
| GCF_029956165 | 392 | 389 | Klebsiella aerogenes |
| GCF_020526085 | 392 | 389 | Klebsiella africana |
| GCF_002806725 | 392 | 389 | Klebsiella electrica |
| GCF_004312065 | 392 | 389 | Klebsiella electrica |
| GCF_006711645 | 392 | 389 | Klebsiella electrica |
| GCF_009825395 | 392 | 389 | Klebsiella electrica |
| GCF_009825425 | 392 | 389 | Klebsiella electrica |
| GCF_014764465 | 392 | 389 | Klebsiella electrica |
| GCF_015135575 | N | 390 | Klebsiella electrica |
| GCF_030253455 | 392 | 389 | Klebsiella electrica |
| SAMD00195896 | 392 | 389 | Klebsiella electrica |
| SAMN31765424 | 392 | 389 | Klebsiella electrica |
| SAMN34191832 | 392 | 389 | Klebsiella electrica |
| DSM105630 | 392 | 389 | Klebsiella grimontii |
| ERS530422 | 392 | 389 | Klebsiella grimontii |
| GCA_000427015 | 392 | 389 | Klebsiella grimontii |
| GCA_001076805 | 392 | 64 | Klebsiella grimontii |
| GCA_001076805 | 392 | 389 | Klebsiella grimontii |
| GCA_001633115 | 392 | 389 | Klebsiella grimontii |
| GCA_002856195 | 392 | 389 | Klebsiella grimontii |
| GCA_019426485 | 392 | 389 | Klebsiella grimontii |
| GCA_019428125 | 392 | 389 | Klebsiella grimontii |
| GCA_019428165 | 392 | 389 | Klebsiella grimontii |
| GCA_019428245 | 392 | 389 | Klebsiella grimontii |
| GCA_902363155 | 392 | 389 | Klebsiella grimontii |
| GCF_001548355 | 392 | 389 | Klebsiella grimontii |
| GCF_002080105 | 392 | 389 | Klebsiella grimontii |
| GCF_002090195 | 392 | 389 | Klebsiella grimontii |
| GCF_002556465 | 87 | 389 | Klebsiella grimontii |
| GCF_002556465 | 392 | 389 | Klebsiella grimontii |
| GCF_002559635 | 392 | 389 | Klebsiella grimontii |
| GCF_002880715 | 87 | 389 | Klebsiella grimontii |
| GCF_002880715 | 392 | 389 | Klebsiella grimontii |
| GCF_003339485 | 392 | 389 | Klebsiella grimontii |
| GCF_003416995 | 87 | 389 | Klebsiella grimontii |
| GCF_003416995 | 392 | 389 | Klebsiella grimontii |
| GCF_003417035 | 87 | 389 | Klebsiella grimontii |
| GCF_003417035 | 392 | 389 | Klebsiella grimontii |
| GCF_004104525 | 392 | 389 | Klebsiella grimontii |
| GCF_004343645 | 392 | 389 | Klebsiella grimontii |
| GCF_008120425 | 392 | 389 | Klebsiella grimontii |
| GCF_008120465 | 392 | 389 | Klebsiella grimontii |
| GCF_008120915 | 392 | 389 | Klebsiella grimontii |
| GCF_008364425 | 87 | 389 | Klebsiella grimontii |
| GCF_008364425 | 392 | 389 | Klebsiella grimontii |
| GCF_009905335 | 87 | 389 | Klebsiella grimontii |
| GCF_009905335 | 392 | 389 | Klebsiella grimontii |
| GCF_014655015 | 392 | 389 | Klebsiella grimontii |
| GCF_015183075 | 392 | 389 | Klebsiella grimontii |
| GCF_015208375 | 392 | 389 | Klebsiella grimontii |
| GCF_016653045 | 392 | 389 | Klebsiella grimontii |
| GCF_017863855 | 87 | 389 | Klebsiella grimontii |
| GCF_017863855 | 392 | 389 | Klebsiella grimontii |
| GCF_017863935 | 87 | 389 | Klebsiella grimontii |
| GCF_017863935 | 392 | 389 | Klebsiella grimontii |
| GCF_019334465 | 392 | 389 | Klebsiella grimontii |
| GCF_019428325 | 392 | 389 | Klebsiella grimontii |
| GCF_019428385 | 392 | 389 | Klebsiella grimontii |
| GCF_019677835 | 392 | 64 | Klebsiella grimontii |
| GCF_019677835 | 392 | 389 | Klebsiella grimontii |
| GCF_019678325 | 392 | 389 | Klebsiella grimontii |
| GCF_019678365 | 87 | 389 | Klebsiella grimontii |
| GCF_019678365 | 392 | 389 | Klebsiella grimontii |
| GCF_019679055 | 392 | 389 | Klebsiella grimontii |
| GCF_019679095 | 87 | 389 | Klebsiella grimontii |
| GCF_019679095 | 392 | 389 | Klebsiella grimontii |
| GCF_020479545 | 87 | 389 | Klebsiella grimontii |
| GCF_020479545 | 392 | 389 | Klebsiella grimontii |
| GCF_020889225 | 392 | 64 | Klebsiella grimontii |
| GCF_020889225 | 392 | 389 | Klebsiella grimontii |
| GCF_022014295 | 392 | 389 | Klebsiella grimontii |
| GCF_022539845 | 392 | 389 | Klebsiella grimontii |
| GCF_022539995 | 392 | 389 | Klebsiella grimontii |
| GCF_022540165 | 392 | 389 | Klebsiella grimontii |
| GCF_022543655 | 392 | 389 | Klebsiella grimontii |
| GCF_022543715 | 392 | 389 | Klebsiella grimontii |
| GCF_022543725 | 392 | 389 | Klebsiella grimontii |
| GCF_022543755 | 392 | 389 | Klebsiella grimontii |
| GCF_022543775 | 392 | 389 | Klebsiella grimontii |
| GCF_022556395 | 392 | 389 | Klebsiella grimontii |
| GCF_022559855 | 392 | 389 | Klebsiella grimontii |
| GCF_026223915 | 392 | 389 | Klebsiella grimontii |
| GCF_026223935 | 392 | 389 | Klebsiella grimontii |
| GCF_028622675 | 392 | 389 | Klebsiella grimontii |
| GCF_028748965 | 392 | 389 | Klebsiella grimontii |
| GCF_028768445 | 392 | 389 | Klebsiella grimontii |
| GCF_030216235 | 87 | 389 | Klebsiella grimontii |
| GCF_030216235 | 392 | 389 | Klebsiella grimontii |
| GCF_030323065 | 392 | 389 | Klebsiella grimontii |
| GCF_030343875 | 87 | 389 | Klebsiella grimontii |
| GCF_030343875 | 392 | 389 | Klebsiella grimontii |
| GCF_030358075 | 87 | 389 | Klebsiella grimontii |
| GCF_030358075 | 392 | 389 | Klebsiella grimontii |
| GCF_030359985 | 87 | 389 | Klebsiella grimontii |
| GCF_030359985 | 392 | 389 | Klebsiella grimontii |
| GCF_030360065 | 87 | 389 | Klebsiella grimontii |
| GCF_030360065 | 392 | 389 | Klebsiella grimontii |
| GCF_030360115 | 87 | 389 | Klebsiella grimontii |
| GCF_030360115 | 392 | 389 | Klebsiella grimontii |
| GCF_030360165 | 392 | 389 | Klebsiella grimontii |
| GCF_032290465 | 392 | 389 | Klebsiella grimontii |
| GCF_032670645 | 392 | 389 | Klebsiella grimontii |
| GCF_032670725 | 392 | 389 | Klebsiella grimontii |
| GCF_032670805 | 392 | 389 | Klebsiella grimontii |
| GCF_032670845 | 392 | 389 | Klebsiella grimontii |
| GCF_032670895 | 392 | 389 | Klebsiella grimontii |
| GCF_032676725 | 392 | 389 | Klebsiella grimontii |
| GCF_032676825 | 392 | 389 | Klebsiella grimontii |
| GCF_032677065 | 392 | 389 | Klebsiella grimontii |
| GCF_032677145 | 392 | 389 | Klebsiella grimontii |
| GCF_032677205 | 392 | 389 | Klebsiella grimontii |
| GCF_032677365 | 392 | 389 | Klebsiella grimontii |
| GCF_032677435 | 87 | 389 | Klebsiella grimontii |
| GCF_032677435 | 392 | 389 | Klebsiella grimontii |
| GCF_032677575 | 392 | 389 | Klebsiella grimontii |
| GCF_032742375 | 392 | 389 | Klebsiella grimontii |
| GCF_032742415 | 392 | 389 | Klebsiella grimontii |
| GCF_032742455 | 392 | 389 | Klebsiella grimontii |
| GCF_032742475 | 392 | 389 | Klebsiella grimontii |
| GCF_032742495 | 392 | 389 | Klebsiella grimontii |
| GCF_032742515 | 392 | 389 | Klebsiella grimontii |
| GCF_032743635 | 392 | 389 | Klebsiella grimontii |
| GCF_032803765 | 392 | 389 | Klebsiella grimontii |
| GCF_035792875 | 392 | 389 | Klebsiella grimontii |
| GCF_035797335 | 392 | 389 | Klebsiella grimontii |
| GCF_036668975 | 87 | 389 | Klebsiella grimontii |
| GCF_036668975 | 392 | 389 | Klebsiella grimontii |
| GCF_037002785 | 392 | 389 | Klebsiella grimontii |
| GCF_038039645 | 392 | 389 | Klebsiella grimontii |
| GCF_038185285 | 392 | 389 | Klebsiella grimontii |
| GCF_038185335 | 392 | 389 | Klebsiella grimontii |
| GCF_038185545 | 392 | 389 | Klebsiella grimontii |
| GCF_038737905 | 392 | 389 | Klebsiella grimontii |
| GCF_038737965 | 392 | 389 | Klebsiella grimontii |
| GCF_038738055 | 392 | 389 | Klebsiella grimontii |
| GCF_040062175 | 392 | 389 | Klebsiella grimontii |
| GCF_040270285 | 392 | 389 | Klebsiella grimontii |
| GCF_040838935 | 392 | 389 | Klebsiella grimontii |
| GCF_042137965 | 87 | 389 | Klebsiella grimontii |
| GCF_042137965 | 392 | 389 | Klebsiella grimontii |
| GCF_043833465 | 87 | 389 | Klebsiella grimontii |
| GCF_043833465 | 392 | 389 | Klebsiella grimontii |
| GCF_046299175 | 392 | 389 | Klebsiella grimontii |
| GCF_046366225 | 392 | 389 | Klebsiella grimontii |
| GCF_046602395 | 87 | 389 | Klebsiella grimontii |
| GCF_046602395 | 392 | 389 | Klebsiella grimontii |
| GCF_046602435 | 87 | 389 | Klebsiella grimontii |
| GCF_046602435 | 392 | 389 | Klebsiella grimontii |
| GCF_046603235 | 392 | 389 | Klebsiella grimontii |
| GCF_900200035 | 392 | 389 | Klebsiella grimontii |
| GCF_900451335 | 81 | 389 | Klebsiella grimontii |
| GCF_900451335 | 170 | 389 | Klebsiella grimontii |
| GCF_901542455 | 392 | 389 | Klebsiella grimontii |
| GCF_902159485 | 392 | 389 | Klebsiella grimontii |
| GCF_902159585 | 392 | 389 | Klebsiella grimontii |
| GCF_902159595 | 392 | 389 | Klebsiella grimontii |
| GCF_902159665 | 392 | 389 | Klebsiella grimontii |
| GCF_902159715 | 392 | 389 | Klebsiella grimontii |
| GCF_902160195 | 392 | 389 | Klebsiella grimontii |
| GCF_902160335 | 392 | 389 | Klebsiella grimontii |
| GCF_902160345 | 392 | 389 | Klebsiella grimontii |
| GCF_902160365 | 392 | 389 | Klebsiella grimontii |
| GCF_902160515 | 392 | 389 | Klebsiella grimontii |
| GCF_902160585 | 392 | 389 | Klebsiella grimontii |
| GCF_902160675 | 87 | 389 | Klebsiella grimontii |
| GCF_902160675 | 392 | 389 | Klebsiella grimontii |
| GCF_902160755 | 392 | 389 | Klebsiella grimontii |
| GCF_902161285 | 392 | 389 | Klebsiella grimontii |
| GCF_902162695 | 392 | 389 | Klebsiella grimontii |
| GCF_902163185 | 392 | 389 | Klebsiella grimontii |
| GCF_902164065 | 87 | 389 | Klebsiella grimontii |
| GCF_902164065 | 392 | 389 | Klebsiella grimontii |
| GCF_902164185 | 392 | 389 | Klebsiella grimontii |
| GCF_902164675 | 392 | 389 | Klebsiella grimontii |
| GCF_902164895 | 392 | 389 | Klebsiella grimontii |
| GCF_902166675 | 392 | 389 | Klebsiella grimontii |
| GCF_902166735 | 392 | 389 | Klebsiella grimontii |
| GFKo12 | 392 | 64 | Klebsiella grimontii |
| GFKo12 | 392 | 389 | Klebsiella grimontii |
| GFKo2 | 392 | 389 | Klebsiella grimontii |
| GFKo20 | 392 | 389 | Klebsiella grimontii |
| GFKo21 | 392 | 389 | Klebsiella grimontii |
| GFKo22 | 392 | 389 | Klebsiella grimontii |
| GFKo26 | 392 | 389 | Klebsiella grimontii |
| GFKo27 | 392 | 389 | Klebsiella grimontii |
| GFKo3 | 392 | 389 | Klebsiella grimontii |
| GFKo33 | 392 | 389 | Klebsiella grimontii |
| GFKo34 | 392 | 389 | Klebsiella grimontii |
| GFKo39 | 87 | 389 | Klebsiella grimontii |
| GFKo39 | 392 | 389 | Klebsiella grimontii |
| GFKo5 | 392 | 389 | Klebsiella grimontii |
| Ko16 | 87 | 389 | Klebsiella grimontii |
| Ko16 | 392 | 389 | Klebsiella grimontii |
| Ko27 | 392 | 64 | Klebsiella grimontii |
| Ko27 | 392 | 389 | Klebsiella grimontii |
| Ko30 | 392 | 389 | Klebsiella grimontii |
| Ko42 | 87 | 389 | Klebsiella grimontii |
| Ko42 | 392 | 389 | Klebsiella grimontii |
| Ko51 | 392 | 389 | Klebsiella grimontii |
| Ko8 | 87 | 389 | Klebsiella grimontii |
| Ko8 | 392 | 389 | Klebsiella grimontii |
| Ko9 | 392 | 389 | Klebsiella grimontii |
| GCF_026735415 | 392 | N | Klebsiella huaxiensis |
| GCF_038737695 | 392 | N | Klebsiella huaxiensis |
| GCF_902158605 | 392 | N | Klebsiella huaxiensis |
| GCF_902158625 | 392 | N | Klebsiella huaxiensis |
| GCF_036441055 | N | N | Klebsiella huaxiensis |
| GCF_005860775 | 392 | N | Klebsiella indica |
| GCF_039409735 | 392 | N | Klebsiella indica |
| GCF_043829405 | 392 | N | Klebsiella indica |
| GCF_043832615 | 392 | N | Klebsiella indica |
| GCF_900198535 | 392 | N | Klebsiella indica |
| GCF_036561965 | 392 | 389 | Klebsiella lignicola |
| SAMD00320942 | 392 | 389 | Klebsiella lignicola |
| SAMD00320943 | 392 | 389 | Klebsiella lignicola |
| SAMD00320977 | 392 | 389 | Klebsiella lignicola |
| SAMEA6574526 | 392 | 389 | Klebsiella lignicola |
| SAMEA6574529 | 392 | 389 | Klebsiella lignicola |
| SAMN10361637 | 392 | 389 | Klebsiella lignicola |
| SAMN13243134 | 392 | 389 | Klebsiella lignicola |
| SAMN15567066 | 392 | 389 | Klebsiella lignicola |
| SAMN15567068 | 392 | 389 | Klebsiella lignicola |
| SAMN15567071 | 392 | 389 | Klebsiella lignicola |
| SAMN15567072 | 392 | 389 | Klebsiella lignicola |
| SAMN20391770 | 392 | 389 | Klebsiella lignicola |
| DSM25444 | 392 | 389 | Klebsiella michiganensis |
| GCA_009173485 | 392 | 389 | Klebsiella michiganensis |
| GCF_000240325 | 392 | 389 | Klebsiella michiganensis |
| GCF_000247835 | 392 | 389 | Klebsiella michiganensis |
| GCF_000276705 | 392 | 389 | Klebsiella michiganensis |
| GCF_000293135 | 392 | 389 | Klebsiella michiganensis |
| GCF_000524315 | 392 | 389 | Klebsiella michiganensis |
| GCF_000632415 | 392 | 389 | Klebsiella michiganensis |
| GCF_000633235 | 392 | 389 | Klebsiella michiganensis |
| GCF_000724525 | 392 | 389 | Klebsiella michiganensis |
| GCF_000735215 | 392 | 389 | Klebsiella michiganensis |
| GCF_000783895 | 392 | 389 | Klebsiella michiganensis |
| GCF_001006545 | 392 | 389 | Klebsiella michiganensis |
| GCF_001028875 | 392 | 389 | Klebsiella michiganensis |
| GCF_001028885 | 392 | 389 | Klebsiella michiganensis |
| GCF_001038305 | 392 | 389 | Klebsiella michiganensis |
| GCF_001051455 | 392 | 389 | Klebsiella michiganensis |
| GCF_001056765 | 392 | 389 | Klebsiella michiganensis |
| GCF_001077175 | 392 | 389 | Klebsiella michiganensis |
| GCF_001583485 | 392 | 389 | Klebsiella michiganensis |
| GCF_001753185 | 392 | 389 | Klebsiella michiganensis |
| GCF_001945455 | 392 | 389 | Klebsiella michiganensis |
| GCF_002111445 | 392 | 389 | Klebsiella michiganensis |
| GCF_002119875 | 392 | 389 | Klebsiella michiganensis |
| GCF_002192755 | 392 | 389 | Klebsiella michiganensis |
| GCF_002216835 | 392 | 389 | Klebsiella michiganensis |
| GCF_002265195 | 392 | 389 | Klebsiella michiganensis |
| GCF_002290285 | 392 | 389 | Klebsiella michiganensis |
| GCF_002856965 | 392 | 155 | Klebsiella michiganensis |
| GCF_002887165 | 392 | 389 | Klebsiella michiganensis |
| GCF_002887605 | 392 | 389 | Klebsiella michiganensis |
| GCF_003011775 | 392 | 389 | Klebsiella michiganensis |
| GCF_003402095 | 392 | 389 | Klebsiella michiganensis |
| GCF_003590255 | 392 | 389 | Klebsiella michiganensis |
| GCF_003598595 | 392 | 389 | Klebsiella michiganensis |
| GCF_004024205 | 392 | 389 | Klebsiella michiganensis |
| GCF_004024445 | 392 | 389 | Klebsiella michiganensis |
| GCF_004102625 | 392 | 389 | Klebsiella michiganensis |
| GCF_004115315 | 392 | 389 | Klebsiella michiganensis |
| GCF_007097115 | 392 | 389 | Klebsiella michiganensis |
| GCF_007106885 | 392 | 389 | Klebsiella michiganensis |
| GCF_007910085 | 392 | 389 | Klebsiella michiganensis |
| GCF_008120085 | 392 | 389 | Klebsiella michiganensis |
| GCF_008120305 | 392 | 389 | Klebsiella michiganensis |
| GCF_008121175 | 392 | 389 | Klebsiella michiganensis |
| GCF_008931605 | 392 | 389 | Klebsiella michiganensis |
| GCF_009025755 | 392 | 389 | Klebsiella michiganensis |
| GCF_009825595 | 392 | 389 | Klebsiella michiganensis |
| GCF_009930855 | 392 | 389 | Klebsiella michiganensis |
| GCF_010093005 | 392 | 389 | Klebsiella michiganensis |
| GCF_013074375 | 392 | 389 | Klebsiella michiganensis |
| GCF_013266825 | 392 | 389 | Klebsiella michiganensis |
| GCF_013392155 | 392 | 389 | Klebsiella michiganensis |
| GCF_013821765 | 392 | 389 | Klebsiella michiganensis |
| GCF_014050515 | 392 | 389 | Klebsiella michiganensis |
| GCF_014050535 | 392 | 389 | Klebsiella michiganensis |
| GCF_014050555 | 392 | 389 | Klebsiella michiganensis |
| GCF_014129405 | 392 | 389 | Klebsiella michiganensis |
| GCF_014330695 | 392 | 389 | Klebsiella michiganensis |
| GCF_014654995 | 392 | 389 | Klebsiella michiganensis |
| GCF_014655005 | 392 | 389 | Klebsiella michiganensis |
| GCF_014655075 | 392 | 389 | Klebsiella michiganensis |
| GCF_014655105 | 392 | 389 | Klebsiella michiganensis |
| GCF_014655115 | 392 | 389 | Klebsiella michiganensis |
| GCF_014855615 | 392 | 389 | Klebsiella michiganensis |
| GCF_014856135 | 392 | 389 | Klebsiella michiganensis |
| GCF_014856145 | 392 | 389 | Klebsiella michiganensis |
| GCF_014856335 | 392 | 389 | Klebsiella michiganensis |
| GCF_014856345 | 392 | 389 | Klebsiella michiganensis |
| GCF_014856365 | 392 | 389 | Klebsiella michiganensis |
| GCF_014856375 | 392 | 389 | Klebsiella michiganensis |
| GCF_014856385 | 392 | 389 | Klebsiella michiganensis |
| GCF_014856435 | 392 | 389 | Klebsiella michiganensis |
| GCF_015356395 | 392 | 389 | Klebsiella michiganensis |
| GCF_015356405 | 392 | 389 | Klebsiella michiganensis |
| GCF_015550995 | 392 | 389 | Klebsiella michiganensis |
| GCF_015773165 | 392 | 389 | Klebsiella michiganensis |
| GCF_015999385 | 392 | 389 | Klebsiella michiganensis |
| GCF_016618215 | 392 | 389 | Klebsiella michiganensis |
| GCF_016652915 | 392 | 389 | Klebsiella michiganensis |
| GCF_016652925 | 392 | 389 | Klebsiella michiganensis |
| GCF_016652965 | 392 | 389 | Klebsiella michiganensis |
| GCF_016652995 | 392 | 389 | Klebsiella michiganensis |
| GCF_016653015 | 392 | 389 | Klebsiella michiganensis |
| GCF_016653035 | 392 | 389 | Klebsiella michiganensis |
| GCF_016653085 | 392 | 389 | Klebsiella michiganensis |
| GCF_016734915 | 392 | 389 | Klebsiella michiganensis |
| GCF_016734965 | 392 | 389 | Klebsiella michiganensis |
| GCF_016734985 | 392 | 389 | Klebsiella michiganensis |
| GCF_016735005 | 392 | 389 | Klebsiella michiganensis |
| GCF_016905825 | 392 | 389 | Klebsiella michiganensis |
| GCF_017114595 | 392 | 389 | Klebsiella michiganensis |
| GCF_017348855 | 392 | 389 | Klebsiella michiganensis |
| GCF_017639895 | 392 | 389 | Klebsiella michiganensis |
| GCF_017798285 | 392 | 389 | Klebsiella michiganensis |
| GCF_017810085 | 392 | 389 | Klebsiella michiganensis |
| GCF_017815775 | 392 | 389 | Klebsiella michiganensis |
| GCF_018092585 | 392 | 389 | Klebsiella michiganensis |
| GCF_018092635 | 392 | 389 | Klebsiella michiganensis |
| GCF_018140945 | 392 | 389 | Klebsiella michiganensis |
| GCF_018422065 | 392 | 389 | Klebsiella michiganensis |
| GCF_018422165 | 69 | 389 | Klebsiella michiganensis |
| GCF_018438865 | 392 | 389 | Klebsiella michiganensis |
| GCF_018439305 | 392 | 389 | Klebsiella michiganensis |
| GCF_018439315 | 392 | 389 | Klebsiella michiganensis |
| GCF_018439335 | 392 | 389 | Klebsiella michiganensis |
| GCF_018439355 | 392 | 389 | Klebsiella michiganensis |
| GCF_018439405 | 392 | 389 | Klebsiella michiganensis |
| GCF_018440985 | 392 | 389 | Klebsiella michiganensis |
| GCF_018442305 | 392 | 389 | Klebsiella michiganensis |
| GCF_018442325 | 392 | 389 | Klebsiella michiganensis |
| GCF_018443125 | 392 | 389 | Klebsiella michiganensis |
| GCF_018604105 | 392 | 389 | Klebsiella michiganensis |
| GCF_019048985 | 392 | 60 | Klebsiella michiganensis |
| GCF_019048985 | 392 | 389 | Klebsiella michiganensis |
| GCF_019050695 | 392 | 389 | Klebsiella michiganensis |
| GCF_019378535 | 392 | 389 | Klebsiella michiganensis |
| GCF_019378655 | 392 | 389 | Klebsiella michiganensis |
| GCF_019378695 | 392 | 389 | Klebsiella michiganensis |
| GCF_019426465 | 392 | 389 | Klebsiella michiganensis |
| GCF_019428105 | 392 | 389 | Klebsiella michiganensis |
| GCF_019428145 | 392 | 389 | Klebsiella michiganensis |
| GCF_019428225 | 392 | 389 | Klebsiella michiganensis |
| GCF_019460145 | 392 | 389 | Klebsiella michiganensis |
| GCF_019678415 | 392 | 389 | Klebsiella michiganensis |
| GCF_019678495 | 392 | 389 | Klebsiella michiganensis |
| GCF_019678505 | 392 | 389 | Klebsiella michiganensis |
| GCF_019678825 | 392 | 389 | Klebsiella michiganensis |
| GCF_019679175 | 392 | 389 | Klebsiella michiganensis |
| GCF_019730455 | 392 | 389 | Klebsiella michiganensis |
| GCF_019730515 | 392 | 389 | Klebsiella michiganensis |
| GCF_019730535 | 392 | 389 | Klebsiella michiganensis |
| GCF_019730985 | 392 | 389 | Klebsiella michiganensis |
| GCF_019731925 | 392 | 389 | Klebsiella michiganensis |
| GCF_019754075 | 392 | 389 | Klebsiella michiganensis |
| GCF_019754135 | 392 | 389 | Klebsiella michiganensis |
| GCF_019803065 | 392 | 389 | Klebsiella michiganensis |
| GCF_020117545 | 392 | 389 | Klebsiella michiganensis |
| GCF_020479805 | 392 | 389 | Klebsiella michiganensis |
| GCF_020695625 | 392 | 389 | Klebsiella michiganensis |
| GCF_021228855 | 392 | 389 | Klebsiella michiganensis |
| GCF_021228995 | 392 | 389 | Klebsiella michiganensis |
| GCF_021391575 | 392 | 389 | Klebsiella michiganensis |
| GCF_021440665 | 392 | 389 | Klebsiella michiganensis |
| GCF_021460075 | 392 | 389 | Klebsiella michiganensis |
| GCF_022163245 | 392 | 389 | Klebsiella michiganensis |
| GCF_022343245 | 392 | 389 | Klebsiella michiganensis |
| GCF_022343265 | 392 | 389 | Klebsiella michiganensis |
| GCF_022501085 | 392 | 389 | Klebsiella michiganensis |
| GCF_022501105 | 392 | 389 | Klebsiella michiganensis |
| GCF_022539825 | 392 | 389 | Klebsiella michiganensis |
| GCF_022539865 | 392 | 389 | Klebsiella michiganensis |
| GCF_022539885 | 392 | 389 | Klebsiella michiganensis |
| GCF_022539905 | 392 | 389 | Klebsiella michiganensis |
| GCF_022539925 | 392 | 389 | Klebsiella michiganensis |
| GCF_022539945 | 392 | 389 | Klebsiella michiganensis |
| GCF_022539965 | 392 | 389 | Klebsiella michiganensis |
| GCF_022540025 | 392 | 389 | Klebsiella michiganensis |
| GCF_022540045 | 392 | 389 | Klebsiella michiganensis |
| GCF_022540065 | 392 | 389 | Klebsiella michiganensis |
| GCF_022540085 | 392 | 389 | Klebsiella michiganensis |
| GCF_022540105 | 392 | 389 | Klebsiella michiganensis |
| GCF_022540125 | 392 | 389 | Klebsiella michiganensis |
| GCF_022543495 | 392 | 389 | Klebsiella michiganensis |
| GCF_022543535 | 392 | 389 | Klebsiella michiganensis |
| GCF_022543555 | 392 | 389 | Klebsiella michiganensis |
| GCF_022543575 | 392 | 389 | Klebsiella michiganensis |
| GCF_022543595 | 392 | 389 | Klebsiella michiganensis |
| GCF_022543615 | N | 389 | Klebsiella michiganensis |
| GCF_022543625 | 392 | 389 | Klebsiella michiganensis |
| GCF_022543675 | 392 | 389 | Klebsiella michiganensis |
| GCF_022543695 | 392 | 389 | Klebsiella michiganensis |
| GCF_022543785 | 392 | 389 | Klebsiella michiganensis |
| GCF_022543815 | 392 | 389 | Klebsiella michiganensis |
| GCF_022543825 | 392 | 389 | Klebsiella michiganensis |
| GCF_022543855 | 392 | 389 | Klebsiella michiganensis |
| GCF_022551435 | 392 | 389 | Klebsiella michiganensis |
| GCF_022569835 | 392 | 389 | Klebsiella michiganensis |
| GCF_022859475 | 392 | 389 | Klebsiella michiganensis |
| GCF_022869885 | 392 | 389 | Klebsiella michiganensis |
| GCF_023093655 | 392 | 389 | Klebsiella michiganensis |
| GCF_023093835 | 392 | 389 | Klebsiella michiganensis |
| GCF_023502385 | 392 | 389 | Klebsiella michiganensis |
| GCF_024499915 | 392 | 389 | Klebsiella michiganensis |
| GCF_024542875 | 392 | 389 | Klebsiella michiganensis |
| GCF_024586865 | N | 389 | Klebsiella michiganensis |
| GCF_025263805 | 392 | 389 | Klebsiella michiganensis |
| GCF_025490535 | 392 | 389 | Klebsiella michiganensis |
| GCF_025665355 | 392 | 389 | Klebsiella michiganensis |
| GCF_025960245 | 392 | 389 | Klebsiella michiganensis |
| GCF_026162305 | 392 | 389 | Klebsiella michiganensis |
| GCF_026162385 | 392 | 389 | Klebsiella michiganensis |
| GCF_026223635 | 392 | 389 | Klebsiella michiganensis |
| GCF_026223955 | 392 | 389 | Klebsiella michiganensis |
| GCF_026223965 | 392 | 389 | Klebsiella michiganensis |
| GCF_026224015 | 392 | 389 | Klebsiella michiganensis |
| GCF_026224035 | 392 | 389 | Klebsiella michiganensis |
| GCF_026224055 | 392 | 389 | Klebsiella michiganensis |
| GCF_026224085 | 392 | 389 | Klebsiella michiganensis |
| GCF_026224385 | 392 | 389 | Klebsiella michiganensis |
| GCF_026224395 | 392 | 389 | Klebsiella michiganensis |
| GCF_026224435 | 392 | 389 | Klebsiella michiganensis |
| GCF_026224455 | 392 | 389 | Klebsiella michiganensis |
| GCF_026224605 | 392 | 389 | Klebsiella michiganensis |
| GCF_026224675 | 392 | 389 | Klebsiella michiganensis |
| GCF_026224695 | 392 | 389 | Klebsiella michiganensis |
| GCF_026224715 | 82 | 389 | Klebsiella michiganensis |
| GCF_026224715 | 392 | 389 | Klebsiella michiganensis |
| GCF_026226195 | N | 389 | Klebsiella michiganensis |
| GCF_026274395 | 392 | 389 | Klebsiella michiganensis |
| GCF_026627765 | 392 | 389 | Klebsiella michiganensis |
| GCF_026967695 | 392 | 389 | Klebsiella michiganensis |
| GCF_027577885 | 392 | 389 | Klebsiella michiganensis |
| GCF_027584345 | 392 | 389 | Klebsiella michiganensis |
| GCF_027584475 | 392 | 389 | Klebsiella michiganensis |
| GCF_027673125 | 392 | 389 | Klebsiella michiganensis |
| GCF_027945215 | 377 | 389 | Klebsiella michiganensis |
| GCF_028736015 | 392 | 389 | Klebsiella michiganensis |
| GCF_028737615 | 392 | 389 | Klebsiella michiganensis |
| GCF_028737625 | 392 | 389 | Klebsiella michiganensis |
| GCF_029027885 | 392 | 389 | Klebsiella michiganensis |
| GCF_029960265 | 392 | 389 | Klebsiella michiganensis |
| GCF_029960425 | 392 | 389 | Klebsiella michiganensis |
| GCF_030164605 | 392 | 389 | Klebsiella michiganensis |
| GCF_030164645 | 392 | 389 | Klebsiella michiganensis |
| GCF_030215845 | 392 | 389 | Klebsiella michiganensis |
| GCF_030225565 | 392 | 389 | Klebsiella michiganensis |
| GCF_030248395 | 392 | 389 | Klebsiella michiganensis |
| GCF_030283225 | 392 | 389 | Klebsiella michiganensis |
| GCF_030283425 | 392 | 389 | Klebsiella michiganensis |
| GCF_030283905 | 392 | 389 | Klebsiella michiganensis |
| GCF_030286175 | 392 | 389 | Klebsiella michiganensis |
| GCF_030291695 | 392 | 389 | Klebsiella michiganensis |
| GCF_030294205 | 392 | 389 | Klebsiella michiganensis |
| GCF_030294285 | 392 | 389 | Klebsiella michiganensis |
| GCF_030294485 | 392 | 389 | Klebsiella michiganensis |
| GCF_030342845 | 392 | 389 | Klebsiella michiganensis |
| GCF_030342925 | 392 | 389 | Klebsiella michiganensis |
| GCF_030342935 | 392 | 389 | Klebsiella michiganensis |
| GCF_030343745 | 392 | 389 | Klebsiella michiganensis |
| GCF_030343755 | 392 | 389 | Klebsiella michiganensis |
| GCF_030344115 | 392 | 389 | Klebsiella michiganensis |
| GCF_030344305 | 392 | 64 | Klebsiella michiganensis |
| GCF_030344305 | 392 | 389 | Klebsiella michiganensis |
| GCF_030344365 | 392 | 389 | Klebsiella michiganensis |
| GCF_030344445 | 392 | 389 | Klebsiella michiganensis |
| GCF_030344485 | 392 | 389 | Klebsiella michiganensis |
| GCF_030358065 | 392 | 389 | Klebsiella michiganensis |
| GCF_030358825 | 392 | 389 | Klebsiella michiganensis |
| GCF_030358865 | 392 | 389 | Klebsiella michiganensis |
| GCF_030358885 | 392 | 389 | Klebsiella michiganensis |
| GCF_030358905 | 392 | 389 | Klebsiella michiganensis |
| GCF_030358925 | 392 | 389 | Klebsiella michiganensis |
| GCF_030358945 | 392 | 389 | Klebsiella michiganensis |
| GCF_030358965 | 392 | 389 | Klebsiella michiganensis |
| GCF_030358985 | 392 | 389 | Klebsiella michiganensis |
| GCF_030359005 | 392 | 389 | Klebsiella michiganensis |
| GCF_030359025 | 392 | 389 | Klebsiella michiganensis |
| GCF_030359045 | 392 | 389 | Klebsiella michiganensis |
| GCF_030359065 | 392 | 389 | Klebsiella michiganensis |
| GCF_030359085 | 392 | 389 | Klebsiella michiganensis |
| GCF_030359105 | 392 | 389 | Klebsiella michiganensis |
| GCF_030359125 | 392 | 389 | Klebsiella michiganensis |
| GCF_030359145 | 392 | 389 | Klebsiella michiganensis |
| GCF_030359155 | 392 | 389 | Klebsiella michiganensis |
| GCF_030359185 | 392 | 389 | Klebsiella michiganensis |
| GCF_030359205 | 392 | 389 | Klebsiella michiganensis |
| GCF_030359225 | 392 | 389 | Klebsiella michiganensis |
| GCF_030359245 | 392 | 389 | Klebsiella michiganensis |
| GCF_030359265 | 392 | 389 | Klebsiella michiganensis |
| GCF_030359285 | 392 | 389 | Klebsiella michiganensis |
| GCF_030359305 | 392 | 389 | Klebsiella michiganensis |
| GCF_030359405 | 392 | 389 | Klebsiella michiganensis |
| GCF_030369335 | 392 | 389 | Klebsiella michiganensis |
| GCF_030490325 | 392 | 389 | Klebsiella michiganensis |
| GCF_030490345 | 392 | 389 | Klebsiella michiganensis |
| GCF_030542275 | 392 | 389 | Klebsiella michiganensis |
| GCF_030542365 | 392 | 389 | Klebsiella michiganensis |
| GCF_030542385 | 392 | 389 | Klebsiella michiganensis |
| GCF_030542395 | 392 | 389 | Klebsiella michiganensis |
| GCF_030542425 | 392 | 389 | Klebsiella michiganensis |
| GCF_030717465 | 392 | 64 | Klebsiella michiganensis |
| GCF_030717465 | 392 | 389 | Klebsiella michiganensis |
| GCF_030850045 | 392 | 389 | Klebsiella michiganensis |
| GCF_030851535 | 392 | 389 | Klebsiella michiganensis |
| GCF_030913565 | 392 | 389 | Klebsiella michiganensis |
| GCF_031081595 | 392 | 389 | Klebsiella michiganensis |
| GCF_031799455 | 392 | 389 | Klebsiella michiganensis |
| GCF_031799495 | 392 | 389 | Klebsiella michiganensis |
| GCF_031799515 | 392 | 389 | Klebsiella michiganensis |
| GCF_031799575 | 392 | 389 | Klebsiella michiganensis |
| GCF_031799615 | 392 | 389 | Klebsiella michiganensis |
| GCF_031799735 | 392 | 389 | Klebsiella michiganensis |
| GCF_031799755 | 392 | 389 | Klebsiella michiganensis |
| GCF_031799765 | 392 | 389 | Klebsiella michiganensis |
| GCF_031799915 | 392 | 389 | Klebsiella michiganensis |
| GCF_031799955 | 392 | 389 | Klebsiella michiganensis |
| GCF_031799995 | 392 | 389 | Klebsiella michiganensis |
| GCF_031800055 | 392 | 389 | Klebsiella michiganensis |
| GCF_031800115 | 392 | 389 | Klebsiella michiganensis |
| GCF_031800135 | 392 | 389 | Klebsiella michiganensis |
| GCF_031800275 | 392 | 389 | Klebsiella michiganensis |
| GCF_031800315 | 392 | 389 | Klebsiella michiganensis |
| GCF_032670665 | 392 | 389 | Klebsiella michiganensis |
| GCF_032670685 | 392 | 389 | Klebsiella michiganensis |
| GCF_032671025 | 392 | 389 | Klebsiella michiganensis |
| GCF_032671165 | 392 | 389 | Klebsiella michiganensis |
| GCF_032671185 | 392 | 389 | Klebsiella michiganensis |
| GCF_032671245 | 392 | 389 | Klebsiella michiganensis |
| GCF_032676685 | 392 | 389 | Klebsiella michiganensis |
| GCF_032676705 | 392 | 389 | Klebsiella michiganensis |
| GCF_032676845 | 392 | 389 | Klebsiella michiganensis |
| GCF_032676965 | 392 | 389 | Klebsiella michiganensis |
| GCF_032677005 | 392 | 389 | Klebsiella michiganensis |
| GCF_032677405 | 392 | 389 | Klebsiella michiganensis |
| GCF_032677425 | 392 | 389 | Klebsiella michiganensis |
| GCF_032677565 | 392 | 389 | Klebsiella michiganensis |
| GCF_032743195 | 392 | 389 | Klebsiella michiganensis |
| GCF_032744695 | 392 | 389 | Klebsiella michiganensis |
| GCF_032744755 | 392 | 389 | Klebsiella michiganensis |
| GCF_032745295 | 392 | 389 | Klebsiella michiganensis |
| GCF_032745375 | 392 | 389 | Klebsiella michiganensis |
| GCF_032746795 | 392 | 389 | Klebsiella michiganensis |
| GCF_033031495 | 392 | 389 | Klebsiella michiganensis |
| GCF_033031515 | 392 | 389 | Klebsiella michiganensis |
| GCF_033031585 | 392 | 389 | Klebsiella michiganensis |
| GCF_033099835 | 392 | 389 | Klebsiella michiganensis |
| GCF_033237845 | 392 | 389 | Klebsiella michiganensis |
| GCF_033345475 | 392 | 389 | Klebsiella michiganensis |
| GCF_033345505 | 392 | 389 | Klebsiella michiganensis |
| GCF_033447165 | 392 | 389 | Klebsiella michiganensis |
| GCF_033447175 | 392 | 389 | Klebsiella michiganensis |
| GCF_033843325 | 392 | 389 | Klebsiella michiganensis |
| GCF_034427635 | 392 | 389 | Klebsiella michiganensis |
| GCF_034427675 | 392 | 389 | Klebsiella michiganensis |
| GCF_035786615 | 392 | 389 | Klebsiella michiganensis |
| GCF_035787195 | 392 | 389 | Klebsiella michiganensis |
| GCF_035792955 | 392 | 389 | Klebsiella michiganensis |
| GCF_035793375 | 392 | 389 | Klebsiella michiganensis |
| GCF_035795375 | 392 | 389 | Klebsiella michiganensis |
| GCF_035797695 | 392 | 389 | Klebsiella michiganensis |
| GCF_036324245 | 392 | 389 | Klebsiella michiganensis |
| GCF_036543365 | 392 | 124 | Klebsiella michiganensis |
| GCF_036543365 | 392 | 277 | Klebsiella michiganensis |
| GCF_036880125 | 392 | 389 | Klebsiella michiganensis |
| GCF_036880185 | 392 | 389 | Klebsiella michiganensis |
| GCF_036906935 | 392 | 389 | Klebsiella michiganensis |
| GCF_036907955 | 392 | 389 | Klebsiella michiganensis |
| GCF_036911815 | 392 | 389 | Klebsiella michiganensis |
| GCF_036950865 | 392 | 389 | Klebsiella michiganensis |
| GCF_036951165 | 392 | 389 | Klebsiella michiganensis |
| GCF_036951205 | 392 | 389 | Klebsiella michiganensis |
| GCF_036951965 | 392 | 389 | Klebsiella michiganensis |
| GCF_036952685 | 392 | 389 | Klebsiella michiganensis |
| GCF_036952705 | 392 | 389 | Klebsiella michiganensis |
| GCF_037030755 | 396 | 389 | Klebsiella michiganensis |
| GCF_037030875 | 392 | 389 | Klebsiella michiganensis |
| GCF_037081885 | 392 | 389 | Klebsiella michiganensis |
| GCF_037081955 | 392 | 389 | Klebsiella michiganensis |
| GCF_037145475 | 392 | 389 | Klebsiella michiganensis |
| GCF_037153215 | 392 | 389 | Klebsiella michiganensis |
| GCF_038185235 | 392 | 389 | Klebsiella michiganensis |
| GCF_038185295 | 392 | 389 | Klebsiella michiganensis |
| GCF_038185345 | 392 | 389 | Klebsiella michiganensis |
| GCF_038185415 | 392 | 389 | Klebsiella michiganensis |
| GCF_038185455 | 392 | 389 | Klebsiella michiganensis |
| GCF_038378065 | 392 | 389 | Klebsiella michiganensis |
| GCF_038448985 | 392 | 389 | Klebsiella michiganensis |
| GCF_038737825 | 392 | 389 | Klebsiella michiganensis |
| GCF_038737875 | 392 | 389 | Klebsiella michiganensis |
| GCF_039571305 | 392 | 389 | Klebsiella michiganensis |
| GCF_039571335 | 392 | 389 | Klebsiella michiganensis |
| GCF_039571365 | 392 | 389 | Klebsiella michiganensis |
| GCF_039571385 | 392 | 389 | Klebsiella michiganensis |
| GCF_039571405 | 392 | 389 | Klebsiella michiganensis |
| GCF_039571425 | 392 | 389 | Klebsiella michiganensis |
| GCF_039571465 | 392 | 389 | Klebsiella michiganensis |
| GCF_039631385 | 392 | 64 | Klebsiella michiganensis |
| GCF_039631385 | 392 | 389 | Klebsiella michiganensis |
| GCF_040061575 | 392 | 389 | Klebsiella michiganensis |
| GCF_040061645 | 392 | 389 | Klebsiella michiganensis |
| GCF_040061735 | 392 | 389 | Klebsiella michiganensis |
| GCF_040061795 | 392 | 389 | Klebsiella michiganensis |
| GCF_040062275 | 392 | 389 | Klebsiella michiganensis |
| GCF_040062625 | 392 | 389 | Klebsiella michiganensis |
| GCF_040064575 | 392 | 389 | Klebsiella michiganensis |
| GCF_040064585 | 392 | 389 | Klebsiella michiganensis |
| GCF_040064615 | 392 | 389 | Klebsiella michiganensis |
| GCF_040064695 | 392 | 389 | Klebsiella michiganensis |
| GCF_040065215 | 392 | 389 | Klebsiella michiganensis |
| GCF_040137155 | 392 | 64 | Klebsiella michiganensis |
| GCF_040137155 | 392 | 389 | Klebsiella michiganensis |
| GCF_040138355 | 392 | 389 | Klebsiella michiganensis |
| GCF_040139465 | 392 | 389 | Klebsiella michiganensis |
| GCF_040215115 | 392 | 389 | Klebsiella michiganensis |
| GCF_040558955 | 392 | 389 | Klebsiella michiganensis |
| GCF_040561405 | 392 | 389 | Klebsiella michiganensis |
| GCF_040563745 | 392 | 389 | Klebsiella michiganensis |
| GCF_040565895 | 392 | 389 | Klebsiella michiganensis |
| GCF_040566355 | 392 | 389 | Klebsiella michiganensis |
| GCF_040571345 | 392 | 389 | Klebsiella michiganensis |
| GCF_040741495 | 392 | 389 | Klebsiella michiganensis |
| GCF_040741945 | 392 | 389 | Klebsiella michiganensis |
| GCF_040742085 | 392 | 389 | Klebsiella michiganensis |
| GCF_040742135 | 392 | 389 | Klebsiella michiganensis |
| GCF_040794845 | 392 | 389 | Klebsiella michiganensis |
| GCF_040797325 | 392 | 389 | Klebsiella michiganensis |
| GCF_040984225 | 392 | 389 | Klebsiella michiganensis |
| GCF_041227195 | 392 | 389 | Klebsiella michiganensis |
| GCF_041283485 | 392 | 389 | Klebsiella michiganensis |
| GCF_041283495 | 392 | 389 | Klebsiella michiganensis |
| GCF_041901425 | 392 | 389 | Klebsiella michiganensis |
| GCF_042271465 | 392 | 389 | Klebsiella michiganensis |
| GCF_042274125 | 392 | 389 | Klebsiella michiganensis |
| GCF_043104395 | 392 | 389 | Klebsiella michiganensis |
| GCF_043796495 | 392 | 389 | Klebsiella michiganensis |
| GCF_043828815 | 392 | 389 | Klebsiella michiganensis |
| GCF_043833205 | 392 | 389 | Klebsiella michiganensis |
| GCF_043945205 | N | 389 | Klebsiella michiganensis |
| GCF_043945335 | 392 | 389 | Klebsiella michiganensis |
| GCF_043945415 | 392 | 389 | Klebsiella michiganensis |
| GCF_044510385 | 392 | 389 | Klebsiella michiganensis |
| GCF_044510405 | 392 | 389 | Klebsiella michiganensis |
| GCF_044510425 | 392 | 389 | Klebsiella michiganensis |
| GCF_044510445 | 392 | 389 | Klebsiella michiganensis |
| GCF_044684105 | 392 | 389 | Klebsiella michiganensis |
| GCF_044684125 | 392 | 389 | Klebsiella michiganensis |
| GCF_044684165 | 392 | 389 | Klebsiella michiganensis |
| GCF_045223035 | 392 | 64 | Klebsiella michiganensis |
| GCF_045223035 | 392 | 389 | Klebsiella michiganensis |
| GCF_045223375 | 392 | 64 | Klebsiella michiganensis |
| GCF_045223375 | 392 | 389 | Klebsiella michiganensis |
| GCF_045570595 | 392 | 389 | Klebsiella michiganensis |
| GCF_045571425 | 392 | 389 | Klebsiella michiganensis |
| GCF_045620305 | 392 | 389 | Klebsiella michiganensis |
| GCF_045620345 | 392 | 389 | Klebsiella michiganensis |
| GCF_045620865 | 392 | 389 | Klebsiella michiganensis |
| GCF_045622225 | 392 | 389 | Klebsiella michiganensis |
| GCF_045624065 | 392 | 389 | Klebsiella michiganensis |
| GCF_045624665 | 392 | 389 | Klebsiella michiganensis |
| GCF_045624845 | 392 | 389 | Klebsiella michiganensis |
| GCF_045624905 | 392 | 389 | Klebsiella michiganensis |
| GCF_045624975 | 392 | 389 | Klebsiella michiganensis |
| GCF_045625645 | 392 | 389 | Klebsiella michiganensis |
| GCF_045626745 | 392 | 389 | Klebsiella michiganensis |
| GCF_046296835 | 392 | 389 | Klebsiella michiganensis |
| GCF_046298195 | 392 | 389 | Klebsiella michiganensis |
| GCF_046366165 | 392 | 389 | Klebsiella michiganensis |
| GCF_046366205 | 392 | 389 | Klebsiella michiganensis |
| GCF_046503775 | 392 | 389 | Klebsiella michiganensis |
| GCF_046578745 | 392 | 389 | Klebsiella michiganensis |
| GCF_046580575 | 392 | 389 | Klebsiella michiganensis |
| GCF_046602675 | 392 | 389 | Klebsiella michiganensis |
| GCF_046603135 | 392 | 389 | Klebsiella michiganensis |
| GCF_046603175 | 392 | 389 | Klebsiella michiganensis |
| GCF_046603215 | 392 | 389 | Klebsiella michiganensis |
| GCF_046603275 | 392 | 389 | Klebsiella michiganensis |
| GCF_046603295 | 392 | 389 | Klebsiella michiganensis |
| GCF_046603315 | 392 | 389 | Klebsiella michiganensis |
| GCF_046603335 | 392 | 389 | Klebsiella michiganensis |
| GCF_046603355 | 392 | 389 | Klebsiella michiganensis |
| GCF_900407165 | 392 | 389 | Klebsiella michiganensis |
| GCF_900407255 | 392 | 389 | Klebsiella michiganensis |
| GCF_900451945 | 392 | 81 | Klebsiella michiganensis |
| GCF_901553745 | 392 | 389 | Klebsiella michiganensis |
| GCF_901556995 | 392 | 389 | Klebsiella michiganensis |
| GCF_901563895 | 392 | 389 | Klebsiella michiganensis |
| GCF_902158845 | 392 | 389 | Klebsiella michiganensis |
| GCF_902158915 | 392 | 389 | Klebsiella michiganensis |
| GCF_902159075 | 392 | 389 | Klebsiella michiganensis |
| GCF_902159515 | 392 | 389 | Klebsiella michiganensis |
| GCF_902159535 | 392 | 389 | Klebsiella michiganensis |
| GCF_902159745 | 392 | 389 | Klebsiella michiganensis |
| GCF_902159765 | 392 | 389 | Klebsiella michiganensis |
| GCF_902159775 | 392 | 389 | Klebsiella michiganensis |
| GCF_902159795 | 392 | 389 | Klebsiella michiganensis |
| GCF_902159805 | 392 | 389 | Klebsiella michiganensis |
| GCF_902159815 | 392 | 389 | Klebsiella michiganensis |
| GCF_902159865 | 392 | 389 | Klebsiella michiganensis |
| GCF_902159905 | 392 | 389 | Klebsiella michiganensis |
| GCF_902160685 | 392 | 389 | Klebsiella michiganensis |
| GCF_902162725 | 392 | 389 | Klebsiella michiganensis |
| GCF_902163395 | 392 | 389 | Klebsiella michiganensis |
| GCF_902164365 | 392 | 389 | Klebsiella michiganensis |
| GCF_902164605 | 392 | 389 | Klebsiella michiganensis |
| GCF_902164635 | 392 | 389 | Klebsiella michiganensis |
| GCF_902164935 | 392 | 389 | Klebsiella michiganensis |
| GCF_902164965 | 392 | 389 | Klebsiella michiganensis |
| GCF_902166295 | 392 | 389 | Klebsiella michiganensis |
| GCF_902166415 | 392 | 389 | Klebsiella michiganensis |
| GCF_902166555 | 392 | 389 | Klebsiella michiganensis |
| GCF_902166585 | 392 | 389 | Klebsiella michiganensis |
| GCF_902166625 | 392 | 389 | Klebsiella michiganensis |
| GCF_902166795 | 392 | 389 | Klebsiella michiganensis |
| GCF_902386125 | 392 | 389 | Klebsiella michiganensis |
| GCF_947047385 | 392 | 389 | Klebsiella michiganensis |
| GCF_963887275 | 392 | 389 | Klebsiella michiganensis |
| GCF_964188435 | 392 | 389 | Klebsiella michiganensis |
| GCF_964188455 | 392 | 389 | Klebsiella michiganensis |
| GCF_964188485 | 392 | 389 | Klebsiella michiganensis |
| GCF_964188495 | 392 | 389 | Klebsiella michiganensis |
| GCF_964188505 | 392 | 389 | Klebsiella michiganensis |
| GCF_964207175 | 392 | 389 | Klebsiella michiganensis |
| GCF_964207295 | 392 | 389 | Klebsiella michiganensis |
| GCF_964207475 | 392 | 389 | Klebsiella michiganensis |
| GCF_964207735 | 392 | 389 | Klebsiella michiganensis |
| GCF_964243925 | 392 | 389 | Klebsiella michiganensis |
| GCF_964248175 | 392 | 389 | Klebsiella michiganensis |
| GCF_964270985 | 392 | 389 | Klebsiella michiganensis |
| GCF_964276435 | 392 | 389 | Klebsiella michiganensis |
| GFKo10 | 392 | 389 | Klebsiella michiganensis |
| GFKo13 | 392 | 389 | Klebsiella michiganensis |
| GFKo14 | 392 | 389 | Klebsiella michiganensis |
| GFKo15 | 392 | 64 | Klebsiella michiganensis |
| GFKo15 | 392 | 389 | Klebsiella michiganensis |
| GFKo18 | 392 | 389 | Klebsiella michiganensis |
| GFKo19 | 392 | 389 | Klebsiella michiganensis |
| GFKo28 | 392 | 389 | Klebsiella michiganensis |
| GFKo29 | 392 | 389 | Klebsiella michiganensis |
| GFKo30 | 392 | 389 | Klebsiella michiganensis |
| GFKo31 | 392 | 389 | Klebsiella michiganensis |
| GFKo32 | 392 | 389 | Klebsiella michiganensis |
| GFKo35 | 392 | 389 | Klebsiella michiganensis |
| GFKo36 | 392 | 389 | Klebsiella michiganensis |
| GFKo37 | 392 | 389 | Klebsiella michiganensis |
| GFKo38 | 392 | 389 | Klebsiella michiganensis |
| GFKo40 | 392 | 389 | Klebsiella michiganensis |
| GFKo41 | 392 | 389 | Klebsiella michiganensis |
| GFKo44 | 392 | 389 | Klebsiella michiganensis |
| GFKo45 | 392 | 389 | Klebsiella michiganensis |
| GFKo7 | 392 | 389 | Klebsiella michiganensis |
| GFKo8 | 392 | 389 | Klebsiella michiganensis |
| Ko10 | 392 | 389 | Klebsiella michiganensis |
| Ko13 | 392 | 389 | Klebsiella michiganensis |
| Ko14 | 392 | 389 | Klebsiella michiganensis |
| Ko18 | 392 | 389 | Klebsiella michiganensis |
| Ko21 | 392 | 389 | Klebsiella michiganensis |
| Ko22 | 392 | 389 | Klebsiella michiganensis |
| Ko23 | 392 | 389 | Klebsiella michiganensis |
| Ko24 | 392 | 389 | Klebsiella michiganensis |
| Ko28 | 392 | 389 | Klebsiella michiganensis |
| Ko29 | 392 | 389 | Klebsiella michiganensis |
| Ko3 | 392 | 389 | Klebsiella michiganensis |
| Ko31 | 392 | 389 | Klebsiella michiganensis |
| Ko32 | 392 | 389 | Klebsiella michiganensis |
| Ko33 | 392 | 389 | Klebsiella michiganensis |
| Ko35 | 392 | 389 | Klebsiella michiganensis |
| Ko36 | 392 | 389 | Klebsiella michiganensis |
| Ko39 | 392 | 389 | Klebsiella michiganensis |
| Ko41 | 392 | 389 | Klebsiella michiganensis |
| Ko43 | 392 | 389 | Klebsiella michiganensis |
| Ko46 | 392 | 389 | Klebsiella michiganensis |
| Ko49 | 392 | 389 | Klebsiella michiganensis |
| Ko5 | 392 | 389 | Klebsiella michiganensis |
| Ko50 | 392 | 389 | Klebsiella michiganensis |
| Ko52 | 392 | 389 | Klebsiella michiganensis |
| Ko56 | 392 | 389 | Klebsiella michiganensis |
| Ko58 | 392 | 389 | Klebsiella michiganensis |
| Ko59 | 392 | 389 | Klebsiella michiganensis |
| GCF_000247895 | 392 | 389 | Klebsiella ornithinolytica |
| GCF_000367425 | 392 | 389 | Klebsiella ornithinolytica |
| GCF_000703485 | 392 | 389 | Klebsiella ornithinolytica |
| GCF_000935285 | 392 | 389 | Klebsiella ornithinolytica |
| GCF_001039315 | 392 | 389 | Klebsiella ornithinolytica |
| GCF_001066395 | 392 | 389 | Klebsiella ornithinolytica |
| GCF_001455225 | 392 | 389 | Klebsiella ornithinolytica |
| GCF_001598295 | 392 | 389 | Klebsiella ornithinolytica |
| GCF_001700855 | 392 | 389 | Klebsiella ornithinolytica |
| GCF_001723565 | 392 | 389 | Klebsiella ornithinolytica |
| GCF_001866535 | 392 | 64 | Klebsiella ornithinolytica |
| GCF_001866535 | 392 | 389 | Klebsiella ornithinolytica |
| GCF_002214825 | 392 | 389 | Klebsiella ornithinolytica |
| GCF_002266585 | 392 | 389 | Klebsiella ornithinolytica |
| GCF_002266645 | 392 | 389 | Klebsiella ornithinolytica |
| GCF_002266665 | 392 | 389 | Klebsiella ornithinolytica |
| GCF_002266675 | 392 | 389 | Klebsiella ornithinolytica |
| GCF_002266685 | 392 | 389 | Klebsiella ornithinolytica |
| GCF_002266695 | 392 | 389 | Klebsiella ornithinolytica |
| GCF_002494395 | 392 | 389 | Klebsiella ornithinolytica |
| GCF_002635365 | 392 | 389 | Klebsiella ornithinolytica |
| GCF_002759495 | 392 | 389 | Klebsiella ornithinolytica |
| GCF_002794395 | 392 | 389 | Klebsiella ornithinolytica |
| GCF_002806925 | 392 | 389 | Klebsiella ornithinolytica |
| GCF_002895535 | 392 | 389 | Klebsiella ornithinolytica |
| GCF_002949305 | 392 | 389 | Klebsiella ornithinolytica |
| GCF_002949335 | 392 | 389 | Klebsiella ornithinolytica |
| GCF_002949375 | 392 | 389 | Klebsiella ornithinolytica |
| GCF_002949385 | 392 | 389 | Klebsiella ornithinolytica |
| GCF_002949435 | 392 | 389 | Klebsiella ornithinolytica |
| GCF_003323835 | 392 | 389 | Klebsiella ornithinolytica |
| GCF_003782005 | 392 | 389 | Klebsiella ornithinolytica |
| GCF_003782015 | 392 | 389 | Klebsiella ornithinolytica |
| GCF_003782025 | 392 | 389 | Klebsiella ornithinolytica |
| GCF_003782085 | 392 | 91 | Klebsiella ornithinolytica |
| GCF_003782095 | 392 | 389 | Klebsiella ornithinolytica |
| GCF_003798165 | 392 | 389 | Klebsiella ornithinolytica |
| GCF_004004825 | 392 | 389 | Klebsiella ornithinolytica |
| GCF_004024085 | 392 | 389 | Klebsiella ornithinolytica |
| GCF_004024535 | 392 | 389 | Klebsiella ornithinolytica |
| GCF_004801135 | 392 | 389 | Klebsiella ornithinolytica |
| GCF_005165545 | 392 | 389 | Klebsiella ornithinolytica |
| GCF_009267985 | 392 | 389 | Klebsiella ornithinolytica |
| GCF_009267995 | 392 | 389 | Klebsiella ornithinolytica |
| GCF_009268015 | 392 | 389 | Klebsiella ornithinolytica |
| GCF_009709685 | 392 | 389 | Klebsiella ornithinolytica |
| GCF_009830315 | 392 | 389 | Klebsiella ornithinolytica |
| GCF_009830325 | 392 | 389 | Klebsiella ornithinolytica |
| GCF_010365445 | 392 | 389 | Klebsiella ornithinolytica |
| GCF_011149255 | 392 | 389 | Klebsiella ornithinolytica |
| GCF_013425975 | 392 | 389 | Klebsiella ornithinolytica |
| GCF_013457615 | 392 | 389 | Klebsiella ornithinolytica |
| GCF_013457875 | 392 | 389 | Klebsiella ornithinolytica |
| GCF_014168795 | 392 | 389 | Klebsiella ornithinolytica |
| GCF_014170855 | 392 | 389 | Klebsiella ornithinolytica |
| GCF_014171135 | 392 | 389 | Klebsiella ornithinolytica |
| GCF_014852555 | 392 | 389 | Klebsiella ornithinolytica |
| GCF_015654305 | 392 | 389 | Klebsiella ornithinolytica |
| GCF_016599735 | 392 | 389 | Klebsiella ornithinolytica |
| GCF_016614655 | 392 | 389 | Klebsiella ornithinolytica |
| GCF_016618175 | 392 | 389 | Klebsiella ornithinolytica |
| GCF_016653655 | 392 | 389 | Klebsiella ornithinolytica |
| GCF_016888105 | 392 | 389 | Klebsiella ornithinolytica |
| GCF_016925155 | 392 | 389 | Klebsiella ornithinolytica |
| GCF_018117465 | 392 | 430 | Klebsiella ornithinolytica |
| GCF_018420695 | 392 | 389 | Klebsiella ornithinolytica |
| GCF_018438815 | 53 | 306 | Klebsiella ornithinolytica |
| GCF_018500205 | 392 | 389 | Klebsiella ornithinolytica |
| GCF_018966505 | 392 | 389 | Klebsiella ornithinolytica |
| GCF_019431225 | 392 | 389 | Klebsiella ornithinolytica |
| GCF_019661065 | 392 | 389 | Klebsiella ornithinolytica |
| GCF_019973635 | 392 | 389 | Klebsiella ornithinolytica |
| GCF_020116345 | 392 | 389 | Klebsiella ornithinolytica |
| GCF_020451065 | 392 | 389 | Klebsiella ornithinolytica |
| GCF_020451145 | 392 | 389 | Klebsiella ornithinolytica |
| GCF_020451165 | 392 | 389 | Klebsiella ornithinolytica |
| GCF_020451185 | 392 | 389 | Klebsiella ornithinolytica |
| GCF_020451205 | 392 | 389 | Klebsiella ornithinolytica |
| GCF_020451265 | 392 | 389 | Klebsiella ornithinolytica |
| GCF_021440775 | 392 | 378 | Klebsiella ornithinolytica |
| GCF_021440985 | 392 | 378 | Klebsiella ornithinolytica |
| GCF_021441325 | 392 | 378 | Klebsiella ornithinolytica |
| GCF_021460745 | 392 | 389 | Klebsiella ornithinolytica |
| GCF_021496525 | 392 | 389 | Klebsiella ornithinolytica |
| GCF_021698235 | 392 | 389 | Klebsiella ornithinolytica |
| GCF_021698245 | 392 | 389 | Klebsiella ornithinolytica |
| GCF_021698375 | 392 | 389 | Klebsiella ornithinolytica |
| GCF_021698415 | 392 | 389 | Klebsiella ornithinolytica |
| GCF_021698535 | 392 | 389 | Klebsiella ornithinolytica |
| GCF_021698545 | 392 | 389 | Klebsiella ornithinolytica |
| GCF_021698595 | 392 | 389 | Klebsiella ornithinolytica |
| GCF_021698635 | 392 | 389 | Klebsiella ornithinolytica |
| GCF_021698645 | 392 | 389 | Klebsiella ornithinolytica |
| GCF_021698685 | 392 | 389 | Klebsiella ornithinolytica |
| GCF_021698795 | 392 | 389 | Klebsiella ornithinolytica |
| GCF_021698805 | 392 | 389 | Klebsiella ornithinolytica |
| GCF_021699015 | 392 | 389 | Klebsiella ornithinolytica |
| GCF_022354265 | 392 | 389 | Klebsiella ornithinolytica |
| GCF_023333185 | 392 | 64 | Klebsiella ornithinolytica |
| GCF_023333185 | 392 | 389 | Klebsiella ornithinolytica |
| GCF_025146705 | 392 | 389 | Klebsiella ornithinolytica |
| GCF_025218455 | 392 | 389 | Klebsiella ornithinolytica |
| GCF_025349945 | 392 | 389 | Klebsiella ornithinolytica |
| GCF_025374775 | 392 | 389 | Klebsiella ornithinolytica |
| GCF_026223855 | 392 | 389 | Klebsiella ornithinolytica |
| GCF_026314315 | 392 | 389 | Klebsiella ornithinolytica |
| GCF_026799605 | 392 | 389 | Klebsiella ornithinolytica |
| GCF_026891915 | 392 | 389 | Klebsiella ornithinolytica |
| GCF_026891955 | 392 | 389 | Klebsiella ornithinolytica |
| GCF_027945845 | 392 | 389 | Klebsiella ornithinolytica |
| GCF_028560195 | 392 | 389 | Klebsiella ornithinolytica |
| GCF_029043495 | 392 | 64 | Klebsiella ornithinolytica |
| GCF_029043495 | 392 | 389 | Klebsiella ornithinolytica |
| GCF_029907795 | 392 | 389 | Klebsiella ornithinolytica |
| GCF_029929835 | 392 | 389 | Klebsiella ornithinolytica |
| GCF_029930115 | 392 | 389 | Klebsiella ornithinolytica |
| GCF_029931375 | 392 | 389 | Klebsiella ornithinolytica |
| GCF_029931395 | 392 | 389 | Klebsiella ornithinolytica |
| GCF_030033675 | 392 | 389 | Klebsiella ornithinolytica |
| GCF_030223405 | 392 | 389 | Klebsiella ornithinolytica |
| GCF_030223465 | 392 | 389 | Klebsiella ornithinolytica |
| GCF_030283645 | 392 | 389 | Klebsiella ornithinolytica |
| GCF_030411025 | 392 | 389 | Klebsiella ornithinolytica |
| GCF_030505655 | 392 | 389 | Klebsiella ornithinolytica |
| GCF_030717445 | 392 | 389 | Klebsiella ornithinolytica |
| GCF_030717685 | 392 | 389 | Klebsiella ornithinolytica |
| GCF_032469895 | 392 | 389 | Klebsiella ornithinolytica |
| GCF_032739325 | 392 | 389 | Klebsiella ornithinolytica |
| GCF_032739505 | 392 | 389 | Klebsiella ornithinolytica |
| GCF_032742335 | 392 | 389 | Klebsiella ornithinolytica |
| GCF_032742435 | 392 | 389 | Klebsiella ornithinolytica |
| GCF_032742555 | 392 | 389 | Klebsiella ornithinolytica |
| GCF_032747375 | 392 | 389 | Klebsiella ornithinolytica |
| GCF_033870295 | 392 | 389 | Klebsiella ornithinolytica |
| GCF_033901075 | 392 | 389 | Klebsiella ornithinolytica |
| GCF_034043555 | 392 | 389 | Klebsiella ornithinolytica |
| GCF_034043575 | 392 | 389 | Klebsiella ornithinolytica |
| GCF_035726845 | 392 | 389 | Klebsiella ornithinolytica |
| GCF_035781275 | 392 | 389 | Klebsiella ornithinolytica |
| GCF_035787015 | 392 | 389 | Klebsiella ornithinolytica |
| GCF_035787155 | 392 | 64 | Klebsiella ornithinolytica |
| GCF_035787155 | 392 | 389 | Klebsiella ornithinolytica |
| GCF_035794615 | 392 | 389 | Klebsiella ornithinolytica |
| GCF_035794715 | 392 | 389 | Klebsiella ornithinolytica |
| GCF_035794915 | 392 | 389 | Klebsiella ornithinolytica |
| GCF_035794955 | 392 | 389 | Klebsiella ornithinolytica |
| GCF_035794975 | 392 | 389 | Klebsiella ornithinolytica |
| GCF_035795215 | 392 | 389 | Klebsiella ornithinolytica |
| GCF_035795255 | 392 | 389 | Klebsiella ornithinolytica |
| GCF_035795295 | 392 | 389 | Klebsiella ornithinolytica |
| GCF_035797155 | 392 | 389 | Klebsiella ornithinolytica |
| GCF_035797455 | 392 | 389 | Klebsiella ornithinolytica |
| GCF_035966365 | 392 | 389 | Klebsiella ornithinolytica |
| GCF_035966495 | 392 | 389 | Klebsiella ornithinolytica |
| GCF_036907275 | 392 | 389 | Klebsiella ornithinolytica |
| GCF_036956365 | 392 | 389 | Klebsiella ornithinolytica |
| GCF_037052765 | 392 | 389 | Klebsiella ornithinolytica |
| GCF_037081915 | 392 | 389 | Klebsiella ornithinolytica |
| GCF_037152235 | 392 | 389 | Klebsiella ornithinolytica |
| GCF_037199105 | 392 | 389 | Klebsiella ornithinolytica |
| GCF_038380145 | 392 | 389 | Klebsiella ornithinolytica |
| GCF_039549945 | 392 | 389 | Klebsiella ornithinolytica |
| GCF_040425645 | 392 | 389 | Klebsiella ornithinolytica |
| GCF_040565875 | 392 | 389 | Klebsiella ornithinolytica |
| GCF_041725675 | 392 | 389 | Klebsiella ornithinolytica |
| GCF_041725755 | 392 | 389 | Klebsiella ornithinolytica |
| GCF_042644305 | 392 | 389 | Klebsiella ornithinolytica |
| GCF_042644455 | 392 | 389 | Klebsiella ornithinolytica |
| GCF_043102325 | 392 | 389 | Klebsiella ornithinolytica |
| GCF_043105325 | 392 | 389 | Klebsiella ornithinolytica |
| GCF_043106145 | 392 | 389 | Klebsiella ornithinolytica |
| GCF_043508675 | 392 | 389 | Klebsiella ornithinolytica |
| GCF_043509305 | 392 | 389 | Klebsiella ornithinolytica |
| GCF_043828725 | 392 | 389 | Klebsiella ornithinolytica |
| GCF_043949005 | 392 | 389 | Klebsiella ornithinolytica |
| GCF_045348735 | 392 | 389 | Klebsiella ornithinolytica |
| GCF_045674925 | 392 | 389 | Klebsiella ornithinolytica |
| GCF_045674935 | 392 | 389 | Klebsiella ornithinolytica |
| GCF_045675685 | 392 | 389 | Klebsiella ornithinolytica |
| GCF_045675695 | 392 | 389 | Klebsiella ornithinolytica |
| GCF_045690405 | 392 | 389 | Klebsiella ornithinolytica |
| GCF_045885775 | 392 | 389 | Klebsiella ornithinolytica |
| GCF_046115475 | 392 | 389 | Klebsiella ornithinolytica |
| GCF_046268845 | 392 | 389 | Klebsiella ornithinolytica |
| GCF_046365865 | 392 | 389 | Klebsiella ornithinolytica |
| GCF_046365905 | 392 | 389 | Klebsiella ornithinolytica |
| GCF_046578775 | 392 | 389 | Klebsiella ornithinolytica |
| GCF_046602315 | 392 | 389 | Klebsiella ornithinolytica |
| GCF_900083685 | 392 | 389 | Klebsiella ornithinolytica |
| GCF_900155375 | 392 | 64 | Klebsiella ornithinolytica |
| GCF_900155375 | 392 | 389 | Klebsiella ornithinolytica |
| GCF_900184005 | 392 | 389 | Klebsiella ornithinolytica |
| GCF_900635915 | 137 | 389 | Klebsiella ornithinolytica |
| GCF_900635915 | 248 | 389 | Klebsiella ornithinolytica |
| GCF_901420905 | 392 | 389 | Klebsiella ornithinolytica |
| GCF_901421005 | 392 | 389 | Klebsiella ornithinolytica |
| GCF_902161225 | 392 | 389 | Klebsiella ornithinolytica |
| GCF_902163305 | 392 | 389 | Klebsiella ornithinolytica |
| GCF_902386325 | 392 | 389 | Klebsiella ornithinolytica |
| GCF_905219255 | 392 | 389 | Klebsiella ornithinolytica |
| GCF_947047355 | 392 | 389 | Klebsiella ornithinolytica |
| GCF_963887305 | 392 | 389 | Klebsiella ornithinolytica |
| GFKo17 | 392 | 389 | Klebsiella ornithinolytica |
| GFKo24 | 392 | 389 | Klebsiella ornithinolytica |
| GFKo25 | 392 | 389 | Klebsiella ornithinolytica |
| GFKo4 | 392 | 389 | Klebsiella ornithinolytica |
| GFKo43 | 392 | 389 | Klebsiella ornithinolytica |
| GFKo9 | 392 | 389 | Klebsiella ornithinolytica |
| SAMD00002693 | 392 | 389 | Klebsiella ornithinolytica |
| SAMD00092916 | 392 | 389 | Klebsiella ornithinolytica |
| SAMD00194387 | 392 | 389 | Klebsiella ornithinolytica |
| SAMD00194388 | 392 | 389 | Klebsiella ornithinolytica |
| SAMD00262490 | 392 | 389 | Klebsiella ornithinolytica |
| SAMD00320838 | 392 | 389 | Klebsiella ornithinolytica |
| SAMD00320921 | 392 | 389 | Klebsiella ornithinolytica |
| SAMD00492892 | N | 389 | Klebsiella ornithinolytica |
| SAMD00492898 | 392 | 389 | Klebsiella ornithinolytica |
| SAMD00492899 | N | 389 | Klebsiella ornithinolytica |
| SAMD00492900 | 392 | 389 | Klebsiella ornithinolytica |
| SAMD00492909 | 392 | 389 | Klebsiella ornithinolytica |
| SAMD00492912 | 392 | 389 | Klebsiella ornithinolytica |
| SAMD00492913 | 392 | 389 | Klebsiella ornithinolytica |
| SAMD00492915 | 392 | 389 | Klebsiella ornithinolytica |
| SAMD00492916 | 392 | 389 | Klebsiella ornithinolytica |
| SAMD00492917 | 392 | 389 | Klebsiella ornithinolytica |
| SAMD00492918 | 392 | 389 | Klebsiella ornithinolytica |
| SAMD00492922 | 392 | 389 | Klebsiella ornithinolytica |
| SAMD00492923 | N | 389 | Klebsiella ornithinolytica |
| SAMD00492926 | N | 389 | Klebsiella ornithinolytica |
| SAMD00492927 | 392 | 389 | Klebsiella ornithinolytica |
| SAMD00492934 | 392 | 389 | Klebsiella ornithinolytica |
| SAMD00492935 | N | 389 | Klebsiella ornithinolytica |
| SAMD00492936 | 392 | 389 | Klebsiella ornithinolytica |
| SAMD00492937 | 392 | 389 | Klebsiella ornithinolytica |
| SAMD00492940 | N | 389 | Klebsiella ornithinolytica |
| SAMD00498302 | 392 | 389 | Klebsiella ornithinolytica |
| SAMD00498736 | 392 | 389 | Klebsiella ornithinolytica |
| SAMD00498885 | 392 | 389 | Klebsiella ornithinolytica |
| SAMD00498996 | 392 | 389 | Klebsiella ornithinolytica |
| SAMD00499993 | 392 | 389 | Klebsiella ornithinolytica |
| SAMD00500506 | 392 | 389 | Klebsiella ornithinolytica |
| SAMD00501192 | 392 | 389 | Klebsiella ornithinolytica |
| SAMD00501195 | 392 | 389 | Klebsiella ornithinolytica |
| SAMD00553967 | 392 | 389 | Klebsiella ornithinolytica |
| SAMD00553996 | 392 | 389 | Klebsiella ornithinolytica |
| SAMD00554007 | 392 | 389 | Klebsiella ornithinolytica |
| SAMD00554015 | 392 | 389 | Klebsiella ornithinolytica |
| SAMD00554021 | 392 | 389 | Klebsiella ornithinolytica |
| SAMD00554027 | 392 | 389 | Klebsiella ornithinolytica |
| SAMD00554034 | 392 | 389 | Klebsiella ornithinolytica |
| SAMEA10303445 | 392 | 389 | Klebsiella ornithinolytica |
| SAMEA104032373 | 392 | 88 | Klebsiella ornithinolytica |
| SAMEA104032373 | 392 | 389 | Klebsiella ornithinolytica |
| SAMEA104032378 | 392 | 389 | Klebsiella ornithinolytica |
| SAMEA104032389 | 392 | 389 | Klebsiella ornithinolytica |
| SAMEA104457996 | 392 | 389 | Klebsiella ornithinolytica |
| SAMEA10468700 | 392 | 389 | Klebsiella ornithinolytica |
| SAMEA110044371 | 392 | 389 | Klebsiella ornithinolytica |
| SAMEA110063381 | 392 | 389 | Klebsiella ornithinolytica |
| SAMEA110063386 | 392 | 389 | Klebsiella ornithinolytica |
| SAMEA110063405 | 392 | 64 | Klebsiella ornithinolytica |
| SAMEA110063405 | 392 | 389 | Klebsiella ornithinolytica |
| SAMEA111347218 | 392 | 389 | Klebsiella ornithinolytica |
| SAMEA111441541 | 392 | 389 | Klebsiella ornithinolytica |
| SAMEA111441543 | 392 | 64 | Klebsiella ornithinolytica |
| SAMEA111441543 | 392 | 389 | Klebsiella ornithinolytica |
| SAMEA111441545 | 392 | 389 | Klebsiella ornithinolytica |
| SAMEA111441555 | 392 | 389 | Klebsiella ornithinolytica |
| SAMEA111441558 | 392 | 389 | Klebsiella ornithinolytica |
| SAMEA111441585 | 392 | 389 | Klebsiella ornithinolytica |
| SAMEA111441586 | 392 | 389 | Klebsiella ornithinolytica |
| SAMEA111441588 | 392 | 389 | Klebsiella ornithinolytica |
| SAMEA111441589 | 392 | 389 | Klebsiella ornithinolytica |
| SAMEA111441590 | 392 | 389 | Klebsiella ornithinolytica |
| SAMEA111441592 | 392 | 389 | Klebsiella ornithinolytica |
| SAMEA111441599 | 392 | 389 | Klebsiella ornithinolytica |
| SAMEA111441608 | 392 | 389 | Klebsiella ornithinolytica |
| SAMEA111503395 | 392 | 389 | Klebsiella ornithinolytica |
| SAMEA111503548 | 392 | 389 | Klebsiella ornithinolytica |
| SAMEA111503905 | 392 | 389 | Klebsiella ornithinolytica |
| SAMEA111504558 | 392 | 389 | Klebsiella ornithinolytica |
| SAMEA111504559 | 392 | 389 | Klebsiella ornithinolytica |
| SAMEA111506337 | 392 | 389 | Klebsiella ornithinolytica |
| SAMEA111506418 | 392 | 389 | Klebsiella ornithinolytica |
| SAMEA112192757 | 392 | 389 | Klebsiella ornithinolytica |
| SAMEA112192759 | 392 | 389 | Klebsiella ornithinolytica |
| SAMEA112192760 | 392 | 389 | Klebsiella ornithinolytica |
| SAMEA112192761 | 392 | 389 | Klebsiella ornithinolytica |
| SAMEA112192762 | 392 | 389 | Klebsiella ornithinolytica |
| SAMEA112192763 | 392 | 389 | Klebsiella ornithinolytica |
| SAMEA11291311 | 392 | 389 | Klebsiella ornithinolytica |
| SAMEA12944378 | 392 | 389 | Klebsiella ornithinolytica |
| SAMEA2046730 | 392 | 389 | Klebsiella ornithinolytica |
| SAMEA3357537 | 392 | 389 | Klebsiella ornithinolytica |
| SAMEA3357538 | 392 | 389 | Klebsiella ornithinolytica |
| SAMEA3357539 | 392 | 389 | Klebsiella ornithinolytica |
| SAMEA3357540 | 392 | 389 | Klebsiella ornithinolytica |
| SAMEA3538634 | 392 | 389 | Klebsiella ornithinolytica |
| SAMEA3726392 | 392 | 389 | Klebsiella ornithinolytica |
| SAMEA47366668 | 392 | 389 | Klebsiella ornithinolytica |
| SAMEA4781101 | 392 | 389 | Klebsiella ornithinolytica |
| SAMEA4781452 | 392 | 389 | Klebsiella ornithinolytica |
| SAMEA4781473 | 392 | 389 | Klebsiella ornithinolytica |
| SAMEA4781952 | 392 | 389 | Klebsiella ornithinolytica |
| SAMEA4781959 | 392 | 389 | Klebsiella ornithinolytica |
| SAMEA4781981 | 392 | 389 | Klebsiella ornithinolytica |
| SAMEA4781994 | 392 | 389 | Klebsiella ornithinolytica |
| SAMEA4781998 | 392 | 389 | Klebsiella ornithinolytica |
| SAMEA4781999 | 392 | 389 | Klebsiella ornithinolytica |
| SAMEA4782000 | 392 | 389 | Klebsiella ornithinolytica |
| SAMEA4782001 | 392 | 389 | Klebsiella ornithinolytica |
| SAMEA4782012 | 392 | 389 | Klebsiella ornithinolytica |
| SAMEA4782014 | 392 | 389 | Klebsiella ornithinolytica |
| SAMEA4782016 | 392 | 389 | Klebsiella ornithinolytica |
| SAMEA4782017 | 392 | 389 | Klebsiella ornithinolytica |
| SAMEA4782021 | 392 | 389 | Klebsiella ornithinolytica |
| SAMEA4782024 | 392 | 389 | Klebsiella ornithinolytica |
| SAMEA4801620 | 392 | 389 | Klebsiella ornithinolytica |
| SAMEA5048722 | 392 | 389 | Klebsiella ornithinolytica |
| SAMEA5048723 | 392 | 389 | Klebsiella ornithinolytica |
| SAMEA5048750 | 392 | 389 | Klebsiella ornithinolytica |
| SAMEA5048751 | 392 | 389 | Klebsiella ornithinolytica |
| SAMEA5048768 | 392 | 389 | Klebsiella ornithinolytica |
| SAMEA5048769 | 392 | 389 | Klebsiella ornithinolytica |
| SAMEA5048784 | 392 | 389 | Klebsiella ornithinolytica |
| SAMEA5048787 | 392 | 389 | Klebsiella ornithinolytica |
| SAMEA5048788 | 392 | 389 | Klebsiella ornithinolytica |
| SAMEA5048794 | 392 | 389 | Klebsiella ornithinolytica |
| SAMEA5048795 | 392 | 389 | Klebsiella ornithinolytica |
| SAMEA5048797 | 392 | 389 | Klebsiella ornithinolytica |
| SAMEA5048798 | 392 | 389 | Klebsiella ornithinolytica |
| SAMEA5048802 | 392 | 389 | Klebsiella ornithinolytica |
| SAMEA5048803 | 392 | 389 | Klebsiella ornithinolytica |
| SAMEA5048811 | 392 | 389 | Klebsiella ornithinolytica |
| SAMEA5048819 | 392 | 389 | Klebsiella ornithinolytica |
| SAMEA5048822 | 392 | 389 | Klebsiella ornithinolytica |
| SAMEA5048831 | 392 | 389 | Klebsiella ornithinolytica |
| SAMEA5048832 | 392 | 389 | Klebsiella ornithinolytica |
| SAMEA5048847 | 392 | 389 | Klebsiella ornithinolytica |
| SAMEA5048849 | 392 | 389 | Klebsiella ornithinolytica |
| SAMEA5048865 | 392 | 389 | Klebsiella ornithinolytica |
| SAMEA5048870 | 392 | 389 | Klebsiella ornithinolytica |
| SAMEA5048873 | 392 | 389 | Klebsiella ornithinolytica |
| SAMEA5048903 | 392 | 389 | Klebsiella ornithinolytica |
| SAMEA5048905 | 392 | 389 | Klebsiella ornithinolytica |
| SAMEA5048920 | 392 | 389 | Klebsiella ornithinolytica |
| SAMEA5048921 | 392 | 389 | Klebsiella ornithinolytica |
| SAMEA5048925 | 392 | 389 | Klebsiella ornithinolytica |
| SAMEA5048928 | 392 | 389 | Klebsiella ornithinolytica |
| SAMEA5048938 | 392 | 389 | Klebsiella ornithinolytica |
| SAMEA5048942 | 392 | 389 | Klebsiella ornithinolytica |
| SAMEA5049003 | 392 | 389 | Klebsiella ornithinolytica |
| SAMEA5049063 | 392 | 64 | Klebsiella ornithinolytica |
| SAMEA5049063 | 392 | 389 | Klebsiella ornithinolytica |
| SAMEA5049064 | 392 | 64 | Klebsiella ornithinolytica |
| SAMEA5049064 | 392 | 389 | Klebsiella ornithinolytica |
| SAMEA5049066 | 392 | 64 | Klebsiella ornithinolytica |
| SAMEA5049066 | 392 | 389 | Klebsiella ornithinolytica |
| SAMEA5049069 | 392 | 389 | Klebsiella ornithinolytica |
| SAMEA5049079 | 392 | 389 | Klebsiella ornithinolytica |
| SAMEA5049082 | 392 | 389 | Klebsiella ornithinolytica |
| SAMEA5049115 | 392 | 389 | Klebsiella ornithinolytica |
| SAMEA5049124 | 392 | 389 | Klebsiella ornithinolytica |
| SAMEA5049149 | 392 | 389 | Klebsiella ornithinolytica |
| SAMEA5049150 | 392 | 389 | Klebsiella ornithinolytica |
| SAMEA5049155 | 392 | 389 | Klebsiella ornithinolytica |
| SAMEA5049161 | 392 | 389 | Klebsiella ornithinolytica |
| SAMEA5049166 | 392 | 389 | Klebsiella ornithinolytica |
| SAMEA5049167 | 392 | 389 | Klebsiella ornithinolytica |
| SAMEA5049176 | 392 | 389 | Klebsiella ornithinolytica |
| SAMEA5049190 | 392 | 389 | Klebsiella ornithinolytica |
| SAMEA5049193 | 392 | 389 | Klebsiella ornithinolytica |
| SAMEA5049203 | 392 | 389 | Klebsiella ornithinolytica |
| SAMEA5049205 | 392 | 389 | Klebsiella ornithinolytica |
| SAMEA5049206 | 392 | 389 | Klebsiella ornithinolytica |
| SAMEA5049225 | 392 | 389 | Klebsiella ornithinolytica |
| SAMEA5049226 | 392 | 389 | Klebsiella ornithinolytica |
| SAMEA5049248 | 392 | 389 | Klebsiella ornithinolytica |
| SAMEA5049253 | 392 | 389 | Klebsiella ornithinolytica |
| SAMEA5049269 | 392 | 389 | Klebsiella ornithinolytica |
| SAMEA5049274 | 392 | 389 | Klebsiella ornithinolytica |
| SAMEA5049278 | 392 | 389 | Klebsiella ornithinolytica |
| SAMEA5049286 | 392 | 389 | Klebsiella ornithinolytica |
| SAMEA5049294 | 392 | 389 | Klebsiella ornithinolytica |
| SAMEA5049307 | 392 | 389 | Klebsiella ornithinolytica |
| SAMEA5049310 | 392 | 389 | Klebsiella ornithinolytica |
| SAMEA5049312 | 392 | 389 | Klebsiella ornithinolytica |
| SAMEA5049315 | 392 | 389 | Klebsiella ornithinolytica |
| SAMEA5049317 | 392 | 389 | Klebsiella ornithinolytica |
| SAMEA5049321 | 392 | 389 | Klebsiella ornithinolytica |
| SAMEA5049327 | 392 | 389 | Klebsiella ornithinolytica |
| SAMEA5049328 | 392 | 389 | Klebsiella ornithinolytica |
| SAMEA5049351 | 392 | 389 | Klebsiella ornithinolytica |
| SAMEA5049388 | 392 | 389 | Klebsiella ornithinolytica |
| SAMEA5049390 | 392 | 389 | Klebsiella ornithinolytica |
| SAMEA5049391 | 392 | 389 | Klebsiella ornithinolytica |
| SAMEA5049419 | 392 | 389 | Klebsiella ornithinolytica |
| SAMEA5049457 | 392 | 389 | Klebsiella ornithinolytica |
| SAMEA5049514 | 392 | 389 | Klebsiella ornithinolytica |
| SAMEA5049518 | 392 | 389 | Klebsiella ornithinolytica |
| SAMEA5049561 | 392 | 389 | Klebsiella ornithinolytica |
| SAMEA5049564 | 392 | 389 | Klebsiella ornithinolytica |
| SAMEA5049566 | 392 | 389 | Klebsiella ornithinolytica |
| SAMEA5049568 | 392 | 389 | Klebsiella ornithinolytica |
| SAMEA5049572 | 392 | 389 | Klebsiella ornithinolytica |
| SAMEA5049577 | 392 | 389 | Klebsiella ornithinolytica |
| SAMEA5049583 | 392 | 389 | Klebsiella ornithinolytica |
| SAMEA5049585 | 392 | 389 | Klebsiella ornithinolytica |
| SAMEA5049588 | 392 | 389 | Klebsiella ornithinolytica |
| SAMEA5049632 | 392 | 389 | Klebsiella ornithinolytica |
| SAMEA5049634 | 392 | 389 | Klebsiella ornithinolytica |
| SAMEA5049642 | 392 | 389 | Klebsiella ornithinolytica |
| SAMEA5049644 | 392 | 389 | Klebsiella ornithinolytica |
| SAMEA5049646 | 392 | 389 | Klebsiella ornithinolytica |
| SAMEA5049650 | 392 | 389 | Klebsiella ornithinolytica |
| SAMEA5049653 | 392 | 389 | Klebsiella ornithinolytica |
| SAMEA5049673 | 392 | 389 | Klebsiella ornithinolytica |
| SAMEA5049686 | 392 | 389 | Klebsiella ornithinolytica |
| SAMEA5049741 | 392 | 389 | Klebsiella ornithinolytica |
| SAMEA5049744 | 392 | 389 | Klebsiella ornithinolytica |
| SAMEA5049763 | 392 | 389 | Klebsiella ornithinolytica |
| SAMEA5049767 | 392 | 389 | Klebsiella ornithinolytica |
| SAMEA5049777 | 392 | 389 | Klebsiella ornithinolytica |
| SAMEA5049783 | 392 | 389 | Klebsiella ornithinolytica |
| SAMEA5049790 | 392 | 64 | Klebsiella ornithinolytica |
| SAMEA5049790 | 392 | 389 | Klebsiella ornithinolytica |
| SAMEA5049794 | 392 | 389 | Klebsiella ornithinolytica |
| SAMEA5049795 | 392 | 389 | Klebsiella ornithinolytica |
| SAMEA5049825 | 392 | 389 | Klebsiella ornithinolytica |
| SAMEA5049850 | 392 | 389 | Klebsiella ornithinolytica |
| SAMEA5049868 | 392 | 389 | Klebsiella ornithinolytica |
| SAMEA5049873 | 392 | 389 | Klebsiella ornithinolytica |
| SAMEA5049885 | 392 | 389 | Klebsiella ornithinolytica |
| SAMEA5049927 | 392 | 389 | Klebsiella ornithinolytica |
| SAMEA5049930 | 392 | 389 | Klebsiella ornithinolytica |
| SAMEA5049946 | 392 | 389 | Klebsiella ornithinolytica |
| SAMEA5050009 | 392 | 389 | Klebsiella ornithinolytica |
| SAMEA5050010 | 392 | 389 | Klebsiella ornithinolytica |
| SAMEA5050020 | 392 | 389 | Klebsiella ornithinolytica |
| SAMEA5050022 | 392 | 389 | Klebsiella ornithinolytica |
| SAMEA5050036 | 392 | 389 | Klebsiella ornithinolytica |
| SAMEA5050068 | 392 | 389 | Klebsiella ornithinolytica |
| SAMEA5050084 | 392 | 389 | Klebsiella ornithinolytica |
| SAMEA5050095 | 392 | 389 | Klebsiella ornithinolytica |
| SAMEA5050098 | 392 | 389 | Klebsiella ornithinolytica |
| SAMEA5050117 | 392 | 389 | Klebsiella ornithinolytica |
| SAMEA5050150 | 392 | 389 | Klebsiella ornithinolytica |
| SAMEA5050158 | 392 | 389 | Klebsiella ornithinolytica |
| SAMEA5050163 | 392 | 389 | Klebsiella ornithinolytica |
| SAMEA5050169 | 392 | 389 | Klebsiella ornithinolytica |
| SAMEA5050173 | 392 | 389 | Klebsiella ornithinolytica |
| SAMEA5050176 | 392 | 389 | Klebsiella ornithinolytica |
| SAMEA5050177 | 392 | 389 | Klebsiella ornithinolytica |
| SAMEA5050226 | 392 | 389 | Klebsiella ornithinolytica |
| SAMEA5050227 | 392 | 389 | Klebsiella ornithinolytica |
| SAMEA5050228 | 392 | 389 | Klebsiella ornithinolytica |
| SAMEA5050231 | 392 | 389 | Klebsiella ornithinolytica |
| SAMEA5050232 | 392 | 389 | Klebsiella ornithinolytica |
| SAMEA5050269 | 392 | 389 | Klebsiella ornithinolytica |
| SAMEA5050270 | 392 | 389 | Klebsiella ornithinolytica |
| SAMEA5050284 | 392 | 389 | Klebsiella ornithinolytica |
| SAMEA5050289 | 392 | 389 | Klebsiella ornithinolytica |
| SAMEA5050290 | 392 | 130 | Klebsiella ornithinolytica |
| SAMEA5050290 | 392 | 244 | Klebsiella ornithinolytica |
| SAMEA5050294 | 392 | 389 | Klebsiella ornithinolytica |
| SAMEA5050303 | 392 | 389 | Klebsiella ornithinolytica |
| SAMEA5050307 | 392 | 389 | Klebsiella ornithinolytica |
| SAMEA5050310 | 392 | 389 | Klebsiella ornithinolytica |
| SAMEA5050311 | 392 | 389 | Klebsiella ornithinolytica |
| SAMEA5050312 | 392 | 389 | Klebsiella ornithinolytica |
| SAMEA5612529 | 392 | 389 | Klebsiella ornithinolytica |
| SAMEA5684234 | 392 | 389 | Klebsiella ornithinolytica |
| SAMEA5684246 | 392 | 389 | Klebsiella ornithinolytica |
| SAMEA5684262 | 392 | 389 | Klebsiella ornithinolytica |
| SAMEA5684270 | 392 | 389 | Klebsiella ornithinolytica |
| SAMEA5684283 | 392 | 389 | Klebsiella ornithinolytica |
| SAMEA5684301 | 392 | 389 | Klebsiella ornithinolytica |
| SAMEA5684315 | 392 | 389 | Klebsiella ornithinolytica |
| SAMEA5684320 | 392 | 389 | Klebsiella ornithinolytica |
| SAMEA5684322 | 392 | 389 | Klebsiella ornithinolytica |
| SAMEA5751256 | 392 | 389 | Klebsiella ornithinolytica |
| SAMEA5751266 | 392 | 389 | Klebsiella ornithinolytica |
| SAMEA5751386 | 392 | 389 | Klebsiella ornithinolytica |
| SAMEA5996249 | 392 | 389 | Klebsiella ornithinolytica |
| SAMEA5996259 | 392 | 389 | Klebsiella ornithinolytica |
| SAMEA5996260 | 392 | 389 | Klebsiella ornithinolytica |
| SAMEA5996263 | 392 | 389 | Klebsiella ornithinolytica |
| SAMEA5996266 | 392 | 389 | Klebsiella ornithinolytica |
| SAMEA5996268 | 392 | 389 | Klebsiella ornithinolytica |
| SAMEA5996269 | 392 | 389 | Klebsiella ornithinolytica |
| SAMEA5996270 | 392 | 389 | Klebsiella ornithinolytica |
| SAMEA5996273 | 392 | 389 | Klebsiella ornithinolytica |
| SAMEA5996278 | 392 | 389 | Klebsiella ornithinolytica |
| SAMEA5996279 | 392 | 389 | Klebsiella ornithinolytica |
| SAMEA5996285 | 392 | 389 | Klebsiella ornithinolytica |
| SAMEA5996287 | 392 | 389 | Klebsiella ornithinolytica |
| SAMEA5996289 | 392 | 389 | Klebsiella ornithinolytica |
| SAMEA5996290 | 392 | 389 | Klebsiella ornithinolytica |
| SAMEA5996295 | 392 | 389 | Klebsiella ornithinolytica |
| SAMEA5996309 | 392 | 389 | Klebsiella ornithinolytica |
| SAMEA5996322 | 392 | 389 | Klebsiella ornithinolytica |
| SAMEA5996323 | 392 | 389 | Klebsiella ornithinolytica |
| SAMEA5996326 | 392 | 389 | Klebsiella ornithinolytica |
| SAMEA5996328 | 392 | 389 | Klebsiella ornithinolytica |
| SAMEA5996329 | 392 | 389 | Klebsiella ornithinolytica |
| SAMEA5996330 | 392 | 389 | Klebsiella ornithinolytica |
| SAMEA6099698 | 392 | 389 | Klebsiella ornithinolytica |
| SAMEA6099699 | 392 | 389 | Klebsiella ornithinolytica |
| SAMEA6099700 | 392 | 389 | Klebsiella ornithinolytica |
| SAMEA6099701 | 392 | 389 | Klebsiella ornithinolytica |
| SAMEA6107543 | 392 | 389 | Klebsiella ornithinolytica |
| SAMEA6107670 | 392 | 389 | Klebsiella ornithinolytica |
| SAMEA6107680 | 392 | 389 | Klebsiella ornithinolytica |
| SAMEA6107687 | 392 | 389 | Klebsiella ornithinolytica |
| SAMEA6107689 | 392 | 389 | Klebsiella ornithinolytica |
| SAMEA6368797 | 392 | 389 | Klebsiella ornithinolytica |
| SAMEA6451122 | 392 | 389 | Klebsiella ornithinolytica |
| SAMEA6656904 | 392 | 389 | Klebsiella ornithinolytica |
| SAMEA6657607 | 392 | 389 | Klebsiella ornithinolytica |
| SAMEA7198809 | 392 | 389 | Klebsiella ornithinolytica |
| SAMEA7198810 | 392 | 389 | Klebsiella ornithinolytica |
| SAMEA7198811 | 392 | 389 | Klebsiella ornithinolytica |
| SAMEA7198812 | 392 | 389 | Klebsiella ornithinolytica |
| SAMEA7198814 | 392 | 389 | Klebsiella ornithinolytica |
| SAMEA7198816 | 392 | 389 | Klebsiella ornithinolytica |
| SAMEA7198817 | 392 | 389 | Klebsiella ornithinolytica |
| SAMEA7198818 | 392 | 389 | Klebsiella ornithinolytica |
| SAMEA7198825 | 392 | 389 | Klebsiella ornithinolytica |
| SAMEA7198826 | 392 | 389 | Klebsiella ornithinolytica |
| SAMEA7198827 | 392 | 389 | Klebsiella ornithinolytica |
| SAMEA7198830 | 392 | 389 | Klebsiella ornithinolytica |
| SAMEA7198831 | 392 | 389 | Klebsiella ornithinolytica |
| SAMEA7198832 | 392 | 389 | Klebsiella ornithinolytica |
| SAMEA7198839 | 392 | 389 | Klebsiella ornithinolytica |
| SAMEA7198840 | 392 | 389 | Klebsiella ornithinolytica |
| SAMEA7198841 | 392 | 389 | Klebsiella ornithinolytica |
| SAMEA7198842 | 392 | 389 | Klebsiella ornithinolytica |
| SAMEA7198847 | 392 | 389 | Klebsiella ornithinolytica |
| SAMEA7198858 | 392 | 389 | Klebsiella ornithinolytica |
| SAMEA7198859 | 392 | 389 | Klebsiella ornithinolytica |
| SAMEA7198867 | 392 | 389 | Klebsiella ornithinolytica |
| SAMEA7198868 | 392 | 389 | Klebsiella ornithinolytica |
| SAMEA7456868 | 392 | 389 | Klebsiella ornithinolytica |
| SAMEA8755900 | 392 | 389 | Klebsiella ornithinolytica |
| SAMN00672462 | 392 | 389 | Klebsiella ornithinolytica |
| SAMN02680230 | 392 | 389 | Klebsiella ornithinolytica |
| SAMN03733691 | 392 | 389 | Klebsiella ornithinolytica |
| SAMN04014975 | 392 | 389 | Klebsiella ornithinolytica |
| SAMN07692515 | 392 | 389 | Klebsiella ornithinolytica |
| SAMN08932888 | 392 | 389 | Klebsiella ornithinolytica |
| SAMN08932889 | 392 | 389 | Klebsiella ornithinolytica |
| SAMN08932890 | 392 | 389 | Klebsiella ornithinolytica |
| SAMN09210746 | 392 | 389 | Klebsiella ornithinolytica |
| SAMN09210750 | 392 | 389 | Klebsiella ornithinolytica |
| SAMN09460063 | 392 | 389 | Klebsiella ornithinolytica |
| SAMN10343276 | 392 | 389 | Klebsiella ornithinolytica |
| SAMN10343277 | 392 | 389 | Klebsiella ornithinolytica |
| SAMN10375007 | 392 | 389 | Klebsiella ornithinolytica |
| SAMN11164559 | 392 | 389 | Klebsiella ornithinolytica |
| SAMN11230993 | 392 | 389 | Klebsiella ornithinolytica |
| SAMN12250677 | 392 | 389 | Klebsiella ornithinolytica |
| SAMN13951920 | 392 | 389 | Klebsiella ornithinolytica |
| SAMN14150169 | 392 | 389 | Klebsiella ornithinolytica |
| SAMN14640330 | 392 | 389 | Klebsiella ornithinolytica |
| SAMN15566985 | 392 | 389 | Klebsiella ornithinolytica |
| SAMN15566986 | 392 | 389 | Klebsiella ornithinolytica |
| SAMN15566987 | 392 | 389 | Klebsiella ornithinolytica |
| SAMN15566988 | 392 | 389 | Klebsiella ornithinolytica |
| SAMN15566989 | 392 | 389 | Klebsiella ornithinolytica |
| SAMN15566990 | 392 | 389 | Klebsiella ornithinolytica |
| SAMN15566996 | 392 | 389 | Klebsiella ornithinolytica |
| SAMN15566997 | 392 | 389 | Klebsiella ornithinolytica |
| SAMN15868781 | 392 | 389 | Klebsiella ornithinolytica |
| SAMN15868863 | 392 | 389 | Klebsiella ornithinolytica |
| SAMN15868895 | 392 | 389 | Klebsiella ornithinolytica |
| SAMN15869077 | 392 | 389 | Klebsiella ornithinolytica |
| SAMN16233152 | 392 | 389 | Klebsiella ornithinolytica |
| SAMN16233158 | 392 | 389 | Klebsiella ornithinolytica |
| SAMN16233194 | 392 | 389 | Klebsiella ornithinolytica |
| SAMN16233230 | 392 | 389 | Klebsiella ornithinolytica |
| SAMN16233239 | 392 | 389 | Klebsiella ornithinolytica |
| SAMN16233240 | 392 | 389 | Klebsiella ornithinolytica |
| SAMN16233310 | 392 | 389 | Klebsiella ornithinolytica |
| SAMN16233312 | 392 | 389 | Klebsiella ornithinolytica |
| SAMN16278201 | 392 | 389 | Klebsiella ornithinolytica |
| SAMN16278268 | 392 | 389 | Klebsiella ornithinolytica |
| SAMN16824625 | 392 | 389 | Klebsiella ornithinolytica |
| SAMN19595661 | 392 | 389 | Klebsiella ornithinolytica |
| SAMN19856777 | 392 | 389 | Klebsiella ornithinolytica |
| SAMN19856778 | 392 | 389 | Klebsiella ornithinolytica |
| SAMN19856779 | 392 | 389 | Klebsiella ornithinolytica |
| SAMN19856781 | 392 | 389 | Klebsiella ornithinolytica |
| SAMN19856782 | 392 | 389 | Klebsiella ornithinolytica |
| SAMN19856783 | 392 | 389 | Klebsiella ornithinolytica |
| SAMN19856784 | 392 | 389 | Klebsiella ornithinolytica |
| SAMN19856785 | 392 | 389 | Klebsiella ornithinolytica |
| SAMN19856786 | 392 | 389 | Klebsiella ornithinolytica |
| SAMN19856787 | 392 | 389 | Klebsiella ornithinolytica |
| SAMN19856788 | 392 | 389 | Klebsiella ornithinolytica |
| SAMN19856789 | 392 | 389 | Klebsiella ornithinolytica |
| SAMN19856790 | 392 | 389 | Klebsiella ornithinolytica |
| SAMN19856791 | 392 | 389 | Klebsiella ornithinolytica |
| SAMN19856792 | 392 | 389 | Klebsiella ornithinolytica |
| SAMN19856793 | 392 | 389 | Klebsiella ornithinolytica |
| SAMN20152571 | 392 | 64 | Klebsiella ornithinolytica |
| SAMN20152571 | 392 | 389 | Klebsiella ornithinolytica |
| SAMN20396754 | 392 | 389 | Klebsiella ornithinolytica |
| SAMN22677368 | 392 | 389 | Klebsiella ornithinolytica |
| SAMN22837213 | 392 | 389 | Klebsiella ornithinolytica |
| SAMN22837220 | 392 | 389 | Klebsiella ornithinolytica |
| SAMN22875454 | 392 | 389 | Klebsiella ornithinolytica |
| SAMN22875455 | 392 | 389 | Klebsiella ornithinolytica |
| SAMN22875457 | 392 | 389 | Klebsiella ornithinolytica |
| SAMN22875460 | 392 | 389 | Klebsiella ornithinolytica |
| SAMN22875461 | 392 | 389 | Klebsiella ornithinolytica |
| SAMN22875462 | 392 | 389 | Klebsiella ornithinolytica |
| SAMN22875463 | 392 | 389 | Klebsiella ornithinolytica |
| SAMN22875465 | 392 | 389 | Klebsiella ornithinolytica |
| SAMN22875467 | 392 | 389 | Klebsiella ornithinolytica |
| SAMN22875469 | 392 | 389 | Klebsiella ornithinolytica |
| SAMN22875474 | 392 | 389 | Klebsiella ornithinolytica |
| SAMN22875475 | 392 | 389 | Klebsiella ornithinolytica |
| SAMN22875477 | 392 | 389 | Klebsiella ornithinolytica |
| SAMN22875479 | 392 | 389 | Klebsiella ornithinolytica |
| SAMN22875480 | 392 | 389 | Klebsiella ornithinolytica |
| SAMN22875488 | 392 | 389 | Klebsiella ornithinolytica |
| SAMN22875489 | 392 | 389 | Klebsiella ornithinolytica |
| SAMN22875490 | 392 | 389 | Klebsiella ornithinolytica |
| SAMN22875492 | 392 | 389 | Klebsiella ornithinolytica |
| SAMN22875496 | 392 | 389 | Klebsiella ornithinolytica |
| SAMN22875499 | 392 | 389 | Klebsiella ornithinolytica |
| SAMN22875500 | 392 | 389 | Klebsiella ornithinolytica |
| SAMN22875501 | 392 | 389 | Klebsiella ornithinolytica |
| SAMN22875503 | 392 | 389 | Klebsiella ornithinolytica |
| SAMN22875504 | 392 | 389 | Klebsiella ornithinolytica |
| SAMN22875508 | 392 | 389 | Klebsiella ornithinolytica |
| SAMN22875511 | 392 | 389 | Klebsiella ornithinolytica |
| SAMN22875514 | 392 | 389 | Klebsiella ornithinolytica |
| SAMN22875516 | 392 | 389 | Klebsiella ornithinolytica |
| SAMN22875520 | 392 | 389 | Klebsiella ornithinolytica |
| SAMN22875521 | 392 | 389 | Klebsiella ornithinolytica |
| SAMN22875523 | 392 | 389 | Klebsiella ornithinolytica |
| SAMN22875527 | 392 | 389 | Klebsiella ornithinolytica |
| SAMN22875529 | 392 | 389 | Klebsiella ornithinolytica |
| SAMN22875530 | 392 | 389 | Klebsiella ornithinolytica |
| SAMN22875531 | 392 | 389 | Klebsiella ornithinolytica |
| SAMN22968076 | 392 | 64 | Klebsiella ornithinolytica |
| SAMN22968076 | 392 | 389 | Klebsiella ornithinolytica |
| SAMN23768814 | 392 | 389 | Klebsiella ornithinolytica |
| SAMN24018969 | 392 | 389 | Klebsiella ornithinolytica |
| SAMN24019066 | 392 | 389 | Klebsiella ornithinolytica |
| SAMN24019228 | 392 | 389 | Klebsiella ornithinolytica |
| SAMN24020802 | 392 | 389 | Klebsiella ornithinolytica |
| SAMN24251563 | 392 | 64 | Klebsiella ornithinolytica |
| SAMN24251563 | 392 | 389 | Klebsiella ornithinolytica |
| SAMN24296925 | 392 | 389 | Klebsiella ornithinolytica |
| SAMN24297207 | 392 | 389 | Klebsiella ornithinolytica |
| SAMN25860413 | 392 | 389 | Klebsiella ornithinolytica |
| SAMN25860416 | 392 | 389 | Klebsiella ornithinolytica |
| SAMN25989382 | 392 | 389 | Klebsiella ornithinolytica |
| SAMN25989395 | 392 | 306 | Klebsiella ornithinolytica |
| SAMN25989671 | 392 | 389 | Klebsiella ornithinolytica |
| SAMN25989723 | 392 | 306 | Klebsiella ornithinolytica |
| SAMN25989788 | 392 | 389 | Klebsiella ornithinolytica |
| SAMN25989844 | 392 | 389 | Klebsiella ornithinolytica |
| SAMN25990043 | 392 | 389 | Klebsiella ornithinolytica |
| SAMN26796510 | 392 | 389 | Klebsiella ornithinolytica |
| SAMN28743851 | 392 | 389 | Klebsiella ornithinolytica |
| SAMN28887035 | 392 | 389 | Klebsiella ornithinolytica |
| SAMN29503370 | 392 | 389 | Klebsiella ornithinolytica |
| SAMN29503514 | 392 | 389 | Klebsiella ornithinolytica |
| SAMN29503521 | 392 | 389 | Klebsiella ornithinolytica |
| SAMN31650698 | 392 | 389 | Klebsiella ornithinolytica |
| SAMN31975269 | 392 | 389 | Klebsiella ornithinolytica |
| SAMN33274621 | 392 | 389 | Klebsiella ornithinolytica |
| SAMN33874247 | 392 | 389 | Klebsiella ornithinolytica |
| SAMN33874248 | 392 | 389 | Klebsiella ornithinolytica |
| SAMN33924912 | 392 | 389 | Klebsiella ornithinolytica |
| SAMN34029912 | 392 | 389 | Klebsiella ornithinolytica |
| SAMN34029915 | 392 | 389 | Klebsiella ornithinolytica |
| SAMN34030207 | 392 | 389 | Klebsiella ornithinolytica |
| SAMN34067561 | 392 | 389 | Klebsiella ornithinolytica |
| SAMN34068372 | 392 | 389 | Klebsiella ornithinolytica |
| SAMN34154773 | 392 | 389 | Klebsiella ornithinolytica |
| SAMN34340408 | 392 | 389 | Klebsiella ornithinolytica |
| SAMN34340411 | 392 | 389 | Klebsiella ornithinolytica |
| SAMN34382074 | 392 | 389 | Klebsiella ornithinolytica |
| SAMN35440192 | 392 | 389 | Klebsiella ornithinolytica |
| SAMN35440194 | 392 | 389 | Klebsiella ornithinolytica |
| DSM5175 | 392 | 389 | Klebsiella oxytoca |
| GCA_900977765 | 392 | 389 | Klebsiella oxytoca |
| GCF_000247855 | 392 | 389 | Klebsiella oxytoca |
| GCF_000247875 | 392 | 389 | Klebsiella oxytoca |
| GCF_000252915 | 392 | 389 | Klebsiella oxytoca |
| GCF_000269585 | 392 | 389 | Klebsiella oxytoca |
| GCF_000507385 | 392 | 389 | Klebsiella oxytoca |
| GCF_000527215 | 392 | 389 | Klebsiella oxytoca |
| GCF_000527235 | 392 | 389 | Klebsiella oxytoca |
| GCF_000607265 | 392 | 389 | Klebsiella oxytoca |
| GCF_001078175 | 392 | 389 | Klebsiella oxytoca |
| GCF_001594375 | 392 | 389 | Klebsiella oxytoca |
| GCF_001598695 | 392 | 389 | Klebsiella oxytoca |
| GCF_002265085 | N | 389 | Klebsiella oxytoca |
| GCF_002508265 | 392 | 389 | Klebsiella oxytoca |
| GCF_003812925 | 392 | 389 | Klebsiella oxytoca |
| GCF_003991285 | 392 | 389 | Klebsiella oxytoca |
| GCF_004005605 | 392 | 64 | Klebsiella oxytoca |
| GCF_004005605 | 392 | 389 | Klebsiella oxytoca |
| GCF_004360035 | 392 | 119 | Klebsiella oxytoca |
| GCF_004360035 | 392 | 389 | Klebsiella oxytoca |
| GCF_004785705 | 392 | 389 | Klebsiella oxytoca |
| GCF_008082015 | 392 | 389 | Klebsiella oxytoca |
| GCF_008082295 | 392 | 389 | Klebsiella oxytoca |
| GCF_009648375 | 392 | 389 | Klebsiella oxytoca |
| GCF_009648435 | 392 | 389 | Klebsiella oxytoca |
| GCF_009832375 | 392 | 389 | Klebsiella oxytoca |
| GCF_010365605 | 392 | 389 | Klebsiella oxytoca |
| GCF_012395905 | 392 | 389 | Klebsiella oxytoca |
| GCF_015265825 | 392 | 389 | Klebsiella oxytoca |
| GCF_015265865 | 392 | 389 | Klebsiella oxytoca |
| GCF_015551825 | 392 | 389 | Klebsiella oxytoca |
| GCF_015554545 | 392 | 389 | Klebsiella oxytoca |
| GCF_015559095 | 392 | 389 | Klebsiella oxytoca |
| GCF_016516145 | 392 | 389 | Klebsiella oxytoca |
| GCF_016529765 | 392 | 389 | Klebsiella oxytoca |
| GCF_016636025 | 392 | 389 | Klebsiella oxytoca |
| GCF_016636045 | 392 | 389 | Klebsiella oxytoca |
| GCF_016636055 | 392 | 389 | Klebsiella oxytoca |
| GCF_016636085 | 392 | 389 | Klebsiella oxytoca |
| GCF_016636105 | 392 | 389 | Klebsiella oxytoca |
| GCF_016636135 | 392 | 389 | Klebsiella oxytoca |
| GCF_016636165 | 392 | 389 | Klebsiella oxytoca |
| GCF_016643945 | 392 | 389 | Klebsiella oxytoca |
| GCF_016734995 | 392 | 389 | Klebsiella oxytoca |
| GCF_017310465 | 392 | 389 | Klebsiella oxytoca |
| GCF_018068785 | 392 | 389 | Klebsiella oxytoca |
| GCF_018441525 | 392 | 389 | Klebsiella oxytoca |
| GCF_018443185 | 392 | 389 | Klebsiella oxytoca |
| GCF_018443495 | 392 | 389 | Klebsiella oxytoca |
| GCF_018443665 | 392 | 389 | Klebsiella oxytoca |
| GCF_018443705 | 392 | 389 | Klebsiella oxytoca |
| GCF_018443785 | 392 | 389 | Klebsiella oxytoca |
| GCF_018443915 | 392 | 389 | Klebsiella oxytoca |
| GCF_018443995 | 392 | 389 | Klebsiella oxytoca |
| GCF_018444055 | 392 | 389 | Klebsiella oxytoca |
| GCF_018444115 | 392 | 389 | Klebsiella oxytoca |
| GCF_018444465 | 392 | 389 | Klebsiella oxytoca |
| GCF_018447075 | 392 | 389 | Klebsiella oxytoca |
| GCF_019677525 | 392 | 389 | Klebsiella oxytoca |
| GCF_019677765 | 392 | 389 | Klebsiella oxytoca |
| GCF_019771925 | 392 | 389 | Klebsiella oxytoca |
| GCF_021264605 | 392 | 389 | Klebsiella oxytoca |
| GCF_021373355 | 392 | 389 | Klebsiella oxytoca |
| GCF_021373415 | 392 | 389 | Klebsiella oxytoca |
| GCF_021398915 | 392 | 389 | Klebsiella oxytoca |
| GCF_022605305 | 392 | 389 | Klebsiella oxytoca |
| GCF_022685985 | 392 | 389 | Klebsiella oxytoca |
| GCF_023572405 | 392 | 389 | Klebsiella oxytoca |
| GCF_025579125 | 392 | 389 | Klebsiella oxytoca |
| GCF_025676565 | 392 | 389 | Klebsiella oxytoca |
| GCF_025732175 | 392 | 389 | Klebsiella oxytoca |
| GCF_025770255 | 392 | 389 | Klebsiella oxytoca |
| GCF_025863555 | 392 | 389 | Klebsiella oxytoca |
| GCF_025863735 | 392 | 389 | Klebsiella oxytoca |
| GCF_025863775 | 392 | 389 | Klebsiella oxytoca |
| GCF_025950225 | 392 | 389 | Klebsiella oxytoca |
| GCF_025950405 | N | 389 | Klebsiella oxytoca |
| GCF_026223815 | 392 | 389 | Klebsiella oxytoca |
| GCF_026223825 | 392 | 389 | Klebsiella oxytoca |
| GCF_026223895 | 392 | 389 | Klebsiella oxytoca |
| GCF_026224075 | 392 | 389 | Klebsiella oxytoca |
| GCF_026224115 | 392 | 389 | Klebsiella oxytoca |
| GCF_026224135 | 392 | 389 | Klebsiella oxytoca |
| GCF_026224155 | 392 | 389 | Klebsiella oxytoca |
| GCF_026224165 | 392 | 389 | Klebsiella oxytoca |
| GCF_026224195 | 392 | 389 | Klebsiella oxytoca |
| GCF_026224215 | 392 | 389 | Klebsiella oxytoca |
| GCF_026224235 | 392 | 389 | Klebsiella oxytoca |
| GCF_026224245 | 392 | 389 | Klebsiella oxytoca |
| GCF_026224275 | 392 | 389 | Klebsiella oxytoca |
| GCF_026224285 | 392 | 389 | Klebsiella oxytoca |
| GCF_026224315 | 392 | 389 | Klebsiella oxytoca |
| GCF_026224335 | 392 | 389 | Klebsiella oxytoca |
| GCF_026224355 | 392 | 389 | Klebsiella oxytoca |
| GCF_026224375 | 392 | 389 | Klebsiella oxytoca |
| GCF_026224475 | 392 | 389 | Klebsiella oxytoca |
| GCF_026224495 | 392 | 389 | Klebsiella oxytoca |
| GCF_026224505 | 392 | 389 | Klebsiella oxytoca |
| GCF_026224535 | 392 | 389 | Klebsiella oxytoca |
| GCF_026224545 | 392 | 389 | Klebsiella oxytoca |
| GCF_026224595 | 392 | 389 | Klebsiella oxytoca |
| GCF_026224645 | 392 | 389 | Klebsiella oxytoca |
| GCF_027257085 | 392 | 389 | Klebsiella oxytoca |
| GCF_027257135 | 392 | 389 | Klebsiella oxytoca |
| GCF_027257255 | 392 | 389 | Klebsiella oxytoca |
| GCF_029226505 | 392 | 389 | Klebsiella oxytoca |
| GCF_030194355 | 392 | 389 | Klebsiella oxytoca |
| GCF_030194365 | 392 | 389 | Klebsiella oxytoca |
| GCF_030194535 | 392 | 389 | Klebsiella oxytoca |
| GCF_030195115 | 392 | 389 | Klebsiella oxytoca |
| GCF_030195175 | 392 | 389 | Klebsiella oxytoca |
| GCF_030195225 | 392 | 389 | Klebsiella oxytoca |
| GCF_030195575 | 392 | 389 | Klebsiella oxytoca |
| GCF_030213545 | 392 | 389 | Klebsiella oxytoca |
| GCF_030225535 | 392 | 389 | Klebsiella oxytoca |
| GCF_030225605 | 392 | 389 | Klebsiella oxytoca |
| GCF_030225715 | 392 | 389 | Klebsiella oxytoca |
| GCF_030342765 | 392 | 389 | Klebsiella oxytoca |
| GCF_030342775 | 392 | 389 | Klebsiella oxytoca |
| GCF_030342805 | 392 | 389 | Klebsiella oxytoca |
| GCF_030342865 | 392 | 389 | Klebsiella oxytoca |
| GCF_030342875 | 392 | 389 | Klebsiella oxytoca |
| GCF_030342905 | 392 | 389 | Klebsiella oxytoca |
| GCF_030342955 | N | 389 | Klebsiella oxytoca |
| GCF_030342985 | 392 | 389 | Klebsiella oxytoca |
| GCF_030342995 | 392 | 389 | Klebsiella oxytoca |
| GCF_030343045 | 392 | 389 | Klebsiella oxytoca |
| GCF_030343055 | 392 | 389 | Klebsiella oxytoca |
| GCF_030343085 | N | 389 | Klebsiella oxytoca |
| GCF_030343095 | 392 | 389 | Klebsiella oxytoca |
| GCF_030343105 | 392 | 389 | Klebsiella oxytoca |
| GCF_030343145 | N | 389 | Klebsiella oxytoca |
| GCF_030343165 | 392 | 389 | Klebsiella oxytoca |
| GCF_030343225 | 392 | 389 | Klebsiella oxytoca |
| GCF_030343235 | 392 | 389 | Klebsiella oxytoca |
| GCF_030343265 | 392 | 389 | Klebsiella oxytoca |
| GCF_030343285 | 392 | 389 | Klebsiella oxytoca |
| GCF_030343295 | 392 | 389 | Klebsiella oxytoca |
| GCF_030343325 | 392 | 389 | Klebsiella oxytoca |
| GCF_030343345 | 392 | 389 | Klebsiella oxytoca |
| GCF_030343365 | 392 | 389 | Klebsiella oxytoca |
| GCF_030343375 | 392 | 389 | Klebsiella oxytoca |
| GCF_030343405 | 392 | 389 | Klebsiella oxytoca |
| GCF_030343425 | 392 | 389 | Klebsiella oxytoca |
| GCF_030343445 | 392 | 389 | Klebsiella oxytoca |
| GCF_030343465 | 392 | 389 | Klebsiella oxytoca |
| GCF_030343485 | 392 | 389 | Klebsiella oxytoca |
| GCF_030343495 | 392 | 389 | Klebsiella oxytoca |
| GCF_030343545 | 392 | 389 | Klebsiella oxytoca |
| GCF_030343565 | 392 | 389 | Klebsiella oxytoca |
| GCF_030343585 | N | 389 | Klebsiella oxytoca |
| GCF_030343625 | 392 | 389 | Klebsiella oxytoca |
| GCF_030343645 | 392 | 389 | Klebsiella oxytoca |
| GCF_030343655 | 392 | 389 | Klebsiella oxytoca |
| GCF_030343685 | 392 | 389 | Klebsiella oxytoca |
| GCF_030343705 | 392 | 389 | Klebsiella oxytoca |
| GCF_030343715 | N | 389 | Klebsiella oxytoca |
| GCF_030343785 | 392 | 389 | Klebsiella oxytoca |
| GCF_030343795 | N | 389 | Klebsiella oxytoca |
| GCF_030343825 | 392 | 389 | Klebsiella oxytoca |
| GCF_030343845 | 392 | 389 | Klebsiella oxytoca |
| GCF_030343865 | 392 | 389 | Klebsiella oxytoca |
| GCF_030343905 | 392 | 389 | Klebsiella oxytoca |
| GCF_030343925 | 392 | 389 | Klebsiella oxytoca |
| GCF_030343945 | 392 | 389 | Klebsiella oxytoca |
| GCF_030343955 | N | 389 | Klebsiella oxytoca |
| GCF_030343985 | N | 389 | Klebsiella oxytoca |
| GCF_030344005 | 392 | 389 | Klebsiella oxytoca |
| GCF_030344025 | N | 389 | Klebsiella oxytoca |
| GCF_030344035 | 392 | 389 | Klebsiella oxytoca |
| GCF_030344065 | 392 | 389 | Klebsiella oxytoca |
| GCF_030344075 | 392 | 389 | Klebsiella oxytoca |
| GCF_030344105 | N | 389 | Klebsiella oxytoca |
| GCF_030344145 | 392 | 389 | Klebsiella oxytoca |
| GCF_030344165 | N | 389 | Klebsiella oxytoca |
| GCF_030344175 | 392 | 389 | Klebsiella oxytoca |
| GCF_030344205 | N | 389 | Klebsiella oxytoca |
| GCF_030344215 | 392 | 389 | Klebsiella oxytoca |
| GCF_030344245 | N | 389 | Klebsiella oxytoca |
| GCF_030344255 | N | 389 | Klebsiella oxytoca |
| GCF_030344265 | 392 | 389 | Klebsiella oxytoca |
| GCF_030344325 | 392 | 389 | Klebsiella oxytoca |
| GCF_030344345 | 392 | 389 | Klebsiella oxytoca |
| GCF_030344385 | N | 389 | Klebsiella oxytoca |
| GCF_030344395 | N | 389 | Klebsiella oxytoca |
| GCF_030344405 | 392 | 389 | Klebsiella oxytoca |
| GCF_030344465 | 392 | 389 | Klebsiella oxytoca |
| GCF_030344505 | 392 | 389 | Klebsiella oxytoca |
| GCF_030344515 | 392 | 389 | Klebsiella oxytoca |
| GCF_030344545 | 392 | 389 | Klebsiella oxytoca |
| GCF_030344565 | 392 | 389 | Klebsiella oxytoca |
| GCF_030344585 | N | 389 | Klebsiella oxytoca |
| GCF_030344595 | N | 389 | Klebsiella oxytoca |
| GCF_030344605 | N | 389 | Klebsiella oxytoca |
| GCF_030344645 | 392 | 389 | Klebsiella oxytoca |
| GCF_030344665 | 392 | 389 | Klebsiella oxytoca |
| GCF_030344685 | N | 389 | Klebsiella oxytoca |
| GCF_030344695 | 392 | 389 | Klebsiella oxytoca |
| GCF_030344725 | 392 | 389 | Klebsiella oxytoca |
| GCF_030344735 | 392 | 389 | Klebsiella oxytoca |
| GCF_030344765 | 392 | 389 | Klebsiella oxytoca |
| GCF_030344785 | 392 | 389 | Klebsiella oxytoca |
| GCF_030344795 | N | 389 | Klebsiella oxytoca |
| GCF_030344825 | 392 | 389 | Klebsiella oxytoca |
| GCF_030344845 | N | 389 | Klebsiella oxytoca |
| GCF_030361405 | N | 389 | Klebsiella oxytoca |
| GCF_030361415 | 392 | 389 | Klebsiella oxytoca |
| GCF_030506015 | 392 | 389 | Klebsiella oxytoca |
| GCF_030506035 | 392 | 389 | Klebsiella oxytoca |
| GCF_030518375 | 392 | 389 | Klebsiella oxytoca |
| GCF_030544885 | 392 | 389 | Klebsiella oxytoca |
| GCF_030658935 | 392 | 389 | Klebsiella oxytoca |
| GCF_031799435 | 392 | 389 | Klebsiella oxytoca |
| GCF_031799655 | 392 | 389 | Klebsiella oxytoca |
| GCF_031799795 | 392 | 389 | Klebsiella oxytoca |
| GCF_031799975 | 392 | 389 | Klebsiella oxytoca |
| GCF_031800355 | 392 | 389 | Klebsiella oxytoca |
| GCF_032461035 | 392 | 389 | Klebsiella oxytoca |
| GCF_032670825 | 392 | 389 | Klebsiella oxytoca |
| GCF_032670945 | 392 | 389 | Klebsiella oxytoca |
| GCF_032671005 | 392 | 389 | Klebsiella oxytoca |
| GCF_032671045 | 392 | 389 | Klebsiella oxytoca |
| GCF_032671085 | 392 | 389 | Klebsiella oxytoca |
| GCF_032671225 | 392 | 389 | Klebsiella oxytoca |
| GCF_032677325 | 392 | 389 | Klebsiella oxytoca |
| GCF_032677545 | 392 | 64 | Klebsiella oxytoca |
| GCF_032677545 | 392 | 389 | Klebsiella oxytoca |
| GCF_033860475 | 392 | 389 | Klebsiella oxytoca |
| GCF_033864495 | 392 | 389 | Klebsiella oxytoca |
| GCF_033865735 | 392 | 389 | Klebsiella oxytoca |
| GCF_033898965 | 392 | 389 | Klebsiella oxytoca |
| GCF_033899945 | 392 | 389 | Klebsiella oxytoca |
| GCF_034506895 | 392 | 389 | Klebsiella oxytoca |
| GCF_035284165 | 392 | 389 | Klebsiella oxytoca |
| GCF_035284175 | 392 | 389 | Klebsiella oxytoca |
| GCF_035284205 | 392 | 389 | Klebsiella oxytoca |
| GCF_035786855 | 392 | 389 | Klebsiella oxytoca |
| GCF_035786895 | 392 | 389 | Klebsiella oxytoca |
| GCF_035794755 | 392 | 389 | Klebsiella oxytoca |
| GCF_036016465 | 392 | 389 | Klebsiella oxytoca |
| GCF_036016495 | 392 | 389 | Klebsiella oxytoca |
| GCF_036016545 | 392 | 389 | Klebsiella oxytoca |
| GCF_036016635 | 392 | 389 | Klebsiella oxytoca |
| GCF_036016645 | 392 | 389 | Klebsiella oxytoca |
| GCF_036542565 | 392 | 389 | Klebsiella oxytoca |
| GCF_036687455 | 392 | 389 | Klebsiella oxytoca |
| GCF_036908015 | 392 | 389 | Klebsiella oxytoca |
| GCF_036911735 | 392 | 389 | Klebsiella oxytoca |
| GCF_036952745 | 392 | 389 | Klebsiella oxytoca |
| GCF_036958305 | 392 | 389 | Klebsiella oxytoca |
| GCF_037151075 | 392 | 389 | Klebsiella oxytoca |
| GCF_038149365 | 392 | 389 | Klebsiella oxytoca |
| GCF_038149585 | 392 | 389 | Klebsiella oxytoca |
| GCF_039024105 | 392 | 389 | Klebsiella oxytoca |
| GCF_039571325 | 392 | 389 | Klebsiella oxytoca |
| GCF_039571445 | 392 | 389 | Klebsiella oxytoca |
| GCF_039605015 | 392 | 389 | Klebsiella oxytoca |
| GCF_040062315 | 392 | 389 | Klebsiella oxytoca |
| GCF_040065135 | 392 | 389 | Klebsiella oxytoca |
| GCF_040065315 | 392 | 389 | Klebsiella oxytoca |
| GCF_040097165 | 392 | 389 | Klebsiella oxytoca |
| GCF_040137435 | 392 | 389 | Klebsiella oxytoca |
| GCF_040138485 | 392 | 389 | Klebsiella oxytoca |
| GCF_040138515 | 392 | 389 | Klebsiella oxytoca |
| GCF_040138595 | 392 | 389 | Klebsiella oxytoca |
| GCF_040138605 | 392 | 389 | Klebsiella oxytoca |
| GCF_040138705 | 392 | 389 | Klebsiella oxytoca |
| GCF_040267715 | 392 | 389 | Klebsiella oxytoca |
| GCF_040561235 | 392 | 389 | Klebsiella oxytoca |
| GCF_040561445 | 392 | 389 | Klebsiella oxytoca |
| GCF_040564465 | 392 | 389 | Klebsiella oxytoca |
| GCF_040565235 | 392 | 389 | Klebsiella oxytoca |
| GCF_040566915 | 392 | 389 | Klebsiella oxytoca |
| GCF_040742065 | 392 | 389 | Klebsiella oxytoca |
| GCF_040742455 | 392 | 389 | Klebsiella oxytoca |
| GCF_041202595 | 392 | 389 | Klebsiella oxytoca |
| GCF_041353115 | 392 | 389 | Klebsiella oxytoca |
| GCF_041698715 | 392 | 389 | Klebsiella oxytoca |
| GCF_041699445 | 392 | 389 | Klebsiella oxytoca |
| GCF_041699465 | 392 | 389 | Klebsiella oxytoca |
| GCF_042137955 | 392 | 389 | Klebsiella oxytoca |
| GCF_043835255 | 392 | 389 | Klebsiella oxytoca |
| GCF_044004165 | 392 | 389 | Klebsiella oxytoca |
| GCF_044004215 | 392 | 389 | Klebsiella oxytoca |
| GCF_044006185 | 392 | 389 | Klebsiella oxytoca |
| GCF_044792955 | 392 | 389 | Klebsiella oxytoca |
| GCF_045261815 | 392 | 389 | Klebsiella oxytoca |
| GCF_045570355 | 392 | 389 | Klebsiella oxytoca |
| GCF_045583105 | 392 | 389 | Klebsiella oxytoca |
| GCF_045584145 | 392 | 389 | Klebsiella oxytoca |
| GCF_046030875 | 392 | 389 | Klebsiella oxytoca |
| GCF_046031235 | 392 | 389 | Klebsiella oxytoca |
| GCF_046366185 | 392 | 389 | Klebsiella oxytoca |
| GCF_046503475 | 392 | 389 | Klebsiella oxytoca |
| GCF_046578785 | 392 | 389 | Klebsiella oxytoca |
| GCF_046578805 | 392 | 389 | Klebsiella oxytoca |
| GCF_046580825 | 392 | 389 | Klebsiella oxytoca |
| GCF_046580835 | 392 | 389 | Klebsiella oxytoca |
| GCF_046602375 | 392 | 389 | Klebsiella oxytoca |
| GCF_046602755 | 392 | 389 | Klebsiella oxytoca |
| GCF_046603195 | 392 | 389 | Klebsiella oxytoca |
| GCF_900407115 | 392 | 389 | Klebsiella oxytoca |
| GCF_900451165 | 392 | 389 | Klebsiella oxytoca |
| GCF_900451235 | 392 | 389 | Klebsiella oxytoca |
| GCF_900451255 | 392 | 389 | Klebsiella oxytoca |
| GCF_900635105 | 392 | 389 | Klebsiella oxytoca |
| GCF_900636985 | 392 | 389 | Klebsiella oxytoca |
| GCF_901212425 | 392 | 389 | Klebsiella oxytoca |
| GCF_902158835 | 392 | 389 | Klebsiella oxytoca |
| GCF_902159605 | 392 | 389 | Klebsiella oxytoca |
| GCF_902159785 | 392 | 389 | Klebsiella oxytoca |
| GCF_902164285 | 392 | 389 | Klebsiella oxytoca |
| GCF_902164825 | 392 | 389 | Klebsiella oxytoca |
| GCF_902166465 | 392 | 389 | Klebsiella oxytoca |
| GCF_902363365 | 392 | 389 | Klebsiella oxytoca |
| GCF_904863345 | 392 | 389 | Klebsiella oxytoca |
| GCF_905232115 | 392 | 389 | Klebsiella oxytoca |
| GCF_905322525 | 392 | 389 | Klebsiella oxytoca |
| GCF_905329555 | 392 | 389 | Klebsiella oxytoca |
| GCF_905330835 | 392 | 389 | Klebsiella oxytoca |
| GCF_905331245 | 392 | 389 | Klebsiella oxytoca |
| GCF_905338025 | 392 | 389 | Klebsiella oxytoca |
| GCF_963887975 | 392 | 389 | Klebsiella oxytoca |
| GCF_964188445 | 392 | 389 | Klebsiella oxytoca |
| GCF_964188465 | 392 | 389 | Klebsiella oxytoca |
| GCF_964188475 | 392 | 389 | Klebsiella oxytoca |
| GCF_964207135 | 392 | 389 | Klebsiella oxytoca |
| GCF_964207205 | 392 | 389 | Klebsiella oxytoca |
| GCF_964207235 | 392 | 389 | Klebsiella oxytoca |
| GCF_964207445 | 392 | 389 | Klebsiella oxytoca |
| GCF_964300735 | 392 | 389 | Klebsiella oxytoca |
| GCF_964301075 | 392 | 389 | Klebsiella oxytoca |
| GCF_964301195 | 392 | 389 | Klebsiella oxytoca |
| GCF_964301215 | 392 | 389 | Klebsiella oxytoca |
| GCF_964301235 | 392 | 389 | Klebsiella oxytoca |
| GCF_964302825 | 392 | 389 | Klebsiella oxytoca |
| GCF_964302905 | 392 | 389 | Klebsiella oxytoca |
| GCF_964303285 | 392 | 389 | Klebsiella oxytoca |
| GCF_964303465 | 392 | 389 | Klebsiella oxytoca |
| GCF_964303765 | 392 | 389 | Klebsiella oxytoca |
| GCF_964303815 | 392 | 389 | Klebsiella oxytoca |
| GCF_964303825 | 392 | 389 | Klebsiella oxytoca |
| GCF_964304095 | 392 | 389 | Klebsiella oxytoca |
| GCF_964304135 | 392 | 389 | Klebsiella oxytoca |
| GCF_964304145 | 392 | 389 | Klebsiella oxytoca |
| GCF_964304165 | 392 | 389 | Klebsiella oxytoca |
| GCF_964304195 | 392 | 389 | Klebsiella oxytoca |
| GCF_964304215 | 392 | 389 | Klebsiella oxytoca |
| GCF_964304245 | 392 | 389 | Klebsiella oxytoca |
| GCF_964304265 | 392 | 389 | Klebsiella oxytoca |
| GCF_964304305 | 392 | 389 | Klebsiella oxytoca |
| GCF_964304335 | 392 | 389 | Klebsiella oxytoca |
| GCF_964304355 | 392 | 389 | Klebsiella oxytoca |
| GCF_964304365 | 392 | 389 | Klebsiella oxytoca |
| GCF_964304395 | 392 | 389 | Klebsiella oxytoca |
| GCF_964304405 | 392 | 389 | Klebsiella oxytoca |
| GCF_964304415 | 392 | 389 | Klebsiella oxytoca |
| GCF_964436405 | 392 | 389 | Klebsiella oxytoca |
| GFKo16 | 392 | 389 | Klebsiella oxytoca |
| GFKo47 | 392 | 389 | Klebsiella oxytoca |
| GFKo6 | 392 | 389 | Klebsiella oxytoca |
| Ko1 | 392 | 389 | Klebsiella oxytoca |
| Ko11 | 392 | 389 | Klebsiella oxytoca |
| Ko12 | 392 | 389 | Klebsiella oxytoca |
| Ko15 | 392 | 389 | Klebsiella oxytoca |
| Ko17 | 392 | 389 | Klebsiella oxytoca |
| Ko19 | 392 | 389 | Klebsiella oxytoca |
| Ko2 | 392 | 389 | Klebsiella oxytoca |
| Ko20 | 392 | 389 | Klebsiella oxytoca |
| Ko25 | 392 | 389 | Klebsiella oxytoca |
| Ko26 | 392 | 389 | Klebsiella oxytoca |
| Ko37 | 392 | 389 | Klebsiella oxytoca |
| Ko38 | 392 | 389 | Klebsiella oxytoca |
| Ko4 | 392 | 389 | Klebsiella oxytoca |
| Ko40 | 392 | 389 | Klebsiella oxytoca |
| Ko44 | 392 | 389 | Klebsiella oxytoca |
| Ko45 | 392 | 389 | Klebsiella oxytoca |
| Ko47 | 392 | 389 | Klebsiella oxytoca |
| Ko48 | 392 | 389 | Klebsiella oxytoca |
| Ko53 | 392 | 389 | Klebsiella oxytoca |
| Ko54 | 392 | 389 | Klebsiella oxytoca |
| Ko55 | 392 | 389 | Klebsiella oxytoca |
| Ko57 | 392 | 389 | Klebsiella oxytoca |
| Ko6 | 392 | 389 | Klebsiella oxytoca |
| Ko7 | 392 | 389 | Klebsiella oxytoca |
| GCA_000247915 | 392 | 389 | Klebsiella pasteurii |
| GCA_009757395 | 392 | 389 | Klebsiella pasteurii |
| GCA_013266985 | 392 | 389 | Klebsiella pasteurii |
| GCA_016616645 | 392 | 389 | Klebsiella pasteurii |
| GCF_012843205 | 392 | 389 | Klebsiella pasteurii |
| GCF_015550565 | 392 | 389 | Klebsiella pasteurii |
| GCF_015601345 | 392 | 389 | Klebsiella pasteurii |
| GCF_018139045 | 392 | 389 | Klebsiella pasteurii |
| GCF_018423175 | 392 | 389 | Klebsiella pasteurii |
| GCF_019661035 | 392 | 389 | Klebsiella pasteurii |
| GCF_026223975 | 392 | 389 | Klebsiella pasteurii |
| GCF_028368715 | 392 | 389 | Klebsiella pasteurii |
| GCF_030169945 | 392 | 389 | Klebsiella pasteurii |
| GCF_030193755 | 392 | 389 | Klebsiella pasteurii |
| GCF_030193795 | 392 | 389 | Klebsiella pasteurii |
| GCF_030343185 | 392 | 389 | Klebsiella pasteurii |
| GCF_030850055 | 392 | 389 | Klebsiella pasteurii |
| GCF_030850095 | 392 | 389 | Klebsiella pasteurii |
| GCF_030850165 | 392 | 389 | Klebsiella pasteurii |
| GCF_031455495 | 392 | 389 | Klebsiella pasteurii |
| GCF_031593235 | 392 | 389 | Klebsiella pasteurii |
| GCF_031799835 | 392 | 389 | Klebsiella pasteurii |
| GCF_031800155 | 392 | 389 | Klebsiella pasteurii |
| GCF_031800225 | 392 | 389 | Klebsiella pasteurii |
| GCF_031800375 | 392 | 389 | Klebsiella pasteurii |
| GCF_032677265 | 392 | 389 | Klebsiella pasteurii |
| GCF_032742035 | 392 | 389 | Klebsiella pasteurii |
| GCF_032742175 | 392 | 389 | Klebsiella pasteurii |
| GCF_032742395 | 392 | 389 | Klebsiella pasteurii |
| GCF_032746275 | 392 | 389 | Klebsiella pasteurii |
| GCF_032746355 | 392 | 389 | Klebsiella pasteurii |
| GCF_033099855 | 392 | 389 | Klebsiella pasteurii |
| GCF_033342075 | 392 | 389 | Klebsiella pasteurii |
| GCF_033899885 | 392 | 118 | Klebsiella pasteurii |
| GCF_033899885 | 392 | 389 | Klebsiella pasteurii |
| GCF_034427715 | 392 | 389 | Klebsiella pasteurii |
| GCF_036287845 | 392 | 389 | Klebsiella pasteurii |
| GCF_037052925 | 392 | 389 | Klebsiella pasteurii |
| GCF_039409695 | 392 | 389 | Klebsiella pasteurii |
| GCF_039409755 | 392 | 389 | Klebsiella pasteurii |
| GCF_040061895 | 392 | 389 | Klebsiella pasteurii |
| GCF_040062705 | 392 | 118 | Klebsiella pasteurii |
| GCF_040062705 | 392 | 389 | Klebsiella pasteurii |
| GCF_040064345 | 392 | 389 | Klebsiella pasteurii |
| GCF_040064435 | 392 | 389 | Klebsiella pasteurii |
| GCF_040139705 | 392 | 389 | Klebsiella pasteurii |
| GCF_041003725 | 392 | 389 | Klebsiella pasteurii |
| GCF_041003735 | 392 | 389 | Klebsiella pasteurii |
| GCF_041226105 | 392 | 389 | Klebsiella pasteurii |
| GCF_041283475 | 392 | 389 | Klebsiella pasteurii |
| GCF_042159875 | 392 | 389 | Klebsiella pasteurii |
| GCF_046299155 | 392 | 389 | Klebsiella pasteurii |
| GCF_046591815 | 392 | 389 | Klebsiella pasteurii |
| GCF_046592175 | 392 | 389 | Klebsiella pasteurii |
| GCF_046592195 | 392 | 389 | Klebsiella pasteurii |
| GCF_046603035 | 392 | 389 | Klebsiella pasteurii |
| GCF_901563825 | 392 | 389 | Klebsiella pasteurii |
| GCF_902158545 | 392 | 389 | Klebsiella pasteurii |
| GCF_902158575 | 392 | 389 | Klebsiella pasteurii |
| GCF_902158585 | 392 | 118 | Klebsiella pasteurii |
| GCF_902158585 | 392 | 389 | Klebsiella pasteurii |
| GCF_902158635 | 392 | 389 | Klebsiella pasteurii |
| GCF_902158645 | 392 | 389 | Klebsiella pasteurii |
| GCF_902158655 | 392 | 64 | Klebsiella pasteurii |
| GCF_902158655 | 392 | 389 | Klebsiella pasteurii |
| GCF_902158665 | 392 | 389 | Klebsiella pasteurii |
| GCF_902158675 | 392 | 389 | Klebsiella pasteurii |
| GCF_902158685 | 392 | 389 | Klebsiella pasteurii |
| GCF_902158695 | 392 | 389 | Klebsiella pasteurii |
| GCF_902158705 | 392 | 389 | Klebsiella pasteurii |
| GCF_902158715 | 392 | 118 | Klebsiella pasteurii |
| GCF_902158715 | 392 | 389 | Klebsiella pasteurii |
| GCF_902158725 | 392 | 389 | Klebsiella pasteurii |
| GCF_000648315 | 392 | 389 | Klebsiella planticola |
| GCF_000735435 | 392 | 389 | Klebsiella planticola |
| GCF_000737915 | 392 | 389 | Klebsiella planticola |
| GCF_000783935 | 392 | 389 | Klebsiella planticola |
| GCF_001049875 | 392 | 389 | Klebsiella planticola |
| GCF_001065685 | 392 | 389 | Klebsiella planticola |
| GCF_001663315 | 392 | 135 | Klebsiella planticola |
| GCF_001663325 | 392 | 389 | Klebsiella planticola |
| GCF_001903595 | 392 | 389 | Klebsiella planticola |
| GCF_002264145 | 392 | 389 | Klebsiella planticola |
| GCF_002554635 | 392 | 389 | Klebsiella planticola |
| GCF_002588355 | 392 | 389 | Klebsiella planticola |
| GCF_002635135 | 392 | 389 | Klebsiella planticola |
| GCF_002761975 | 392 | 389 | Klebsiella planticola |
| GCF_002906195 | 392 | 389 | Klebsiella planticola |
| GCF_003699975 | 392 | 389 | Klebsiella planticola |
| GCF_004024155 | 392 | 135 | Klebsiella planticola |
| GCF_004312145 | 392 | 389 | Klebsiella planticola |
| GCF_004312325 | 392 | 389 | Klebsiella planticola |
| GCF_004345285 | 392 | 389 | Klebsiella planticola |
| GCF_004365995 | 392 | 389 | Klebsiella planticola |
| GCF_004366295 | 392 | 389 | Klebsiella planticola |
| GCF_005048625 | 392 | 64 | Klebsiella planticola |
| GCF_005048625 | 392 | 389 | Klebsiella planticola |
| GCF_006757685 | 392 | 389 | Klebsiella planticola |
| GCF_008694045 | 392 | 389 | Klebsiella planticola |
| GCF_010598615 | 392 | 389 | Klebsiella planticola |
| GCF_010598665 | 392 | 389 | Klebsiella planticola |
| GCF_011290675 | 392 | 389 | Klebsiella planticola |
| GCF_013462275 | 392 | 64 | Klebsiella planticola |
| GCF_013462275 | 392 | 389 | Klebsiella planticola |
| GCF_014856235 | 392 | 389 | Klebsiella planticola |
| GCF_018443145 | 392 | 389 | Klebsiella planticola |
| GCF_018443165 | 392 | 135 | Klebsiella planticola |
| GCF_019856295 | 392 | 389 | Klebsiella planticola |
| GCF_019968885 | 392 | 389 | Klebsiella planticola |
| GCF_020115725 | 392 | 135 | Klebsiella planticola |
| GCF_021245905 | 392 | 389 | Klebsiella planticola |
| GCF_021441395 | 392 | 389 | Klebsiella planticola |
| GCF_022637595 | 392 | 64 | Klebsiella planticola |
| GCF_022637595 | 392 | 389 | Klebsiella planticola |
| GCF_024494745 | 392 | 389 | Klebsiella planticola |
| GCF_024609735 | 392 | 135 | Klebsiella planticola |
| GCF_025092595 | 392 | 389 | Klebsiella planticola |
| GCF_029635815 | 392 | 389 | Klebsiella planticola |
| GCF_032742975 | 392 | 389 | Klebsiella planticola |
| GCF_032742995 | 392 | 389 | Klebsiella planticola |
| GCF_032744815 | 392 | 389 | Klebsiella planticola |
| GCF_032745695 | 392 | 389 | Klebsiella planticola |
| GCF_033342015 | 392 | 389 | Klebsiella planticola |
| GCF_033436785 | 392 | 389 | Klebsiella planticola |
| GCF_033870315 | 392 | 389 | Klebsiella planticola |
| GCF_034318605 | 392 | 135 | Klebsiella planticola |
| GCF_034506695 | 392 | 389 | Klebsiella planticola |
| GCF_034506705 | 392 | 389 | Klebsiella planticola |
| GCF_034506835 | 392 | 389 | Klebsiella planticola |
| GCF_034930895 | 392 | 389 | Klebsiella planticola |
| GCF_036440415 | 392 | 389 | Klebsiella planticola |
| GCF_036440655 | 392 | 389 | Klebsiella planticola |
| GCF_036947115 | 392 | 224 | Klebsiella planticola |
| GCF_036948625 | 392 | 135 | Klebsiella planticola |
| GCF_036949845 | 392 | 135 | Klebsiella planticola |
| GCF_036956645 | 392 | 389 | Klebsiella planticola |
| GCF_036957185 | 392 | 209 | Klebsiella planticola |
| GCF_039519595 | 392 | 389 | Klebsiella planticola |
| GCF_040329095 | 392 | 135 | Klebsiella planticola |
| GCF_040529445 | 392 | 389 | Klebsiella planticola |
| GCF_040561385 | 392 | 389 | Klebsiella planticola |
| GCF_040838815 | 392 | 389 | Klebsiella planticola |
| GCF_041002795 | 392 | 389 | Klebsiella planticola |
| GCF_041002845 | 392 | 389 | Klebsiella planticola |
| GCF_041024025 | 392 | 389 | Klebsiella planticola |
| GCF_041092745 | 392 | 389 | Klebsiella planticola |
| GCF_041092835 | 392 | 389 | Klebsiella planticola |
| GCF_041092885 | 392 | 389 | Klebsiella planticola |
| GCF_041357935 | 392 | 135 | Klebsiella planticola |
| GCF_041357955 | 392 | 389 | Klebsiella planticola |
| GCF_041357965 | 392 | 389 | Klebsiella planticola |
| GCF_042945525 | 392 | 389 | Klebsiella planticola |
| GCF_045690375 | 392 | 389 | Klebsiella planticola |
| GCF_045690385 | 392 | 64 | Klebsiella planticola |
| GCF_045690385 | 392 | 389 | Klebsiella planticola |
| GCF_045690415 | 392 | 389 | Klebsiella planticola |
| GCF_045690425 | 392 | 389 | Klebsiella planticola |
| GCF_046578755 | 392 | 389 | Klebsiella planticola |
| GCF_046580845 | 392 | 389 | Klebsiella planticola |
| GCF_046580955 | 392 | 389 | Klebsiella planticola |
| GCF_046580965 | 392 | 389 | Klebsiella planticola |
| GCF_900083755 | 392 | 389 | Klebsiella planticola |
| GCF_900455785 | 392 | 389 | Klebsiella planticola |
| GCF_901420695 | 392 | 389 | Klebsiella planticola |
| GCF_901420725 | 392 | 389 | Klebsiella planticola |
| GCF_902160325 | 392 | 389 | Klebsiella planticola |
| GCF_902807195 | 392 | 389 | Klebsiella planticola |
| GCF_903935845 | 392 | 389 | Klebsiella planticola |
| SAMD00498608 | 69 | 389 | Klebsiella planticola |
| SAMD00498608 | 392 | 389 | Klebsiella planticola |
| SAMD00553992 | 392 | 389 | Klebsiella planticola |
| SAMD00553995 | 392 | 389 | Klebsiella planticola |
| SAMD00554001 | 392 | 389 | Klebsiella planticola |
| SAMEA10417050 | 392 | 389 | Klebsiella planticola |
| SAMEA111441560 | 392 | 389 | Klebsiella planticola |
| SAMEA111441566 | 392 | 389 | Klebsiella planticola |
| SAMEA111441598 | 392 | 389 | Klebsiella planticola |
| SAMEA111441612 | 392 | 389 | Klebsiella planticola |
| SAMEA111441613 | 392 | 389 | Klebsiella planticola |
| SAMEA11350655 | 392 | 199 | Klebsiella planticola |
| SAMEA11350656 | 392 | 135 | Klebsiella planticola |
| SAMEA1920250 | 392 | 389 | Klebsiella planticola |
| SAMEA2053118 | 392 | 389 | Klebsiella planticola |
| SAMEA2273635 | 392 | 389 | Klebsiella planticola |
| SAMEA2273831 | 392 | 389 | Klebsiella planticola |
| SAMEA4603025 | 392 | 389 | Klebsiella planticola |
| SAMEA4680411 | 392 | 389 | Klebsiella planticola |
| SAMEA4781547 | 392 | 389 | Klebsiella planticola |
| SAMEA4781982 | 392 | 389 | Klebsiella planticola |
| SAMEA4781985 | 392 | 389 | Klebsiella planticola |
| SAMEA4781993 | 392 | 389 | Klebsiella planticola |
| SAMEA4781995 | 392 | 389 | Klebsiella planticola |
| SAMEA4781997 | 392 | 389 | Klebsiella planticola |
| SAMEA4782010 | 392 | 389 | Klebsiella planticola |
| SAMEA4782020 | 392 | 389 | Klebsiella planticola |
| SAMEA5048749 | 392 | 389 | Klebsiella planticola |
| SAMEA5048758 | 392 | 389 | Klebsiella planticola |
| SAMEA5048782 | 392 | 389 | Klebsiella planticola |
| SAMEA5048783 | 392 | 389 | Klebsiella planticola |
| SAMEA5048786 | 392 | 389 | Klebsiella planticola |
| SAMEA5048789 | 392 | 389 | Klebsiella planticola |
| SAMEA5048791 | 392 | 389 | Klebsiella planticola |
| SAMEA5048850 | 392 | 389 | Klebsiella planticola |
| SAMEA5048880 | 392 | 389 | Klebsiella planticola |
| SAMEA5048881 | 392 | 389 | Klebsiella planticola |
| SAMEA5048924 | 392 | 389 | Klebsiella planticola |
| SAMEA5048930 | 392 | 389 | Klebsiella planticola |
| SAMEA5048975 | 392 | 389 | Klebsiella planticola |
| SAMEA5049016 | 392 | 389 | Klebsiella planticola |
| SAMEA5049039 | 392 | 389 | Klebsiella planticola |
| SAMEA5049081 | 392 | 389 | Klebsiella planticola |
| SAMEA5049169 | 392 | 389 | Klebsiella planticola |
| SAMEA5049170 | 392 | 389 | Klebsiella planticola |
| SAMEA5049194 | 392 | 389 | Klebsiella planticola |
| SAMEA5049198 | 392 | 389 | Klebsiella planticola |
| SAMEA5049217 | 392 | 389 | Klebsiella planticola |
| SAMEA5049306 | 392 | 389 | Klebsiella planticola |
| SAMEA5049314 | 392 | 389 | Klebsiella planticola |
| SAMEA5049316 | 392 | 389 | Klebsiella planticola |
| SAMEA5049323 | 392 | 389 | Klebsiella planticola |
| SAMEA5049326 | 392 | 389 | Klebsiella planticola |
| SAMEA5049356 | 392 | 389 | Klebsiella planticola |
| SAMEA5049396 | 392 | 64 | Klebsiella planticola |
| SAMEA5049396 | 392 | 389 | Klebsiella planticola |
| SAMEA5049409 | 392 | 389 | Klebsiella planticola |
| SAMEA5049414 | 392 | 389 | Klebsiella planticola |
| SAMEA5049451 | 392 | 389 | Klebsiella planticola |
| SAMEA5049453 | 392 | 389 | Klebsiella planticola |
| SAMEA5049454 | 392 | 389 | Klebsiella planticola |
| SAMEA5049456 | 392 | 389 | Klebsiella planticola |
| SAMEA5049519 | 392 | 389 | Klebsiella planticola |
| SAMEA5049569 | 392 | 389 | Klebsiella planticola |
| SAMEA5049579 | 392 | 389 | Klebsiella planticola |
| SAMEA5049581 | 392 | 389 | Klebsiella planticola |
| SAMEA5049586 | 392 | 389 | Klebsiella planticola |
| SAMEA5049602 | 392 | 389 | Klebsiella planticola |
| SAMEA5049603 | 392 | 389 | Klebsiella planticola |
| SAMEA5049627 | 392 | 389 | Klebsiella planticola |
| SAMEA5049635 | 392 | 389 | Klebsiella planticola |
| SAMEA5049648 | 392 | 389 | Klebsiella planticola |
| SAMEA5049649 | 392 | 389 | Klebsiella planticola |
| SAMEA5049656 | 392 | 389 | Klebsiella planticola |
| SAMEA5049657 | 392 | 389 | Klebsiella planticola |
| SAMEA5049662 | 392 | 389 | Klebsiella planticola |
| SAMEA5049666 | 392 | 389 | Klebsiella planticola |
| SAMEA5049704 | 392 | 389 | Klebsiella planticola |
| SAMEA5049711 | 392 | 389 | Klebsiella planticola |
| SAMEA5049712 | 392 | 389 | Klebsiella planticola |
| SAMEA5049713 | 392 | 389 | Klebsiella planticola |
| SAMEA5049714 | 392 | 389 | Klebsiella planticola |
| SAMEA5049715 | 392 | 389 | Klebsiella planticola |
| SAMEA5049716 | 392 | 389 | Klebsiella planticola |
| SAMEA5049717 | 392 | 389 | Klebsiella planticola |
| SAMEA5049718 | 392 | 389 | Klebsiella planticola |
| SAMEA5049719 | 392 | 389 | Klebsiella planticola |
| SAMEA5049720 | 392 | 389 | Klebsiella planticola |
| SAMEA5049721 | 392 | 389 | Klebsiella planticola |
| SAMEA5049739 | 392 | 389 | Klebsiella planticola |
| SAMEA5049786 | 392 | 64 | Klebsiella planticola |
| SAMEA5049786 | 392 | 389 | Klebsiella planticola |
| SAMEA5049881 | 392 | 389 | Klebsiella planticola |
| SAMEA5049884 | 392 | 389 | Klebsiella planticola |
| SAMEA5049931 | 392 | 389 | Klebsiella planticola |
| SAMEA5050216 | 392 | 389 | Klebsiella planticola |
| SAMEA5050223 | 392 | 389 | Klebsiella planticola |
| SAMEA5050257 | 392 | 389 | Klebsiella planticola |
| SAMEA5050266 | 392 | 389 | Klebsiella planticola |
| SAMEA5050267 | 392 | 389 | Klebsiella planticola |
| SAMEA5050285 | 392 | 389 | Klebsiella planticola |
| SAMEA5050287 | 392 | 389 | Klebsiella planticola |
| SAMEA5050288 | 392 | 389 | Klebsiella planticola |
| SAMEA5050301 | 392 | 389 | Klebsiella planticola |
| SAMEA5050302 | 392 | 389 | Klebsiella planticola |
| SAMEA5684198 | 392 | 389 | Klebsiella planticola |
| SAMEA5684208 | 392 | 389 | Klebsiella planticola |
| SAMEA5684222 | 392 | 389 | Klebsiella planticola |
| SAMEA5684231 | 392 | 389 | Klebsiella planticola |
| SAMEA5684248 | 392 | 389 | Klebsiella planticola |
| SAMEA5684281 | 392 | 389 | Klebsiella planticola |
| SAMEA5684293 | 392 | 389 | Klebsiella planticola |
| SAMEA5684307 | 392 | 389 | Klebsiella planticola |
| SAMEA5684312 | 392 | 389 | Klebsiella planticola |
| SAMEA5684314 | 392 | 389 | Klebsiella planticola |
| SAMEA5751433 | 392 | 389 | Klebsiella planticola |
| SAMEA5751871 | 392 | 389 | Klebsiella planticola |
| SAMEA5996264 | 392 | 389 | Klebsiella planticola |
| SAMEA5996267 | 392 | 389 | Klebsiella planticola |
| SAMEA6368766 | 392 | 389 | Klebsiella planticola |
| SAMEA6544381 | 392 | 389 | Klebsiella planticola |
| SAMEA6656491 | 392 | 389 | Klebsiella planticola |
| SAMEA7198819 | 392 | 389 | Klebsiella planticola |
| SAMEA7198820 | 392 | 389 | Klebsiella planticola |
| SAMN04448496 | 392 | 389 | Klebsiella planticola |
| SAMN07692508 | 392 | 389 | Klebsiella planticola |
| SAMN12138074 | 392 | 389 | Klebsiella planticola |
| SAMN12250659 | 392 | 135 | Klebsiella planticola |
| SAMN12289331 | 392 | 389 | Klebsiella planticola |
| SAMN12289377 | 392 | 389 | Klebsiella planticola |
| SAMN12774118 | 392 | 389 | Klebsiella planticola |
| SAMN12774253 | 392 | 389 | Klebsiella planticola |
| SAMN12774268 | 392 | 389 | Klebsiella planticola |
| SAMN15566992 | 392 | 156 | Klebsiella planticola |
| SAMN15566992 | 392 | 196 | Klebsiella planticola |
| SAMN15566994 | 392 | 135 | Klebsiella planticola |
| SAMN15868157 | 392 | 389 | Klebsiella planticola |
| SAMN15868169 | 392 | 195 | Klebsiella planticola |
| SAMN15868699 | 392 | 196 | Klebsiella planticola |
| SAMN15869005 | 392 | 135 | Klebsiella planticola |
| SAMN16233066 | 392 | 64 | Klebsiella planticola |
| SAMN16233066 | 392 | 389 | Klebsiella planticola |
| SAMN16233069 | 392 | 64 | Klebsiella planticola |
| SAMN16233069 | 392 | 389 | Klebsiella planticola |
| SAMN16233071 | 392 | 64 | Klebsiella planticola |
| SAMN16233071 | 392 | 389 | Klebsiella planticola |
| SAMN16233102 | 392 | 64 | Klebsiella planticola |
| SAMN16233102 | 392 | 389 | Klebsiella planticola |
| SAMN16233112 | 392 | 64 | Klebsiella planticola |
| SAMN16233112 | 392 | 389 | Klebsiella planticola |
| SAMN16233129 | 392 | 64 | Klebsiella planticola |
| SAMN16233129 | 392 | 389 | Klebsiella planticola |
| SAMN16233148 | 392 | 64 | Klebsiella planticola |
| SAMN16233148 | 392 | 389 | Klebsiella planticola |
| SAMN16233169 | 392 | 64 | Klebsiella planticola |
| SAMN16233169 | 392 | 389 | Klebsiella planticola |
| SAMN16233351 | 392 | 389 | Klebsiella planticola |
| SAMN16278278 | 392 | 156 | Klebsiella planticola |
| SAMN18207222 | 392 | 389 | Klebsiella planticola |
| SAMN19667690 | 392 | 389 | Klebsiella planticola |
| SAMN19856776 | 392 | 195 | Klebsiella planticola |
| SAMN19856780 | 392 | 389 | Klebsiella planticola |
| SAMN22968114 | 392 | 135 | Klebsiella planticola |
| SAMN24019754 | 392 | 194 | Klebsiella planticola |
| SAMN24297127 | 392 | 64 | Klebsiella planticola |
| SAMN24297127 | 392 | 389 | Klebsiella planticola |
| SAMN25538142 | 392 | 389 | Klebsiella planticola |
| SAMN25988452 | 392 | 389 | Klebsiella planticola |
| SAMN25988455 | 392 | 389 | Klebsiella planticola |
| SAMN25988471 | 392 | 389 | Klebsiella planticola |
| SAMN25988654 | 392 | 389 | Klebsiella planticola |
| SAMN25988681 | 392 | 389 | Klebsiella planticola |
| SAMN25988685 | 392 | 389 | Klebsiella planticola |
| SAMN25988732 | 392 | 389 | Klebsiella planticola |
| SAMN25988746 | 392 | 135 | Klebsiella planticola |
| SAMN25988747 | 392 | 389 | Klebsiella planticola |
| SAMN25988758 | 392 | 389 | Klebsiella planticola |
| SAMN25988769 | 392 | 389 | Klebsiella planticola |
| SAMN25988842 | 392 | 389 | Klebsiella planticola |
| SAMN25988870 | 392 | 389 | Klebsiella planticola |
| SAMN25988882 | 392 | 389 | Klebsiella planticola |
| SAMN25988886 | 392 | 389 | Klebsiella planticola |
| SAMN25988903 | 392 | 389 | Klebsiella planticola |
| SAMN25988916 | 392 | 389 | Klebsiella planticola |
| SAMN25988980 | 392 | 389 | Klebsiella planticola |
| SAMN25988988 | 392 | 389 | Klebsiella planticola |
| SAMN25988991 | 392 | 389 | Klebsiella planticola |
| SAMN25989018 | 392 | 389 | Klebsiella planticola |
| SAMN25989024 | 392 | 389 | Klebsiella planticola |
| SAMN25989171 | 392 | 389 | Klebsiella planticola |
| SAMN25989173 | 392 | 244 | Klebsiella planticola |
| SAMN25989198 | 392 | 389 | Klebsiella planticola |
| SAMN25989201 | 392 | 389 | Klebsiella planticola |
| SAMN25989217 | 392 | 389 | Klebsiella planticola |
| SAMN25989256 | 392 | 389 | Klebsiella planticola |
| SAMN25989301 | 392 | 389 | Klebsiella planticola |
| SAMN25989304 | 392 | 389 | Klebsiella planticola |
| SAMN25989350 | 392 | 389 | Klebsiella planticola |
| SAMN25989367 | 392 | 389 | Klebsiella planticola |
| SAMN25989424 | 392 | 244 | Klebsiella planticola |
| SAMN25989470 | 392 | 389 | Klebsiella planticola |
| SAMN25989506 | 392 | 389 | Klebsiella planticola |
| SAMN25989539 | 392 | 389 | Klebsiella planticola |
| SAMN25989553 | 392 | 389 | Klebsiella planticola |
| SAMN25989593 | 392 | 389 | Klebsiella planticola |
| SAMN25989596 | 392 | 389 | Klebsiella planticola |
| SAMN25989628 | 392 | 389 | Klebsiella planticola |
| SAMN25989636 | 392 | 389 | Klebsiella planticola |
| SAMN25989663 | 392 | 389 | Klebsiella planticola |
| SAMN25989696 | 392 | 389 | Klebsiella planticola |
| SAMN25989722 | 392 | 389 | Klebsiella planticola |
| SAMN25989731 | 392 | 389 | Klebsiella planticola |
| SAMN25989787 | 392 | 389 | Klebsiella planticola |
| SAMN25989837 | 392 | 389 | Klebsiella planticola |
| SAMN25989871 | 392 | 389 | Klebsiella planticola |
| SAMN25989881 | 392 | 389 | Klebsiella planticola |
| SAMN25989953 | 392 | 389 | Klebsiella planticola |
| SAMN25989955 | 392 | 389 | Klebsiella planticola |
| SAMN25989958 | 392 | 389 | Klebsiella planticola |
| SAMN25989996 | 392 | 389 | Klebsiella planticola |
| SAMN25990009 | 392 | 389 | Klebsiella planticola |
| SAMN25990015 | 392 | 389 | Klebsiella planticola |
| SAMN26419480 | 392 | 389 | Klebsiella planticola |
| SAMN26891831 | 392 | 389 | Klebsiella planticola |
| SAMN26891832 | 392 | 389 | Klebsiella planticola |
| SAMN26891833 | 392 | 389 | Klebsiella planticola |
| SAMN29503595 | 392 | 64 | Klebsiella planticola |
| SAMN29503595 | 392 | 195 | Klebsiella planticola |
| SAMN29503829 | 392 | 199 | Klebsiella planticola |
| SAMN29503829 | 392 | 200 | Klebsiella planticola |
| SAMN30498012 | 392 | 389 | Klebsiella planticola |
| SAMN30498055 | 392 | 389 | Klebsiella planticola |
| SAMN34403743 | 392 | 389 | Klebsiella planticola |
| ERR024818 | 229 | 389 | Klebsiella pneumoniae |
| ERR024818 | 392 | 389 | Klebsiella pneumoniae |
| ERR024819 | 229 | 389 | Klebsiella pneumoniae |
| ERR024819 | 392 | 389 | Klebsiella pneumoniae |
| ERR024821 | 229 | 389 | Klebsiella pneumoniae |
| ERR024821 | 392 | 389 | Klebsiella pneumoniae |
| ERR024822 | 229 | 389 | Klebsiella pneumoniae |
| ERR024822 | 392 | 389 | Klebsiella pneumoniae |
| ERR024823 | 229 | 389 | Klebsiella pneumoniae |
| ERR024823 | 392 | 389 | Klebsiella pneumoniae |
| ERR024824 | 229 | 389 | Klebsiella pneumoniae |
| ERR024824 | 392 | 389 | Klebsiella pneumoniae |
| ERR024828 | 229 | 389 | Klebsiella pneumoniae |
| ERR024828 | 392 | 389 | Klebsiella pneumoniae |
| ERR024831 | 392 | 389 | Klebsiella pneumoniae |
| ERR024832 | 392 | 389 | Klebsiella pneumoniae |
| ERR024833 | 229 | 389 | Klebsiella pneumoniae |
| ERR024833 | 392 | 389 | Klebsiella pneumoniae |
| ERR024835 | 229 | 389 | Klebsiella pneumoniae |
| ERR024835 | 392 | 389 | Klebsiella pneumoniae |
| ERR024836 | 229 | 389 | Klebsiella pneumoniae |
| ERR024836 | 392 | 389 | Klebsiella pneumoniae |
| ERR024837 | 392 | 389 | Klebsiella pneumoniae |
| ERR024838 | 229 | 389 | Klebsiella pneumoniae |
| ERR024838 | 392 | 389 | Klebsiella pneumoniae |
| ERR024839 | 229 | 389 | Klebsiella pneumoniae |
| ERR024839 | 392 | 389 | Klebsiella pneumoniae |
| ERR024840 | 229 | 389 | Klebsiella pneumoniae |
| ERR024840 | 392 | 389 | Klebsiella pneumoniae |
| ERR024843 | 229 | 389 | Klebsiella pneumoniae |
| ERR024843 | 392 | 389 | Klebsiella pneumoniae |
| ERR024844 | 229 | 389 | Klebsiella pneumoniae |
| ERR024844 | 392 | 389 | Klebsiella pneumoniae |
| ERR024845 | 229 | 389 | Klebsiella pneumoniae |
| ERR024845 | 392 | 389 | Klebsiella pneumoniae |
| ERR024847 | 45 | 389 | Klebsiella pneumoniae |
| ERR024847 | 157 | 389 | Klebsiella pneumoniae |
| ERR024847 | 229 | 389 | Klebsiella pneumoniae |
| ERR024847 | 342 | 389 | Klebsiella pneumoniae |
| ERR024848 | 229 | 389 | Klebsiella pneumoniae |
| ERR024848 | 392 | 389 | Klebsiella pneumoniae |
| ERR024850 | 229 | 389 | Klebsiella pneumoniae |
| ERR024850 | 392 | 389 | Klebsiella pneumoniae |
| ERR024851 | 392 | 389 | Klebsiella pneumoniae |
| ERR024852 | 229 | 389 | Klebsiella pneumoniae |
| ERR024852 | 392 | 389 | Klebsiella pneumoniae |
| ERR024854 | 392 | 389 | Klebsiella pneumoniae |
| ERR025098 | 392 | 389 | Klebsiella pneumoniae |
| ERR025099 | 229 | 389 | Klebsiella pneumoniae |
| ERR025099 | 392 | 389 | Klebsiella pneumoniae |
| ERR025100 | 392 | 389 | Klebsiella pneumoniae |
| ERR025101 | 392 | 389 | Klebsiella pneumoniae |
| ERR025103 | 392 | 389 | Klebsiella pneumoniae |
| ERR025104 | 229 | 389 | Klebsiella pneumoniae |
| ERR025104 | 392 | 389 | Klebsiella pneumoniae |
| ERR025105 | 229 | 389 | Klebsiella pneumoniae |
| ERR025105 | 392 | 389 | Klebsiella pneumoniae |
| ERR025107 | 229 | 389 | Klebsiella pneumoniae |
| ERR025107 | 392 | 389 | Klebsiella pneumoniae |
| ERR025108 | 229 | 389 | Klebsiella pneumoniae |
| ERR025108 | 392 | 389 | Klebsiella pneumoniae |
| ERR025109 | 229 | 389 | Klebsiella pneumoniae |
| ERR025109 | 392 | 389 | Klebsiella pneumoniae |
| ERR025111 | 392 | 389 | Klebsiella pneumoniae |
| ERR025112 | 229 | 389 | Klebsiella pneumoniae |
| ERR025112 | 392 | 389 | Klebsiella pneumoniae |
| ERR025113 | 392 | 389 | Klebsiella pneumoniae |
| ERR025115 | 229 | 389 | Klebsiella pneumoniae |
| ERR025115 | 392 | 389 | Klebsiella pneumoniae |
| ERR025116 | 229 | 389 | Klebsiella pneumoniae |
| ERR025116 | 392 | 389 | Klebsiella pneumoniae |
| ERR025117 | 229 | 389 | Klebsiella pneumoniae |
| ERR025117 | 392 | 389 | Klebsiella pneumoniae |
| ERR025118 | 229 | 389 | Klebsiella pneumoniae |
| ERR025118 | 392 | 389 | Klebsiella pneumoniae |
| ERR025119 | 392 | 389 | Klebsiella pneumoniae |
| ERR025121 | 229 | 389 | Klebsiella pneumoniae |
| ERR025121 | 392 | 389 | Klebsiella pneumoniae |
| ERR025125 | 229 | 64 | Klebsiella pneumoniae |
| ERR025125 | 229 | 389 | Klebsiella pneumoniae |
| ERR025125 | 392 | 64 | Klebsiella pneumoniae |
| ERR025125 | 392 | 389 | Klebsiella pneumoniae |
| ERR025126 | 392 | 389 | Klebsiella pneumoniae |
| ERR025128 | 392 | 389 | Klebsiella pneumoniae |
| ERR025131 | 229 | 389 | Klebsiella pneumoniae |
| ERR025131 | 392 | 389 | Klebsiella pneumoniae |
| ERR025132 | 229 | 389 | Klebsiella pneumoniae |
| ERR025132 | 392 | 389 | Klebsiella pneumoniae |
| ERR025133 | 229 | 389 | Klebsiella pneumoniae |
| ERR025133 | 392 | 389 | Klebsiella pneumoniae |
| ERR025135 | 229 | 389 | Klebsiella pneumoniae |
| ERR025135 | 392 | 389 | Klebsiella pneumoniae |
| ERR025137 | 229 | 389 | Klebsiella pneumoniae |
| ERR025137 | 392 | 389 | Klebsiella pneumoniae |
| ERR025139 | 392 | 389 | Klebsiella pneumoniae |
| ERR025140 | 229 | 389 | Klebsiella pneumoniae |
| ERR025140 | 392 | 389 | Klebsiella pneumoniae |
| ERR025141 | 392 | 389 | Klebsiella pneumoniae |
| ERR025142 | 229 | 389 | Klebsiella pneumoniae |
| ERR025142 | 392 | 389 | Klebsiella pneumoniae |
| ERR025143 | 229 | 389 | Klebsiella pneumoniae |
| ERR025143 | 392 | 389 | Klebsiella pneumoniae |
| ERR025144 | 229 | 389 | Klebsiella pneumoniae |
| ERR025144 | 392 | 389 | Klebsiella pneumoniae |
| ERR025146 | 229 | 389 | Klebsiella pneumoniae |
| ERR025146 | 392 | 389 | Klebsiella pneumoniae |
| ERR025147 | 229 | 389 | Klebsiella pneumoniae |
| ERR025147 | 392 | 389 | Klebsiella pneumoniae |
| ERR025150 | 226 | 389 | Klebsiella pneumoniae |
| ERR025150 | 229 | 389 | Klebsiella pneumoniae |
| ERR025151 | 229 | 389 | Klebsiella pneumoniae |
| ERR025151 | 392 | 389 | Klebsiella pneumoniae |
| ERR025152 | 229 | 389 | Klebsiella pneumoniae |
| ERR025152 | 392 | 389 | Klebsiella pneumoniae |
| ERR025154 | 229 | 389 | Klebsiella pneumoniae |
| ERR025154 | 392 | 389 | Klebsiella pneumoniae |
| ERR025155 | 392 | 64 | Klebsiella pneumoniae |
| ERR025155 | 392 | 389 | Klebsiella pneumoniae |
| ERR025156 | 392 | 389 | Klebsiella pneumoniae |
| ERR025157 | 229 | 389 | Klebsiella pneumoniae |
| ERR025157 | 392 | 389 | Klebsiella pneumoniae |
| ERR025158 | 392 | 389 | Klebsiella pneumoniae |
| ERR025159 | 229 | 389 | Klebsiella pneumoniae |
| ERR025159 | 392 | 389 | Klebsiella pneumoniae |
| ERR025160 | 229 | 389 | Klebsiella pneumoniae |
| ERR025160 | 392 | 389 | Klebsiella pneumoniae |
| ERR025161 | 229 | 389 | Klebsiella pneumoniae |
| ERR025161 | 392 | 389 | Klebsiella pneumoniae |
| ERR025462 | 392 | 389 | Klebsiella pneumoniae |
| ERR025464 | 392 | 389 | Klebsiella pneumoniae |
| ERR025465 | 229 | 389 | Klebsiella pneumoniae |
| ERR025465 | 392 | 389 | Klebsiella pneumoniae |
| ERR025468 | 229 | 389 | Klebsiella pneumoniae |
| ERR025468 | 392 | 389 | Klebsiella pneumoniae |
| ERR025469 | 392 | 389 | Klebsiella pneumoniae |
| ERR025470 | 229 | 389 | Klebsiella pneumoniae |
| ERR025470 | 392 | 389 | Klebsiella pneumoniae |
| ERR025471 | 229 | 389 | Klebsiella pneumoniae |
| ERR025471 | 392 | 389 | Klebsiella pneumoniae |
| ERR025472 | 229 | 389 | Klebsiella pneumoniae |
| ERR025472 | 392 | 389 | Klebsiella pneumoniae |
| ERR025473 | 392 | 389 | Klebsiella pneumoniae |
| ERR025475 | 229 | 389 | Klebsiella pneumoniae |
| ERR025475 | 392 | 389 | Klebsiella pneumoniae |
| ERR025477 | 229 | 389 | Klebsiella pneumoniae |
| ERR025477 | 392 | 389 | Klebsiella pneumoniae |
| ERR025478 | 392 | 389 | Klebsiella pneumoniae |
| ERR025479 | 229 | 389 | Klebsiella pneumoniae |
| ERR025479 | 392 | 389 | Klebsiella pneumoniae |
| ERR025482 | 392 | 389 | Klebsiella pneumoniae |
| ERR025483 | 392 | 389 | Klebsiella pneumoniae |
| ERR025484 | 229 | 389 | Klebsiella pneumoniae |
| ERR025484 | 392 | 389 | Klebsiella pneumoniae |
| ERR025485 | 392 | 389 | Klebsiella pneumoniae |
| ERR025486 | 229 | 389 | Klebsiella pneumoniae |
| ERR025486 | 392 | 389 | Klebsiella pneumoniae |
| ERR025489 | 392 | 389 | Klebsiella pneumoniae |
| ERR025491 | 392 | 389 | Klebsiella pneumoniae |
| ERR025494 | 229 | 389 | Klebsiella pneumoniae |
| ERR025494 | 392 | 389 | Klebsiella pneumoniae |
| ERR025495 | 229 | 389 | Klebsiella pneumoniae |
| ERR025495 | 392 | 389 | Klebsiella pneumoniae |
| ERR025496 | 229 | 389 | Klebsiella pneumoniae |
| ERR025496 | 392 | 389 | Klebsiella pneumoniae |
| ERR025497 | 392 | 389 | Klebsiella pneumoniae |
| ERR025498 | 229 | 389 | Klebsiella pneumoniae |
| ERR025498 | 392 | 389 | Klebsiella pneumoniae |
| ERR025499 | 392 | 389 | Klebsiella pneumoniae |
| ERR025501 | 229 | 389 | Klebsiella pneumoniae |
| ERR025501 | 392 | 389 | Klebsiella pneumoniae |
| ERR025502 | 229 | 389 | Klebsiella pneumoniae |
| ERR025502 | 392 | 389 | Klebsiella pneumoniae |
| ERR025503 | 229 | 64 | Klebsiella pneumoniae |
| ERR025503 | 229 | 389 | Klebsiella pneumoniae |
| ERR025503 | 392 | 64 | Klebsiella pneumoniae |
| ERR025503 | 392 | 389 | Klebsiella pneumoniae |
| ERR025504 | 229 | 389 | Klebsiella pneumoniae |
| ERR025504 | 392 | 389 | Klebsiella pneumoniae |
| ERR025505 | 229 | 389 | Klebsiella pneumoniae |
| ERR025505 | 392 | 389 | Klebsiella pneumoniae |
| ERR025506 | 392 | 389 | Klebsiella pneumoniae |
| ERR025510 | 392 | 389 | Klebsiella pneumoniae |
| ERR025511 | 392 | 389 | Klebsiella pneumoniae |
| ERR025512 | 229 | 389 | Klebsiella pneumoniae |
| ERR025512 | 392 | 389 | Klebsiella pneumoniae |
| ERR025515 | 392 | 389 | Klebsiella pneumoniae |
| ERR025515 | 436 | 389 | Klebsiella pneumoniae |
| ERR025516 | 392 | 389 | Klebsiella pneumoniae |
| ERR025517 | 392 | 389 | Klebsiella pneumoniae |
| ERR025518 | 392 | 389 | Klebsiella pneumoniae |
| ERR025519 | 229 | 389 | Klebsiella pneumoniae |
| ERR025519 | 392 | 389 | Klebsiella pneumoniae |
| ERR025520 | 229 | 389 | Klebsiella pneumoniae |
| ERR025520 | 392 | 389 | Klebsiella pneumoniae |
| ERR025521 | 392 | 389 | Klebsiella pneumoniae |
| ERR025522 | 229 | 64 | Klebsiella pneumoniae |
| ERR025522 | 229 | 389 | Klebsiella pneumoniae |
| ERR025522 | 392 | 64 | Klebsiella pneumoniae |
| ERR025522 | 392 | 389 | Klebsiella pneumoniae |
| ERR025523 | 229 | 389 | Klebsiella pneumoniae |
| ERR025523 | 392 | 389 | Klebsiella pneumoniae |
| ERR025524 | 229 | 389 | Klebsiella pneumoniae |
| ERR025524 | 392 | 389 | Klebsiella pneumoniae |
| ERR025525 | 392 | 389 | Klebsiella pneumoniae |
| ERR025527 | 392 | 389 | Klebsiella pneumoniae |
| ERR025531 | 229 | 389 | Klebsiella pneumoniae |
| ERR025531 | 392 | 389 | Klebsiella pneumoniae |
| ERR025532 | 229 | 389 | Klebsiella pneumoniae |
| ERR025532 | 392 | 389 | Klebsiella pneumoniae |
| ERR025534 | 229 | 389 | Klebsiella pneumoniae |
| ERR025534 | 392 | 389 | Klebsiella pneumoniae |
| ERR025535 | 229 | 389 | Klebsiella pneumoniae |
| ERR025535 | 392 | 389 | Klebsiella pneumoniae |
| ERR025535 | 436 | 389 | Klebsiella pneumoniae |
| ERR025536 | 392 | 389 | Klebsiella pneumoniae |
| ERR025538 | 392 | 389 | Klebsiella pneumoniae |
| ERR025540 | 229 | 389 | Klebsiella pneumoniae |
| ERR025540 | 392 | 389 | Klebsiella pneumoniae |
| ERR025541 | 229 | 306 | Klebsiella pneumoniae |
| ERR025541 | 392 | 306 | Klebsiella pneumoniae |
| ERR025542 | 392 | 389 | Klebsiella pneumoniae |
| ERR025543 | 229 | 389 | Klebsiella pneumoniae |
| ERR025543 | 392 | 389 | Klebsiella pneumoniae |
| ERR025544 | 392 | 389 | Klebsiella pneumoniae |
| ERR025545 | 229 | N | Klebsiella pneumoniae |
| ERR025545 | 392 | N | Klebsiella pneumoniae |
| ERR025546 | 229 | 389 | Klebsiella pneumoniae |
| ERR025546 | 392 | 389 | Klebsiella pneumoniae |
| ERR025547 | 229 | 389 | Klebsiella pneumoniae |
| ERR025547 | 392 | 389 | Klebsiella pneumoniae |
| ERR025548 | 229 | 389 | Klebsiella pneumoniae |
| ERR025548 | 392 | 389 | Klebsiella pneumoniae |
| ERR025550 | 392 | 389 | Klebsiella pneumoniae |
| ERR025553 | 392 | 389 | Klebsiella pneumoniae |
| ERR025554 | 392 | 389 | Klebsiella pneumoniae |
| ERR025555 | 229 | 389 | Klebsiella pneumoniae |
| ERR025555 | 392 | 389 | Klebsiella pneumoniae |
| ERR025557 | 392 | 389 | Klebsiella pneumoniae |
| ERR025558 | 392 | 389 | Klebsiella pneumoniae |
| ERR025561 | 229 | 389 | Klebsiella pneumoniae |
| ERR025561 | 392 | 389 | Klebsiella pneumoniae |
| ERR025562 | 229 | 389 | Klebsiella pneumoniae |
| ERR025562 | 392 | 389 | Klebsiella pneumoniae |
| ERR025563 | 229 | 389 | Klebsiella pneumoniae |
| ERR025563 | 392 | 389 | Klebsiella pneumoniae |
| ERR025564 | 229 | 389 | Klebsiella pneumoniae |
| ERR025564 | 392 | 389 | Klebsiella pneumoniae |
| ERR025566 | 229 | 389 | Klebsiella pneumoniae |
| ERR025566 | 392 | 389 | Klebsiella pneumoniae |
| ERR025566 | 436 | 389 | Klebsiella pneumoniae |
| ERR025570 | 229 | 389 | Klebsiella pneumoniae |
| ERR025570 | 392 | 389 | Klebsiella pneumoniae |
| ERR025572 | 229 | 389 | Klebsiella pneumoniae |
| ERR025572 | 392 | 389 | Klebsiella pneumoniae |
| ERR025574 | 229 | 389 | Klebsiella pneumoniae |
| ERR025574 | 392 | 389 | Klebsiella pneumoniae |
| ERR025575 | 229 | 389 | Klebsiella pneumoniae |
| ERR025575 | 392 | 389 | Klebsiella pneumoniae |
| ERR025576 | 229 | 389 | Klebsiella pneumoniae |
| ERR025576 | 392 | 389 | Klebsiella pneumoniae |
| ERR025579 | 392 | 389 | Klebsiella pneumoniae |
| ERR025581 | 229 | 389 | Klebsiella pneumoniae |
| ERR025581 | 392 | 389 | Klebsiella pneumoniae |
| ERR025582 | 229 | 389 | Klebsiella pneumoniae |
| ERR025582 | 392 | 389 | Klebsiella pneumoniae |
| ERR025583 | 229 | 389 | Klebsiella pneumoniae |
| ERR025583 | 392 | 389 | Klebsiella pneumoniae |
| ERR025585 | 392 | 389 | Klebsiella pneumoniae |
| ERR025587 | 229 | 389 | Klebsiella pneumoniae |
| ERR025587 | 392 | 389 | Klebsiella pneumoniae |
| ERR025588 | 229 | 389 | Klebsiella pneumoniae |
| ERR025588 | 392 | 389 | Klebsiella pneumoniae |
| ERR025589 | 392 | 389 | Klebsiella pneumoniae |
| ERR025592 | 229 | 389 | Klebsiella pneumoniae |
| ERR025592 | 392 | 389 | Klebsiella pneumoniae |
| ERR025593 | 392 | 389 | Klebsiella pneumoniae |
| ERR025594 | 229 | 389 | Klebsiella pneumoniae |
| ERR025594 | 392 | 389 | Klebsiella pneumoniae |
| ERR025595 | 392 | 389 | Klebsiella pneumoniae |
| ERR025596 | 229 | 389 | Klebsiella pneumoniae |
| ERR025596 | 392 | 389 | Klebsiella pneumoniae |
| ERR025597 | 229 | 389 | Klebsiella pneumoniae |
| ERR025597 | 392 | 389 | Klebsiella pneumoniae |
| ERR025599 | 229 | 64 | Klebsiella pneumoniae |
| ERR025599 | 229 | 389 | Klebsiella pneumoniae |
| ERR025599 | 392 | 64 | Klebsiella pneumoniae |
| ERR025599 | 392 | 389 | Klebsiella pneumoniae |
| ERR025600 | 229 | 64 | Klebsiella pneumoniae |
| ERR025600 | 229 | 389 | Klebsiella pneumoniae |
| ERR025600 | 392 | 64 | Klebsiella pneumoniae |
| ERR025600 | 392 | 389 | Klebsiella pneumoniae |
| ERR025601 | 229 | 389 | Klebsiella pneumoniae |
| ERR025601 | 392 | 389 | Klebsiella pneumoniae |
| ERR025602 | 392 | 389 | Klebsiella pneumoniae |
| ERR025603 | 392 | 389 | Klebsiella pneumoniae |
| ERR025605 | 229 | 389 | Klebsiella pneumoniae |
| ERR025605 | 392 | 389 | Klebsiella pneumoniae |
| ERR025606 | 229 | 389 | Klebsiella pneumoniae |
| ERR025606 | 392 | 389 | Klebsiella pneumoniae |
| ERR025607 | 229 | 389 | Klebsiella pneumoniae |
| ERR025607 | 392 | 389 | Klebsiella pneumoniae |
| ERR025608 | 229 | 389 | Klebsiella pneumoniae |
| ERR025608 | 392 | 389 | Klebsiella pneumoniae |
| ERR025609 | 229 | 389 | Klebsiella pneumoniae |
| ERR025609 | 392 | 389 | Klebsiella pneumoniae |
| ERR025610 | 392 | 389 | Klebsiella pneumoniae |
| ERR025612 | 175 | 244 | Klebsiella pneumoniae |
| ERR025612 | 229 | 244 | Klebsiella pneumoniae |
| ERR025613 | 229 | 389 | Klebsiella pneumoniae |
| ERR025613 | 392 | 389 | Klebsiella pneumoniae |
| ERR025614 | 392 | 389 | Klebsiella pneumoniae |
| ERR025615 | 229 | 389 | Klebsiella pneumoniae |
| ERR025615 | 392 | 389 | Klebsiella pneumoniae |
| ERR025616 | 229 | 389 | Klebsiella pneumoniae |
| ERR025616 | 392 | 389 | Klebsiella pneumoniae |
| ERR025618 | 392 | 389 | Klebsiella pneumoniae |
| ERR025619 | 229 | 389 | Klebsiella pneumoniae |
| ERR025619 | 392 | 389 | Klebsiella pneumoniae |
| ERR025620 | 229 | 389 | Klebsiella pneumoniae |
| ERR025620 | 392 | 389 | Klebsiella pneumoniae |
| ERR025621 | 229 | 389 | Klebsiella pneumoniae |
| ERR025621 | 392 | 389 | Klebsiella pneumoniae |
| ERR025622 | 229 | 389 | Klebsiella pneumoniae |
| ERR025622 | 392 | 389 | Klebsiella pneumoniae |
| ERR025624 | 392 | 389 | Klebsiella pneumoniae |
| ERR025625 | 392 | 389 | Klebsiella pneumoniae |
| ERR025626 | 229 | 389 | Klebsiella pneumoniae |
| ERR025626 | 392 | 389 | Klebsiella pneumoniae |
| ERR025627 | 229 | 389 | Klebsiella pneumoniae |
| ERR025627 | 392 | 389 | Klebsiella pneumoniae |
| ERR025628 | 229 | 389 | Klebsiella pneumoniae |
| ERR025628 | 392 | 389 | Klebsiella pneumoniae |
| ERR025629 | 392 | 389 | Klebsiella pneumoniae |
| ERR025633 | 392 | 389 | Klebsiella pneumoniae |
| ERR025635 | 229 | 389 | Klebsiella pneumoniae |
| ERR025635 | 392 | 389 | Klebsiella pneumoniae |
| ERR025638 | 229 | 389 | Klebsiella pneumoniae |
| ERR025638 | 392 | 389 | Klebsiella pneumoniae |
| ERR025641 | 229 | 389 | Klebsiella pneumoniae |
| ERR025641 | 392 | 389 | Klebsiella pneumoniae |
| ERR025645 | 392 | 389 | Klebsiella pneumoniae |
| ERR025646 | 392 | 389 | Klebsiella pneumoniae |
| ERR025647 | 229 | 366 | Klebsiella pneumoniae |
| ERR025647 | 392 | 366 | Klebsiella pneumoniae |
| ERR025648 | 229 | 389 | Klebsiella pneumoniae |
| ERR025648 | 392 | 389 | Klebsiella pneumoniae |
| ERR025649 | 392 | 389 | Klebsiella pneumoniae |
| ERR025652 | 392 | 389 | Klebsiella pneumoniae |
| ERR025653 | 392 | 389 | Klebsiella pneumoniae |
| ERR025654 | 229 | 389 | Klebsiella pneumoniae |
| ERR025654 | 392 | 389 | Klebsiella pneumoniae |
| ERR025655 | 392 | 389 | Klebsiella pneumoniae |
| ERR025657 | 229 | 389 | Klebsiella pneumoniae |
| ERR025657 | 392 | 389 | Klebsiella pneumoniae |
| ERR025658 | 229 | 389 | Klebsiella pneumoniae |
| ERR025658 | 392 | 389 | Klebsiella pneumoniae |
| ERR025659 | 229 | 389 | Klebsiella pneumoniae |
| ERR025659 | 392 | 389 | Klebsiella pneumoniae |
| ERR025660 | 392 | 389 | Klebsiella pneumoniae |
| ERR025661 | 229 | 389 | Klebsiella pneumoniae |
| ERR025661 | 392 | 389 | Klebsiella pneumoniae |
| ERR025662 | 229 | 389 | Klebsiella pneumoniae |
| ERR025662 | 392 | 389 | Klebsiella pneumoniae |
| ERR025663 | 229 | 389 | Klebsiella pneumoniae |
| ERR025663 | 392 | 389 | Klebsiella pneumoniae |
| ERR025664 | 229 | 389 | Klebsiella pneumoniae |
| ERR025664 | 392 | 389 | Klebsiella pneumoniae |
| ERR025665 | 229 | 389 | Klebsiella pneumoniae |
| ERR025665 | 392 | 389 | Klebsiella pneumoniae |
| ERR025666 | 229 | 389 | Klebsiella pneumoniae |
| ERR025666 | 392 | 389 | Klebsiella pneumoniae |
| ERR025667 | 229 | 389 | Klebsiella pneumoniae |
| ERR025667 | 392 | 389 | Klebsiella pneumoniae |
| ERR025668 | 229 | 389 | Klebsiella pneumoniae |
| ERR025668 | 392 | 389 | Klebsiella pneumoniae |
| ERR025670 | 392 | 389 | Klebsiella pneumoniae |
| ERR025673 | 392 | 389 | Klebsiella pneumoniae |
| ERR025674 | 229 | 389 | Klebsiella pneumoniae |
| ERR025674 | 392 | 389 | Klebsiella pneumoniae |
| ERR025675 | 229 | 389 | Klebsiella pneumoniae |
| ERR025675 | 392 | 389 | Klebsiella pneumoniae |
| ERR025676 | 392 | 389 | Klebsiella pneumoniae |
| ERR025677 | 392 | 389 | Klebsiella pneumoniae |
| ERR025678 | 392 | 389 | Klebsiella pneumoniae |
| ERR025679 | 229 | 389 | Klebsiella pneumoniae |
| ERR025679 | 392 | 389 | Klebsiella pneumoniae |
| ERR025680 | 229 | 389 | Klebsiella pneumoniae |
| ERR025680 | 392 | 389 | Klebsiella pneumoniae |
| ERR025979 | 392 | 389 | Klebsiella pneumoniae |
| ERR025980 | 229 | 389 | Klebsiella pneumoniae |
| ERR025980 | 392 | 389 | Klebsiella pneumoniae |
| ERR025981 | 229 | 389 | Klebsiella pneumoniae |
| ERR025981 | 392 | 389 | Klebsiella pneumoniae |
| ERR025983 | 229 | 389 | Klebsiella pneumoniae |
| ERR025983 | 392 | 389 | Klebsiella pneumoniae |
| ERR025984 | 392 | 389 | Klebsiella pneumoniae |
| ERR025985 | 229 | 389 | Klebsiella pneumoniae |
| ERR025985 | 392 | 389 | Klebsiella pneumoniae |
| ERR025988 | 229 | 389 | Klebsiella pneumoniae |
| ERR025988 | 392 | 389 | Klebsiella pneumoniae |
| ERR025989 | 229 | 389 | Klebsiella pneumoniae |
| ERR025989 | 392 | 389 | Klebsiella pneumoniae |
| ERR025989 | 436 | 389 | Klebsiella pneumoniae |
| ERR025990 | 229 | 64 | Klebsiella pneumoniae |
| ERR025990 | 229 | 389 | Klebsiella pneumoniae |
| ERR025990 | 392 | 64 | Klebsiella pneumoniae |
| ERR025990 | 392 | 389 | Klebsiella pneumoniae |
| ERR025992 | 229 | 389 | Klebsiella pneumoniae |
| ERR025992 | 392 | 389 | Klebsiella pneumoniae |
| ERR025993 | 392 | 389 | Klebsiella pneumoniae |
| ERR025994 | 229 | 389 | Klebsiella pneumoniae |
| ERR025994 | 392 | 389 | Klebsiella pneumoniae |
| ERR025995 | 229 | 389 | Klebsiella pneumoniae |
| ERR025995 | 392 | 389 | Klebsiella pneumoniae |
| ERR025996 | 229 | 389 | Klebsiella pneumoniae |
| ERR025996 | 392 | 389 | Klebsiella pneumoniae |
| ERR025997 | 229 | 389 | Klebsiella pneumoniae |
| ERR025997 | 392 | 389 | Klebsiella pneumoniae |
| ERR025998 | 229 | 389 | Klebsiella pneumoniae |
| ERR025998 | 392 | 389 | Klebsiella pneumoniae |
| ERR025999 | 392 | 389 | Klebsiella pneumoniae |
| ERR026000 | 392 | 389 | Klebsiella pneumoniae |
| ERR026001 | 392 | 389 | Klebsiella pneumoniae |
| GCF_000742135 | 229 | 389 | Klebsiella pneumoniae |
| GCF_000742135 | 392 | 389 | Klebsiella pneumoniae |
| GCF_900451275 | 229 | 389 | Klebsiella pneumoniae |
| GCF_900451275 | 392 | 389 | Klebsiella pneumoniae |
| Me103 | 392 | 389 | Klebsiella pneumoniae |
| Me106 | 229 | 389 | Klebsiella pneumoniae |
| Me106 | 392 | 389 | Klebsiella pneumoniae |
| Me107 | 229 | 389 | Klebsiella pneumoniae |
| Me107 | 392 | 389 | Klebsiella pneumoniae |
| Me19 | 229 | 389 | Klebsiella pneumoniae |
| Me19 | 392 | 389 | Klebsiella pneumoniae |
| Me33 | 229 | 389 | Klebsiella pneumoniae |
| Me5 | 229 | 389 | Klebsiella pneumoniae |
| Me5 | 392 | 389 | Klebsiella pneumoniae |
| Me74 | 229 | 389 | Klebsiella pneumoniae |
| Me74 | 392 | 389 | Klebsiella pneumoniae |
| Me78 | 392 | 389 | Klebsiella pneumoniae |
| Me81 | 229 | 389 | Klebsiella pneumoniae |
| Me81 | 392 | 389 | Klebsiella pneumoniae |
| Me96 | 229 | 389 | Klebsiella pneumoniae |
| Me96 | 392 | 389 | Klebsiella pneumoniae |
| PSKpn1 | 229 | 389 | Klebsiella pneumoniae |
| PSKpn1 | 392 | 389 | Klebsiella pneumoniae |
| PSKpn10 | N | 389 | Klebsiella pneumoniae |
| PSKpn11 | 229 | 389 | Klebsiella pneumoniae |
| PSKpn11 | 392 | 389 | Klebsiella pneumoniae |
| PSKpn12 | 229 | 389 | Klebsiella pneumoniae |
| PSKpn12 | 392 | 389 | Klebsiella pneumoniae |
| PSKpn13 | N | 389 | Klebsiella pneumoniae |
| PSKpn14 | N | 389 | Klebsiella pneumoniae |
| PSKpn15 | 229 | 389 | Klebsiella pneumoniae |
| PSKpn15 | 392 | 389 | Klebsiella pneumoniae |
| PSKpn16 | 229 | 389 | Klebsiella pneumoniae |
| PSKpn16 | 392 | 389 | Klebsiella pneumoniae |
| PSKpn2 | 229 | 389 | Klebsiella pneumoniae |
| PSKpn2 | 392 | 389 | Klebsiella pneumoniae |
| PSKpn24 | N | 389 | Klebsiella pneumoniae |
| PSKpn25 | N | 389 | Klebsiella pneumoniae |
| PSKpn26 | 229 | 389 | Klebsiella pneumoniae |
| PSKpn26 | 392 | 389 | Klebsiella pneumoniae |
| PSKpn27 | 229 | 389 | Klebsiella pneumoniae |
| PSKpn27 | 392 | 389 | Klebsiella pneumoniae |
| PSKpn28 | 392 | 389 | Klebsiella pneumoniae |
| PSKpn29 | 392 | 389 | Klebsiella pneumoniae |
| PSKpn3 | 229 | 389 | Klebsiella pneumoniae |
| PSKpn3 | 392 | 389 | Klebsiella pneumoniae |
| PSKpn30 | 392 | 389 | Klebsiella pneumoniae |
| PSKpn31 | 392 | 389 | Klebsiella pneumoniae |
| PSKpn32 | 229 | 389 | Klebsiella pneumoniae |
| PSKpn32 | 392 | 389 | Klebsiella pneumoniae |
| PSKpn32 | 436 | 389 | Klebsiella pneumoniae |
| PSKpn33 | N | 389 | Klebsiella pneumoniae |
| PSKpn35 | N | 389 | Klebsiella pneumoniae |
| PSKpn36 | 229 | 389 | Klebsiella pneumoniae |
| PSKpn36 | 392 | 389 | Klebsiella pneumoniae |
| PSKpn37 | 229 | 389 | Klebsiella pneumoniae |
| PSKpn37 | 392 | 389 | Klebsiella pneumoniae |
| PSKpn38 | 392 | 389 | Klebsiella pneumoniae |
| PSKpn39 | 392 | 389 | Klebsiella pneumoniae |
| PSKpn4 | 229 | 389 | Klebsiella pneumoniae |
| PSKpn4 | 392 | 389 | Klebsiella pneumoniae |
| PSKpn40 | 229 | 389 | Klebsiella pneumoniae |
| PSKpn40 | 392 | 389 | Klebsiella pneumoniae |
| PSKpn41 | 392 | 389 | Klebsiella pneumoniae |
| PSKpn7 | N | 389 | Klebsiella pneumoniae |
| PSKpn9 | 392 | 389 | Klebsiella pneumoniae |
| ERR024830 | 392 | 389 | Klebsiella quasipneumoniae |
| ERR024841 | 392 | 389 | Klebsiella quasipneumoniae |
| ERR025122 | 392 | 389 | Klebsiella quasipneumoniae |
| ERR025476 | 392 | 389 | Klebsiella quasipneumoniae |
| ERR025480 | 392 | 389 | Klebsiella quasipneumoniae |
| ERR025481 | 392 | 389 | Klebsiella quasipneumoniae |
| ERR025488 | 392 | 389 | Klebsiella quasipneumoniae |
| ERR025490 | 392 | 389 | Klebsiella quasipneumoniae |
| ERR025507 | 392 | 389 | Klebsiella quasipneumoniae |
| ERR025509 | 392 | 389 | Klebsiella quasipneumoniae |
| ERR025514 | 392 | 389 | Klebsiella quasipneumoniae |
| ERR025528 | 392 | 389 | Klebsiella quasipneumoniae |
| ERR025537 | 392 | 389 | Klebsiella quasipneumoniae |
| ERR025551 | 392 | 389 | Klebsiella quasipneumoniae |
| ERR025556 | 392 | 389 | Klebsiella quasipneumoniae |
| ERR025559 | 392 | 389 | Klebsiella quasipneumoniae |
| ERR025560 | 392 | 389 | Klebsiella quasipneumoniae |
| ERR025584 | 392 | 389 | Klebsiella quasipneumoniae |
| ERR025586 | 392 | 389 | Klebsiella quasipneumoniae |
| ERR025611 | 392 | 389 | Klebsiella quasipneumoniae |
| ERR025631 | 392 | 389 | Klebsiella quasipneumoniae |
| ERR025986 | 392 | 389 | Klebsiella quasipneumoniae |
| GCF_000751755 | 392 | 389 | Klebsiella quasipneumoniae |
| GCF_002269255 | N | 389 | Klebsiella quasivariicola |
| GCF_030343205 | 392 | N | Klebsiella spallanzanii |
| GCF_041414535 | 417 | N | Klebsiella spallanzanii |
| GCF_901563875 | 392 | N | Klebsiella spallanzanii |
| GCF_902158555 | 417 | N | Klebsiella spallanzanii |
| GCF_902158565 | 392 | N | Klebsiella spallanzanii |
| GCF_902158595 | 392 | 64 | Klebsiella spallanzanii |
| GCF_013705725 | 417 | N | Klebsiella taxon 3 |
| GCF_000829965 | 392 | 389 | Klebsiella terrigena |
| GCF_000964285 | 392 | 389 | Klebsiella terrigena |
| GCF_001975865 | 392 | 389 | Klebsiella terrigena |
| GCF_003752475 | 392 | 389 | Klebsiella terrigena |
| GCF_003752645 | 392 | 389 | Klebsiella terrigena |
| GCF_004362605 | 392 | 389 | Klebsiella terrigena |
| GCF_008080425 | 392 | 64 | Klebsiella terrigena |
| GCF_008080425 | 392 | 389 | Klebsiella terrigena |
| GCF_012029655 | 392 | 389 | Klebsiella terrigena |
| GCF_013393135 | 392 | 389 | Klebsiella terrigena |
| GCF_015571975 | 392 | 389 | Klebsiella terrigena |
| GCF_021440425 | 392 | 389 | Klebsiella terrigena |
| GCF_021698755 | 392 | 389 | Klebsiella terrigena |
| GCF_022605325 | 392 | 389 | Klebsiella terrigena |
| GCF_022627975 | 392 | 389 | Klebsiella terrigena |
| GCF_024809955 | 392 | 64 | Klebsiella terrigena |
| GCF_024809955 | 392 | 389 | Klebsiella terrigena |
| GCF_024810075 | 392 | 64 | Klebsiella terrigena |
| GCF_024810075 | 392 | 389 | Klebsiella terrigena |
| GCF_030078455 | 392 | 389 | Klebsiella terrigena |
| GCF_030389145 | 392 | 389 | Klebsiella terrigena |
| GCF_032086955 | 392 | 389 | Klebsiella terrigena |
| GCF_032086995 | 392 | 389 | Klebsiella terrigena |
| GCF_035793115 | 392 | 389 | Klebsiella terrigena |
| GCF_035797295 | 392 | 389 | Klebsiella terrigena |
| GCF_036561925 | 392 | 389 | Klebsiella terrigena |
| GCF_040838775 | 392 | 389 | Klebsiella terrigena |
| GCF_040838805 | 392 | 389 | Klebsiella terrigena |
| GCF_040838825 | 392 | 64 | Klebsiella terrigena |
| GCF_040838825 | 392 | 389 | Klebsiella terrigena |
| GCF_040838865 | 392 | 64 | Klebsiella terrigena |
| GCF_040838865 | 392 | 389 | Klebsiella terrigena |
| GCF_040838875 | 392 | 389 | Klebsiella terrigena |
| GCF_040838905 | 392 | 389 | Klebsiella terrigena |
| GCF_040838925 | 392 | 64 | Klebsiella terrigena |
| GCF_040838925 | 392 | 389 | Klebsiella terrigena |
| GCF_040838955 | 392 | 389 | Klebsiella terrigena |
| GCF_045290325 | 392 | 389 | Klebsiella terrigena |
| GCF_900476485 | 392 | 389 | Klebsiella terrigena |
| GCF_900636285 | 392 | 389 | Klebsiella terrigena |
| GCF_900706855 | 52 | 389 | Klebsiella terrigena |
| GCF_900706855 | 175 | 389 | Klebsiella terrigena |
| GCF_900706855 | 216 | 389 | Klebsiella terrigena |
| GCF_901420265 | 392 | 389 | Klebsiella terrigena |
| GCF_902109485 | 392 | 389 | Klebsiella terrigena |
| GCF_902705905 | 392 | 389 | Klebsiella terrigena |
| GCF_902705915 | 392 | 389 | Klebsiella terrigena |
| GCF_902705925 | 392 | 389 | Klebsiella terrigena |
| GCF_902705935 | 392 | 389 | Klebsiella terrigena |
| GCF_902705945 | 392 | 389 | Klebsiella terrigena |
| GCF_902705965 | 392 | 389 | Klebsiella terrigena |
| GCF_902705975 | 392 | 389 | Klebsiella terrigena |
| GCF_902705985 | 392 | 389 | Klebsiella terrigena |
| GCF_902705995 | 392 | 389 | Klebsiella terrigena |
| GCF_902706005 | 392 | 389 | Klebsiella terrigena |
| GCF_902706015 | 392 | 389 | Klebsiella terrigena |
| GCF_902706025 | 392 | 389 | Klebsiella terrigena |
| GCF_902706045 | 392 | 389 | Klebsiella terrigena |
| GCF_902706055 | 392 | 389 | Klebsiella terrigena |
| GCF_902706065 | 392 | 389 | Klebsiella terrigena |
| GCF_902706075 | 392 | 389 | Klebsiella terrigena |
| GCF_902706085 | 392 | 389 | Klebsiella terrigena |
| GCF_902706115 | 392 | 389 | Klebsiella terrigena |
| GCF_902706135 | 392 | 389 | Klebsiella terrigena |
| GCF_902706215 | 392 | 389 | Klebsiella terrigena |
| GCF_902706305 | 392 | 389 | Klebsiella terrigena |
| GCF_902706315 | 392 | 389 | Klebsiella terrigena |
| GCF_902706325 | 392 | 389 | Klebsiella terrigena |
| GCF_902706335 | 392 | 389 | Klebsiella terrigena |
| GCF_902706345 | 392 | 389 | Klebsiella terrigena |
| GCF_902706355 | 392 | 389 | Klebsiella terrigena |
| GCF_902706365 | 392 | 389 | Klebsiella terrigena |
| GCF_902706375 | 392 | 389 | Klebsiella terrigena |
| GCF_902706385 | 392 | 389 | Klebsiella terrigena |
| GCF_902706395 | 392 | 389 | Klebsiella terrigena |
| GCF_902706405 | 392 | 389 | Klebsiella terrigena |
| GCF_902706415 | 392 | 389 | Klebsiella terrigena |
| GCF_902706425 | 392 | 389 | Klebsiella terrigena |
| GCF_902706435 | 392 | 389 | Klebsiella terrigena |
| GCF_902706445 | 392 | 389 | Klebsiella terrigena |
| GCF_902706455 | 392 | 389 | Klebsiella terrigena |
| GCF_902706465 | 392 | 389 | Klebsiella terrigena |
| GCF_902706475 | 392 | 389 | Klebsiella terrigena |
| GCF_902706485 | 392 | 389 | Klebsiella terrigena |
| GCF_902706495 | 392 | 389 | Klebsiella terrigena |
| GCF_902706505 | 392 | 389 | Klebsiella terrigena |
| GCF_902706515 | 392 | 389 | Klebsiella terrigena |
| GCF_902706525 | 392 | 389 | Klebsiella terrigena |
| GCF_902706535 | 392 | 389 | Klebsiella terrigena |
| GCF_902706545 | 392 | 389 | Klebsiella terrigena |
| GCF_902706555 | 392 | 389 | Klebsiella terrigena |
| GCF_902706565 | 392 | 389 | Klebsiella terrigena |
| GCF_902706575 | 392 | 389 | Klebsiella terrigena |
| GFKo52 | 392 | 389 | Klebsiella terrigena |
| SAMD00008836 | 392 | 389 | Klebsiella terrigena |
| SAMEA2298771 | 392 | 389 | Klebsiella terrigena |
| SAMEA5048780 | 392 | 389 | Klebsiella terrigena |
| SAMEA5049071 | 392 | 389 | Klebsiella terrigena |
| SAMEA5049072 | 392 | 389 | Klebsiella terrigena |
| SAMEA5049073 | 392 | 389 | Klebsiella terrigena |
| SAMEA5049210 | 392 | 389 | Klebsiella terrigena |
| SAMEA5049211 | 392 | 389 | Klebsiella terrigena |
| SAMEA5049213 | 392 | 389 | Klebsiella terrigena |
| SAMEA5049214 | 392 | 389 | Klebsiella terrigena |
| SAMEA5049215 | 392 | 64 | Klebsiella terrigena |
| SAMEA5049215 | 392 | 389 | Klebsiella terrigena |
| SAMEA5049216 | 392 | 389 | Klebsiella terrigena |
| SAMEA5049366 | 392 | 389 | Klebsiella terrigena |
| SAMEA5049375 | 392 | 389 | Klebsiella terrigena |
| SAMEA5049547 | 392 | 389 | Klebsiella terrigena |
| SAMEA5049598 | 392 | 389 | Klebsiella terrigena |
| SAMEA5049599 | 392 | 389 | Klebsiella terrigena |
| SAMEA5049600 | 392 | 389 | Klebsiella terrigena |
| SAMEA5050198 | 392 | 389 | Klebsiella terrigena |
| SAMEA5050282 | 392 | 389 | Klebsiella terrigena |
| SAMEA5050291 | 392 | 389 | Klebsiella terrigena |
| SAMEA5050293 | 392 | 389 | Klebsiella terrigena |
| SAMEA5050295 | 392 | 389 | Klebsiella terrigena |
| SAMEA5684196 | 392 | 389 | Klebsiella terrigena |
| SAMEA5684272 | 392 | 389 | Klebsiella terrigena |
| SAMEA5684273 | 392 | 389 | Klebsiella terrigena |
| SAMEA6511050 | 392 | 389 | Klebsiella terrigena |
| SAMN10361115 | 392 | 389 | Klebsiella terrigena |
| ERR024826 | 392 | 389 | Klebsiella variicola |
| ERR024827 | 392 | 389 | Klebsiella variicola |
| ERR024834 | 392 | 389 | Klebsiella variicola |
| ERR024853 | 392 | 389 | Klebsiella variicola |
| ERR025102 | 392 | 389 | Klebsiella variicola |
| ERR025114 | 392 | 389 | Klebsiella variicola |
| ERR025120 | 392 | 389 | Klebsiella variicola |
| ERR025124 | 392 | 389 | Klebsiella variicola |
| ERR025134 | 392 | 389 | Klebsiella variicola |
| ERR025138 | 392 | 389 | Klebsiella variicola |
| ERR025145 | 392 | 389 | Klebsiella variicola |
| ERR025148 | 392 | 389 | Klebsiella variicola |
| ERR025153 | 392 | 389 | Klebsiella variicola |
| ERR025463 | 392 | 389 | Klebsiella variicola |
| ERR025549 | 392 | 389 | Klebsiella variicola |
| ERR025573 | 392 | 389 | Klebsiella variicola |
| ERR025598 | 392 | 96 | Klebsiella variicola |
| ERR025598 | 392 | 306 | Klebsiella variicola |
| ERR025671 | 392 | 389 | Klebsiella variicola |
| ERR025982 | 392 | 389 | Klebsiella variicola |
| GCF_000828055 | 392 | 389 | Klebsiella variicola |
| GCF_009758445 | 392 | 389 | Klebsiellal scottii |
| GCF_036561905 | 392 | 389 | Klebsiellal scottii |
| GCF_036561945 | 392 | 389 | Klebsiellal scottii |
| GCF_036562005 | N | 389 | Klebsiellal scottii |
| GCF_042055345 | 392 | 389 | Klebsiellal scottii |
| SAMEA5049058 | 392 | 389 | Klebsiellal scottii |
| SAMEA5049367 | 392 | 389 | Klebsiellal scottii |
| SAMEA5049368 | 392 | 389 | Klebsiellal scottii |
| SAMEA5049636 | 392 | 389 | Klebsiellal scottii |
| SAMEA5684275 | 392 | 389 | Klebsiellal scottii |
| SAMEA5684292 | 392 | 389 | Klebsiellal scottii |

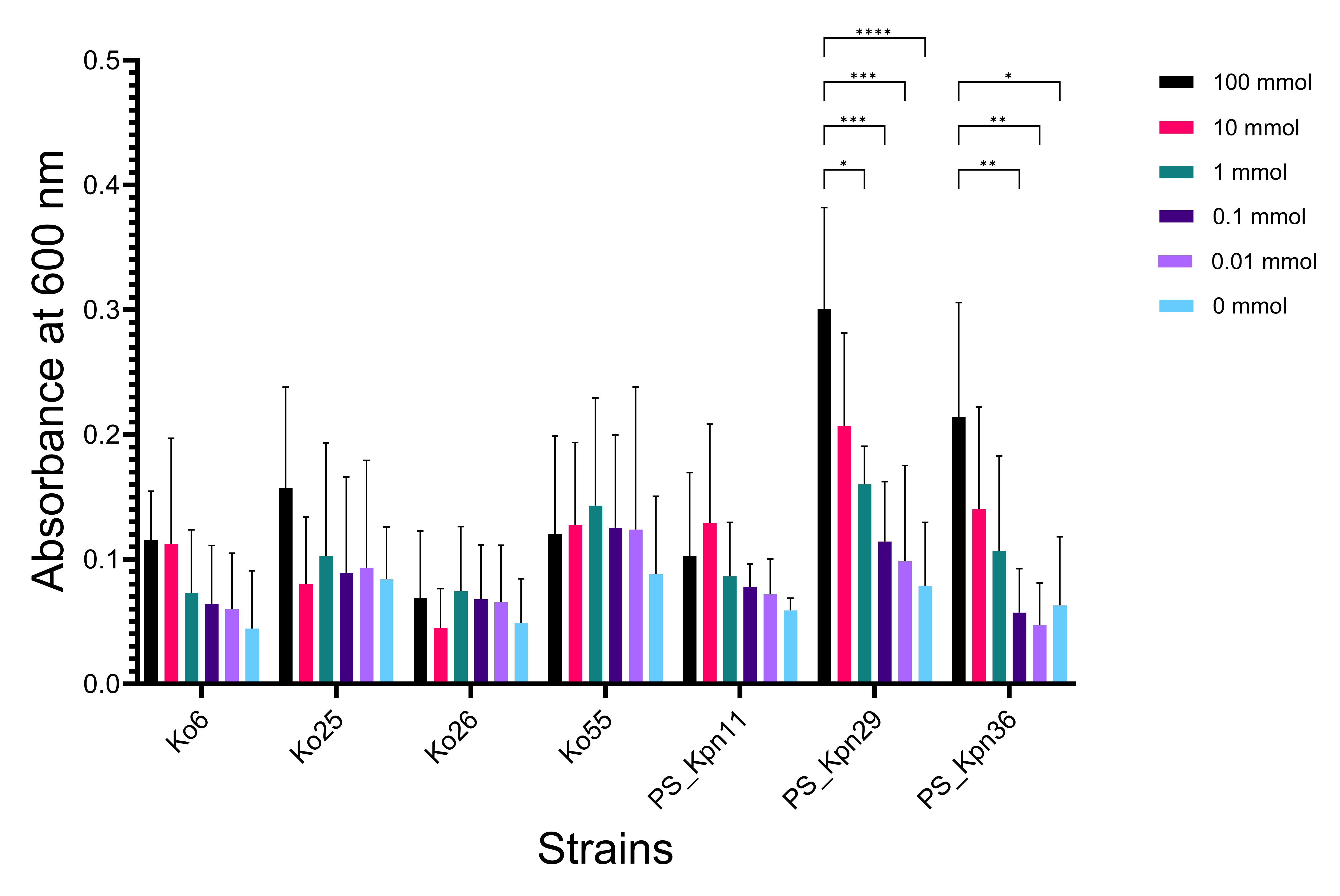

**Supplementary Figure 1.** Sodium hippurate does not inhibit growth of *Klebsiella* species. Substrate utilization assays were performed with four *Klebsiella* *oxytoca* (Ko6, Ko25, Ko26, Ko55) and three *Klebsiella pneumoniae* (PS_Kpn11, PS_Kpn29, and PS_Kpn36) strains isolated from urine. They were exposed to sodium hippurate (100, 10, 1, 0.1, 0.01 and 0 mmol). Growth was assessed by measuring optical density at 600 nm. Data are presented as mean + SD from four biological replicates (three technical replicates each). Statistical analysis was performed by two-way ANOVA with Tukey's post hoc test for multiple comparisons, comparing values across all concentration pairs within each strain, using GraphPad Prism (v10.4.2, GraphPad Software, Boston, MA, USA). Only statistically significant pairwise comparisons are indicated: *, P < 0.05; **, P < 0.01; ***, P < 0.001; ****, P < 0.0001.

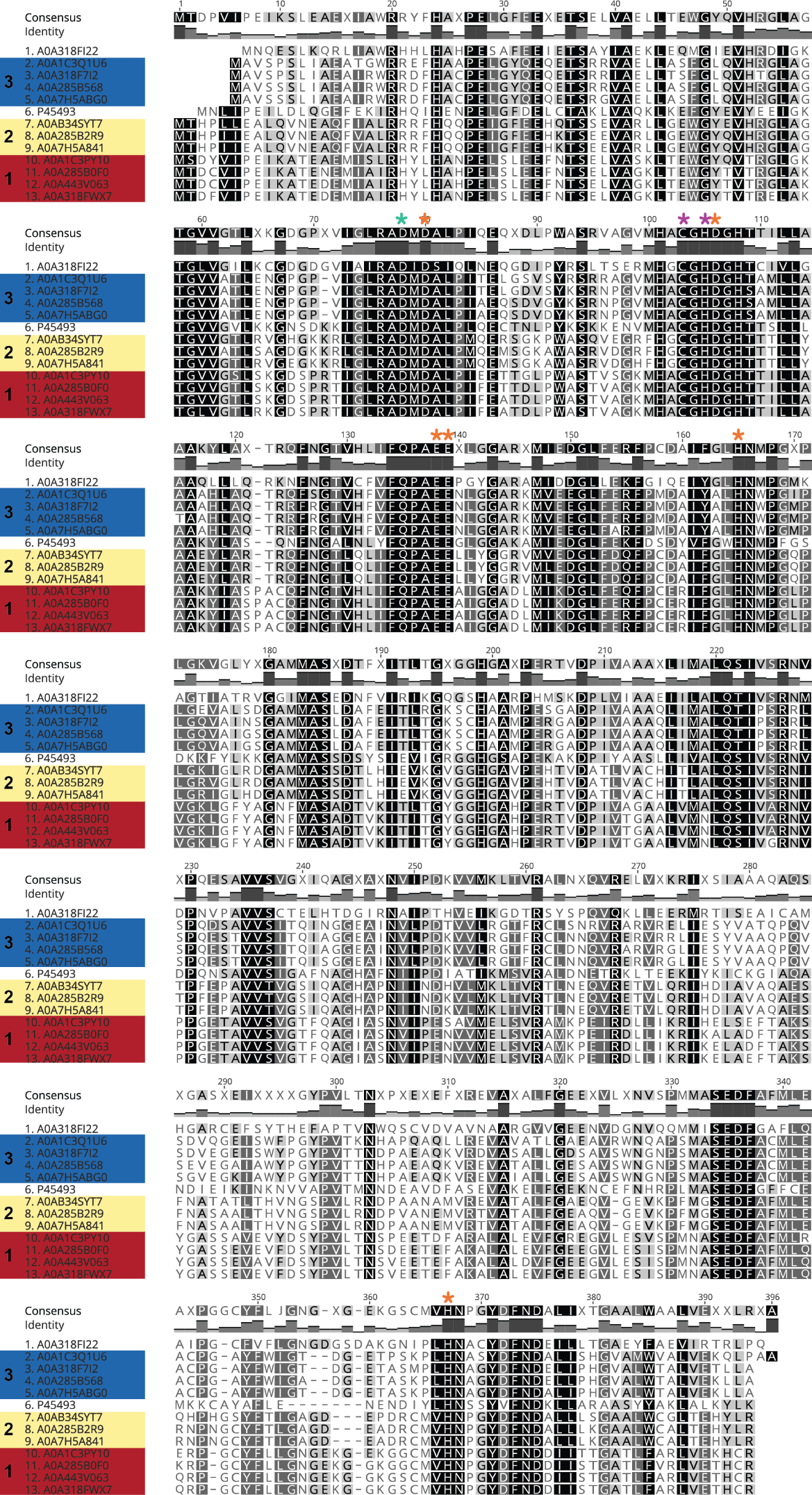

**Supplementary Figure 2.** Comparison of UniProt-identified hippurate hydrolase sequences. The zinc-containing metallocarboxypeptidase of *C. jejuni* (P45493) represents the well-characterized reference sequence. Sites (orange * above sequence) essential for the *C. jejuni* enzyme’s activity were previously identified using site-directed mutagenesis ^18^: complete loss of detectable hippurate hydrolytic activity for the individual Asp_104_, Glu_134_, Glu_135_, His_161_ and His_356_ mutants; the Asp_76_ mutant (green * above sequence) showed 72 % activity of the wild-type enzyme; all listed residues except for Glu_134_ are involved in binding zinc; Asp78, Glu_134_ and Glu_135_ are part of the enzyme active site ^18^. Predicted metal-binding sites for the *Klebsiella* sequences are shown by the purple *; in addition to these predicted sites, those corresponding to Glu_135_, His_161_ and His_356_ (*C. jejuni* numbering) are predicted to be metal-binding sites in *Klebsiella*. Hippurate hydrolase group affiliations for the *Klebsiella*-derived protein sequences are as shown in **Figure 1**.

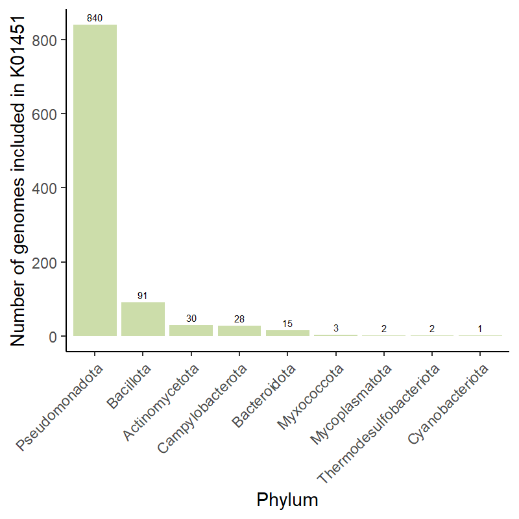

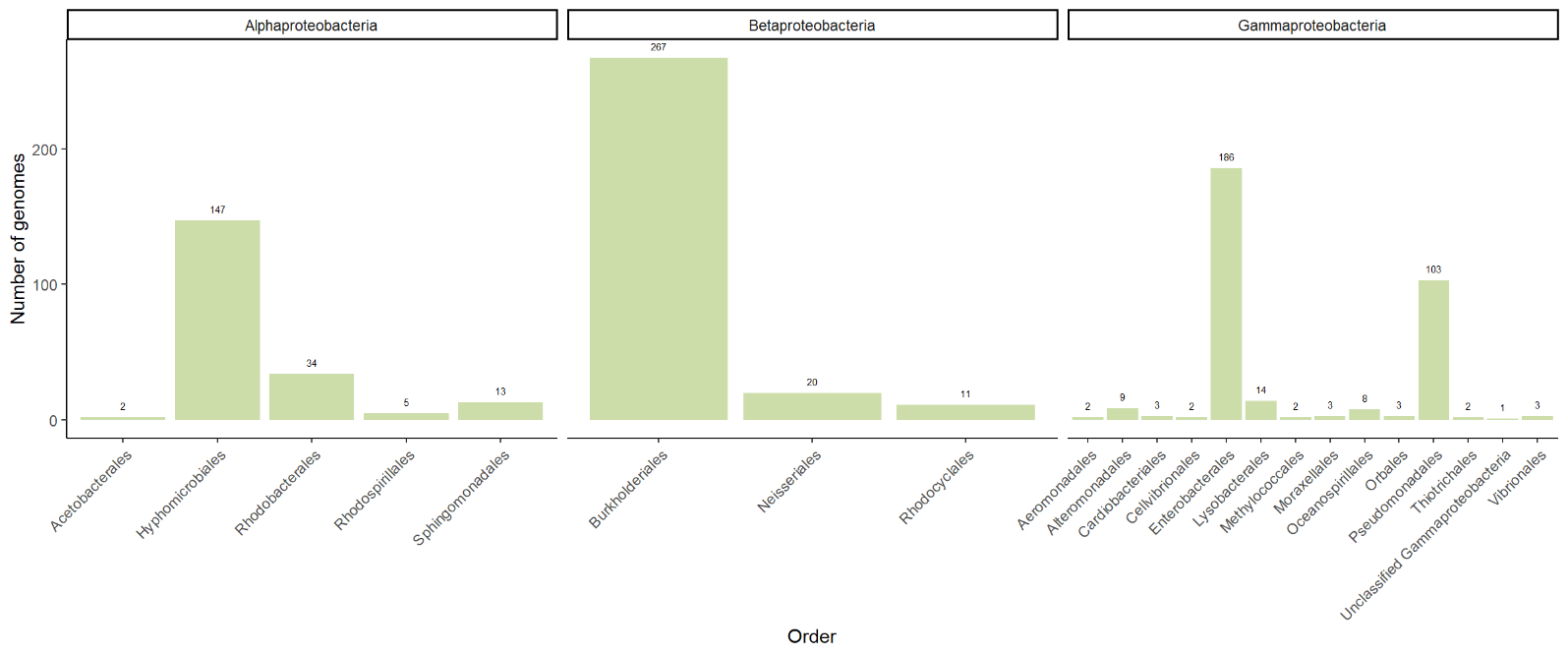

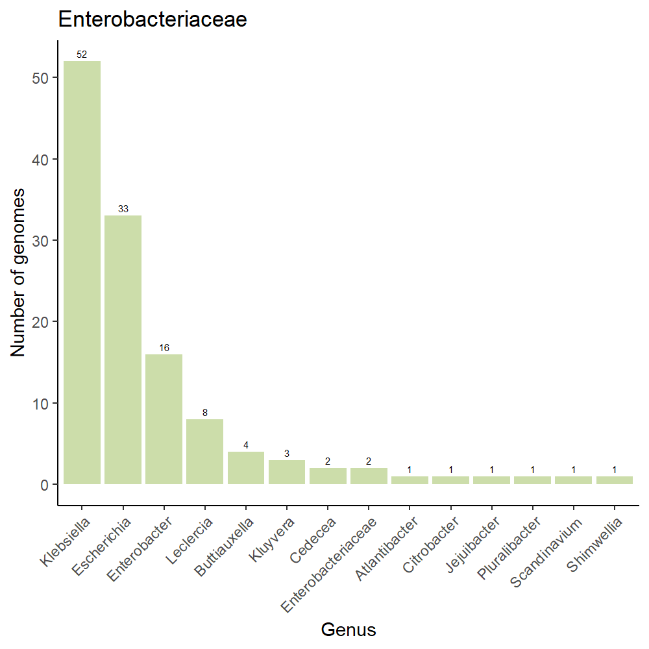

**Supplementary Figure 3.** Breakdown of bacterial taxa represented in KEGG orthology K01451 (hippurate hydrolase; KEGG) from phylum to genus level, with a focus on the phylum *Pseudomonadota*. The genus *Klebsiella* belongs to the family *Enterobacteriaceae* (order *Enterobacterales*). Only data for bacteria are presented here. The full list of organisms associated with K01451 at the time our analyses were undertaken can be found in **Supplementary Table 6**.

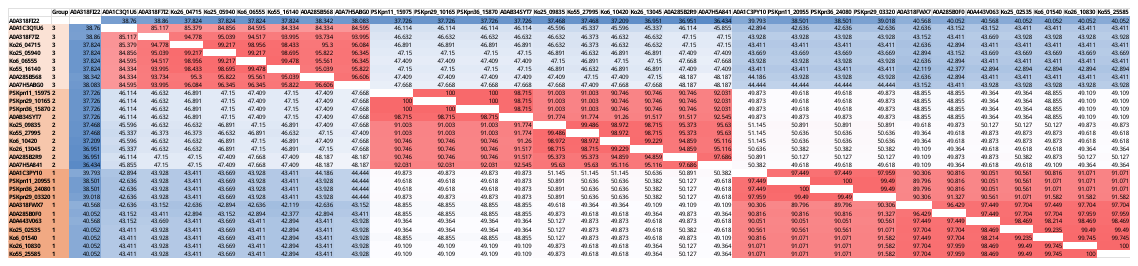

**Supplementary Figure 4.** Comparison of UniProt *Klebsiella* hippurate hydrolase protein sequences with those detected in the genomes of our seven in-house *Klebsiella* strains isolated from urine. Three clusters of proteins can be seen, with group numbering matching that shown in **Figure 2A**. PSKpn prefix, *K. pneumoniae*. Ko prefix, *K. oxytoca*. *K. quasipneumoniae* A0A1C3PY10, A0A1C3Q1U6 and A0AB34SYT7; *K. michiganensis* A0A7H5A841, A0A7H5ABG0 and A0A443V063; *K. grimontii* A0A285B0F0, A0A285B2R9 and A0A285B568; *K. oxytoca* A0A318FWX7, A0A318F7I2 and A0A318FI22. Values shown indicate amino acid identity (%), as determined from a multiple-sequence alignment (Clustal Omega) created in Geneious Prime.
